# Mechanical Properties of Native and Decellularized Reproductive Tissues: Insights for Tissue Engineering Strategies

**DOI:** 10.1101/2023.10.09.561536

**Authors:** R. Franko, Y. Franko, E. Ribes Martinez, G.A. Ferronato, I. Heinzelmann, N. Grechi, S. Devkota, P.K. Fontes, R. Coeti, T.S.I. Oshiro, M.A.M.M. Ferraz

## Abstract

Understanding the mechanical properties and porosity of reproductive tissues is vital for regenerative medicine in tissue engineering. This study investigated the changes in Young’s modulus (YM), storage modulus (E′), loss modulus (E″), and porosity of native and decellularized bovine reproductive tissues during the estrous cycle. Testis tunica albuginea had significantly higher YM, E′, and E″ than the inner testis, indicating greater stiffness and viscoelasticity. Endometrium showed no distinct differences in YM, E′, or E′ across the estrous cycle or between horns. Ovaries exhibited significant variations in YM, E′, E″, and porosity, with higher YM and E′ in the ipsilateral cortex and medulla during the luteal phase. Decellularized ovarian tissues displayed increased porosity. The oviduct displayed no significant differences in YM or E′ in the isthmus, but the contralateral ampulla had reduced YM and E′ in the luteal phase. These findings offer valuable insights into the dynamic mechanical properties and porosity of reproductive tissues, facilitating the development of biomimetic scaffolds for tissue engineering applications.

## Introduction

Tissue engineering, a multidisciplinary area combining principles of biology, chemistry, and engineering, has been revolutionizing the regenerative medicine field over the past few decades ^1^. One specific aspect of tissue engineering that has garnered significant attention is the engineering of reproductive tissues. Reproductive tissues play a crucial role in the biology of reproduction, serving as the environment for germ cell maturation, fertilization, and embryo development. Engineered male and female biomimetic reproductive tissues are being developed as autonomous *in vitro* units^2–6^ or as integrated multi-organ *in vitro* systems to support germ cell and embryo function and to display characteristic endocrine phenotypic patterns, such as the 28-day human menstrual cycle^7^. These engineered tissues are facilitating research in reproductive biology and may eventually allow the restoration of reproductive capacity in patients^8^.

Among the key properties to consider in tissue engineering, tissue stiffness, elasticity, viscoelasticity, and porosity are of fundamental importance^9,10^. The mechanical properties of biological tissues, from nanoscale to macroscale dimensions, have been shown to be critical for cellular behavior and tissue functionality^9,11,12^. Approaches to engineering biological tissues must, therefore, integrate and approximate the mechanics, both static and dynamic, of native tissues. Porosity also plays a significant role in enabling the manufacture of complex three-dimensional structures, recreating mechanical properties close to those of the host tissues, facilitating interconnected structures for the transport of macromolecules and cells, and behaving as biocompatible inserts^10^. The porosity of biomaterials has a direct impact on their behavior and performance, posing both advantages and challenges for each class of materials^10^.

The reproductive tract is a dynamic environment, characterized by constant change in response to hormonal cues. These changes are particularly notable in female reproductive tissues, which undergo cyclical modifications throughout the menstrual/estrous cycle^13,14^. Hormones like follicle-stimulating hormone (FSH) and luteinizing hormone (LH), produced by the pituitary gland, stimulate the development and ovulation of oocytes and also induce the production of estradiol and progesterone by the ovaries. These steroid hormones prepare the body for pregnancy, with estradiol also producing secondary sex characteristics in females^15^. For example, progesterone and estradiol play a crucial role in regulating the menstrual cycle, during which the uterus undergoes phases of proliferation, secretion, and menstruation, in response to these hormonal changes^13,14^.

In addition to their role in reproductive function, hormones can also influence the mechanical properties of reproductive tissues. For instance, there is evidence suggesting that female sex hormones may affect the structure and mechanical properties of the musculoskeletal system, potentially altering injury risk during certain phases of the menstrual cycle or during pregnancy^16^. From a tissue engineering perspective, it is essential to consider these dynamic mechanical properties while designing biomimetic reproductive tissues. The mechanical properties of biological tissues are fundamental for cellular behavior and consequent tissue functionality, and this understanding has greatly advanced the field of tissue engineering and regenerative medicine^9,17,18^. However, designing reproductive tissues that can mimic the complex and demanding mechanical environments *in vivo* remains a challenge, mainly due to a poor understanding of such properties in native tissues.

In this study, we investigated the mechanical properties and porosity of both native and decellularized reproductive tissues. The aim was to understand how these properties, which are critical for tissue function, change in response to hormonal fluctuations such as those in the estrous cycles. By comparing these properties between native and decellularized tissues, we hope to gain insights that will inform the creation of more accurate *in vitro* models for reproductive tissue engineering, potentially advancing regenerative medicine and the restoration of reproductive capacity in patients.

## Results

### Elastic and viscoelastic properties of reproductive tissues are location, tissue, and estrous-cycle dependent

In the biomechanical analysis of the testis, there were clear differences between the *tunica albuginea* and the inner testicular tissue in terms of their storage modulus (E′) and loss modulus (E″). Both E′ and E″ were found to be significantly higher in the *tunica albuginea* compared to the inner testis (p<0.001 for both E′ and E″), indicating that the *tunica albuginea* is both more elastic and more viscoelastic than the inner testicular tissue. Furthermore, the difference between the storage and loss moduli (E′-E″) was found to be greater in the *tunica albuginea* than in the inner testis, suggesting a relative increase in elastic response in the *tunica albuginea* compared to the inner testicular tissue (Fig. 1a).

**Figure 1.**
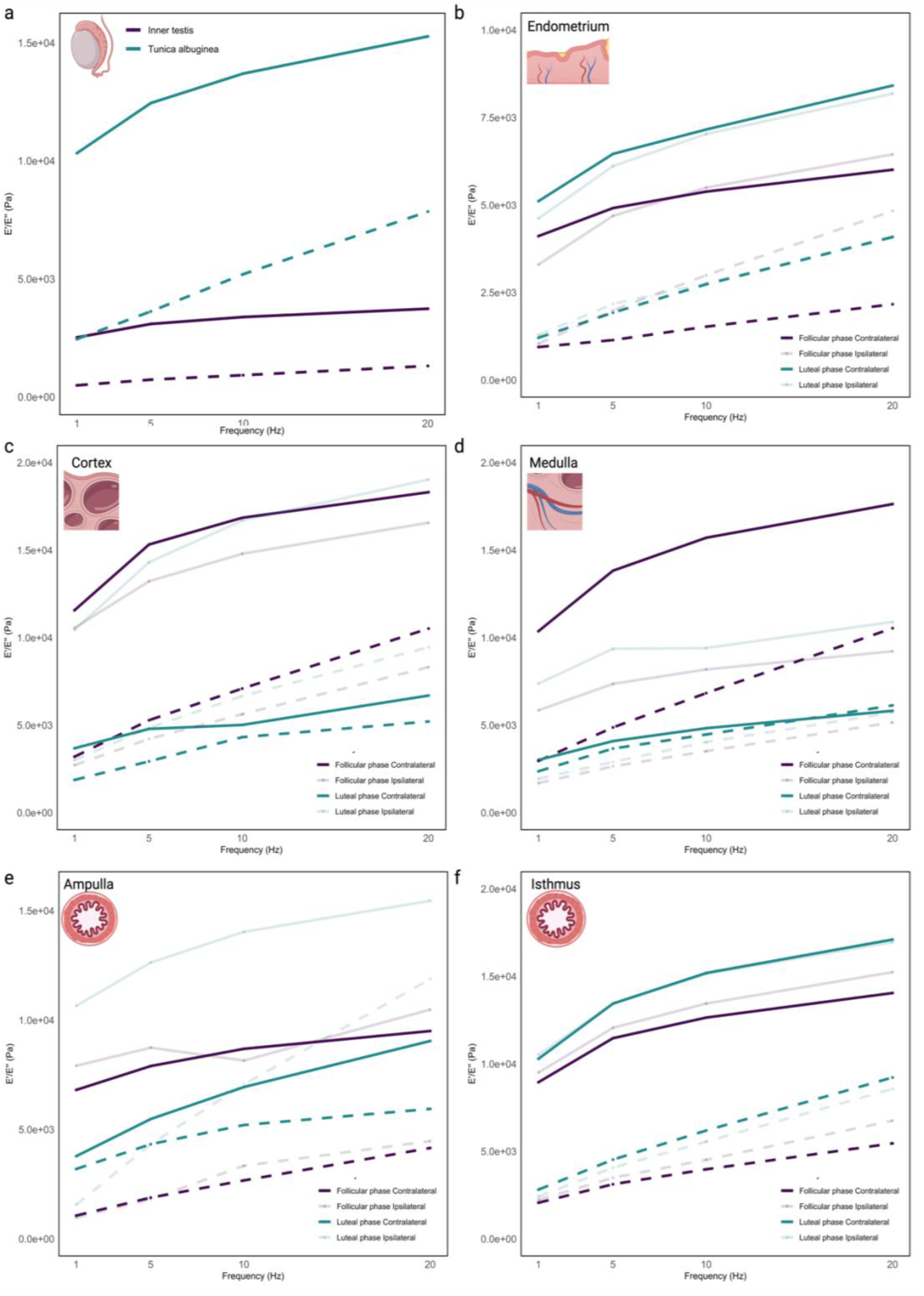
Elastic and viscoelastic properties of reproductive tissues. Storage (E′, continuous lines) and Loss (E″, dashed lines) modulus of native inner testis and *tunica albuginea* (**a**), endometrium (**b**), ovarian cortex (**c**), ovarian medulla (**d**), oviductal ampulla (**e**), and oviductal isthmus (**f**). Female tissues were analyzed for both follicular and luteal phases and ipsi- and contralateral positions.

In the biomechanical analysis of the endometrial tissue, non-distinct differences were observed across the luteal and follicular phases and between the ipsi- and contralateral horns for E′ and E″ individually. Nevertheless, a decrease in the difference between the storage modulus and loss modulus (E′-E″) in the ipsilateral horn during the follicular phase, compared to the other phases and locations, was observed (Fig. 1b).

The biomechanical properties of the ovarian cortex (Fig. 1c) and medulla (Fig. 1d) were also analyzed, with several significant findings observed. In the luteal phase cortex, the storage modulus from the ipsilateral horn was significantly higher than the contralateral horn (p<0.001). Same observation was found in the luteal phase medulla for both E′ (p<0.01) and E″ (p=0.0399). The difference E′-E″ in the luteal contralateral cortex and medulla were reduced when compared to their respective luteal ipsilateral tissues. This decrease in the difference between E′ and E″ might suggest an increased viscoelasticity of the contralateral ovarian tissues during the luteal phase. Regarding the oviduct, no significant difference on E′ and E″ when comparing estrous stage and position were observed in the isthmus (Fig. 1f). Nevertheless, there was a reduction in storage modulus (E′) of the contralateral ampulla when compared to the ipsilateral ampulla in the luteal phase (p<0.001, Fig. 1e). Moreover, a reduction in the E′-E″ was observed in the luteal ipsilateral ampulla (specially under higher frequencies) when compared to the follicular phase, suggesting an increased viscoelasticity of the ipsilateral ampulla during the luteal phase (Fig. 1e).

### Validation of decellularization method

Following the implementation of a sequential decellularization protocol involving sodium deoxycholate and DNase I, the effectiveness of the employed method was validated through the application of Hoechst 33342 nuclear staining to evaluate the decellularized and native tissues (Fig. 2a illustrates the comprehensive results of the decellularization process). To quantify the depletion of DNA in the samples subsequent to decellularization, DNA extraction was performed prior to and after the procedure. Analysis of the obtained data exhibited a remarkable DNA removal of 97.0, 97.6, 96.6, and 96.3% for endometrium, ovary, oviduct, and testis, respectively, during the decellularization process (Fig. 2b).

**Figure 2.**
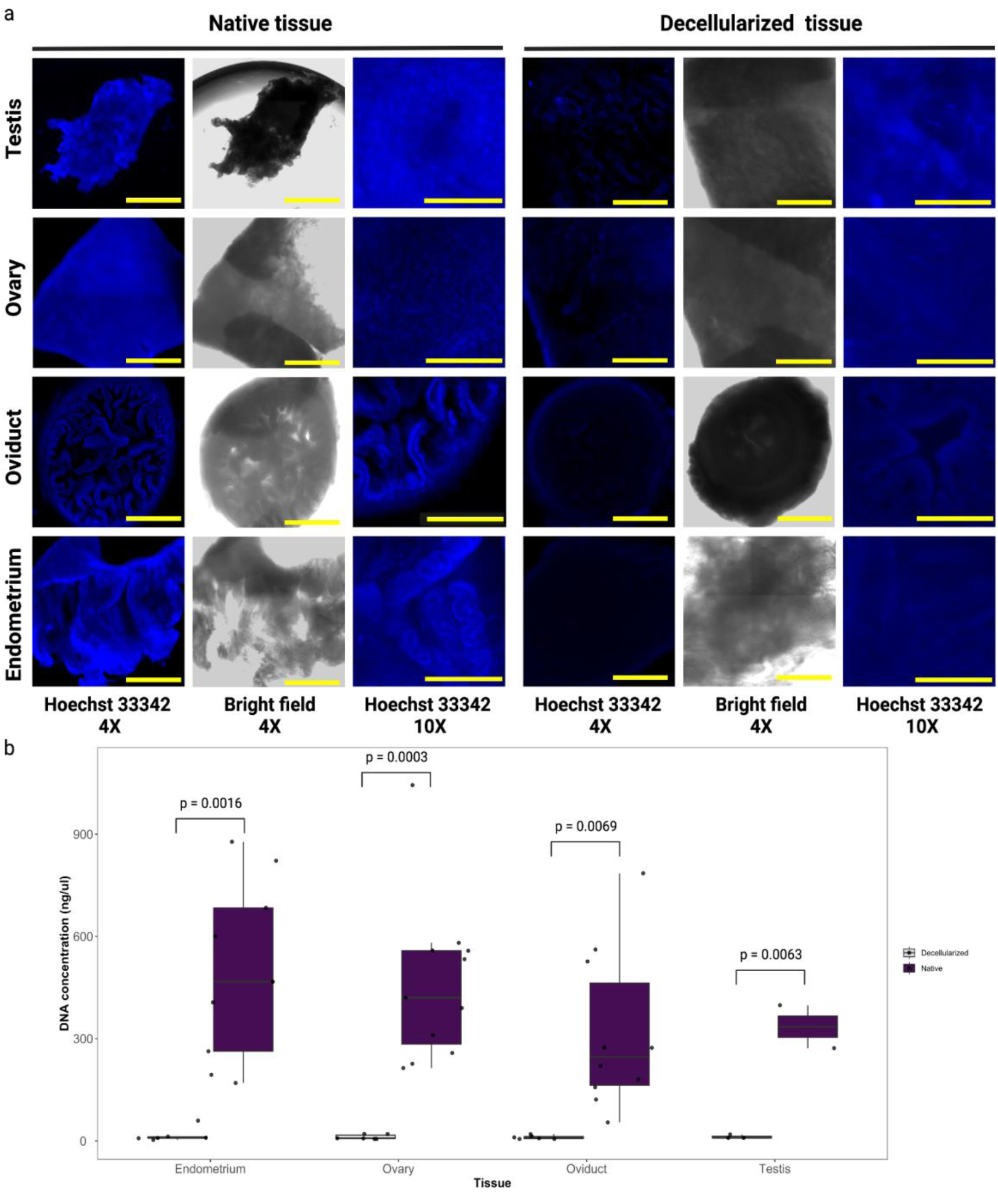
Validation of decellularization protocol. Representative images of nuclear staining (Hoechst 33342) of native and decellularized tissues are shown (**a**). Decellularization efficiency was also validated by DNA (**b**) quantification of native and decellularized tissues. Scale bar for 4x images = 2,000 µm. Scale bar for 10x images = 500 µm.

### Young’s modulus and porosity of testicles are affected by tissue decellularization

Nanoindentation was conducted on pieces of the *tunica albuginea* and inner testis to evaluate the Young’s modulus of both native and decellularized tissues. Each tissue fragment underwent multiple random indentations as seen in Fig. 3a. The native inner testis exhibited a reduced mean Young’s modulus (YM) of 1.39±0.44 kPa when compared to the decellularized tissue (mean YM of 12.83±7.70 kPa, p<0.001; Fig. 3b). Differently from the inner testis, no significant changes on Young’s modulus were observed after decellularization for the *tunica albuginea* (5.88±1.52 *vs* 12.71±9.83 kPa, native and decellularized, respectively, p=0.1737; Fig. 3b). Additionally, the native inner testis had a reduced mean Young’s modulus compared to the native *tunica albuginea* (1.39±0.44 *vs* 5.88±1.52 kPa, respectively; p=0.0220). Nevertheless, after decellularization, a lack of difference between inner testis and *tunica albuginea* YM was observed (p=1.00).

**Figure 3.**
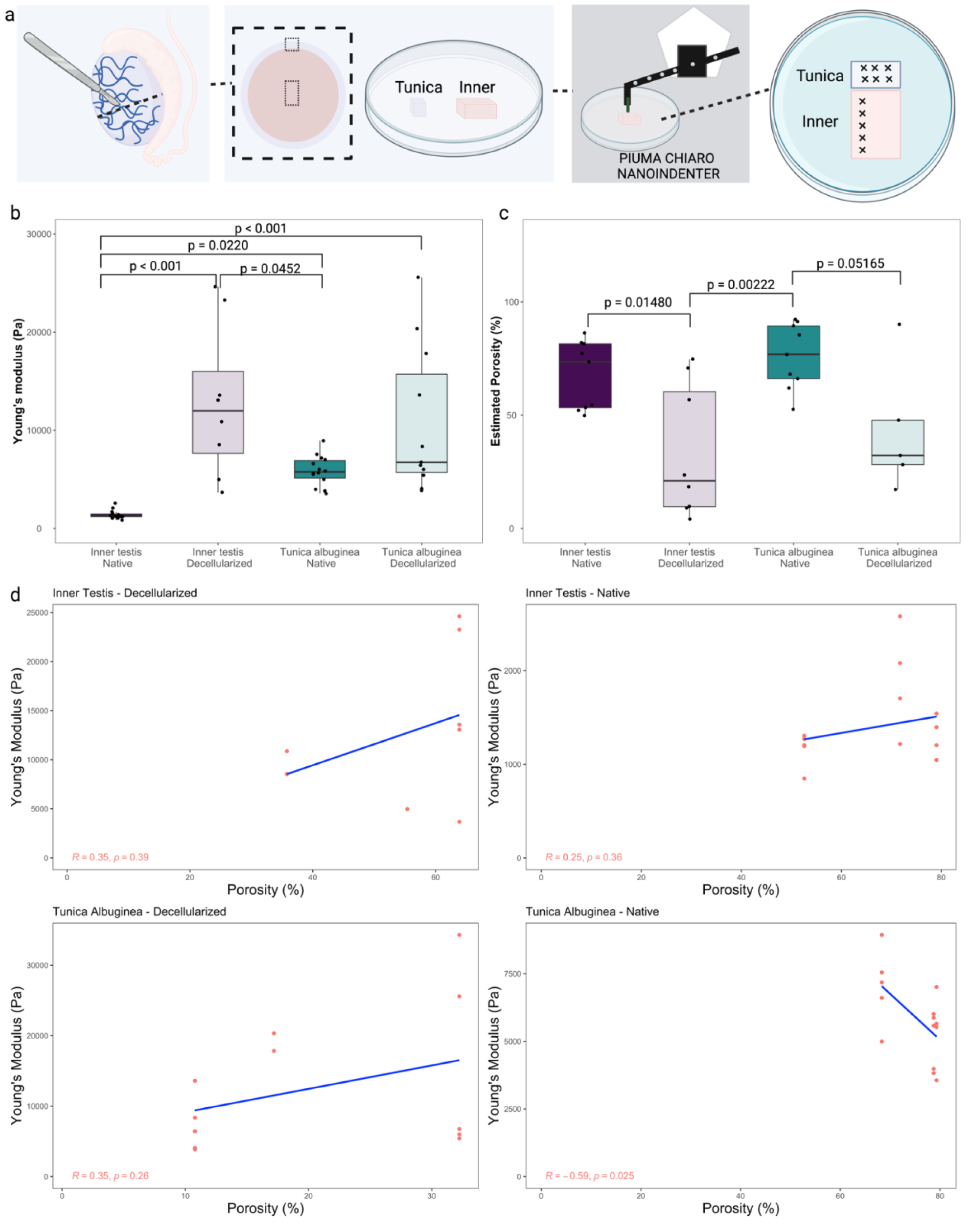
Young’s modulus and porosity analysis of native and decellularized testicles. In (**a**) details of sampling for nanoindentation and porosity analysis, testes were cut transversely in half, the *tunica vaginalis* was removed to allow the isolation of the *tunica albuginea* and the inner part of the testis, the samples were mounted on a petri dish using 8 mg mL^-1^ agarose solution. In (**b**) Young’s modulus of testis (n = 3 bulls). In (**c**) porosity data (n = 3 bulls). In (**d**) correlation plots of Porosity and Young’s modulus.

The estimated porosity of the native inner testis was higher than the decellularized inner testis (67.5±15.0 *vs* 33.4±29.3%, respectively, p=0.01480; Fig. 3c). Similarly, the decellularization process reduced the porosity of the native *tunica albuginea* (76.0±14.5% *vs* 43.1±28.5%, respectively, p=0.05165; Fig. 3c). Contrary to the YM data, no differences on porosity were observed when comparing inner testis and *tunica albuginea*, for both native and decellularized tissues (p=0.8543 and p=0.8617, respectively). To explore the relationship between porosity and Young’s modulus, a correlation study was conducted. A weak negative correlation between porosity and Young’s modulus was observed only in the native *tunica albuginea* (R=-0.59 and p=0.025, Fig. 3d), while no significant correlations were detected for the other groups.

In summary, the results showed that the decellularized inner testis had a significantly higher YM compared to the native tissue, while no significant changes in YM were observed in the decellularized *tunica albuginea*. The porosity was higher in the native compared to the decellularized tissues, for both inner testis and *tunica albuginea*.

### Young’s modulus and porosity of female reproductive tissues are estrous cycle dependent and affected by tissue decellularization

Three female reproductive tissues were analyzed from the same animal (endometrium, ovary, and oviduct). In total, six cows were studied, three at follicular phase and three at luteal phase of the estrous cycle. Additionally, samples were analyzed considering the position: ipsi- and contralateral, to the dominant follicle in follicular phase animals or the corpus luteum in luteal phase 2 animals.

#### Young’s modulus and porosity of endometrium

Tissue sections from contra- and ipsilateral horns of the endometrium in the luteal and follicular phases of the estrous cycle were evaluated (Fig. 4a). An increase in the YM was observed in decellularized compared to native tissues only at luteal phase (Fig. 4b), in both contralateral (9.24±5.22 *vs* 2.75±3.6 kPa, p=0.00837) and ipsilateral horns (16.25±9.69 *vs* 2.51±1.55 kPa, p<0.0001), but not at follicular phase, neither in contralateral (7.51±6.40 *vs* 2.75±3.6 kPa, decellularized and native, respectively, p>0.05) nor ipsilateral (4.22±1.65 *vs* 2.13±1.62, decellularized and native, respectively, p>0.05). An increase in YM in the luteal compared to the follicular phase was observed for both native and decellularized ipsilateral horns (p=0.0208 and p<0.0001, respectively), while no significant differences were observed for the contralateral horns. Nevertheless, no differences on YM were observed between native contralateral and ipsilateral horns (for a full comparison of groups see Suppl. Data 1).

**Figure 4.**
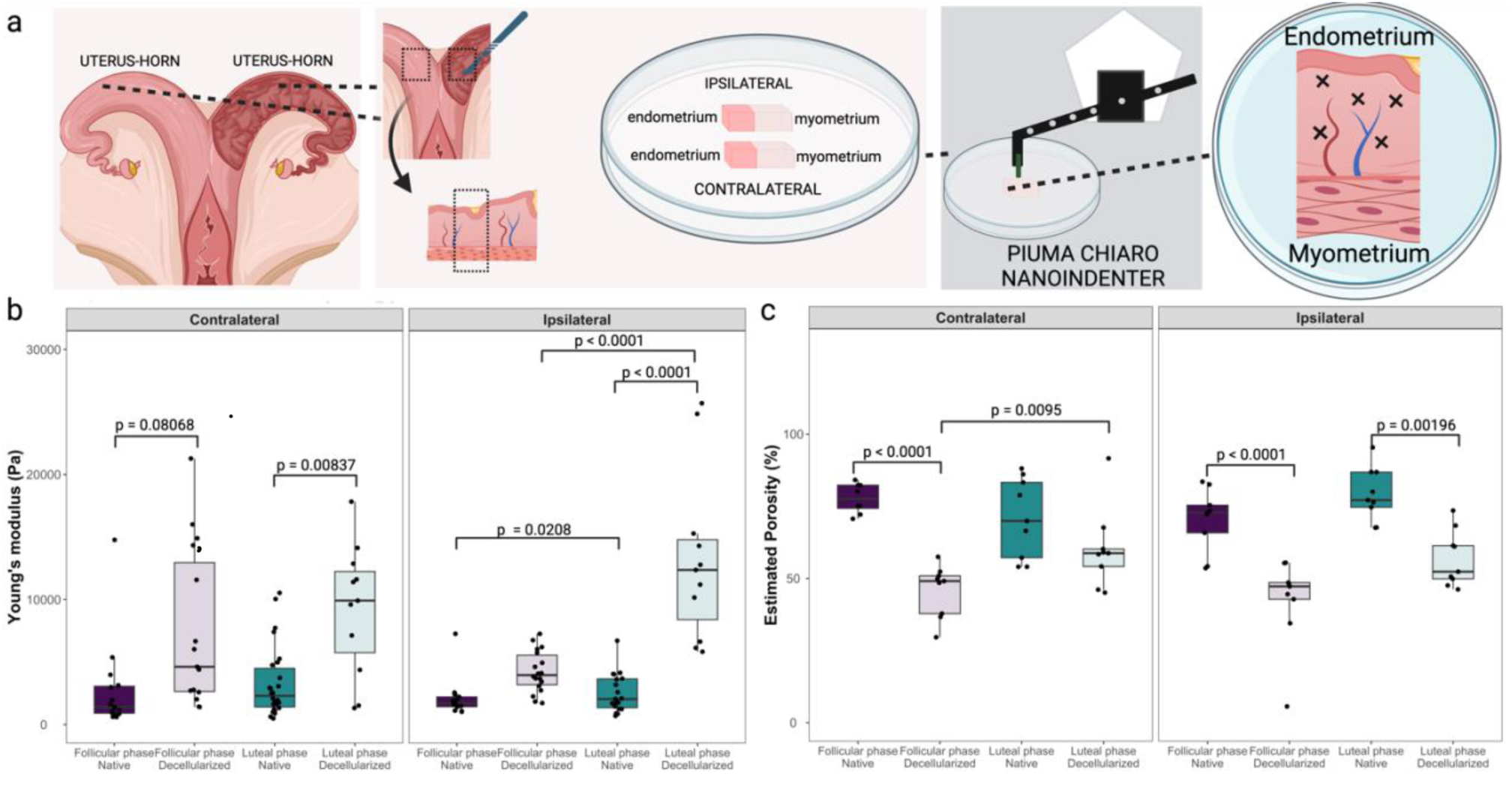
Young’s modulus and porosity analysis of native and decellularized endometrium. In (**a**) details of sampling for nanoindentation and porosity analysis, uteri were opened and a transversal sample was collected of each horn (ipsi- and contralateral), the endometrium was collected with the myometrium to stabilize the positioning of the tissue (myometrium was not evaluated) and the sample was mounted on a petri dish using 8 mg mL^-1^ agarose solution. In (**b**) Young’s modulus of endometrial tissue from native and decellularized tissue (n = 3 cows in follicular phase; n = 3 cows in luteal phase). In (**c**) porosity data from endometrial tissue from native and decellularized tissue (n = 3 cows in follicular phase; n = 3 cows in luteal phase).

There was a reduction of estimated porosity values for contralateral native tissues compared to the decellularized tissues in the follicular phase (77.7±5.1 vs 45.8±9.0%, p<0.0001), which was not significant in the luteal phase (70.9±13.7 vs 60.1±13.7%, p=0.28177, Fig. 4c). Same reduction was observed in the ipsilateral samples between native and decellularized tissues during follicular (70.4±10.8 vs 42.5±15.2%, p<0.0001) and luteal (79.2±9.2 vs 56.8±9.7%, p=0.00196) phases. No significant differences were observed when contralateral and ipsilateral horns were compared (for a full comparison of groups see Suppl. Data 1). Similarly to the testis, a correlation study between YM and porosity was performed, weak negative correlations for contralateral decellularized luteal and ipsilateral native luteal endometrium were observed (R=-0.59 and p=0.02; R=-0.1 and p=0.03, respectively; Suppl. Fig. 1), while all other groups had no significant correlations.

In summary, the native endometrium of the ipsilateral horn showed lower YM in the follicular phase compared to the luteal phase. After decellularization, the stiffness increased only in horns collected at luteal phase, in both ipsi- and contralateral horns. The porosity of the decellularized tissues was lower than that of the native tissues in follicular-contralateral and both phases in ipsilateral horns.

#### Young’s modulus and porosity of ovary

Ipsi- and contralateral ovaries from luteal and follicular phases of the estrous cycle were evaluated. For nanoindentation analysis, thin slices were cut transversely to analyze all layers of the ovary. Multiple indentations were performed with a 150 μm interval in the x-axis starting with a distance of 1,500 μm (600 μm estimated cortex and tunica layers and 900 μm of medulla layer) to accurately measure all layers of the ovary (*Tunica albuginea*, cortex, and medulla, as shown in Fig. 5a). Because a non-precise distinction between *Tunica albuginea* and cortex was possible, data was separated and presented as cortex and medulla. For porosity measurements, the cortex and the medulla were separately dissected as described in the methods.

**Figure 5.**
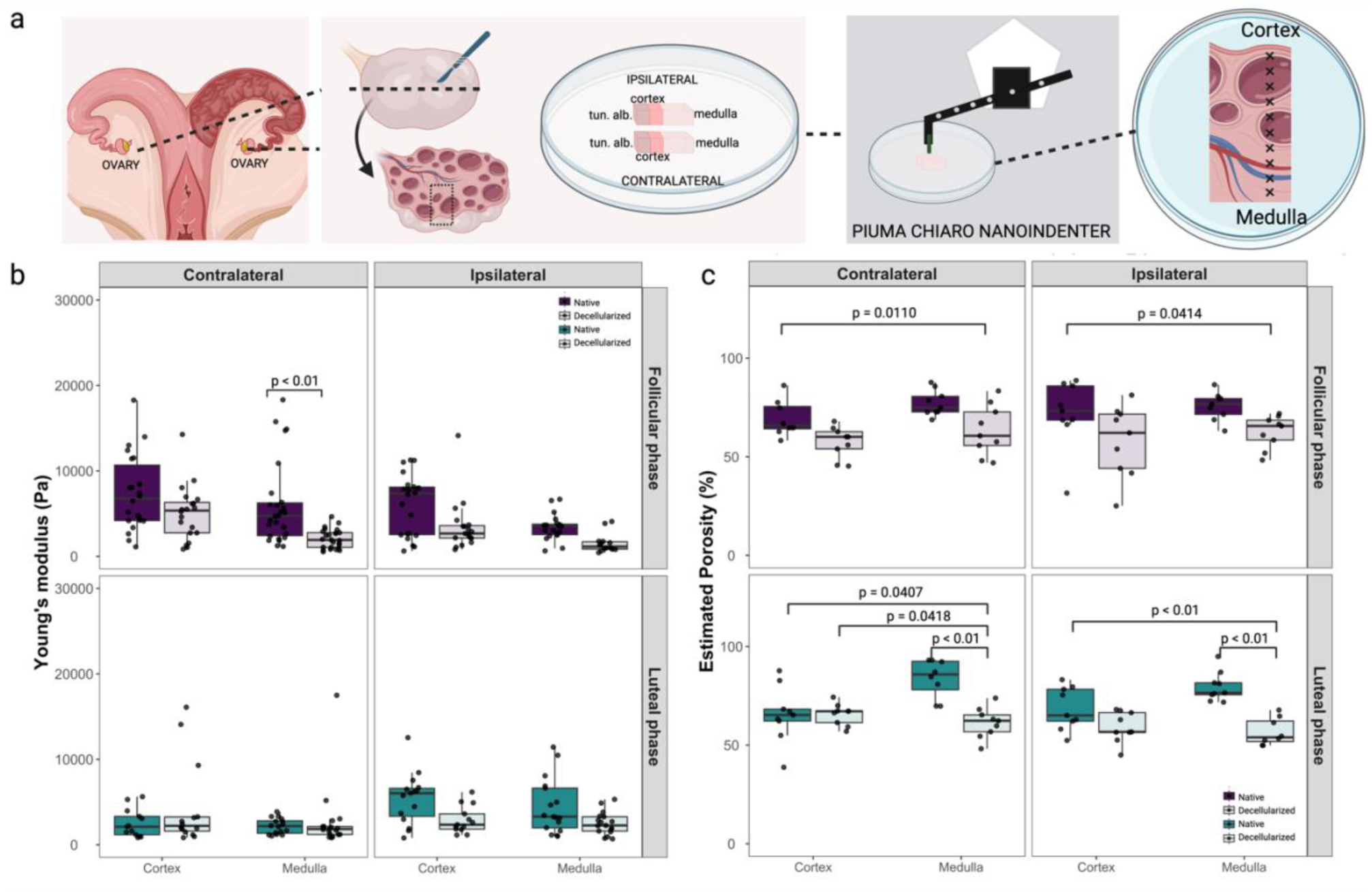
Young’s modulus and porosity analysis of native and decellularized ovaries. In (**a**) details of sampling for nanoindentation, ipsi- and contralateral ovaries to the dominant follicle in follicular phase animals (n = 3 cows) or to the corpus luteum in luteal animals (n = 3 cows) were collected for each measurement and were cut transversely in half to evaluate the different layers of the ovary separately (cortex and medulla). For porosity measurements, the cortex and the medulla were separately dissected as described in the methods. In (**b**) boxplot analysis of Young’s modulus measurements of ipsi- and contralateral ovaries of follicular and luteal phases of native (n = 6 ovaries) and decellularized (n = 6 ovaries) tissues. In (**c**) boxplot analysis of porosity data of ipsi- and contralateral ovaries of follicular and luteal phases of native (n = 6 ovaries) and decellularized tissues (n = 6 ovaries).

The only significant difference on YM was observed when comparing the native *versus* decellularized tissues in the contralateral medulla from the follicular phase (6.02±4.84 *vs* 2.06±1.12 kPa, p<0.01, Fig. 5b). All the other analysis were similar when comparing native *versus* decellularized, respectively, 7.37±4.44 *vs* 5.07±3.19 kPa (contralateral cortex at follicular phase), 2.51±1.64 *vs* 4.38±5.00 kPa (contralateral cortex at luteal phase), 6.03±3.54 *vs* 3.47±2.94 kPa (ipsilateral cortex at follicular phase), 5.43±3.01 *vs* 2.91±1.58 kPa (ipsilateral cortex at luteal phase), 2.18±0.91 *vs* 2.78±3.81 kPa (contralateral medulla at luteal phase), 3.41±1.55 *vs* 1.54±1.11 kPa (ipsilateral medulla at follicular phase), and 4.45±3.22 *vs* 2.51±1.33 kPa (ipsilateral medulla at luteal phase). No effects of estrous cycle or contra-ipsilateral positions on YM were observed for native tissues (for a full comparison of groups see Suppl. Data 1).

Although not statistically different, native cortex mean YM were higher than native medulla YM for both follicular ipsilateral (6.0 *vs* 3.4 kPa), follicular contralateral (7.4 *vs* 6.0 kPa), luteal ipsilateral (5.4 *vs* 4.4 kPa), and luteal contralateral (2.5 vs 2.2 kPa). Interestingly, a reduction in Young’s modulus was observed in all analyzed native tissues, when transitioning from medulla to cortex (900 - 1,050 μm distance; Suppl. Fig. 2).

Porosity was significantly higher in native than decellularized medulla during luteal phase for both contralateral (83.8±9.7 *vs* 61.3±12.8%) and ipsilateral (79.7±7.4 *vs* 56.7±6.9%) tissues (p<0.01 for both, Fig. 5c), but not during follicular phase (77.2±6.4 *vs* 63.3±12.8% in contralateral medulla and 75.7±7.0 *vs* 62.5±8.3% in ipsilateral medulla). Similarly, no differences were observed in cortex samples when comparing native *versus* decellularized, respectively, 69.5±9.2 *vs* 57.4±7.9% (contralateral-follicular), 65.6.5±14.3 *vs* 65.6±5.4% (contralateral-luteal), 71.8±17.3 *vs* 57.9±18.1% (ipsilateral-follicular), and 68.6±10.8 *vs* 59.1±7.8% (ipsilateral-luteal). No effects of estrous cycle or contra-ipsilateral positions on porosity were observed for native tissues (for a full comparison of groups see Suppl. Data 1).

A correlation study between YM and porosity was performed for cortex and medulla separately, a weak negative correlation for native cortex contralateral luteal (R=-0.59, p=0.034) and a strong positive correlation for decellularized cortex ipsilateral luteal (R=0.77, p=0.0021) were observed (Suppl. Fig. 3), while all other groups had no significant correlations. Similarly, weak correlations for native ipsi- (R=-0.49, p=0.033) and contralateral (R=0.48, p=0.014) follicular medulla samples were observed (Suppl. Fig. 4), while all other groups had no significant correlations.

In summary, the results showed that the decellularized tissues generally had lower YM values compared to the native tissues, but the differences were not statistically significant except for the contralateral medulla in the follicular phase. No significant effects of the estrous cycle or contra-ipsilateral positions were observed on YM or porosity in the native tissues, except for porosity differences between native and decellularized medulla during the luteal phase. Additionally, a reduction in YM was observed when transitioning from the medulla to the cortex in all analyzed native tissues, but no statistical differences were observed.

#### Young’s modulus and porosity of oviduct

For the analysis of oviductal tissue, ipsi- and contralateral oviducts from follicular and luteal phases of the estrous cycle were evaluated. Both nanoindentation and porosity were performed separately in the ampulla and isthmus segments. Multiple nanoindentation measurements were conducted with 200 μm increments for the analysis of luminal epithelium, stroma, muscular, and tunica layers as shown in Fig. 6a and Suppl. Fig. 5. Due to adherence to the nanoindentation probe and the difficulties of precisely measuring the thin oviductal samples, it was not possible to obtain sufficient measurements to analyze the tissuE′s layers as first planned (lumen-stroma surface *vs* muscle-*tunica* surface, Suppl. Fig. 6). Therefore, results are presented only separated by ampulla *vs* isthmus. For porosity measurements, all the tissue layers were analyzed as one, either in ampulla and isthmus segments.

**Figure 6.**
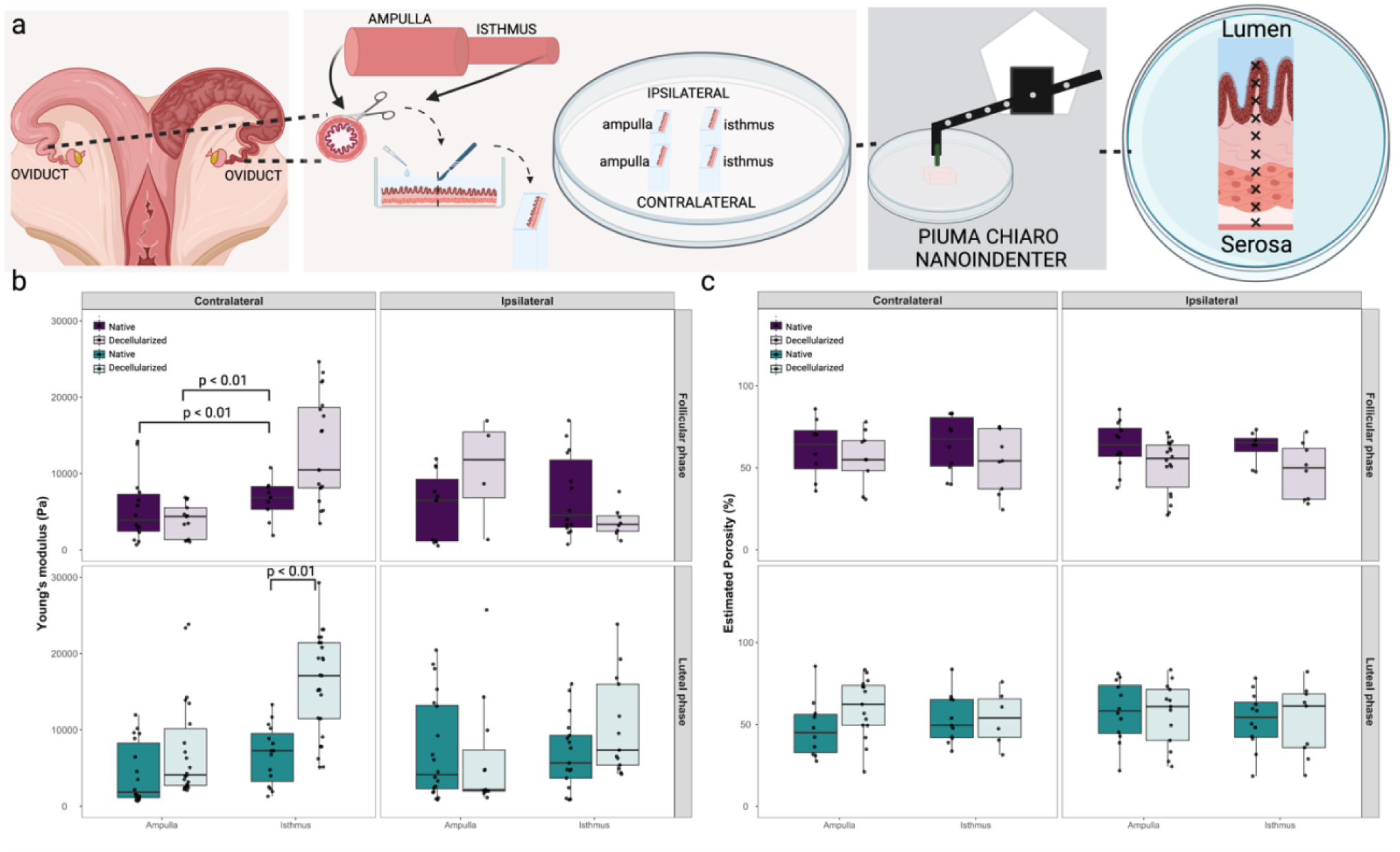
Young’s modulus and porosity analysis of native and decellularized oviducts. In (**a**) details of sampling for nanoindentation, ipsilateral (n = 3 cows) and contralateral (n = 3 cows) thin transversal slices of the ampulla and isthmus segments of the oviducts (ampullary-isthmic junction was discarded). Boxplots depicting Young’s modulus (**b**) and Porosity (**c**) measurements of ipsi- and contralateral ampulla and isthmus of follicular and luteal phases, for both native (n = 6 oviducts) and decellularized (n = 6 oviducts) tissues are shown.

The YM was only significantly different in the contralateral oviducts, either when considering the estrous cycle phase, in which the isthmus was stiffer than the ampulla at follicular phase (6.55±2.69 *vs* 5.36±4.37 kPa, p<0.01, Fig. 6b), or considering the decellularization process, in which the decellularized isthmus was stiffer than the native one at luteal phase (16.80±7.56 *vs* 6.74±3.82 kPa, p<0.01, Fig. 6b). All the other analysis were similar when comparing native *versus* decellularized, respectively, 5.36±4.37 *vs* 3.92±2.19 kPa (contralateral ampulla at follicular phase), 6.55±2.69 *vs* 14.12±7.73 kPa (contralateral isthmus at follicular phase), 5.46±4.57 *vs* 15.93±13.63 kPa (ipsilateral ampulla at follicular phase), 7.06±5.38 *vs* 3.69±1.98 kPa (ipsilateral isthmus at follicular phase), 4.27±4.00 *vs* 7.38±6.58 kPa (contralateral ampulla at luteal phase), 7.35±6.65 *vs* 12.64±14.08 kPa (ipsilateral ampulla at luteal phase), and 6.88±4.74 *vs* 13.32±9.60 kPa (ipsilateral isthmus at luteal phase). No other effects of estrus stage nor position were observed in native tissues (for a full comparison of groups see Suppl. Data 1). Nevertheless, although significance could not be determined, higher YM were observed in muscle-*tunica* surfaces than in lumen-stroma surfaces for native ampulla contralateral follicular (8.6 *vs* 2.1 kPa), native ampulla contralateral luteal (5.9 *vs* 1.7 kPa), native ampulla ipsilateral follicular (9.2 *vs* 1.0 kPa), native ampulla ipsilateral luteal (9.9 *vs* 2.6 kPa), which was less evident in native isthmus contralateral follicular (6.7 *vs* 5.3 kPa), native isthmus contralateral luteal (7.7 *vs* 4.9 kPa), native isthmus ipsilateral follicular (8.3 *vs* 4.0 kPa), and native isthmus ipsilateral luteal (7.7 *vs* 4.8 kPa; Suppl. Fig. 6). Regarding the porosity, no effects of estrous cycle or contra-ipsilateral positions on porosity were observed for all tissues (for a full comparison of groups see Suppl. Data 1).

A correlation study between YM and porosity was also performed for ampulla and isthmus separately. In the ampulla, a weak negative correlation for decellularized ampulla contralateral follicular (R=-0.56, p=0.045) and a weak positive correlation for native ampulla ipsilateral follicular (R=0.61, p=0.049) were observed (Suppl. Fig. 7), while all other groups had no significant correlations. In isthmus, a positive correlation for native ipsi- (R=0.63, p=0.017) and contralateral (R=0.71, p=0.031) follicular isthmus and a negative correlation for decellularized ipsilateral luteal (R=-0.65, p=0.0089) were observed (Suppl. Fig. 8), while all other groups showed no significant correlations.

In summary, a significant YM increase was observed in the isthmus than in the ampulla at follicular phase in contralateral native tissues, with decellularization leading to increased YM in contralateral luteal isthmus. Ipsilateral native ampulla tended to be stiffer than contralateral in both phases.

## Discussion

Tissue engineering holds great potential in regenerative medicine, particularly for reproductive tissues. The mechanical properties and porosity of tissues are fundamental considerations for successful tissue engineering approaches^9,10,18–20^. Here, we demonstrated that estrous cycle and/or decellularization can lead to alterations in mechanical properties and porosity of testis, endometrium, ovary, and oviduct, providing valuable insights into the dynamic changes of these properties and offer implications for tissue engineering strategies in reproductive medicine. By understanding and manipulating these characteristics, we can advance regenerative therapies and potentially restore reproductive capacity in patients.

Currently, various methods are employed to investigate the mechanical properties of biological tissues. These methods include techniques such as tensile testing, shear rheometry, and nanoindentation (reviewed by^21^). While each method has its advantages, nanoindentation offers distinct benefits for distinguishing different tissue components. Nanoindentation provides localized mechanical characterization by indenting the tissue surface with a small probe, allowing for precise measurements of mechanical properties at the microscale^22^. This makes it particularly suitable for distinguishing different tissue parts. By analyzing parameters like YM, storage modulus (E′), loss modulus (E″), and the E′-E″ difference, nanoindentation enables a comprehensive understanding of the mechanical behavior and relative contributions of elastic and viscous properties within specific tissue regions^22^. Moreover, it is important to consider that mounting tissue samples for rheological measurements can introduce challenges, particularly for porous tissues. The mounting process, which involves securing the tissue in place, by using holders or mounting solutions (such as agarose, agar, and commercially available glues) can potentially alter the tissuE′s mechanical properties. In the case of porous tissues, mounting solutions can penetrate the pores and introduce inaccuracies in the measurement of properties, like YM, and alter the interpretation of tissue rheology^23^. Here, a careful optimization of the mounting techniques was performed to minimize these effects and ensure accurate characterization of tissue rheology.

Adherence was a significant challenge in our nanoindentation measurements, particularly in the analysis of the oviductal tissue. Despite employing adherence mode, a technique aimed at reducing adherence effects by retracting the probe and holding it for a brief period, and using BSA to prevent adherence, the decellularized oviductal tissues exhibited heightened stickiness and stronger adhesion compared to the native tissues. This increased adherence posed difficulties in accurately assessing the mechanical properties of the oviduct, compromising the reliability and precision of the measurements. Careful consideration of alternative approaches or modifications to the testing methodology may be required to overcome the adherence issues and ensure accurate characterization of the mechanical behavior of adherent tissues. Despite the challenges posed by adherence and sample mounting in nanoindentation measurements, we were able to partially overcome these obstacles and perform accurate assessments of the mechanical properties of the analyzed tissues. Nevertheless, the sticky nature and increased adherence of decellularized tissues required careful consideration during the mounting process and we were not able to measure the different layers of the oviductal decellularized tissues.

In the male biomechanical analysis conducted, stark differences were observed between the *tunica albuginea* and the inner testicular tissue in terms of their storage and loss moduli. Both E′ and E″ were found to be significantly higher in the *tunica albuginea* compared to the inner testis. This suggests a greater elasticity and viscoelasticity in the *tunica albuginea*, indicative of a higher resistance to deformation and greater energy dissipation, respectively^24^. Furthermore, when evaluating the difference between the storage and loss moduli (E′-E″, also known as loss tangent), it was found to be greater in the *tunica albuginea* than in the inner testis. This finding implies that, relative to its viscous behavior, the *tunica albuginea* exhibits a stronger elastic response compared to the inner testicular tissue, which might suggest that the tunica is more resilient or resistant to permanent deformation, as it can store and release more of the energy from mechanical stresses rather than losing it^24^. This could be important in contexts where the tissue needs to undergo repeated or cyclical deformation. Collectively, these observations shed light on the distinct biomechanical properties of these two testicular components, which could have significant implications for their physiological roles and responses to mechanical stress.

Our analysis of the endometrial tissue yielded a significant finding wherein we observed a decrease in the difference between the E′-E″ in the ipsilateral horn during the follicular phase, in comparison to other phases and locations. This reduction in the E′-E′’ difference suggests a transition towards a more viscous mechanical behavior of the endometrial tissue during this phase. In essence, the tissue appears less resistant to deformation and exhibits enhanced capability for energy dissipation when subjected to applied stress. One plausible explanation for this observed biomechanical shift could be the fluctuating estradiol levels, known to peak during the follicular phase. Estradiol, a potent estrogen, is instrumental in modulating the structural and functional attributes of the endometrium, thereby priming it for implantation^25^. This hormone is known to influence endometrial proliferation, vascularization, and secretion. During the proliferative phase, estrogen interacts with estrogen receptors to induce mucosal proliferation, preparing the endometrium for the secretory phase^25,26^. Furthermore, estrogen also plays a role in endometrial vascularization by affecting the secretion of vascular endothelial growth factor (VEGF) from endometrial epithelial cells, which can increase subendometrial vascularity and blood flow^27^. Thus, the observed increase in viscous behavior in the ipsilateral horn during the follicular phase may be interpreted as a consequence of heightened estradiol levels. This could result in an endometrium that is more compliant and receptive, thereby effectively preparing the tissue for possible embryo implantation.

An intriguing aspect of the findings was the observation that the difference between the storage and loss moduli in the contralateral cortex and medulla was reduced during the luteal phase compared to the ipsilateral tissues. This decrease in the difference between E′ and E″ might suggest an increased viscoelasticity of the contralateral ovarian tissues during the luteal phase. This could potentially be linked to enhanced vascularization in these tissues. Enhanced vascularization, or the formation of new blood vessels, can influence the biomechanical properties of tissues^17^. The presence of more blood vessels can make a tissue more viscoelastic due to the additional fluid (blood) and complex network of flexible, compliant vessels^17^. Another interesting aspect to consider is the role of ovulation in these biomechanical changes. The side of ovulation (ipsilateral or contralateral) could influence the biomechanical properties of the ovarian tissues. In preparation for ovulation, the ovarian tissue undergoes significant changes, including increased vascularization and remodeling, to release the mature ovum (reviewed by^28^). These changes can significantly affect the tissuE′s mechanical properties, possibly explaining the reduced E′-E″ in the luteal phase of contralateral ovarian tissues. While these correlations between vascularization, ovulation, and increased viscoelasticity are plausible, it is important to note that these are hypotheses that would need to be confirmed with further studies, such as histological analysis or imaging studies to directly assess vascularization in these tissues during the different phases of the estrous cycle.

The oviduct was the least affected tissue when comparing storage and loss moduli. There were no significant differences in E′ and E″ in the isthmus when comparing estrous stage and position. However, in the luteal phase, a reduction in storage modulus was observed in the contralateral ampulla compared to the ipsilateral ampulla, and the luteal ipsilateral ampulla exhibited a decreased difference between storage modulus and loss modulus, suggesting increased viscoelasticity, particularly at higher frequencies. The lack of significant differences in E′ and E′’ observed in the oviduct may be attributed to the challenges in analyzing the tunica-muscle and lumen-stroma layers separately. As previously described, due to adherence to the probe, it was difficult to nanoindent these layers, and the data obtained represented a combination of all layers. When we analyzed the tunica-muscle and lumen-stroma layers separately (Suppl. Fig. 9), an evident effect of the estrous cycle and the position was observed, for both ampulla (Suppl. Table 7) and isthmus (Suppl. Table 8). Nevertheless, we should be cautious when analyzing these results due to the limited number of points analyzed (see Suppl. Fig. 5 for missing data points).

We subsequently focused on evaluating the YM of decellularized tissues and comparing them to their native counterparts. Decellularization plays a pivotal role in tissue engineering and regenerative medicine, as it aims to remove cellular components and derive extracellular matrix (ECM)-based scaffolds or hydrogels. Understanding the changes in mechanical properties resulting from decellularization is essential for optimizing tissue engineering strategies and designing biomimetic materials for different applications. Various decellularization methods include physical techniques like freeze-thawing cycles, chemical agents such as detergents, and biological agents like enzymes, each with varying impacts on ECM structure and the potential for recellularization or its use to form hydrogels^29–32^. In this process, detergents are commonly used and can have an impact on the structural integrity of the ECM. Components of the ECM, including collagen, glycosaminoglycans, and glycoproteins, like laminin, play crucial roles in cellular responses within engineered tissues^33^. Nevertheless, some detergents, such as sodium dodecyl sulfate (SDS) and Triton X-100, have been shown to denature collagen and not preserve laminin molecules, potentially influencing cellular behavior within the decellularized tissues^34,35^. In a previous study, we employed a decellularization protocol involving both SDS and Triton X-100 detergents as previously described^36^. This method resulted in inadequate preservation of certain ECM components, notably laminin (data not shown). Therefore, here, we used sodium deoxycholate (SDC) as the sole detergent, as it has been demonstrated to better preserve ECM components^37^. Moreover, the presence of residual detergents, genetic material, and cellular debris can adversely affect the behavior and functionality of scaffolds, impeding their intended applications^38^. Ensuring thorough removal of these components is essential for creating biocompatible and immunologically compatible tissue-engineered constructs that promote optimal cell growth, differentiation, and integration within the host tissue environment^29,38^. Here, a reduction of more than 96% was observed on DNA contents of the decellularized tissues, which was similar to other studies^36,37,39,40^, demonstrating the suitability of our decellularization protocol.

The decellularization process can significantly impact the mechanical properties of tissues, altering their stiffness and porosity. In here, decellularization had varying effects on the mechanical properties of different tissues. In the inner testis, decellularization resulted in a significant increase in YM, indicating increased stiffness. This trend was also observed in the decellularized endometrium and contralateral isthmus, where YM also increased. Conversely, decellularization led to a decrease in YM in the contralateral follicular medulla of the ovaries. These findings suggest that decellularization can have tissue-specific effects on YM. Several studies have demonstrated similar effects of decellularization on mechanical properties across various tissues. For instance, in decellularized cardiac tissue, a decrease in Young’s modulus and tensile strength was observed compared to native tissue^41^. Similarly, decellularized tendon constructs showed reduced stiffness compared to their native counterparts^42^. On the other hand, decellularized liver scaffolds exhibited increased YM^43^, and, in the case of arteries, the tensile modulus was found to be significantly higher in decellularized compared to native arteries, indicating an increased stiffness after decellularization^44^. These data highlight the tissue-specific responses to decellularization and emphasize the need for careful characterization of mechanical properties following the decellularization process.

Additionally, the decellularization process had varying effects on the porosity of the different tissues examined. Generally, the porosity of the decellularized tissues decreased compared to the native tissues. However, it is interesting to highlight that in the oviduct, no changes in porosity were observed after decellularization. These findings suggest that decellularization can significantly impact the porosity of tissues, and this effect may differ depending on the tissue type and the phase of the estrous cycle. Understanding the implications of changes in porosity is crucial for tissue engineering strategies, as it can influence cell infiltration, nutrient exchange, and overall tissue functionality (reviewed by^10,20^). Further investigation into the implications of altered porosity in decellularized tissues is warranted to optimize tissue engineering approaches for regenerative medicine applications. Despite discovering certain correlations between porosity and YM, the analysis conducted in the present study did not yield definitive conclusions regarding the precise impact of porosity on YM. This limitation arose due to the observed variability across different tissues included in this study.

In conclusion, this study provides valuable insights into the mechanical properties and porosity of reproductive tissues during different phases of the estrous cycle and following decellularization. The use of nanoindentation allowed for precise characterization of tissue mechanical properties at the microscale, revealing distinct biomechanical differences between tissue components. Despite challenges associated with adherence and sample mounting, accurate measurements of the mechanical properties were obtained, illuminating the unique characteristics of each tissue. Decellularization exhibited tissue-specific effects on stiffness and porosity, underscoring the importance of meticulous characterization of decellularized tissues for tissue engineering applications. These findings highlight the dynamic nature of tissue properties and their potential implications for tissue engineering strategies in reproductive medicine. The knowledge gained from this study can contribute to the development of biomimetic materials and regenerative therapies tailored to the specific needs of reproductive tissues. This customization ensures that engineered tissue constructs closely resemble their natural counterparts, advancing the field of reproductive medicine and regenerative therapies.

## Materials and Methods

### Tissue collection and preparation

Testicular samples were collected from bulls aged 22 to 23 months (n=3) from a local abattoir and were transported to the laboratory at room temperature (RT) within 2-3 hours of collection. Upon arrival, the testes were first rinsed with distilled water and then with 70% ethanol. The *tunica vaginalis* was incised to reveal the underlying *tunica albuginea* and seminiferous tubules. Following this, the *tunica vaginalis* was carefully removed to permit dissection of the *tunica albuginea*. Tissue samples of the *tunica albuginea* were subsequently excised, thinly sliced using a microtome blade, and set aside for further analyses. Similarly, internal slices of the testicular tubules and underlying stroma were collected. After dissection, the tissues were placed in a petri dish containing Phosphate Buffered Saline (PBS) and prepared for: (1) rheological analysis (Figure 3a); (2) porosity analysis; (3) DNA extraction and quantification; (4) DAPI staining; and (5) decellularization (Figure 7). All samples were kept on ice until further processing.

**Figure 7.**
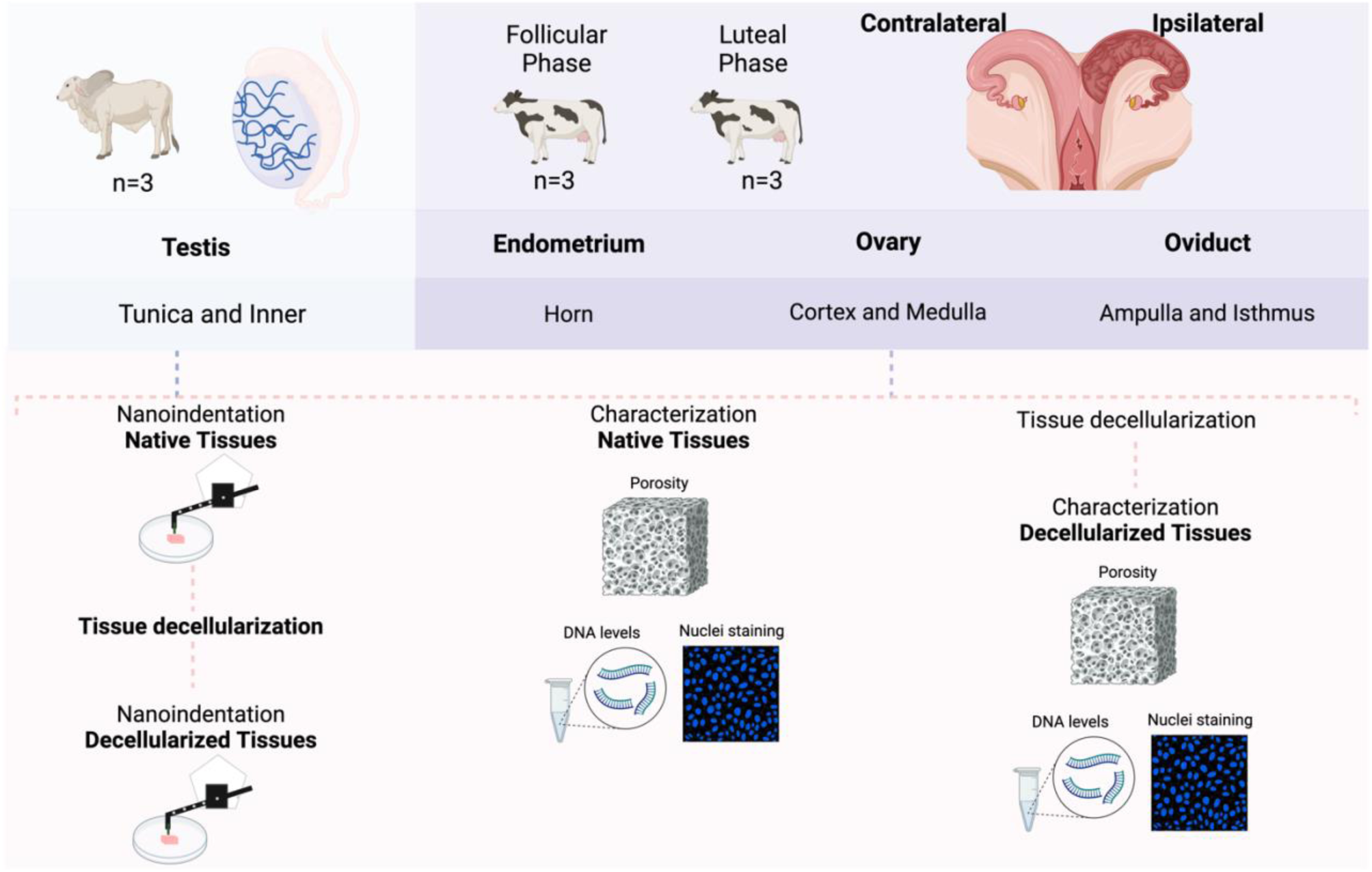
Experimental design: Testicular samples (n=3 bulls) and female reproductive tracts (n=6 cows) were collected from a local abattoir. The female reproductive tracts were identified either at follicular or luteal phase (n=3 per phase), ipsilateral to the dominant follicle (follicular phase) or to the *corpus luteum* (luteal phase). Samples of endometrium were collected from both horns, medulla and cortex samples from the ovaries, and ampulla and isthmus segments from the oviducts. All the samples were submitted to nanoindentation as native tissues, porosity analysis (n=3 technical replicate per tissue), characterization (including nuclei staining and DNA levels, n=1 sample per organ), and decellularization process. The same native samples analyzed in the nanoindenter were submitted to decellularization and analyzed again in the nanoindenter as decellularized tissue. After decellularization, samples were submitted to porosity analysis (n=3 technical replicate per tissue) and characterization (including nuclei staining and DNA levels, n=1 sample per organ) to validate the decellularization process.

Female reproductive tracts from cows aged 33 to 67 months old (n=6) were also collected from the local abattoir and carried to the laboratory at RT within 2-3 h of collection. The estrous cycle phase was determined in female tracts according to ovarian morphology based on follicle size and *corpus luteum* (CL) presence as previously described^45^. All female samples were separated according to their position in reference to the dominant follicle (> 11 mm; follicular phase animals, n=3) or the *corpus luteum* (luteal phase animals, n=3), being divided in ipsi- and contralateral (Figure 7). For oviductal tissue, the surrounding connective tissues were removed and the ampulla and isthmus were separated (ampullary-isthmic junction was discarded). For porosity, DNA, nuclei measurements, and decellularization, thin transversal slices of the tubes (ampulla and isthmus) were cut using a microtome blade. For nanoindentation, a 5 mm slice was separated, then the oviduct was cut open, an 8 mg/ml agarose solution (at 40°C) was poured on the lumen top and, after agarose hardening, a thin section was cut and positioned on a petri dish for analysis (Figure 6a). Uteri were opened and the endometrium dissected from the two horns (ipsi- and contralateral), pieces were immediately processed for porosity, DNA, nuclei measurements, and decellularization, or thin transversal slices were cut and mounted on a petri dish for nanoindentation (Figure 4a).

Ovaries were first cut transversionally in half, and then thin transversal slices were cut and mounted in a petri dish for nanoindentation of all ovarian layers (*Tunica albuginea*, cortex, and medulla, Figure 5a). For porosity, DNA, nuclei measurements, and decellularization, cortex and medulla were isolated separately. For isolating the cortical part of the ovaries, a cutting mold based on the design by Kagawa et al.^46^, with a 0.5 x 10 x 10 mm (HxWxL) space cutting area and a plate was designed using Adobe Fusion 360 (supplementary figure 10). The .stl 3D file was then sliced for 3D printing by using a 3D printing preprocessing software (Chitubox). The cutting mold was then printed using an Elegoo Mars Pro 3D printer. Molds were washed in isopropanol for 5 min and cured by UV-light exposure for 5 min, using the Elegoo Mercury Plus. Before use, the 3D-printed molds were sterilized by UV light for 20 min. The mold was, then, used to cut thin slices of cortex (outside 0.5 mm layer) and medulla separately. All tissues used for nanoindentation were collected and processed for decellularization, to be used for decellularized tissue nanoindentation. A schematic representation of the experimental design is illustrated in Figure 7.

### Nanoindentation analysis

In general, each tissue piece was indented using a Piuma Chiaro nanoindenter (Optics 11). Indentation depth was set to 5 μm. The probe had a glass spherical tip (diameter 28 μm for native tissues and 8.5 μm for decellularized tissues) mounted on an individually calibrated cantilever with a spring constant of ~0.5 N m^−1^. Deformation of the cantilever following contact with the biological sample was measured by tracking the phase-shift in light, reflected from the back of the cantilever. The Young’s modulus (YM) and the effective Young’s modulus (efYM) were calculated using the built-in PIUMA software based on a linear Hertzian contact model build on the first 10% of the force– distance curve. The elastic and viscoelastic moduli (E′ and E”, respectively) were determined by using the same probe and equipment using the indentation mode, with frequencies of 1, 5, 10, and 20 Hz, 300 nm amplitude, 1 s slope time, initial relaxation of 10 s, relaxation of 2 s and 5 μm indentation depth. Furthermore, in all experiments, the Piuma attachment mode with a 650 μm distance and 5 s wait was used to prevent probes from starting in contact with tissues, a z-above surface of 20 μm, a speed of 80 μm s^-1^ and a threshold of 0.0001 were used in all measurements. All experiments were performed in PBS with 5% bovine serum albumin to reduce sample attachment, at RT.

For testicular tissues, a matrix indentation was used for determining both YM, efYM, E′, and E”, using parameters described above. Four measurement points were taken with a 100 μm distance in the x-axis. For the oviduct, 8-12 measurement points were taken with a 200 μm distance in the x-axis, being the first 400 μm estimated to correspond to the luminal epithelium and stroma layers, the following measurements to the muscular and tunica layers. For the endometrium indentation, 5-10 points were randomly selected and measured. For the ovary, 8-12 measurement points were taken with a 150 μm distance in the x-axis, being the first 750 μm estimated to correspond to the medulla layer, and the following measurements to the cortex and tunica albuginea layers.

### Porosity measurements

The tissue slice volume was determined using a liquid displacement method^47^. Following volume measurement, the samples were frozen at −80°C for 1 h and subjected to lyophilization overnight. The lyophilized samples were then weighed to obtain their dry weight before being immersed in pure ethanol for an overnight incubation at RT. Post-incubation, samples were carefully removed from the ethanol, the excess liquid was blotted off, and the samples were weighed again. The relative porosity (%) was then calculated based on these measurements using the equation provided below:

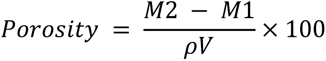

where M2 represents the mass of the samples after ethanol incubation (g), M1 means the dry weight after lyophilization (g), *ρ* is the ethanol density (0.789 g/cm^3^), and V is the volume (cm^3^).

### Decellularization using sodium deoxycholate detergent

After dissection of tissues, tissue pieces were placed in sterile ultrapure water and incubated overnight at 4°C on a rotator, at a volume of 10 mL solution per 1 g of tissue, to remove red blood cells. The next day, washed tissue pieces were placed in decellularization solution with 4% Sodium Deoxycholate (SDC) in sterile ultrapure water and incubated at 4°C for 24 h on a rotator at a volume of 10 mL solution per 1 g of tissue. After 24 h of decellularization, tissues were washed at RT on a rotator 3 times for 5 min in sterile ultrapure water to remove all remaining detergent. Next, tissues were treated with DNase I solution (5 μg mL^-1^ in ultra-pure water) for 3 h at RT on a rotating rack. Afterwards tissues were transferred to sterile ultrapure water and washed on a rotator 3 times for 5 min.

### Nuclear staining of tissues for determining efficiency of decellularization protocol

Native and decellularized tissue pieces were washed in PBS and incubated for 3 h at RT with 4% paraformaldehyde (PFA). After PFA removal, samples were washed and kept in PBS at 4°C. For staining, samples were incubated for 30 min in a 0.5 µL mL^-1^ Hoechst 33342 (20 mM) solution in PBS at RT, then washed and placed in a 48-well dish for imaging using an EVOS M700 fluorescence microscope (Thermo Fisher Scientific) with ×4 and ×10 objectives.

### DNA extraction and quantification

DNA purification of native and decellularized tissues was performed using the DNA Purification Thermo Scientific Kit from ThermoFisher (K0512) according to manufacturer’s instructions, using between 26-38 mg of tissue. Tissue pieces were first homogenized in a lysis buffer for 1 min prior to the start of extraction. Quantification of DNA in samples was performed using Qubit 1X dsDNA BR Kit with Qubit 4 Fluorometer (Invitrogen), according to manufacturer’s instructions.

### Statistics and Reproducibility

We were interested in studying the differences on porosity and mechanical properties of different native and decellularized reproductive tissues. To assess the effects of tissue, decellularization (Native vs. Decellularized), estrous cycle (Follicular phase vs. Luteal phase), and position (Contralateral vs. Ipsilateral) on Young’s modulus (YM), porosity, or storage and loss moduli (E′ and E″) we used a linear model with a Tukey’s post hoc. First, a factor variable named “Comparison” was created in the dataset to represent the combinations of “Estrous_cycle,” “Position,” and/or “Cellularization” using the paste() function. The “Comparison” variable was converted to a factor to ensure appropriate handling of categorical data. Next, a linear model was fitted using the lm() function, with “Indentation” (for E′ and E″ data analysis), “Porosity”, or “YM” as the response variable and “Comparison” as the predictor variable. Each tissue specific dataset “df” was used as the data source for the model. To perform pairwise comparisons using Tukey’s method, the glht() function from the “multcomp” package was employed. The linfct argument within glht() was set to mcp(Comparison = “Tukey”), indicating that Tukey’s method should be used for the pairwise comparisons. Finally, the results were summarized using the summary() function applied to the “glht” object. The test argument within summary() was set to adjusted(type = methods), where “methods” is a vector containing the desired methods for the pairwise comparisons. The common methods used for controlling the family-wise error rate, such as “adjusted”, “single-step”, “sidak”, “bonferroni”, and “holm”, were included in the vector. The significance level was set at α = 0.05 for all statistical tests.

To assess the relationship between YM and porosity, a correlation analysis was performed separately for each combination of tissue type and cellular category. The Pearson correlation coefficient was calculated to measure the strength and direction of the linear relationship between the variables. To further investigate the relationship between YM and porosity, linear regression analysis was conducted for each combination of tissue type and decellularization category. The linear regression model was fitted to the data, with YM as the dependent variable and Porosity as the independent variable. The model provided estimates of the slope and intercept, indicating the direction and magnitude of the relationship. The significance of the regression coefficients was assessed using p-values. All data analysis and visualization were carried out in R (ver. 4.3.1), and the scripts and packages used for carrying out our analyses are described in supplementary file 2.

## Availability of data and materials

All data produced and analyzed in this project is available within the main manuscript and as supplementary files.

## Supplementary material

Supplementary file 1: Suppl. Figures 1-10

Supplementary file 2: All R code to replicate the analysis performed in this study

Supplementary data 1: full set of statistical comparisons of all analyzed data.

## Author’s contributions

Conceptualization: all; Methodology: all; Investigation: all; Supervision: MAMMF; Writing—original draft: all; Writing—review & editing: all authors. This was a team effort Project SLAM, and the order of co-first authors provided here was decided through a Murder Mystery SLAM Party. Co-first authors can prioritize their names when adding this paper’s reference to their résumés.

## Supporting information

R code for all analyses

## Acknowledgements

This work was supported by the Alexander von Humboldt Foundation in the framework of the Sofja Kovalevskaja Award endowed by the German Federal Ministry of Education and Research.

