## Supplementary file 1 for "Mechanical Properties of Native and Decellularized Reproductive Tissues: Insights for Tissue Engineering Strategies"

### Supplementary files

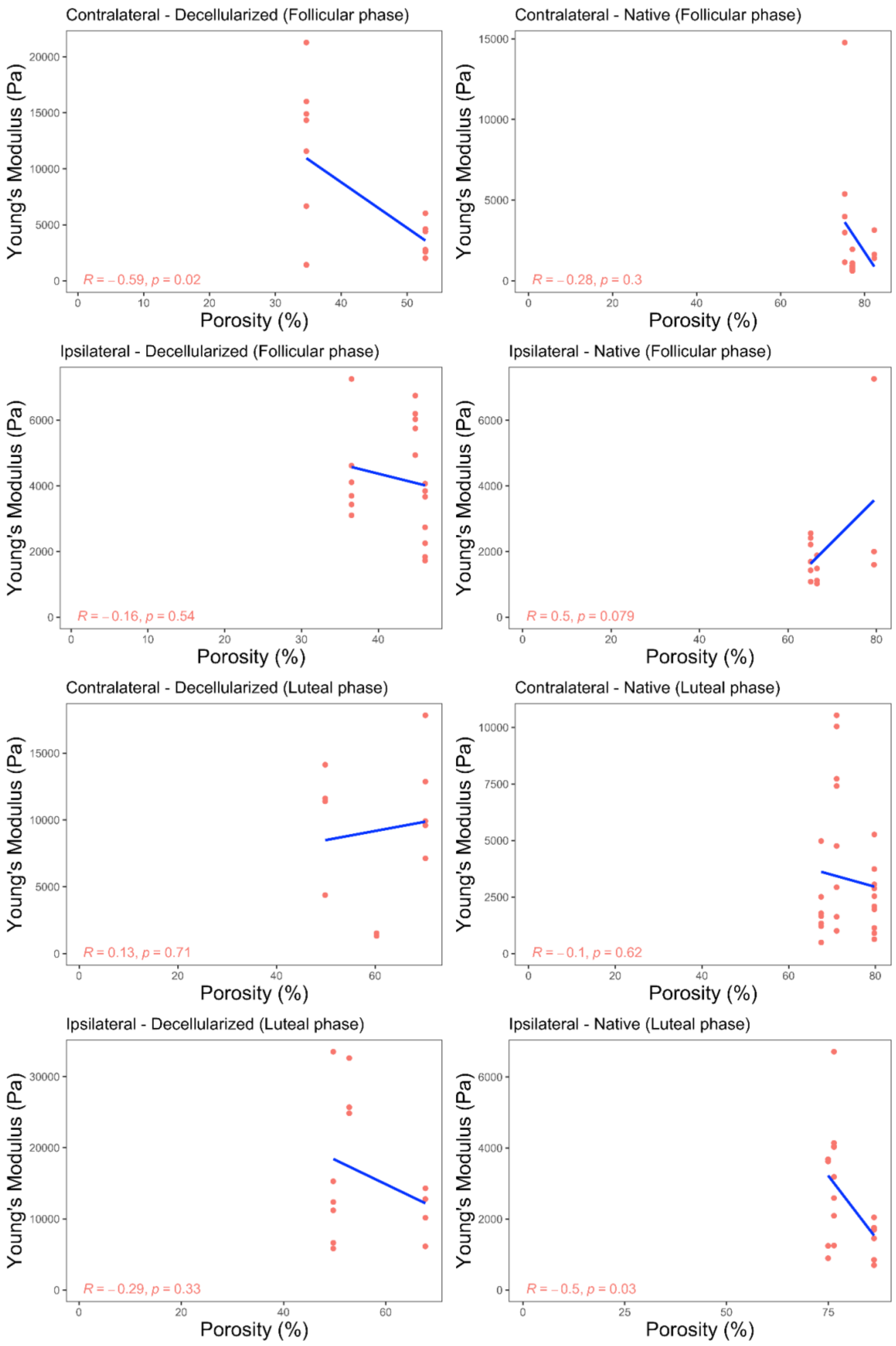


**Supplementary Figure 1.** Correlation plots of porosity and Young's modulus in endometrium tissues. Analysis was performed in native and decellularized tissues collected from ipsi- and contralateral horns of endometrium tissues from cows at luteal (n = 3 cows) and follicular (n = 3 cows) phases of the estrous cycle

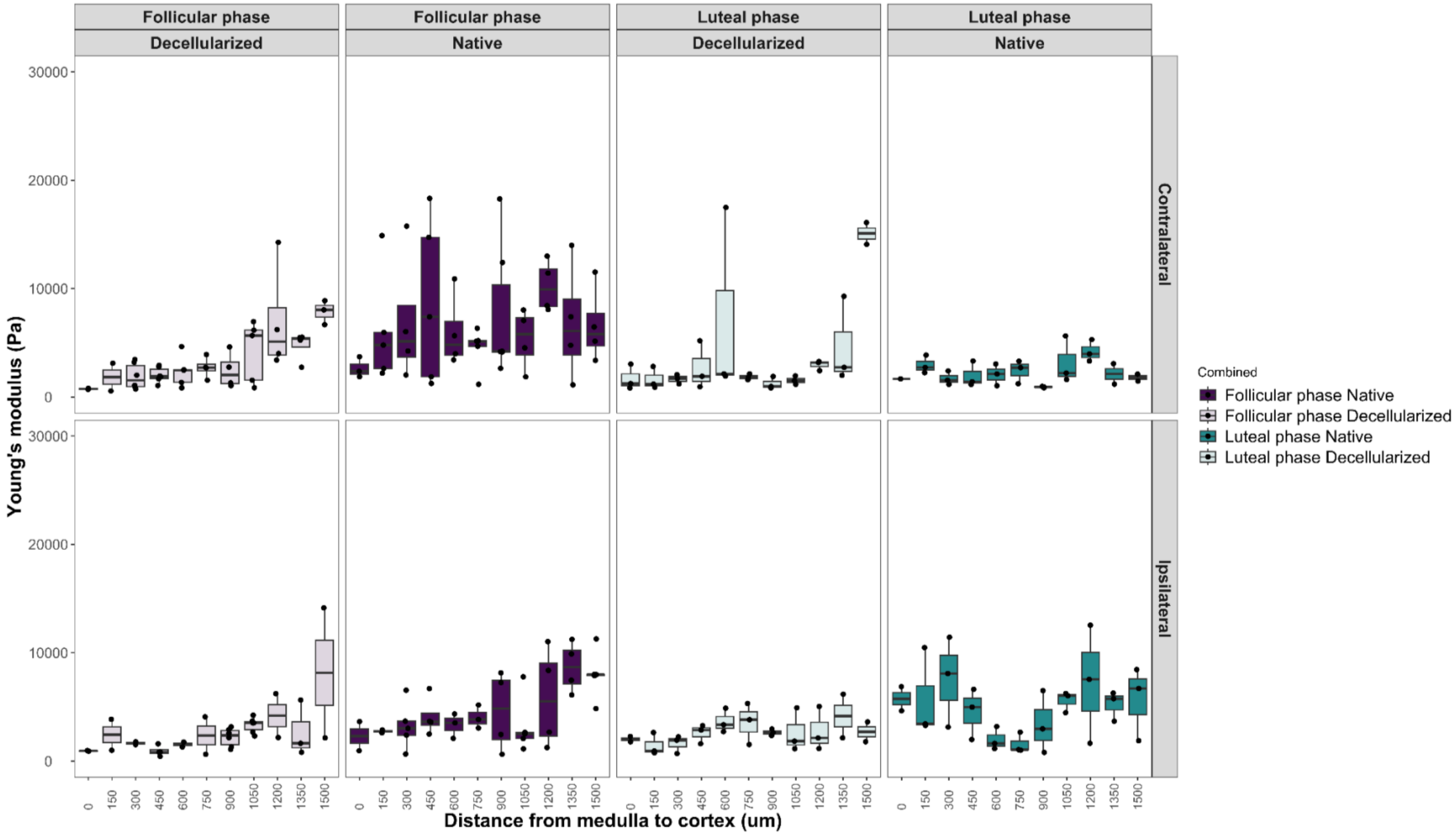


**Supplementary Figure 2.** Young's modulus analysis in native and decellularized ovaries through nanoindentation specific distance points. After a transversal cut, the ovary tissues were mounted on a petri dish using 8 mg $mL^{-1}$ agarose solution as a single piece, where the nanoindentation analysis was performed in a straightness acquisition mode with a 150 μm distance between analysis points, resulting in 1,500 μm x-axis distance in total, which 0 μm was the most external point and 1,500 μm the most inner. Analysis was performed in native and decellularized tissues collected from ipsi- and contralateral ovaries from cows at luteal (n = 3 cows) and follicular (n = 3 cows) phases of the estrous cycle.

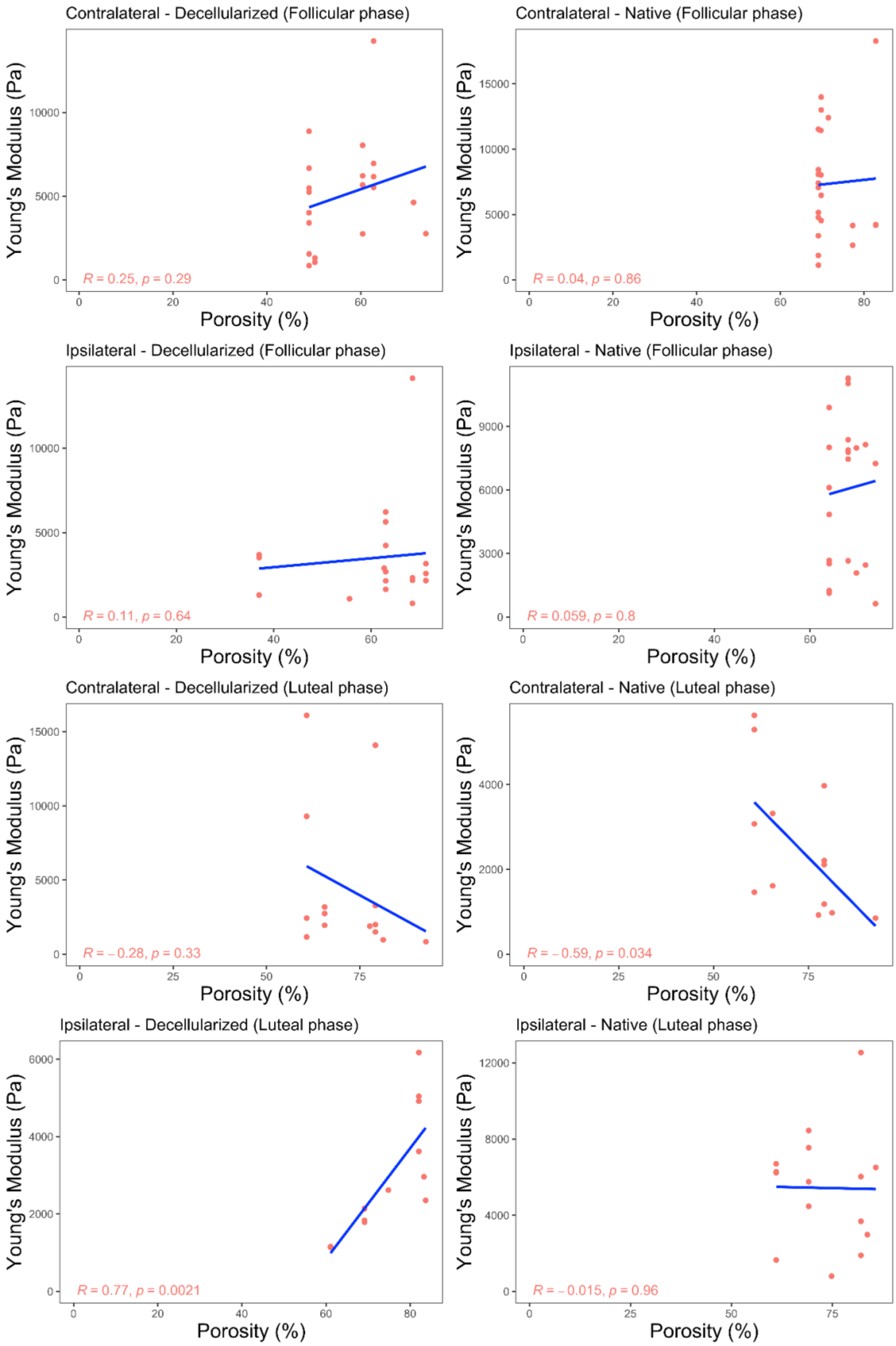


**Supplementary Figure 3.** Correlation plots of porosity and Young's modulus in the cortical segment of the ovary. Analysis was performed in native and decellularized tissues collected from ipsi- and contralateral ovaries from cows at luteal (n = 3 cows) and follicular (n = 3 cows) phases of the estrous cycle.

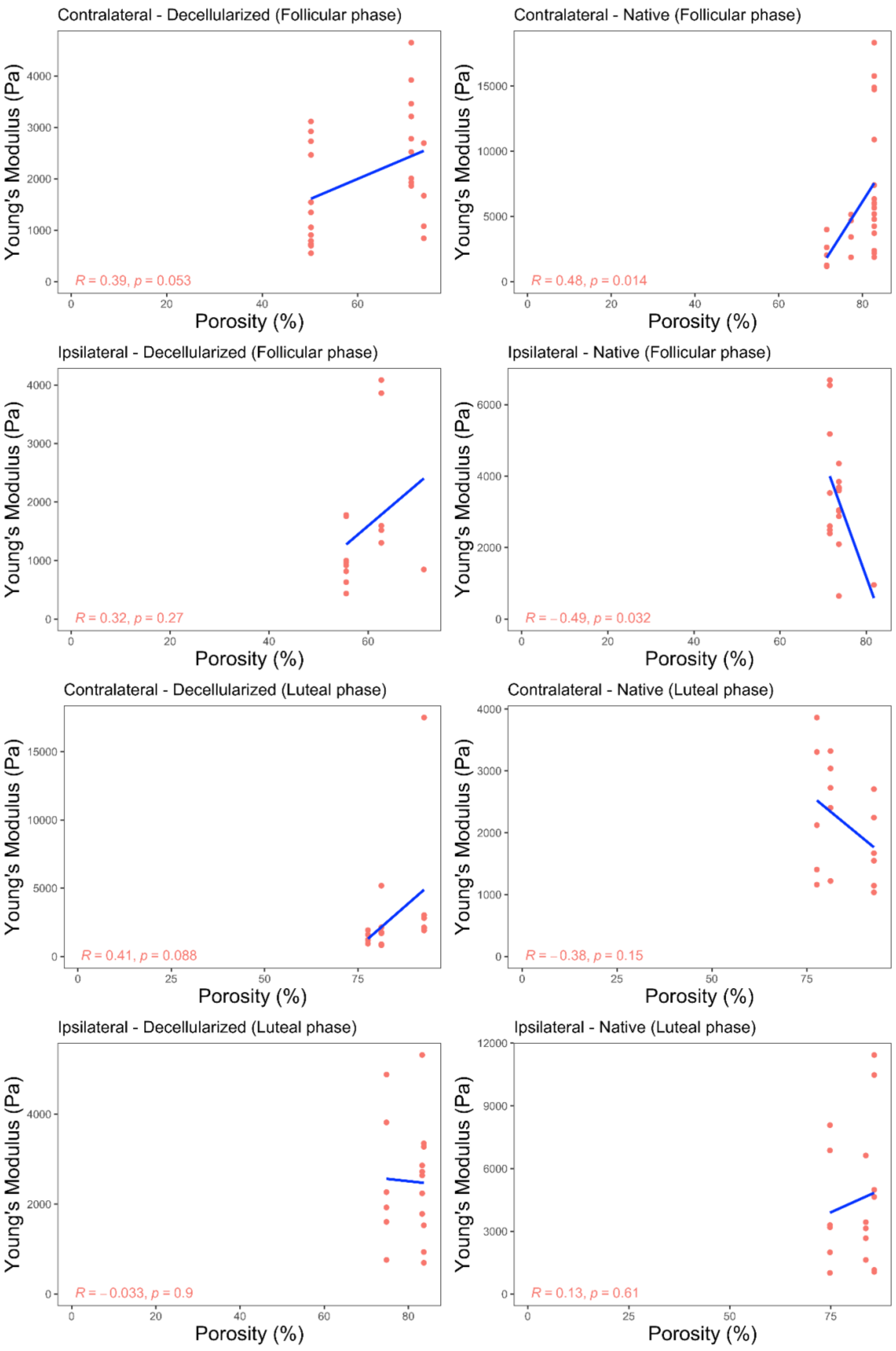


**Supplementary Figure 4.** Correlation plots of porosity and Young's modulus in the medullar segment of the ovary. Analysis was performed in native and decellularized tissues collected from ipsi- and contralateral ovaries from cows at luteal (n = 3 cows) and follicular (n = 3 cows) phases of the estrous cycle.

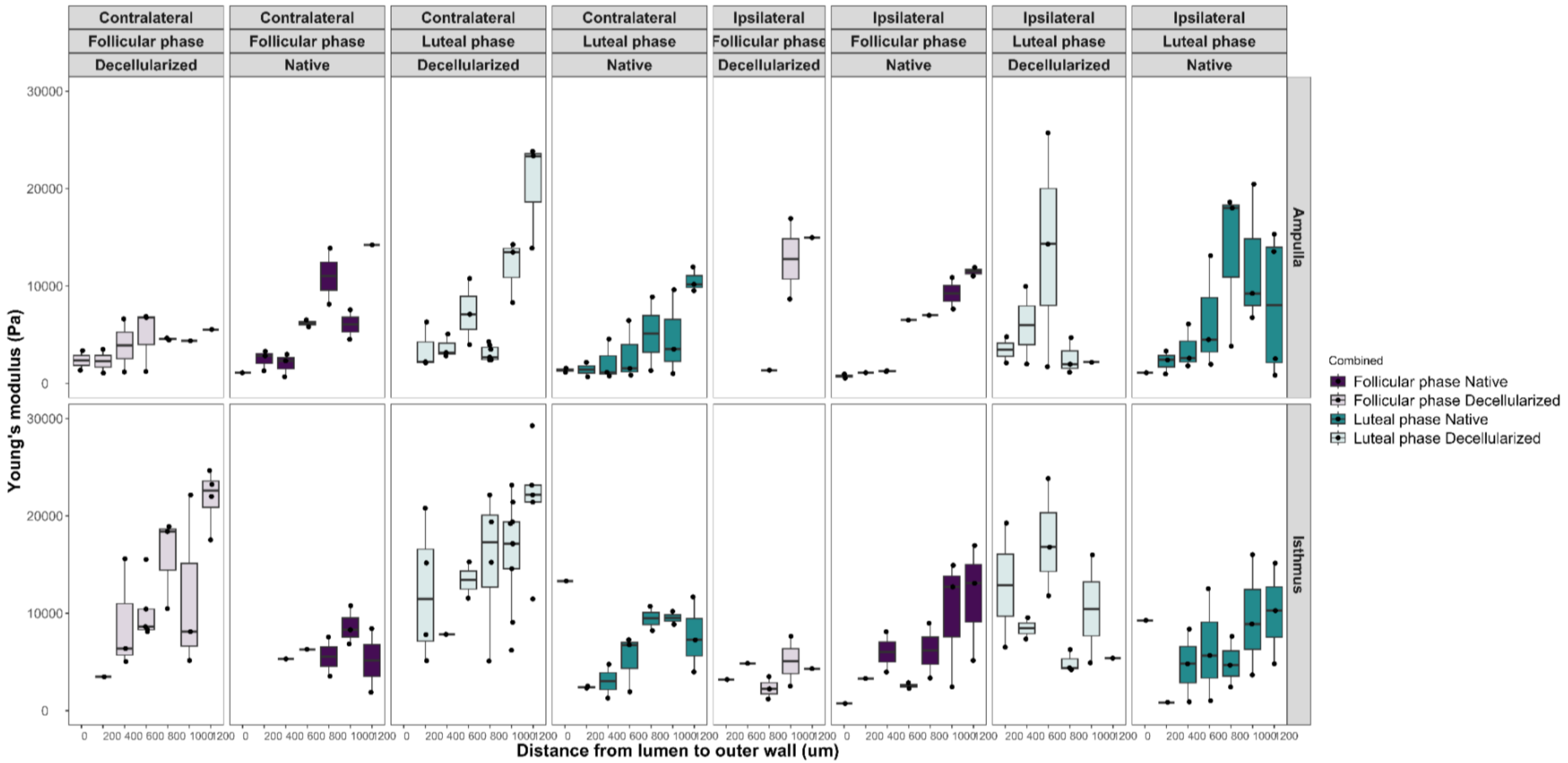


**Supplementary Figure 5.** Young's modulus analysis in native and decellularized oviductal samples through nanoindentation specific distance points. After a transversal cut, the oviductal tissues were longitudinally opened and mounted sideway on a petri dish using 8 mg $mL^{-1}$ agarose solution, this way all the tissue's layers were free for nanoindentation. For nanoindentor analysis, a straightness acquisition mode with a 200 μm distance between analysis points was performed, resulting in 1,200 μm x-axis distance in total, which 0 μm was the most internal point (luminal oviductal side) and 1,200 μm the most external (tunica oviductal side). Analysis was performed in native and decellularized tissues collected from ipsi- and contralateral ovaries from cows at luteal (n = 3 cows) and follicular (n = 3 cows) phases of the estrous cycle.

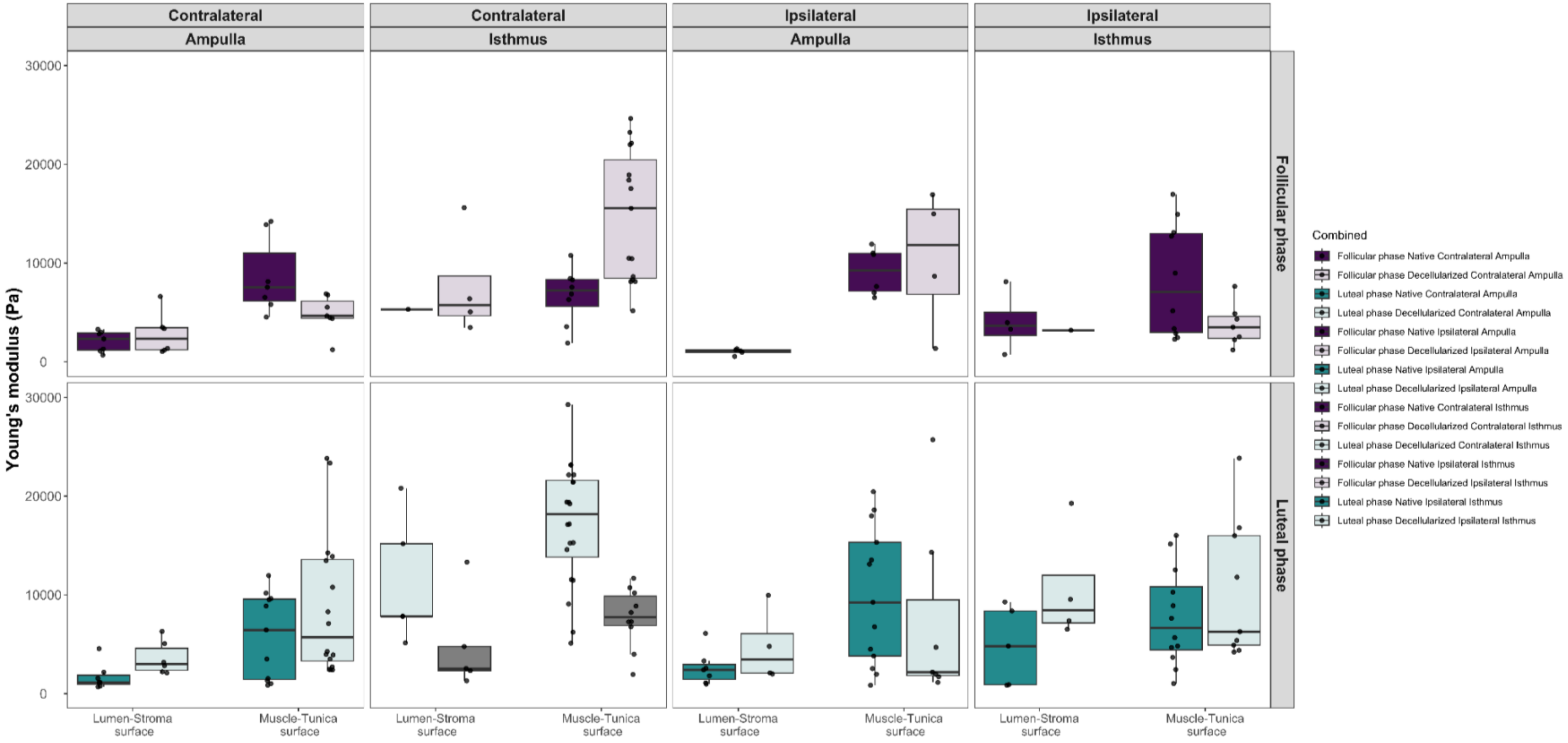


**Supplementary Figure 6.** Young's modulus analysis in native and decellularized oviductal samples derived from supplementary figure 5. All the specific distance points presented in Supplementary Figure 5 were divided into two segments with 400 µm as threshold, which points lower than 400 µm were classified as Lumen-Stroma surface, and values higher than it was classified as Muscle-Tunica surface. Analysis was performed in native and decellularized tissues collected from ipsi- and contralateral ovaries from cows at luteal (n = 3 cows) and follicular (n = 3 cows) phases of the estrous cycle.

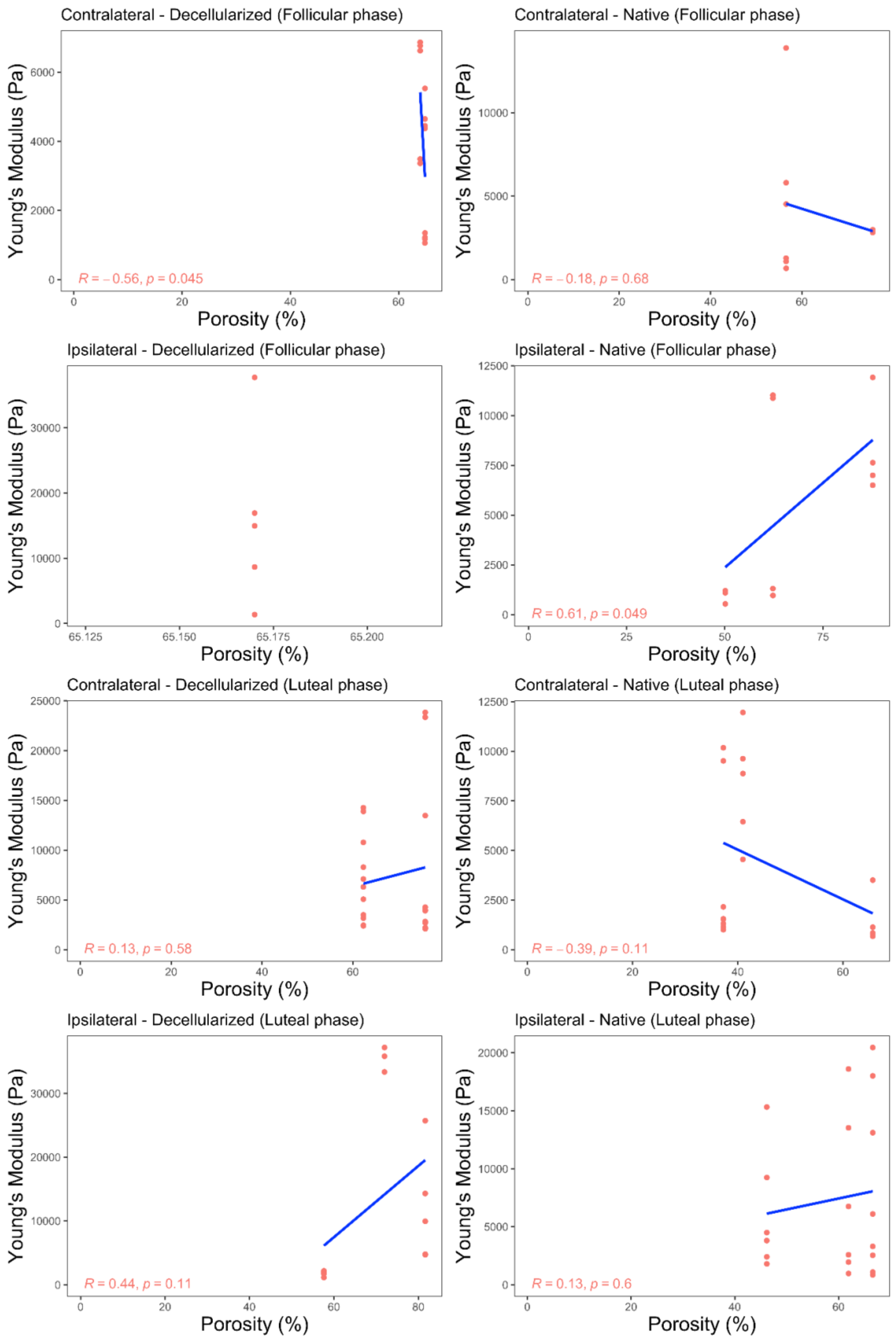


**Supplementary Figure 7.** Correlation plots of porosity and Young's modulus in the ampullary segment of the oviduct. Analysis was performed in native and decellularized tissues collected from ipsi- and contralateral oviducts from cows at luteal (n = 3 cows) and follicular (n = 3 cows) phases of the estrous cycle.

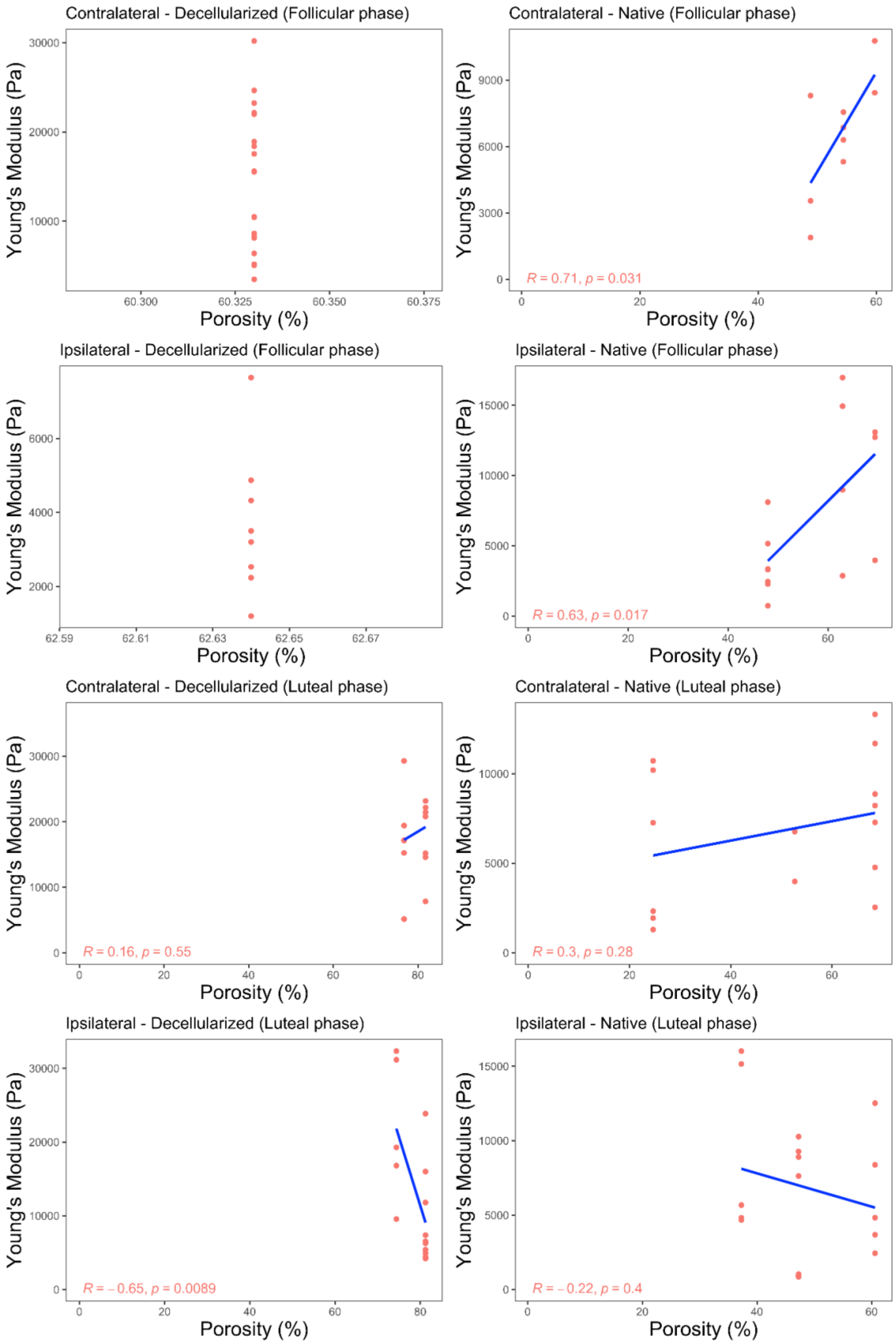


**Supplementary Figure 8.** Correlation plots of porosity and Young's modulus in the isthmic segment of the oviduct. Analysis was performed in native and decellularized tissues collected from ipsi- and contralateral oviducts from cows at luteal (n = 3 cows) and follicular (n = 3 cows) phases of the estrous cycle.

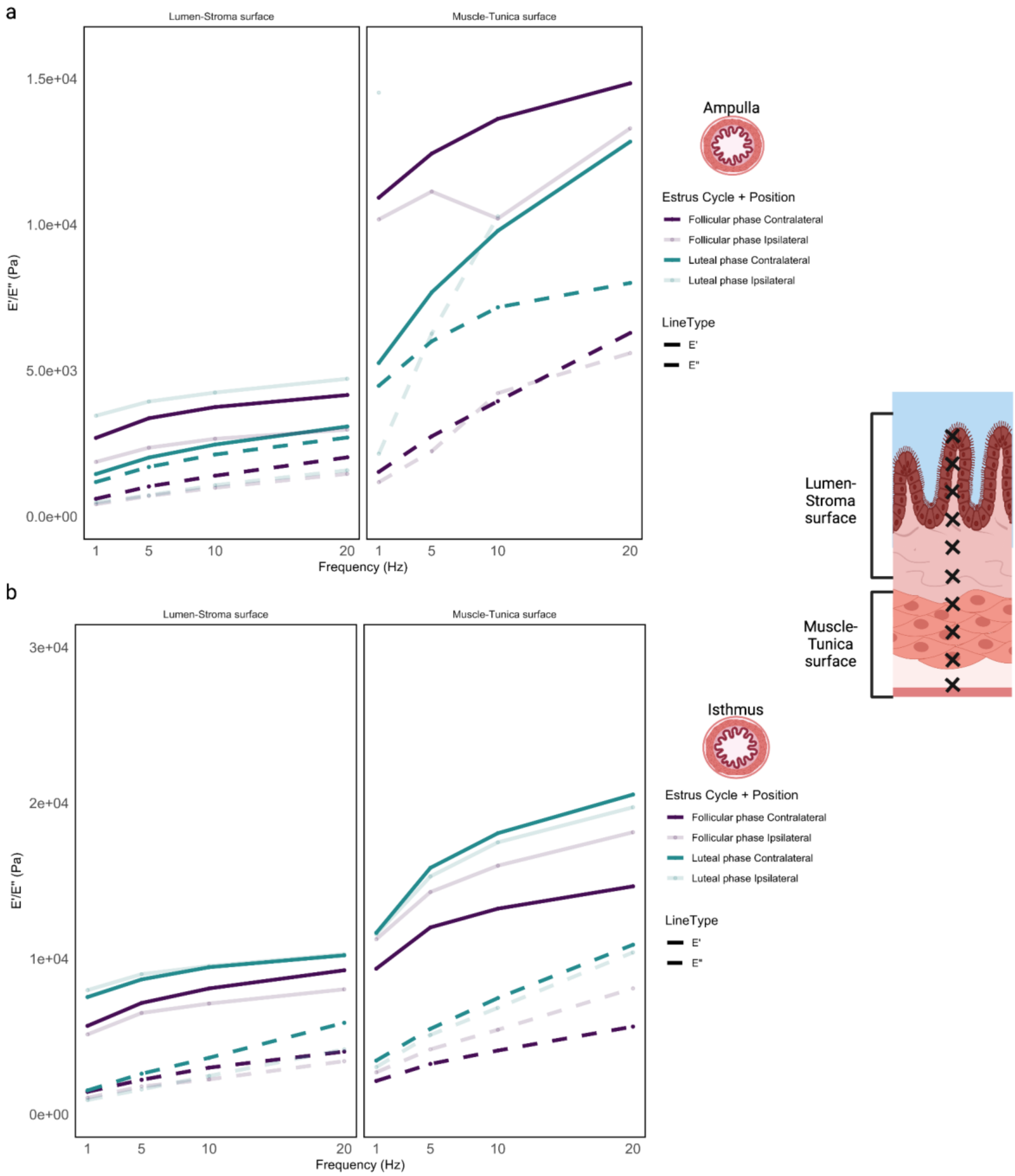


**Supplementary Figure 9.** Elastic and visco-elastic properties of the oviduct differentiated by specific distance points presented in Supplementary Figure 5 . Storage (E', continuous lines) and Loss (E", dashed lines) modulus of native oviductal ampulla (**a**) and isthmus (**b**). Oviduct layers were divided as Luteal-Stroma surface and Muscle-Tunica surface, and samples were analyzed for both follicular and luteal phases and ipsi- and contralateral positions.

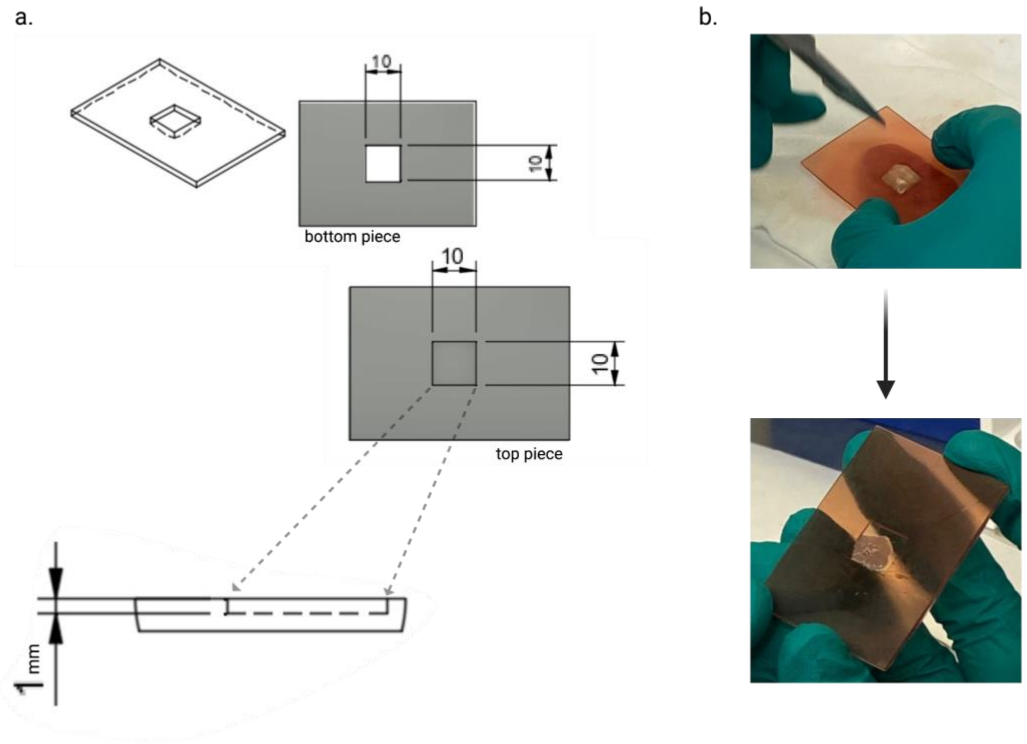


**Supplementary Figure 10.** Ovary slicer 3D print model (a) and example of the printed construct with sliced cortex (b).
