## Supplementary material for "Mechanical Properties of Native and Decellularized Reproductive Tissues: Insights for Tissue Engineering Strategies": R code for all analyses

### Project_SLAM.R

ra98gom

2023-10-09

```r
setwd("~/Desktop/R/Project SLAM 2023/Rheology")

library(reshape2)
library(nlme)
library(multcomp)
```

```
## Loading required package: mvtnorm

## Loading required package: survival

## Loading required package: TH.data

## Loading required package: MASS

##
## Attaching package: 'TH.data'

## The following object is masked from 'package:MASS':
##
##     geyser
```

```r
library(viridis)
```

```
## Loading required package: viridisLite
```

```r
library(ggplot2)
library(gridExtra)
library(lme4)
```

```
## Loading required package: Matrix

##
## Attaching package: 'lme4'

## The following object is masked from 'package:nlme':
##
##     lmList
```

```r
library(scales)
```

```
##
## Attaching package: 'scales'

## The following object is masked from 'package:viridis':
##
##     viridis_pal
```

```r
library(ggplot2)
library(cowplot)
library(ggpubr)
```

```
##
## Attaching package: 'ggpubr'

## The following object is masked from 'package:cowplot':
##
##     get_legend
```

```r
library(broom)
library(dplyr)
```

```
##
## Attaching package: 'dplyr'

## The following object is masked from 'package:gridExtra':
##
##     combine

## The following object is masked from 'package:MASS':
##
##     select

## The following object is masked from 'package:nlme':
##
##     collapse

## The following objects are masked from 'package:stats':
##
##     filter, lag

## The following objects are masked from 'package:base':
##
##     intersect, setdiff, setequal, union
```

```r
############################################################################
###################### Decellularization DNA analysis ##########################
############################################################################

# Read the dataset from a CSV file
df <- read.csv("DNA.csv")

#Summary data DNA
summary_df <- df %>%
  group_by(Cellular, Sample) %>%
  summarise(mean_DNA = mean(DNA),
            sd_DNA = sd(DNA))
```

```
## `summarise()` has grouped output by 'Cellular'. You can override using the
## `.groups` argument.
```

```r
#PLot DNA


png(filename="DNA.png",
    width = 15, height = 7, units = "in",
    res = 600)
```

```r
par(mgp=c(1.5,0.5,0),
    mar = c(4, #bottom
            4, #left
            3, #top
            2),
    cex.main = 0.8,
    family = "sans" #right
)


# Define color palette
color_cellular_native <- viridis_pal(option = "D")(1)
color_cellular_decellularized <- alpha(viridis_pal(option = "D")(1), 0.2)

# Create the plot using ggplot
ggplot(data = df, aes(x = Sample, y = DNA, fill = Cellular)) +
  geom_boxplot(width = 0.7, outlier.shape = NA) +
  geom_jitter(position = position_jitterdodge(), alpha = 0.8) +
  xlab("Tissue") +
  ylab("DNA concentration (ng/ul)") +

  # Color scale for 'fill' aesthetic
  scale_fill_manual(values = c("Native" = color_cellular_native,
                              "Decellularized" = color_cellular_decellularized)) +

  theme_bw() +
  ylim(0, 1100) +
  theme(panel.grid.major = element_blank(), panel.grid.minor = element_blank(),
        axis.text.x = element_text(size = 14, angle = 0, hjust = 0.5),
        axis.text.y = element_text(size = 14, angle = 0, hjust = 0.5),
        axis.title.x = element_text(size = 14, face = "bold"),
        axis.title.y = element_text(size = 14, face = "bold"),
        legend.position = "right",
        strip.text = element_text(size = 14, face = "bold"))
```

```
## Warning: Removed 1 rows containing non-finite values (`stat_boxplot()`).

## Warning: Removed 1 rows containing missing values (`geom_point()`).
```

```r
dev.off()
```

```
## pdf
##   2
```

```r
# Perform Tukey's HSD test for each Sample
tukey_results <- list()
for (sample in unique(df$Sample)) {
  subset_df <- df[df$Sample == sample, ]
  glm_model <- aov(DNA ~ Cellular, data = subset_df)
  tukey_result <- TukeyHSD(glm_model, "Cellular")
  tukey_results[[sample]] <- tukey_result
}

# Summarize the Tukey's HSD results
for (sample in names(tukey_results)) {
```

```r
  cat("Tukey's HSD results for Sample:", sample, "\n")
  print(tukey_results[[sample]])
  cat("\n")
}
```

```
## Tukey's HSD results for Sample: Oviduct
##   Tukey multiple comparisons of means
##     95% family-wise confidence level
##
## Fit: aov(formula = DNA ~ Cellular, data = subset_df)
##
## $Cellular
##                          diff      lwr      upr     p adj
## Native-Decellularized 304.803 98.10747 511.4985 0.0069134
##
##
## Tukey's HSD results for Sample: Ovary
##   Tukey multiple comparisons of means
##     95% family-wise confidence level
##
## Fit: aov(formula = DNA ~ Cellular, data = subset_df)
##
## $Cellular
##                           diff      lwr      upr     p adj
## Native-Decellularized 451.8935 241.8231 661.9639 0.0003574
##
##
## Tukey's HSD results for Sample: Endometrium
##   Tukey multiple comparisons of means
##     95% family-wise confidence level
##
## Fit: aov(formula = DNA ~ Cellular, data = subset_df)
##
## $Cellular
##                           diff      lwr      upr     p adj
## Native-Decellularized 557.0943 250.2549 863.9337 0.0016206
##
##
## Tukey's HSD results for Sample: Testis
##   Tukey multiple comparisons of means
##     95% family-wise confidence level
##
## Fit: aov(formula = DNA ~ Cellular, data = subset_df)
##
## $Cellular
##                           diff      lwr      upr     p adj
## Native-Decellularized 350.3433 112.0012 588.6855 0.0129092
```

```r
#################################################################
#################### Male Analysis ##############################
#################################################################

################### E' E" #######################################
```

```r
# Read data
data <- read.csv("E1_E2_male.csv")

# Set up the plot dimensions and margins
png(filename = "E1_E2_male.png",
    width = 8.86, height = 6.86, units = "in",
    res = 600)

par(mgp = c(1.5, 0.5, 0),
    mar = c(4, 4, 3, 2),
    cex.main = 0.8,
    family = "sans")

# Calculate means and standard deviations
data_summary <- data %>%
  group_by(Tissue, Module, Hz) %>%
  summarise(mean_Indentation = mean(Indentation, na.rm = TRUE))
```

```
## `summarise()` has grouped output by 'Tissue', 'Module'. You can override using
## the `.groups` argument.
```

```r
# Define fill colors for each factor level
fill_colors <- c("Inner testis" = viridis_pal(option = "D")(1),
                 "Tunica albuginea" = "#238A8DFF")

# Create the line plot
ggplot(data_summary, aes(x = Hz, y = mean_Indentation, linetype = Module, color = Tissue)) +
  geom_line(size = 1.5) +
  geom_point(size = 1) +
  scale_color_manual(values = fill_colors) +
  scale_linetype_manual(values = c("E1" = "solid", "E2" = "dashed"),
                        labels = c("E1" = "E'", "E2" = "E\"")) +
  scale_x_continuous(breaks = c(1, 5, 10, 20)) +
  scale_y_continuous(labels = scientific_format(), limits = c(0, 15500)) +
  labs(y = expression(paste("E'/E'' (Pa)")), x = "Frequency (Hz)", color = "Tissue", linetype = "Module
  theme_minimal() +
  theme(panel.grid.major = element_blank(),
        panel.grid.minor = element_blank(),
        panel.border = element_rect(colour = "black", fill = NA, size = 1),
        axis.text = element_text(size = 12))
```

```
## Warning: Using `size` aesthetic for lines was deprecated in ggplot2 3.4.0.
## i Please use `linewidth` instead.
## This warning is displayed once every 8 hours.
## Call `lifecycle::last_lifecycle_warnings()` to see where this warning was
## generated.

## Warning: The `size` argument of `element_rect()` is deprecated as of ggplot2 3.4.0.
## i Please use the `linewidth` argument instead.
## This warning is displayed once every 8 hours.
## Call `lifecycle::last_lifecycle_warnings()` to see where this warning was
## generated.
```

```r
# Save the plot
dev.off()
```

```
## pdf
##   2
```

```r
# Stats

df <- data

# Create a factor variable for the comparisons
df$Comparison <- factor(paste(df$Module, df$Tissue, sep = " - "))

# Perform pairwise comparisons using Tukey's method
model <- lm(Indentation ~ Comparison, data = df)
comp <- glht(model, linfct = mcp(Comparison = "Tukey"))

# Summarize the results
summary(comp)
```

```
##
##   Simultaneous Tests for General Linear Hypotheses
##
## Multiple Comparisons of Means: Tukey Contrasts
##
##
## Fit: lm(formula = Indentation ~ Comparison, data = df)
##
## Linear Hypotheses:
##                                                  Estimate Std. Error t value
## E1 - Tunica albuginea - E1 - Inner testis == 0       9753.0      617.5  15.794
## E2 - Inner testis - E1 - Inner testis == 0          -2319.2      605.0  -3.833
## E2 - Tunica albuginea - E1 - Inner testis == 0       1596.7      617.5   2.586
## E2 - Inner testis - E1 - Tunica albuginea == 0     -12072.3      617.5 -19.550
## E2 - Tunica albuginea - E1 - Tunica albuginea == 0  -8156.3      629.7 -12.952
## E2 - Tunica albuginea - E2 - Inner testis == 0       3916.0      617.5   6.342
##                                                  Pr(>|t|)
## E1 - Tunica albuginea - E1 - Inner testis == 0       <0.001 ***
## E2 - Inner testis - E1 - Inner testis == 0          <0.001 ***
## E2 - Tunica albuginea - E1 - Inner testis == 0       0.0507 .
## E2 - Inner testis - E1 - Tunica albuginea == 0       <0.001 ***
## E2 - Tunica albuginea - E1 - Tunica albuginea == 0   <0.001 ***
## E2 - Tunica albuginea - E2 - Inner testis == 0       <0.001 ***
## ---
## Signif. codes:  0 '***' 0.001 '**' 0.01 '*' 0.05 '.' 0.1 ' ' 1
## (Adjusted p values reported -- single-step method)
```

```r
################### YM ######################################

# Read the dataset from a CSV file
df <- read.csv("YM_male.csv")

# Summary data
summary_df <- df %>%
  group_by(Cellular, Tissue) %>%
  summarise(mean_YM = mean(YM),
            sd_YM = sd(YM))
```

```
## `summarise()` has grouped output by 'Cellular'. You can override using the
```

```
## `.groups` argument.
```

```r
# Plot YM

# Update the labels for Tissue
df$Tissue <- factor(df$Tissue, labels = c("Inner testis", "Tunica albuginea"))


png(filename="YM_male.png",
    width = 8.86, height = 6.86, units = "in",
    res = 600)

par(mgp=c(1.5,0.5,0),
    mar = c(4, #bottom
            4, #left
            3, #top
            2),
    cex.main = 0.8,
    family = "sans" #right
)

# Create a combined factor variable for Tissue and Cellular
df$Combined <- factor(paste(df$Tissue, df$Cellular, sep = " "),
                      levels = c("Inner testis Native", "Inner testis Decellularized",
                                 "Tunica albuginea Native", "Tunica albuginea Decellularized"))

# Define fill colors for each factor level
fill_colors <- c("Inner testis Native" = viridis_pal(option = "D")(1),
                 "Inner testis Decellularized" = alpha(viridis_pal(option = "D")(1), 0.2),
                 "Tunica albuginea Native" = "#238A8DFF",
                 "Tunica albuginea Decellularized" = alpha("#238A8DFF", 0.2))

# Create the plot using ggplot
ggplot(data = df, aes(x = Combined, y = YM, fill = Combined)) +
  geom_boxplot(width = 0.5, outlier.shape = NA) +
  geom_jitter(position = position_jitterdodge(), alpha = 1) +
  xlab("") +
  ylab("Young's modulus (Pa)") +
  ggtitle("") +
  scale_fill_manual(values = fill_colors, guide = "none") +
  theme_bw() +
  scale_x_discrete(labels = c("Inner testis\n Native", "Inner testis\n Decellularized",
                              "Tunica albuginea\n Native", "Tunica albuginea\n Decellularized")) +
  ylim(0, 30000) +
  theme(panel.grid.major = element_blank(), panel.grid.minor = element_blank(),
        axis.text.x = element_text(size = 14, angle = 0, hjust = 0.5, vjust = 0.5),
        axis.text.y = element_text(size = 12, angle = 0, hjust = 0.5),
        axis.title.x = element_blank(),
        axis.title.y = element_text(size = 16, face = "bold"),
        legend.position = "right",
        strip.text = element_text(size = 14, face = "bold"))  # Increase the size of facet labels
```

```
## Warning: Removed 1 rows containing non-finite values (`stat_boxplot()`).

## Warning: Removed 1 rows containing missing values (`geom_point()`).
```

```r
dev.off()
```

```
## pdf
##   2
```

```r
# Stats

# Create a factor variable for the comparisons
df$Comparison <- factor(paste(df$Tissue, df$Cellular, sep = " - "))

# Perform pairwise comparisons using Tukey's method
model <- lm(YM ~ Comparison, data = df)
comp <- glht(model, linfct = mcp(Comparison = "Tukey"))

# Summarize the results
summary(comp)
```

```
##
##   Simultaneous Tests for General Linear Hypotheses
##
## Multiple Comparisons of Means: Tukey Contrasts
##
##
## Fit: lm(formula = YM ~ Comparison, data = df)
##
## Linear Hypotheses:
##                                                                        Estimate
## Inner testis - Native - Inner testis - Decellularized == 0             -11434.4
## Tunica albuginea - Decellularized - Inner testis - Decellularized == 0   -115.8
## Tunica albuginea - Native - Inner testis - Decellularized == 0          -6951.4
## Tunica albuginea - Decellularized - Inner testis - Native == 0          11318.6
## Tunica albuginea - Native - Inner testis - Native == 0                   4482.9
## Tunica albuginea - Native - Tunica albuginea - Decellularized == 0      -6835.7
##                                                                        Std. Error
## Inner testis - Native - Inner testis - Decellularized == 0                 2537.4
## Tunica albuginea - Decellularized - Inner testis - Decellularized == 0     2645.4
## Tunica albuginea - Native - Inner testis - Decellularized == 0             2568.7
## Tunica albuginea - Decellularized - Inner testis - Native == 0             2244.7
## Tunica albuginea - Native - Inner testis - Native == 0                     2153.8
## Tunica albuginea - Native - Tunica albuginea - Decellularized == 0         2280.0
##                                                                        t value
## Inner testis - Native - Inner testis - Decellularized == 0              -4.506
## Tunica albuginea - Decellularized - Inner testis - Decellularized == 0  -0.044
## Tunica albuginea - Native - Inner testis - Decellularized == 0          -2.706
## Tunica albuginea - Decellularized - Inner testis - Native == 0           5.042
## Tunica albuginea - Native - Inner testis - Native == 0                   2.081
## Tunica albuginea - Native - Tunica albuginea - Decellularized == 0      -2.998
##                                                                        Pr(>|t|)
## Inner testis - Native - Inner testis - Decellularized == 0               <0.001
## Tunica albuginea - Decellularized - Inner testis - Decellularized == 0   1.0000
## Tunica albuginea - Native - Inner testis - Decellularized == 0           0.0451
## Tunica albuginea - Decellularized - Inner testis - Native == 0           <0.001
## Tunica albuginea - Native - Inner testis - Native == 0                   0.1737
## Tunica albuginea - Native - Tunica albuginea - Decellularized == 0       0.0218
##
```

```
## Inner testis - Native - Inner testis - Decellularized == 0              ***
## Tunica albuginea - Decellularized - Inner testis - Decellularized == 0
## Tunica albuginea - Native - Inner testis - Decellularized == 0         *
## Tunica albuginea - Decellularized - Inner testis - Native == 0         ***
## Tunica albuginea - Native - Inner testis - Native == 0
## Tunica albuginea - Native - Tunica albuginea - Decellularized == 0     *
## ---
## Signif. codes:  0 '***' 0.001 '**' 0.01 '*' 0.05 '.' 0.1 ' ' 1
## (Adjusted p values reported -- single-step method)
```

```r
################### Porosity #######################################

# Read the dataset from a CSV file
df <- read.csv("Porosity_male.csv")

# Summary data
summary_df <- df %>%
  group_by(Cellular, Tissue) %>%
  summarise(mean_Porosity = mean(Porosity),
            sd_Porosity = sd(Porosity))
```

```
## `summarise()` has grouped output by 'Cellular'. You can override using the
## `.groups` argument.
```

```r
# Update the labels for Tissue
df$Tissue <- factor(df$Tissue, labels = c("Inner testis", "Tunica albuginea"))

# Plot Porosity

png(filename="porosity_male.png",
    width = 8.86, height = 6.86, units = "in",
    res = 600)

par(mgp=c(1.5,0.5,0),
    mar = c(4, #bottom
            4, #left
            3, #top
            2),
    cex.main = 0.8,
    family = "sans" #right
)

# Create a combined factor variable for Tissue and Cellular
df$Combined <- factor(paste(df$Tissue, df$Cellular, sep = " "),
                      levels = c("Inner testis Native", "Inner testis Decellularized",
                                 "Tunica albuginea Native", "Tunica albuginea Decellularized"))

# Define fill colors for each factor level
fill_colors <- c("Inner testis Native" = viridis_pal(option = "D")(1),
                 "Inner testis Decellularized" = alpha(viridis_pal(option = "D")(1), 0.2),
                 "Tunica albuginea Native" = "#238A8DFF",
                 "Tunica albuginea Decellularized" = alpha("#238A8DFF", 0.2))

# Create the plot using ggplot
ggplot(data = df, aes(x = Combined, y = Porosity, fill = Combined)) +
```

```r
  geom_boxplot(width = 0.5, outlier.shape = NA) +
  geom_jitter(position = position_jitterdodge(), alpha = 1) +
  xlab("") +
  ylab("Estimated Porosity (%)") +
  ggtitle("") +
  scale_fill_manual(values = fill_colors, guide = "none") +
  theme_bw() +
  scale_x_discrete(labels = c("Inner testis\n Native", "Inner testis\n Decellularized",
                              "Tunica albuginea\n Native", "Tunica albuginea\n Decellularized")) +
  ylim(0, 130) +
  theme(panel.grid.major = element_blank(), panel.grid.minor = element_blank(),
        axis.text.x = element_text(size = 14, angle = 0, hjust = 0.5, vjust = 0.5),
        axis.text.y = element_text(size = 12, angle = 0, hjust = 0.5),
        axis.title.x = element_blank(),
        axis.title.y = element_text(size = 16, face = "bold"),
        legend.position = "right",
        strip.text = element_text(size = 14, face = "bold"))  # Increase the size of facet labels

dev.off()
```

```
## pdf
##   2
```

```r
# Stats

# Read the dataset from a CSV file
df <- read.csv("Porosity_male.csv")

# Create a factor variable for the comparisons
df$Comparison <- factor(paste(df$Tissue, df$Cellular, sep = " - "))

# Perform pairwise comparisons using Tukey's method
model <- lm(Porosity ~ Comparison, data = df)
comp <- glht(model, linfct = mcp(Comparison = "Tukey"))

# Summarize the results
summary(comp)
```

```
##
##   Simultaneous Tests for General Linear Hypotheses
##
## Multiple Comparisons of Means: Tukey Contrasts
##
##
## Fit: lm(formula = Porosity ~ Comparison, data = df)
##
## Linear Hypotheses:
##                                                          Estimate Std. Error
## Testis - Native - Testis - Decellularized == 0             34.419     10.547
## Tunica - Decellularized - Testis - Decellularized == 0      9.684     12.374
## Tunica - Native - Testis - Decellularized == 0             42.593     10.547
## Tunica - Decellularized - Testis - Native == 0            -24.735     12.107
## Tunica - Native - Testis - Native == 0                      8.174     10.232
## Tunica - Native - Tunica - Decellularized == 0             32.909     12.107
##                                                          t value Pr(>|t|)
```

```
## Testis - Native - Testis - Decellularized == 0          3.263  0.01500 *
## Tunica - Decellularized - Testis - Decellularized == 0   0.783  0.86088
## Tunica - Native - Testis - Decellularized == 0          4.038  0.00217 **
## Tunica - Decellularized - Testis - Native == 0         -2.043  0.19653
## Tunica - Native - Testis - Native == 0                  0.799  0.85351
## Tunica - Native - Tunica - Decellularized == 0          2.718  0.05153 .
## ---
## Signif. codes:  0 '***' 0.001 '**' 0.01 '*' 0.05 '.' 0.1 ' ' 1
## (Adjusted p values reported -- single-step method)
```

```
################### YM-Porosity correlation ########################################


# Read the dataset from a CSV file
df <- read.csv("YM_male.csv")

# Update the labels for Tissue
df$Tissue <- factor(df$Tissue, labels = c("Inner testis", "Tunica albuginea"))


png(filename="correlation_male.png",
    width = 15, height = 10, units = "in",
    res = 600)

par(mgp=c(1.5,0.5,0),
    mar = c(4, #bottom
            4, #left
            3, #top
            2),
    cex.main = 0.8,
    family = "sans" #right
)


# Create a scatter plot for Inner testis and Decellularized
p_inner_testis_decell <- ggplot(df, aes(x = Porosity, y = YM, color = Cellular)) +
  geom_point(data = subset(df, Tissue == "Inner testis" & Cellular == "Decellularized"), show.legend = F
  geom_smooth(data = subset(df, Tissue == "Inner testis" & Cellular == "Decellularized"), method = "lm"
  labs(x = "Porosity (%)", y = "Young's Modulus (Pa)") +
  ggtitle("Inner Testis - Decellularized") +
  stat_cor(data = subset(df, Tissue == "Inner testis" & Cellular == "Decellularized"), method = "pearso
  theme_bw() +
  theme(panel.grid.major = element_blank(), panel.grid.minor = element_blank(),
        legend.position = "none",
        axis.title.x = element_text(size = 16),
        axis.title.y = element_text(size = 16))

# Create a scatter plot for Inner testis and Native
p_inner_testis_native <- ggplot(df, aes(x = Porosity, y = YM, color = Cellular)) +
  geom_point(data = subset(df, Tissue == "Inner testis" & Cellular == "Native"), show.legend = FALSE) +
  geom_smooth(data = subset(df, Tissue == "Inner testis" & Cellular == "Native"), method = "lm", se = F
  labs(x = "Porosity (%)", y = "Young's Modulus (Pa)") +
  ggtitle("Inner Testis - Native") +
  stat_cor(data = subset(df, Tissue == "Inner testis" & Cellular == "Native"), method = "pearson", label
```

```
  theme_bw() +
  theme(panel.grid.major = element_blank(), panel.grid.minor = element_blank(),
        legend.position = "none",
        axis.title.x = element_text(size = 16),
        axis.title.y = element_text(size = 16))

# Create a scatter plot for Tunica albuginea and Decellularized
p_tunica_decell <- ggplot(df, aes(x = Porosity, y = YM, color = Cellular)) +
  geom_point(data = subset(df, Tissue == "Tunica albuginea" & Cellular == "Decellularized"), show.legend
  geom_smooth(data = subset(df, Tissue == "Tunica albuginea" & Cellular == "Decellularized"), method =
  labs(x = "Porosity (%)", y = "Young's Modulus (Pa)") +
  ggtitle("Tunica Albuginea - Decellularized") +
  stat_cor(data = subset(df, Tissue == "Tunica albuginea" & Cellular == "Decellularized"), method = "pe
  theme_bw() +
  theme(panel.grid.major = element_blank(), panel.grid.minor = element_blank(),
        legend.position = "none",
        axis.title.x = element_text(size = 16),
        axis.title.y = element_text(size = 16))

# Create a scatter plot for Tunica albuginea and Native
p_tunica_native <- ggplot(df, aes(x = Porosity, y = YM, color = Cellular)) +
  geom_point(data = subset(df, Tissue == "Tunica albuginea" & Cellular == "Native"), show.legend = FALS
  geom_smooth(data = subset(df, Tissue == "Tunica albuginea" & Cellular == "Native"), method = "lm", se
  labs(x = "Porosity (%)", y = "Young's Modulus (Pa)") +
  ggtitle("Tunica Albuginea - Native") +
  stat_cor(data = subset(df, Tissue == "Tunica albuginea" & Cellular == "Native"), method = "pearson", 
  theme_bw() +
  theme(panel.grid.major = element_blank(), panel.grid.minor = element_blank(),
        legend.position = "none",
        axis.title.x = element_text(size = 16),
        axis.title.y = element_text(size = 16))

# Combine the plots
p_combined <- cowplot::plot_grid(
  cowplot::plot_grid(p_inner_testis_decell, p_inner_testis_native, nrow = 1),
  cowplot::plot_grid(p_tunica_decell, p_tunica_native, nrow = 1),
  nrow = 2
)
```

```
## `geom_smooth()` using formula = 'y ~ x'

## `geom_smooth()` using formula = 'y ~ x'
## `geom_smooth()` using formula = 'y ~ x'
## `geom_smooth()` using formula = 'y ~ x'
```

```
# Display the combined plot
p_combined

dev.off()
```

```
## pdf
##   2
```

```
#################################################################
#################### Endometrium Analysis #######################
#################################################################
```

```
################### E' E" ########################################


#data input
data <- read.csv("Endo_rheo.csv")

# Plot

png(filename="E1_E2_endo.png",
    width = 9.86, height = 6.86, units = "in",
    res = 600)

par(mgp=c(1.5,0.5,0),
    mar = c(4, #bottom
            4, #left
            3, #top
            2),
    cex.main = 0.8,
    family = "sans" #right
)

# Calculate means and standard deviations
data_summary <- data %>%
  group_by(Module, Estrus_cycle, Position, Hz) %>%
  summarise(mean_Indentation = mean(Indentation, na.rm = TRUE),
            sd_Indentation = sd(Indentation, na.rm = TRUE))
```

```
## `summarise()` has grouped output by 'Module', 'Estrus_cycle', 'Position'. You
## can override using the `.groups` argument.
```

```
# Update the labels for Position, Estrus_cycle
data_summary$Position <- factor(data_summary$Position, labels = c("Contralateral", "Ipsilateral"))
data_summary$Estrus_cycle <- factor(data_summary$Estrus_cycle, labels = c("Follicular phase", "Luteal pl

# Create a new variable for line type
data_summary <- data_summary %>%
  mutate(LineType = ifelse(Module == "E1", "E1", "E2"))

# Define fill colors for each factor level
fill_colors <- c("Follicular phase Contralateral" = viridis_pal(option = "D")(1),
                 "Follicular phase Ipsilateral" = alpha(viridis_pal(option = "D")(1), 0.2),
                 "Luteal phase Contralateral" = "#238A8DFF",
                 "Luteal phase Ipsilateral" = alpha("#238A8DFF", 0.2))

# Create the line plot with separate lines for each Estrus_cycle and Position
ggplot(data_summary, aes(x = Hz, y = mean_Indentation, group = interaction(Module, Estrus_cycle, Positi
                         color = paste(Estrus_cycle, Position), linetype = LineType)) +
  geom_line(size = 1.5) +  # Adjust line thickness
  geom_point(size = 1) +  # Add dots at each Hz point
  scale_color_manual(values = fill_colors) +  # Use custom fill colors
  scale_linetype_manual(values = c("E1" = "solid", "E2" = "dashed"),
                        labels = c("E1" = "E'", "E2" = "E\"")) +
  scale_x_continuous(breaks = c(1, 5, 10, 20)) +
  scale_y_continuous(labels = scientific_format(), limits = c(0, 10000)) +  # Adjust Y-axis limits
```

```r
  labs(y = expression(paste("E'/E'' (Pa)")), x = "Frequency (Hz)", color = "Estrus Cycle + Position") +
  theme_minimal() +
  theme(panel.grid.major = element_blank(),
        panel.grid.minor = element_blank(),
        panel.border = element_rect(colour = "black", fill = NA, size = 1),
        axis.text = element_text(size = 12))  # Increase font size of axis labels

dev.off()
```

```
## pdf
##   2
```

```r
# Stats

df <- data

# Create a factor variable for the comparisons
df$Comparison <- factor(paste(df$Estrus_cycle, df$Position, df$Module, sep = " - "))

# Perform pairwise comparisons using Tukey's method
model <- lm(Indentation ~ Comparison, data = df)
comp <- glht(model, linfct = mcp(Comparison = "Tukey"))

# Summarize the results
summary(comp)
```

```
## Warning in RET$pfunction("adjusted", ...): Completion with error > abseps

##
##   Simultaneous Tests for General Linear Hypotheses
##
## Multiple Comparisons of Means: Tukey Contrasts
##
##
## Fit: lm(formula = Indentation ~ Comparison, data = df)
##
## Linear Hypotheses:
##                                              Estimate Std. Error t value Pr(>|t|)
## Fol - Contra - E2 - Fol - Contra - E1 == 0   -3660.3      761.2  -4.808  < 0.001
## Fol - Ipsi - E1 - Fol - Contra - E1 == 0      -120.8      798.4  -0.151  1.00000
## Fol - Ipsi - E2 - Fol - Contra - E1 == 0     -2390.9      798.4  -2.995  0.05538
## Lut - Contra - E1 - Fol - Contra - E1 == 0    1699.7      648.8   2.620  0.14690
## Lut - Contra - E2 - Fol - Contra - E1 == 0   -2634.2      652.8  -4.035  0.00154
## Lut - Ipsi - E1 - Fol - Contra - E1 == 0      1380.8      687.6   2.008  0.46966
## Lut - Ipsi - E2 - Fol - Contra - E1 == 0     -2520.9      687.6  -3.666  0.00614
## Fol - Ipsi - E1 - Fol - Contra - E2 == 0      3539.5      798.4   4.433  < 0.001
## Fol - Ipsi - E2 - Fol - Contra - E2 == 0      1269.4      798.4   1.590  0.75039
## Lut - Contra - E1 - Fol - Contra - E2 == 0    5359.9      648.8   8.261  < 0.001
## Lut - Contra - E2 - Fol - Contra - E2 == 0    1026.1      652.8   1.572  0.76136
## Lut - Ipsi - E1 - Fol - Contra - E2 == 0      5041.0      687.6   7.332  < 0.001
## Lut - Ipsi - E2 - Fol - Contra - E2 == 0      1139.3      687.6   1.657  0.70861
## Fol - Ipsi - E2 - Fol - Ipsi - E1 == 0       -2270.1      833.9  -2.722  0.11422
## Lut - Contra - E1 - Fol - Ipsi - E1 == 0      1820.4      692.0   2.631  0.14358
## Lut - Contra - E2 - Fol - Ipsi - E1 == 0     -2513.4      695.7  -3.613  0.00758
## Lut - Ipsi - E1 - Fol - Ipsi - E1 == 0        1501.5      728.5   2.061  0.43390
```

```
## Lut - Ipsi - E2 - Fol - Ipsi - E1 == 0      -2400.2      728.5  -3.295  0.02275
## Lut - Contra - E1 - Fol - Ipsi - E2 == 0     4090.5      692.0   5.911  < 0.001
## Lut - Contra - E2 - Fol - Ipsi - E2 == 0     -243.3      695.7  -0.350  0.99997
## Lut - Ipsi - E1 - Fol - Ipsi - E2 == 0       3771.6      728.5   5.177  < 0.001
## Lut - Ipsi - E2 - Fol - Ipsi - E2 == 0       -130.1      728.5  -0.179  1.00000
## Lut - Contra - E2 - Lut - Contra - E1 == 0  -4333.8      517.3  -8.378  < 0.001
## Lut - Ipsi - E1 - Lut - Contra - E1 == 0     -318.9      560.5  -0.569  0.99917
## Lut - Ipsi - E2 - Lut - Contra - E1 == 0    -4220.6      560.5  -7.530  < 0.001
## Lut - Ipsi - E1 - Lut - Contra - E2 == 0     4014.9      565.1   7.105  < 0.001
## Lut - Ipsi - E2 - Lut - Contra - E2 == 0      113.2      565.1   0.200  1.00000
## Lut - Ipsi - E2 - Lut - Ipsi - E1 == 0      -3901.7      605.0  -6.449  < 0.001
##
## Fol - Contra - E2 - Fol - Contra - E1 == 0 ***
## Fol - Ipsi - E1 - Fol - Contra - E1 == 0
## Fol - Ipsi - E2 - Fol - Contra - E1 == 0   .
## Lut - Contra - E1 - Fol - Contra - E1 == 0
## Lut - Contra - E2 - Fol - Contra - E1 == 0 **
## Lut - Ipsi - E1 - Fol - Contra - E1 == 0
## Lut - Ipsi - E2 - Fol - Contra - E1 == 0   **
## Fol - Ipsi - E1 - Fol - Contra - E2 == 0   ***
## Fol - Ipsi - E2 - Fol - Contra - E2 == 0
## Lut - Contra - E1 - Fol - Contra - E2 == 0 ***
## Lut - Contra - E2 - Fol - Contra - E2 == 0
## Lut - Ipsi - E1 - Fol - Contra - E2 == 0   ***
## Lut - Ipsi - E2 - Fol - Contra - E2 == 0
## Fol - Ipsi - E2 - Fol - Ipsi - E1 == 0
## Lut - Contra - E1 - Fol - Ipsi - E1 == 0
## Lut - Contra - E2 - Fol - Ipsi - E1 == 0   **
## Lut - Ipsi - E1 - Fol - Ipsi - E1 == 0
## Lut - Ipsi - E2 - Fol - Ipsi - E1 == 0     *
## Lut - Contra - E1 - Fol - Ipsi - E2 == 0   ***
## Lut - Contra - E2 - Fol - Ipsi - E2 == 0
## Lut - Ipsi - E1 - Fol - Ipsi - E2 == 0     ***
## Lut - Ipsi - E2 - Fol - Ipsi - E2 == 0
## Lut - Contra - E2 - Lut - Contra - E1 == 0 ***
## Lut - Ipsi - E1 - Lut - Contra - E1 == 0
## Lut - Ipsi - E2 - Lut - Contra - E1 == 0   ***
## Lut - Ipsi - E1 - Lut - Contra - E2 == 0   ***
## Lut - Ipsi - E2 - Lut - Contra - E2 == 0
## Lut - Ipsi - E2 - Lut - Ipsi - E1 == 0     ***
## ---
## Signif. codes:  0 '***' 0.001 '**' 0.01 '*' 0.05 '.' 0.1 ' ' 1
## (Adjusted p values reported -- single-step method)
```

```r
################### YM ######################################

# Read the dataset from a CSV file
df <- read.csv("Endometrium_YM3.csv")

# Summary data
summary_df <- df %>%
  group_by(Cellular, Tissue, Position, Estrus_cycle) %>%
  summarise(mean_YM = mean(YM),
            sd_YM = sd(YM))
```

```
## `summarise()` has grouped output by 'Cellular', 'Tissue', 'Position'. You can
## override using the `.groups` argument.
```

```r
# Plot


png(filename="Endometirum_YM.png",
    width = 10, height = 6.86, units = "in",
    res = 600)

par(mgp=c(1.5,0.5,0),
    mar = c(4, #bottom
            4, #left
            3, #top
            2),
    cex.main = 0.8,
    family = "sans" #right
)

# Read the dataset from a CSV file
df <- read.csv("Endometrium_YM3.csv")

# Update the labels for Position, Estrus_cycle
df$Position <- factor(df$Position, labels = c("Contralateral", "Ipsilateral"))
df$Estrus_cycle <- factor(df$Estrus_cycle, labels = c("Follicular phase", "Luteal phase"))

# Convert Animal_ID to a factor
df$Animal_ID <- as.factor(df$Animal_ID)

# Create a combined factor variable for Estrus_cycle and Cellular
df$Combined <- factor(paste(df$Estrus_cycle, df$Cellular, sep = " "),
                      levels = c("Follicular phase Native", "Follicular phase Decellularized",
                                 "Luteal phase Native", "Luteal phase Decellularized"))

# Define fill colors for each factor level
fill_colors <- c("Follicular phase Native" = viridis_pal(option = "D")(1),
                 "Follicular phase Decellularized" = alpha(viridis_pal(option = "D")(1), 0.2),
                 "Luteal phase Native" = "#238A8DFF",
                 "Luteal phase Decellularized" = alpha("#238A8DFF", 0.2))

# Create the plot using ggplot
ggplot(data = df, aes(x = Combined, y = YM, fill = Combined)) +
  geom_boxplot(width = 0.5, outlier.shape = NA) +
  geom_jitter(position = position_jitterdodge(), alpha = 1) +
  xlab("") +
  ylab("Young's modulus (Pa)") +
  ggtitle("Boxplot and Individual Data Points of YM by Estrus_cycle, Cellular, and Position") +
  scale_fill_manual(values = fill_colors, guide = "none") +
  theme_bw() +
  scale_x_discrete(labels = c("Follicular phase\nNative", "Follicular phase\nDecellularized",
                              "Luteal phase\nNative", "Luteal phase\nDecellularized")) +
  ylim(0, 30000) +
  facet_grid(. ~ Position) +
  theme(panel.grid.major = element_blank(), panel.grid.minor = element_blank(),
```

```r
        axis.text.x = element_text(size = 11, angle = 0, hjust = 0.5),
        axis.text.y = element_text(size = 12, angle = 0, hjust = 0.5),
        axis.title.x = element_text(size = 18, face = "bold"),
        axis.title.y = element_text(size = 16, face = "bold"),
        legend.position = "right",
        strip.text = element_text(size = 14, face = "bold"))  # Increase the size of facet labels
```

```
## Warning: Removed 2 rows containing non-finite values (`stat_boxplot()`).
```

```
## Warning: Removed 2 rows containing missing values (`geom_point()`).
```

```r
dev.off()
```

```
## pdf
##   2
```

```r
# Stats

# Read the dataset from a CSV file
df <- read.csv("Endometrium_YM.csv")

# Create a factor variable for the comparisons
df$Comparison <- factor(paste(df$Estrus_cycle, df$Position, df$Cellular, sep = " - "))

# Perform pairwise comparisons using Tukey's method
model <- lm(YM ~ Comparison, data = df)
comp <- glht(model, linfct = mcp(Comparison = "Tukey"))

# Summarize the results
summary(comp)
```

```
##
##   Simultaneous Tests for General Linear Hypotheses
##
## Multiple Comparisons of Means: Tukey Contrasts
##
##
## Fit: lm(formula = YM ~ Comparison, data = df)
##
## Linear Hypotheses:
##                                                                  Estimate
## Fol - Contra - Native - Fol - Contra - Decellularized == 0         -4765.0
## Fol - Ipsi - Decellularized - Fol - Contra - Decellularized == 0   -3293.3
## Fol - Ipsi - Native - Fol - Contra - Decellularized == 0           -5380.8
## Lut - Contra - Decellularized - Fol - Contra - Decellularized == 0  1728.1
## Lut - Contra - Native - Fol - Contra - Decellularized == 0         -4211.3
## Lut - Ipsi - Decellularized - Fol - Contra - Decellularized == 0    8738.8
## Lut - Ipsi - Native - Fol - Contra - Decellularized == 0           -5005.5
## Fol - Ipsi - Decellularized - Fol - Contra - Native == 0            1471.7
## Fol - Ipsi - Native - Fol - Contra - Native == 0                    -615.8
## Lut - Contra - Decellularized - Fol - Contra - Native == 0          6493.1
## Lut - Contra - Native - Fol - Contra - Native == 0                   553.7
## Lut - Ipsi - Decellularized - Fol - Contra - Native == 0           13503.8
## Lut - Ipsi - Native - Fol - Contra - Native == 0                    -240.5
## Fol - Ipsi - Native - Fol - Ipsi - Decellularized == 0             -2087.5
## Lut - Contra - Decellularized - Fol - Ipsi - Decellularized == 0    5021.4
```

```
## Lut - Contra - Native - Fol - Ipsi - Decellularized == 0            -918.1
## Lut - Ipsi - Decellularized - Fol - Ipsi - Decellularized == 0     12032.1
## Lut - Ipsi - Native - Fol - Ipsi - Decellularized == 0            -1712.2
## Lut - Contra - Decellularized - Fol - Ipsi - Native == 0            7108.9
## Lut - Contra - Native - Fol - Ipsi - Native == 0                    1169.4
## Lut - Ipsi - Decellularized - Fol - Ipsi - Native == 0            14119.6
## Lut - Ipsi - Native - Fol - Ipsi - Native == 0                       375.3
## Lut - Contra - Native - Lut - Contra - Decellularized == 0         -5939.4
## Lut - Ipsi - Decellularized - Lut - Contra - Decellularized == 0    7010.7
## Lut - Ipsi - Native - Lut - Contra - Decellularized == 0           -6733.6
## Lut - Ipsi - Decellularized - Lut - Contra - Native == 0           12950.2
## Lut - Ipsi - Native - Lut - Contra - Native == 0                    -794.1
## Lut - Ipsi - Native - Lut - Ipsi - Decellularized == 0            -13744.3
##                                                                  Std. Error
## Fol - Contra - Native - Fol - Contra - Decellularized == 0            1644.1
## Fol - Ipsi - Decellularized - Fol - Contra - Decellularized == 0      1574.1
## Fol - Ipsi - Native - Fol - Contra - Decellularized == 0              1706.2
## Lut - Contra - Decellularized - Fol - Contra - Decellularized == 0    1787.3
## Lut - Contra - Native - Fol - Contra - Decellularized == 0            1459.9
## Lut - Ipsi - Decellularized - Fol - Contra - Decellularized == 0      1706.2
## Lut - Ipsi - Native - Fol - Contra - Decellularized == 0              1555.2
## Fol - Ipsi - Decellularized - Fol - Contra - Native == 0              1574.1
## Fol - Ipsi - Native - Fol - Contra - Native == 0                      1706.2
## Lut - Contra - Decellularized - Fol - Contra - Native == 0            1787.3
## Lut - Contra - Native - Fol - Contra - Native == 0                    1459.9
## Lut - Ipsi - Decellularized - Fol - Contra - Native == 0              1706.2
## Lut - Ipsi - Native - Fol - Contra - Native == 0                      1555.2
## Fol - Ipsi - Native - Fol - Ipsi - Decellularized == 0                1638.8
## Lut - Contra - Decellularized - Fol - Ipsi - Decellularized == 0      1723.2
## Lut - Contra - Native - Fol - Ipsi - Decellularized == 0              1380.6
## Lut - Ipsi - Decellularized - Fol - Ipsi - Decellularized == 0        1638.8
## Lut - Ipsi - Native - Fol - Ipsi - Decellularized == 0                1481.0
## Lut - Contra - Decellularized - Fol - Ipsi - Native == 0              1844.6
## Lut - Contra - Native - Fol - Ipsi - Native == 0                      1529.4
## Lut - Ipsi - Decellularized - Fol - Ipsi - Native == 0                1766.0
## Lut - Ipsi - Native - Fol - Ipsi - Native == 0                        1620.6
## Lut - Contra - Native - Lut - Contra - Decellularized == 0            1619.5
## Lut - Ipsi - Decellularized - Lut - Contra - Decellularized == 0      1844.6
## Lut - Ipsi - Native - Lut - Contra - Decellularized == 0              1705.9
## Lut - Ipsi - Decellularized - Lut - Contra - Native == 0              1529.4
## Lut - Ipsi - Native - Lut - Contra - Native == 0                      1358.9
## Lut - Ipsi - Native - Lut - Ipsi - Decellularized == 0                1620.6
##                                                                  t value
## Fol - Contra - Native - Fol - Contra - Decellularized == 0         -2.898
## Fol - Ipsi - Decellularized - Fol - Contra - Decellularized == 0   -2.092
## Fol - Ipsi - Native - Fol - Contra - Decellularized == 0           -3.154
## Lut - Contra - Decellularized - Fol - Contra - Decellularized == 0   0.967
## Lut - Contra - Native - Fol - Contra - Decellularized == 0         -2.885
## Lut - Ipsi - Decellularized - Fol - Contra - Decellularized == 0    5.122
## Lut - Ipsi - Native - Fol - Contra - Decellularized == 0           -3.219
## Fol - Ipsi - Decellularized - Fol - Contra - Native == 0            0.935
## Fol - Ipsi - Native - Fol - Contra - Native == 0                   -0.361
## Lut - Contra - Decellularized - Fol - Contra - Native == 0          3.633
## Lut - Contra - Native - Fol - Contra - Native == 0                  0.379
```

```
## Lut - Ipsi - Decellularized - Fol - Contra - Native == 0           7.915
## Lut - Ipsi - Native - Fol - Contra - Native == 0                  -0.155
## Fol - Ipsi - Native - Fol - Ipsi - Decellularized == 0            -1.274
## Lut - Contra - Decellularized - Fol - Ipsi - Decellularized == 0   2.914
## Lut - Contra - Native - Fol - Ipsi - Decellularized == 0          -0.665
## Lut - Ipsi - Decellularized - Fol - Ipsi - Decellularized == 0     7.342
## Lut - Ipsi - Native - Fol - Ipsi - Decellularized == 0            -1.156
## Lut - Contra - Decellularized - Fol - Ipsi - Native == 0           3.854
## Lut - Contra - Native - Fol - Ipsi - Native == 0                   0.765
## Lut - Ipsi - Decellularized - Fol - Ipsi - Native == 0             7.995
## Lut - Ipsi - Native - Fol - Ipsi - Native == 0                     0.232
## Lut - Contra - Native - Lut - Contra - Decellularized == 0        -3.667
## Lut - Ipsi - Decellularized - Lut - Contra - Decellularized == 0   3.801
## Lut - Ipsi - Native - Lut - Contra - Decellularized == 0          -3.947
## Lut - Ipsi - Decellularized - Lut - Contra - Native == 0           8.467
## Lut - Ipsi - Native - Lut - Contra - Native == 0                  -0.584
## Lut - Ipsi - Native - Lut - Ipsi - Decellularized == 0            -8.481
##                                                                  Pr(>|t|)
## Fol - Contra - Native - Fol - Contra - Decellularized == 0         0.08079 .
## Fol - Ipsi - Decellularized - Fol - Contra - Decellularized == 0   0.42243
## Fol - Ipsi - Native - Fol - Contra - Decellularized == 0           0.04104 *
## Lut - Contra - Decellularized - Fol - Contra - Decellularized == 0 0.97779
## Lut - Contra - Native - Fol - Contra - Decellularized == 0         0.08373 .
## Lut - Ipsi - Decellularized - Fol - Contra - Decellularized == 0   < 0.001 ***
## Lut - Ipsi - Native - Fol - Contra - Decellularized == 0           0.03377 *
## Fol - Ipsi - Decellularized - Fol - Contra - Native == 0           0.98165
## Fol - Ipsi - Native - Fol - Contra - Native == 0                   0.99996
## Lut - Contra - Decellularized - Fol - Contra - Native == 0         0.00919 **
## Lut - Contra - Native - Fol - Contra - Native == 0                 0.99994
## Lut - Ipsi - Decellularized - Fol - Contra - Native == 0           < 0.001 ***
## Lut - Ipsi - Native - Fol - Contra - Native == 0                   1.00000
## Fol - Ipsi - Native - Fol - Ipsi - Decellularized == 0             0.90534
## Lut - Contra - Decellularized - Fol - Ipsi - Decellularized == 0   0.07764 .
## Lut - Contra - Native - Fol - Ipsi - Decellularized == 0           0.99768
## Lut - Ipsi - Decellularized - Fol - Ipsi - Decellularized == 0     < 0.001 ***
## Lut - Ipsi - Native - Fol - Ipsi - Decellularized == 0             0.94161
## Lut - Contra - Decellularized - Fol - Ipsi - Native == 0           0.00456 **
## Lut - Contra - Native - Fol - Ipsi - Native == 0                   0.99445
## Lut - Ipsi - Decellularized - Fol - Ipsi - Native == 0             < 0.001 ***
## Lut - Ipsi - Native - Fol - Ipsi - Native == 0                     1.00000
## Lut - Contra - Native - Lut - Contra - Decellularized == 0         0.00838 **
## Lut - Ipsi - Decellularized - Lut - Contra - Decellularized == 0   0.00542 **
## Lut - Ipsi - Native - Lut - Contra - Decellularized == 0           0.00325 **
## Lut - Ipsi - Decellularized - Lut - Contra - Native == 0           < 0.001 ***
## Lut - Ipsi - Native - Lut - Contra - Native == 0                   0.99899
## Lut - Ipsi - Native - Lut - Ipsi - Decellularized == 0             < 0.001 ***
## ---
## Signif. codes:  0 '***' 0.001 '**' 0.01 '*' 0.05 '.' 0.1 ' ' 1
## (Adjusted p values reported -- single-step method)
```

```
################### Porosity #######################################


# Read the dataset from a CSV file
```

```r
df <- read.csv("Endometrium_porosity.csv")

# Summary data

summary_df <- df %>%
  group_by(Cellular, Position, Estrus_cycle) %>%
  summarise(mean_Porosity = mean(Porosity),
            sd_Porosity = sd(Porosity))
```

```
## `summarise()` has grouped output by 'Cellular', 'Position'. You can override
## using the `.groups` argument.
```

```r
#Plot

png(filename="Endometirum_porosity.png",
    width = 10, height = 6.86, units = "in",
    res = 600)

par(mgp=c(1.5,0.5,0),
    mar = c(4, #bottom
            4, #left
            3, #top
            2),
    cex.main = 0.8,
    family = "sans" #right
)

# Update the labels for Position and Estrus_cycle
df$Position <- factor(df$Position, labels = c("Contralateral", "Ipsilateral"))
df$Estrus_cycle <- factor(df$Estrus_cycle, labels = c("Follicular phase", "Luteal phase"))

# Convert Animal_ID to a factor
df$Animal_ID <- as.factor(df$Animal_ID)

# Create a combined factor variable for Estrus_cycle and Cellular
df$Combined <- factor(paste(df$Estrus_cycle, df$Cellular, sep = " "),
                      levels = c("Follicular phase Native", "Follicular phase Decellularized",
                                 "Luteal phase Native", "Luteal phase Decellularized"))

# Define fill colors for each factor level
fill_colors <- c("Follicular phase Native" = viridis_pal(option = "D")(1),
                 "Follicular phase Decellularized" = alpha(viridis_pal(option = "D")(1), 0.2),
                 "Luteal phase Native" = "#238A8DFF",
                 "Luteal phase Decellularized" = alpha("#238A8DFF", 0.2))

# Create the plot using ggplot
ggplot(data = df, aes(x = Combined, y = Porosity, fill = Combined)) +
  geom_boxplot(width = 0.5, outlier.shape = NA) +
  geom_jitter(position = position_jitterdodge(), alpha = 1) +
  xlab("") +
  ylab("Estimated Porosity (%)") +
  ggtitle("Boxplot and Individual Data Points of Porosity by Estrus_cycle, Cellular, and Position") +
  scale_fill_manual(values = fill_colors, guide = "none") +
  theme_bw() +
```

```r
  scale_x_discrete(labels = c("Follicular phase\nNative", "Follicular phase\nDecellularized",
                              "Luteal phase\nNative", "Luteal phase\nDecellularized")) +
  ylim(0, 130) +
  facet_grid(. ~ Position) +
  theme(panel.grid.major = element_blank(), panel.grid.minor = element_blank(),
        axis.text.x = element_text(size = 11, angle = 0, hjust = 0.5),
        axis.text.y = element_text(size = 12, angle = 0, hjust = 0.5),
        axis.title.x = element_text(size = 18, face = "bold"),
        axis.title.y = element_text(size = 16, face = "bold"),
        legend.position = "right",
        strip.text = element_text(size = 14, face = "bold"))  # Increase the size of facet labels

dev.off()
```

```
## pdf
##   2
```

```r
# Stats

# Read the dataset from a CSV file
df <- read.csv("Endometrium_porosity.csv")

# Create a factor variable for the comparisons
df$Comparison <- factor(paste(df$Estrus_cycle, df$Position, df$Cellular, sep = " - "))

# Perform pairwise comparisons using Tukey's method
model <- lm(Porosity ~ Comparison, data = df)
comp <- glht(model, linfct = mcp(Comparison = "Tukey"))

# Summarize the results
summary(comp)
```

```
##
##   Simultaneous Tests for General Linear Hypotheses
##
## Multiple Comparisons of Means: Tukey Contrasts
##
##
## Fit: lm(formula = Porosity ~ Comparison, data = df)
##
## Linear Hypotheses:
##                                                                Estimate
## Fol - Contra - Native - Fol - Contra - Decellularized == 0      31.9042
## Fol - Ipsi - Decellularized - Fol - Contra - Decellularized == 0    -3.3390
## Fol - Ipsi - Native - Fol - Contra - Decellularized == 0        24.5681
## Lut - Contra - Decellularized - Fol - Contra - Decellularized == 0  14.2973
## Lut - Contra - Native - Fol - Contra - Decellularized == 0      25.0808
## Lut - Ipsi - Decellularized - Fol - Contra - Decellularized == 0    10.9570
## Lut - Ipsi - Native - Fol - Contra - Decellularized == 0        33.3700
## Fol - Ipsi - Decellularized - Fol - Contra - Native == 0       -35.2432
## Fol - Ipsi - Native - Fol - Contra - Native == 0                -7.3360
## Lut - Contra - Decellularized - Fol - Contra - Native == 0     -17.6069
## Lut - Contra - Native - Fol - Contra - Native == 0              -6.8234
## Lut - Ipsi - Decellularized - Fol - Contra - Native == 0       -20.9471
## Lut - Ipsi - Native - Fol - Contra - Native == 0                 1.4658
```

```
## Fol - Ipsi - Native - Fol - Ipsi - Decellularized == 0            27.9071
## Lut - Contra - Decellularized - Fol - Ipsi - Decellularized == 0    17.6363
## Lut - Contra - Native - Fol - Ipsi - Decellularized == 0            28.4198
## Lut - Ipsi - Decellularized - Fol - Ipsi - Decellularized == 0      14.2960
## Lut - Ipsi - Native - Fol - Ipsi - Decellularized == 0              36.7090
## Lut - Contra - Decellularized - Fol - Ipsi - Native == 0           -10.2709
## Lut - Contra - Native - Fol - Ipsi - Native == 0                     0.5126
## Lut - Ipsi - Decellularized - Fol - Ipsi - Native == 0             -13.6111
## Lut - Ipsi - Native - Fol - Ipsi - Native == 0                       8.8019
## Lut - Contra - Native - Lut - Contra - Decellularized == 0          10.7835
## Lut - Ipsi - Decellularized - Lut - Contra - Decellularized == 0    -3.3402
## Lut - Ipsi - Native - Lut - Contra - Decellularized == 0            19.0727
## Lut - Ipsi - Decellularized - Lut - Contra - Native == 0           -14.1237
## Lut - Ipsi - Native - Lut - Contra - Native == 0                     8.2892
## Lut - Ipsi - Native - Lut - Ipsi - Decellularized == 0              22.4130
##                                                                   Std. Error
## Fol - Contra - Native - Fol - Contra - Decellularized == 0              5.5033
## Fol - Ipsi - Decellularized - Fol - Contra - Decellularized == 0        5.3390
## Fol - Ipsi - Native - Fol - Contra - Decellularized == 0                5.3390
## Lut - Contra - Decellularized - Fol - Contra - Decellularized == 0      5.3390
## Lut - Contra - Native - Fol - Contra - Decellularized == 0              5.3390
## Lut - Ipsi - Decellularized - Fol - Contra - Decellularized == 0        5.3390
## Lut - Ipsi - Native - Fol - Contra - Decellularized == 0                5.3390
## Fol - Ipsi - Decellularized - Fol - Contra - Native == 0                5.5033
## Fol - Ipsi - Native - Fol - Contra - Native == 0                        5.5033
## Lut - Contra - Decellularized - Fol - Contra - Native == 0              5.5033
## Lut - Contra - Native - Fol - Contra - Native == 0                      5.5033
## Lut - Ipsi - Decellularized - Fol - Contra - Native == 0                5.5033
## Lut - Ipsi - Native - Fol - Contra - Native == 0                        5.5033
## Fol - Ipsi - Native - Fol - Ipsi - Decellularized == 0                  5.3390
## Lut - Contra - Decellularized - Fol - Ipsi - Decellularized == 0        5.3390
## Lut - Contra - Native - Fol - Ipsi - Decellularized == 0                5.3390
## Lut - Ipsi - Decellularized - Fol - Ipsi - Decellularized == 0          5.3390
## Lut - Ipsi - Native - Fol - Ipsi - Decellularized == 0                  5.3390
## Lut - Contra - Decellularized - Fol - Ipsi - Native == 0                5.3390
## Lut - Contra - Native - Fol - Ipsi - Native == 0                        5.3390
## Lut - Ipsi - Decellularized - Fol - Ipsi - Native == 0                  5.3390
## Lut - Ipsi - Native - Fol - Ipsi - Native == 0                          5.3390
## Lut - Contra - Native - Lut - Contra - Decellularized == 0              5.3390
## Lut - Ipsi - Decellularized - Lut - Contra - Decellularized == 0        5.3390
## Lut - Ipsi - Native - Lut - Contra - Decellularized == 0                5.3390
## Lut - Ipsi - Decellularized - Lut - Contra - Native == 0                5.3390
## Lut - Ipsi - Native - Lut - Contra - Native == 0                        5.3390
## Lut - Ipsi - Native - Lut - Ipsi - Decellularized == 0                  5.3390
##                                                                   t value
## Fol - Contra - Native - Fol - Contra - Decellularized == 0           5.797
## Fol - Ipsi - Decellularized - Fol - Contra - Decellularized == 0    -0.625
## Fol - Ipsi - Native - Fol - Contra - Decellularized == 0             4.602
## Lut - Contra - Decellularized - Fol - Contra - Decellularized == 0   2.678
## Lut - Contra - Native - Fol - Contra - Decellularized == 0           4.698
## Lut - Ipsi - Decellularized - Fol - Contra - Decellularized == 0     2.052
## Lut - Ipsi - Native - Fol - Contra - Decellularized == 0             6.250
## Fol - Ipsi - Decellularized - Fol - Contra - Native == 0            -6.404
## Fol - Ipsi - Native - Fol - Contra - Native == 0                    -1.333
```

```
## Lut - Contra - Decellularized - Fol - Contra - Native == 0          -3.199
## Lut - Contra - Native - Fol - Contra - Native == 0                  -1.240
## Lut - Ipsi - Decellularized - Fol - Contra - Native == 0            -3.806
## Lut - Ipsi - Native - Fol - Contra - Native == 0                     0.266
## Fol - Ipsi - Native - Fol - Ipsi - Decellularized == 0               5.227
## Lut - Contra - Decellularized - Fol - Ipsi - Decellularized == 0     3.303
## Lut - Contra - Native - Fol - Ipsi - Decellularized == 0             5.323
## Lut - Ipsi - Decellularized - Fol - Ipsi - Decellularized == 0       2.678
## Lut - Ipsi - Native - Fol - Ipsi - Decellularized == 0               6.876
## Lut - Contra - Decellularized - Fol - Ipsi - Native == 0            -1.924
## Lut - Contra - Native - Fol - Ipsi - Native == 0                     0.096
## Lut - Ipsi - Decellularized - Fol - Ipsi - Native == 0              -2.549
## Lut - Ipsi - Native - Fol - Ipsi - Native == 0                       1.649
## Lut - Contra - Native - Lut - Contra - Decellularized == 0           2.020
## Lut - Ipsi - Decellularized - Lut - Contra - Decellularized == 0    -0.626
## Lut - Ipsi - Native - Lut - Contra - Decellularized == 0             3.572
## Lut - Ipsi - Decellularized - Lut - Contra - Native == 0            -2.645
## Lut - Ipsi - Native - Lut - Contra - Native == 0                     1.553
## Lut - Ipsi - Native - Lut - Ipsi - Decellularized == 0               4.198
##                                                                    Pr(>|t|)
## Fol - Contra - Native - Fol - Contra - Decellularized == 0          < 0.001 ***
## Fol - Ipsi - Decellularized - Fol - Contra - Decellularized == 0    0.99838
## Fol - Ipsi - Native - Fol - Contra - Decellularized == 0            < 0.001 ***
## Lut - Contra - Decellularized - Fol - Contra - Decellularized == 0  0.14803
## Lut - Contra - Native - Fol - Contra - Decellularized == 0          < 0.001 ***
## Lut - Ipsi - Decellularized - Fol - Contra - Decellularized == 0    0.45667
## Lut - Ipsi - Native - Fol - Contra - Decellularized == 0            < 0.001 ***
## Fol - Ipsi - Decellularized - Fol - Contra - Native == 0            < 0.001 ***
## Fol - Ipsi - Native - Fol - Contra - Native == 0                    0.88270
## Lut - Contra - Decellularized - Fol - Contra - Native == 0          0.04204 *
## Lut - Contra - Native - Fol - Contra - Native == 0                  0.91657
## Lut - Ipsi - Decellularized - Fol - Contra - Native == 0            0.00730 **
## Lut - Ipsi - Native - Fol - Contra - Native == 0                    0.99999
## Fol - Ipsi - Native - Fol - Ipsi - Decellularized == 0              < 0.001 ***
## Lut - Contra - Decellularized - Fol - Ipsi - Decellularized == 0    0.03207 *
## Lut - Contra - Native - Fol - Ipsi - Decellularized == 0            < 0.001 ***
## Lut - Ipsi - Decellularized - Fol - Ipsi - Decellularized == 0      0.14883
## Lut - Ipsi - Native - Fol - Ipsi - Decellularized == 0              < 0.001 ***
## Lut - Contra - Decellularized - Fol - Ipsi - Native == 0            0.54004
## Lut - Contra - Native - Fol - Ipsi - Native == 0                    1.00000
## Lut - Ipsi - Decellularized - Fol - Ipsi - Native == 0              0.19439
## Lut - Ipsi - Native - Fol - Ipsi - Native == 0                      0.71889
## Lut - Contra - Native - Lut - Contra - Decellularized == 0          0.47695
## Lut - Ipsi - Decellularized - Lut - Contra - Decellularized == 0    0.99838
## Lut - Ipsi - Native - Lut - Contra - Decellularized == 0            0.01479 *
## Lut - Ipsi - Decellularized - Lut - Contra - Native == 0            0.15941
## Lut - Ipsi - Native - Lut - Contra - Native == 0                    0.77563
## Lut - Ipsi - Native - Lut - Ipsi - Decellularized == 0              0.00206 **
## ---
## Signif. codes:  0 '***' 0.001 '**' 0.01 '*' 0.05 '.' 0.1 ' ' 1
## (Adjusted p values reported -- single-step method)
```

```
################### YM-Porosity correlation ########################################
```

```r
# Read the dataset from a CSV file
df <- read.csv("Endometrium_YM3.csv")

# Update the labels for Position and Estrus_cycle
df$Position <- factor(df$Position, labels = c("Contralateral", "Ipsilateral"))
df$Estrus_cycle <- factor(df$Estrus_cycle, labels = c("Follicular phase", "Luteal phase"))

png(filename="Endometirum_correlation.png",
    width = 10, height = 15, units = "in",
    res = 600)

par(mgp=c(1.5,0.5,0),
    mar = c(4, #bottom
            4, #left
            3, #top
            2),
    cex.main = 0.8,
    family = "sans" #right
)

# Create scatter plot for Contra-lateral and Decellularized in the Follicular phase
p_contra_decell_follicular <- ggplot(df, aes(x = Porosity, y = YM, color = Cellular)) +
  geom_point(data = subset(df, Position == "Contralateral" & Estrus_cycle == "Follicular phase" & Cellul
  geom_smooth(data = subset(df, Position == "Contralateral" & Estrus_cycle == "Follicular phase" & Cellu
  labs(x = "Porosity (%)", y = "Young's Modulus (Pa)") +
  ggtitle("Contralateral - Decellularized (Follicular phase)") +
  stat_cor(data = subset(df, Position == "Contralateral" & Estrus_cycle == "Follicular phase" & Cellula
  theme_bw() +
  theme(panel.grid.major = element_blank(),
        panel.grid.minor = element_blank(),
        legend.position = "none",
        axis.title.x = element_text(size = 16),
        axis.title.y = element_text(size = 16))

# Create scatter plot for Contra-lateral and Native in the Follicular phase
p_contra_native_follicular <- ggplot(df, aes(x = Porosity, y = YM, color = Cellular)) +
  geom_point(data = subset(df, Position == "Contralateral" & Estrus_cycle == "Follicular phase" & Cellul
  geom_smooth(data = subset(df, Position == "Contralateral" & Estrus_cycle == "Follicular phase" & Cellu
  labs(x = "Porosity (%)", y = "Young's Modulus (Pa)") +
  ggtitle("Contralateral - Native (Follicular phase)") +
  stat_cor(data = subset(df, Position == "Contralateral" & Estrus_cycle == "Follicular phase" & Cellula
  theme_bw() +
  theme(panel.grid.major = element_blank(),
        panel.grid.minor = element_blank(),
        legend.position = "none",
        axis.title.x = element_text(size = 16),
        axis.title.y = element_text(size = 16))

# Create scatter plot for Ipsi-lateral and Decellularized in the Follicular phase
p_ipsi_decell_follicular <- ggplot(df, aes(x = Porosity, y = YM, color = Cellular)) +
  geom_point(data = subset(df, Position == "Ipsilateral" & Estrus_cycle == "Follicular phase" & Cellula
  geom_smooth(data = subset(df, Position == "Ipsilateral" & Estrus_cycle == "Follicular phase" & Cellula
  labs(x = "Porosity (%)", y = "Young's Modulus (Pa)") +
```

```r
  ggtitle("Ipsilateral - Decellularized (Follicular phase)") +
  stat_cor(data = subset(df, Position == "Ipsilateral" & Estrus_cycle == "Follicular phase" & Cellular =
  theme_bw() +
  theme(panel.grid.major = element_blank(),
        panel.grid.minor = element_blank(),
        legend.position = "none",
        axis.title.x = element_text(size = 16),
        axis.title.y = element_text(size = 16))

# Create scatter plot for Ipsi-lateral and Native in the Follicular phase
p_ipsi_native_follicular <- ggplot(df, aes(x = Porosity, y = YM, color = Cellular)) +
  geom_point(data = subset(df, Position == "Ipsilateral" & Estrus_cycle == "Follicular phase" & Cellula
  geom_smooth(data = subset(df, Position == "Ipsilateral" & Estrus_cycle == "Follicular phase" & Cellul
  labs(x = "Porosity (%)", y = "Young's Modulus (Pa)") +
  ggtitle("Ipsilateral - Native (Follicular phase)") +
  stat_cor(data = subset(df, Position == "Ipsilateral" & Estrus_cycle == "Follicular phase" & Cellular =
  theme_bw() +
  theme(panel.grid.major = element_blank(),
        panel.grid.minor = element_blank(),
        legend.position = "none",
        axis.title.x = element_text(size = 16),
        axis.title.y = element_text(size = 16))

# Create scatter plot for Contra-lateral and Decellularized in the Luteal phase
p_contra_decell_luteal <- ggplot(df, aes(x = Porosity, y = YM, color = Cellular)) +
  geom_point(data = subset(df, Position == "Contralateral" & Estrus_cycle == "Luteal phase" & Cellular =
  geom_smooth(data = subset(df, Position == "Contralateral" & Estrus_cycle == "Luteal phase" & Cellular
  labs(x = "Porosity (%)", y = "Young's Modulus (Pa)") +
  ggtitle("Contralateral - Decellularized (Luteal phase)") +
  stat_cor(data = subset(df, Position == "Contralateral" & Estrus_cycle == "Luteal phase" & Cellular ==
  theme_bw() +
  theme(panel.grid.major = element_blank(),
        panel.grid.minor = element_blank(),
        legend.position = "none",
        axis.title.x = element_text(size = 16),
        axis.title.y = element_text(size = 16))

# Create scatter plot for Contra-lateral and Native in the Luteal phase
p_contra_native_luteal <- ggplot(df, aes(x = Porosity, y = YM, color = Cellular)) +
  geom_point(data = subset(df, Position == "Contralateral" & Estrus_cycle == "Luteal phase" & Cellular =
  geom_smooth(data = subset(df, Position == "Contralateral" & Estrus_cycle == "Luteal phase" & Cellular
  labs(x = "Porosity (%)", y = "Young's Modulus (Pa)") +
  ggtitle("Contralateral - Native (Luteal phase)") +
  stat_cor(data = subset(df, Position == "Contralateral" & Estrus_cycle == "Luteal phase" & Cellular ==
  theme_bw() +
  theme(panel.grid.major = element_blank(),
        panel.grid.minor = element_blank(),
        legend.position = "none",
        axis.title.x = element_text(size = 16),
        axis.title.y = element_text(size = 16))

# Create scatter plot for Ipsi-lateral and Decellularized in the Luteal phase
p_ipsi_decell_luteal <- ggplot(df, aes(x = Porosity, y = YM, color = Cellular)) +
```

```
  geom_point(data = subset(df, Position == "Ipsilateral" & Estrus_cycle == "Luteal phase" & Cellular ==
  geom_smooth(data = subset(df, Position == "Ipsilateral" & Estrus_cycle == "Luteal phase" & Cellular =
  labs(x = "Porosity (%)", y = "Young's Modulus (Pa)") +
  ggtitle("Ipsilateral - Decellularized (Luteal phase)") +
  stat_cor(data = subset(df, Position == "Ipsilateral" & Estrus_cycle == "Luteal phase" & Cellular == "
  theme_bw() +
  theme(panel.grid.major = element_blank(),
        panel.grid.minor = element_blank(),
        legend.position = "none",
        axis.title.x = element_text(size = 16),
        axis.title.y = element_text(size = 16))

# Create scatter plot for Ipsi-lateral and Native in the Luteal phase
p_ipsi_native_luteal <- ggplot(df, aes(x = Porosity, y = YM, color = Cellular)) +
  geom_point(data = subset(df, Position == "Ipsilateral" & Estrus_cycle == "Luteal phase" & Cellular ==
  geom_smooth(data = subset(df, Position == "Ipsilateral" & Estrus_cycle == "Luteal phase" & Cellular =
  labs(x = "Porosity (%)", y = "Young's Modulus (Pa)") +
  ggtitle("Ipsilateral - Native (Luteal phase)") +
  stat_cor(data = subset(df, Position == "Ipsilateral" & Estrus_cycle == "Luteal phase" & Cellular == "
  theme_bw() +
  theme(panel.grid.major = element_blank(),
        panel.grid.minor = element_blank(),
        legend.position = "none",
        axis.title.x = element_text(size = 16),
        axis.title.y = element_text(size = 16))

# Combine the plots using cowplot
p_combined <- cowplot::plot_grid(
  cowplot::plot_grid(p_contra_decell_follicular, p_contra_native_follicular, nrow = 1),
  cowplot::plot_grid(p_ipsi_decell_follicular, p_ipsi_native_follicular, nrow = 1),
  cowplot::plot_grid(p_contra_decell_luteal, p_contra_native_luteal, nrow = 1),
  cowplot::plot_grid(p_ipsi_decell_luteal, p_ipsi_native_luteal, nrow = 1),
  nrow = 4
)
```

```
## `geom_smooth()` using formula = 'y ~ x'
## `geom_smooth()` using formula = 'y ~ x'
## `geom_smooth()` using formula = 'y ~ x'
## `geom_smooth()` using formula = 'y ~ x'
## `geom_smooth()` using formula = 'y ~ x'
## `geom_smooth()` using formula = 'y ~ x'
## `geom_smooth()` using formula = 'y ~ x'
## `geom_smooth()` using formula = 'y ~ x'
```

```
# Display the combined plot
p_combined


dev.off()
```

```
## pdf
##   2
```

```
#################################################################
#################### Ovary Analysis ##############################
```

```r
################################################################

################### E' E" ########################################

#data input
data <- read.csv("Ovary_rheo.csv")

# Create X_position_group based on X_position values
data$X_position_group <- ifelse(data$X_position <= 750, "Medulla", "Cortex")
data$X_position_group <- factor(data$X_position_group, levels = c("Cortex", "Medulla"))

# Subset the data based on X_position_group
data_cortex <- subset(data, X_position_group == "Cortex")
data_medulla <- subset(data, X_position_group == "Medulla")


# Plot Cortex

png(filename="E1_E2_cortex.png",
    width = 9.86, height = 6.86, units = "in",
    res = 600)

par(mgp=c(1.5,0.5,0),
    mar = c(4, #bottom
            4, #left
            3, #top
            2),
    cex.main = 0.8,
    family = "sans" #right
)


# Calculate means and standard deviations
data_summary <- data_cortex %>%
  group_by(Module, Estrus_cycle, Position, Hz) %>%
  summarise(mean_Indentation = mean(Indentation, na.rm = TRUE),
            sd_Indentation = sd(Indentation, na.rm = TRUE))
```

```
## `summarise()` has grouped output by 'Module', 'Estrus_cycle', 'Position'. You
## can override using the `.groups` argument.
```

```r
# Create a new variable for line type
data_summary <- data_summary %>%
  mutate(LineType = ifelse(Module == "E1", "E1", "E2"))

# Define fill colors for each factor level
fill_colors <- c("Follicular phase Contralateral" = viridis_pal(option = "D")(1),
                 "Follicular phase Ipsilateral" = alpha(viridis_pal(option = "D")(1), 0.2),
                 "Luteal phase Contralateral" = "#238A8DFF",
                 "Luteal phase Ipsilateral" = alpha("#238A8DFF", 0.2))

# Create the line plot with separate lines for each Estrus_cycle and Position
ggplot(data_summary, aes(x = Hz, y = mean_Indentation, group = interaction(Module, Estrus_cycle, Positi
                         color = paste(Estrus_cycle, Position), linetype = LineType)) +
```

```r
  geom_line(size = 1.5) +  # Adjust line thickness
  geom_point(size = 1) +  # Add dots at each Hz point
  scale_color_manual(values = fill_colors) +  # Use custom fill colors
  scale_linetype_manual(values = c("E1" = "solid", "E2" = "dashed"),
                        labels = c("E1" = "E'", "E2" = "E\"")) +
  scale_x_continuous(breaks = c(1, 5, 10, 20)) +
  scale_y_continuous(labels = scientific_format(), limits = c(0, 20000)) +  # Adjust Y-axis limits
  labs(y = expression(paste("E'/E'' (Pa)")), x = "Frequency (Hz)", color = "Estrus Cycle + Position") +
  theme_minimal() +
  theme(panel.grid.major = element_blank(),
        panel.grid.minor = element_blank(),
        panel.border = element_rect(colour = "black", fill = NA, size = 1),
        axis.text = element_text(size = 12))  # Increase font size of axis labels


dev.off()
```

```
## pdf
##   2
```

```r
# Plot Medulla

png(filename="E1_E2_medulla.png",
    width = 9.86, height = 6.86, units = "in",
    res = 600)

par(mgp=c(1.5,0.5,0),
    mar = c(4, #bottom
            4, #left
            3, #top
            2),
    cex.main = 0.8,
    family = "sans" #right
)


# Calculate means and standard deviations
data_summary <- data_medulla %>%
  group_by(Module, Estrus_cycle, Position, Hz) %>%
  summarise(mean_Indentation = mean(Indentation, na.rm = TRUE),
            sd_Indentation = sd(Indentation, na.rm = TRUE))
```

```
## `summarise()` has grouped output by 'Module', 'Estrus_cycle', 'Position'. You
## can override using the `.groups` argument.
```

```r
# Create a new variable for line type
data_summary <- data_summary %>%
  mutate(LineType = ifelse(Module == "E1", "E1", "E2"))

# Define fill colors for each factor level
fill_colors <- c("Follicular phase Contralateral" = viridis_pal(option = "D")(1),
                 "Follicular phase Ipsilateral" = alpha(viridis_pal(option = "D")(1), 0.2),
                 "Luteal phase Contralateral" = "#238A8DFF",
                 "Luteal phase Ipsilateral" = alpha("#238A8DFF", 0.2))
```

```r
# Create the line plot with separate lines for each Estrus_cycle and Position
ggplot(data_summary, aes(x = Hz, y = mean_Indentation, group = interaction(Module, Estrus_cycle, Positi
                         color = paste(Estrus_cycle, Position), linetype = LineType)) +
  geom_line(size = 1.5) +  # Adjust line thickness
  geom_point(size = 1) +  # Add dots at each Hz point
  scale_color_manual(values = fill_colors) +  # Use custom fill colors
  scale_linetype_manual(values = c("E1" = "solid", "E2" = "dashed"),
                        labels = c("E1" = "E'", "E2" = "E\"")) +
  scale_x_continuous(breaks = c(1, 5, 10, 20)) +
  scale_y_continuous(labels = scientific_format(), limits = c(0, 20000)) +  # Adjust Y-axis limits
  labs(y = expression(paste("E'/E'' (Pa)")), x = "Frequency (Hz)", color = "Estrus Cycle + Position") +
  theme_minimal() +
  theme(panel.grid.major = element_blank(),
        panel.grid.minor = element_blank(),
        panel.border = element_rect(colour = "black", fill = NA, size = 1),
        axis.text = element_text(size = 12))  # Increase font size of axis labels


dev.off()
```

```
## pdf
##   2
```

```r
# Stats

# Cortex
df <- data_cortex

# Create a factor variable for the comparisons
df$Comparison <- factor(paste(df$Estrus_cycle, df$Position, df$Module, sep = " - "))

# Perform pairwise comparisons using Tukey's method
model <- lm(Indentation ~ Comparison, data = df)
comp <- glht(model, linfct = mcp(Comparison = "Tukey"))

# Summarize the results
summary(comp)
```

```
##
##   Simultaneous Tests for General Linear Hypotheses
##
## Multiple Comparisons of Means: Tukey Contrasts
##
##
## Fit: lm(formula = Indentation ~ Comparison, data = df)
##
## Linear Hypotheses:
##                                                                          Estimate
## Follicular phase - Contralateral - E2 - Follicular phase - Contralateral - E1 == 0  -8989.8
## Follicular phase - Ipsilateral - E1 - Follicular phase - Contralateral - E1 == 0    -1727.6
## Follicular phase - Ipsilateral - E2 - Follicular phase - Contralateral - E1 == 0   -10294.1
## Luteal phase - Contralateral - E1 - Follicular phase - Contralateral - E1 == 0     -10469.8
## Luteal phase - Contralateral - E2 - Follicular phase - Contralateral - E1 == 0     -11960.3
## Luteal phase - Ipsilateral - E1 - Follicular phase - Contralateral - E1 == 0         -324.0
## Luteal phase - Ipsilateral - E2 - Follicular phase - Contralateral - E1 == 0        -9586.4
```

```
## Follicular phase - Ipsilateral - E1 - Follicular phase - Contralateral - E2 == 0     7262.2
## Follicular phase - Ipsilateral - E2 - Follicular phase - Contralateral - E2 == 0    -1304.3
## Luteal phase - Contralateral - E1 - Follicular phase - Contralateral - E2 == 0      -1480.0
## Luteal phase - Contralateral - E2 - Follicular phase - Contralateral - E2 == 0      -2970.5
## Luteal phase - Ipsilateral - E1 - Follicular phase - Contralateral - E2 == 0         8665.9
## Luteal phase - Ipsilateral - E2 - Follicular phase - Contralateral - E2 == 0         -596.6
## Follicular phase - Ipsilateral - E2 - Follicular phase - Ipsilateral - E1 == 0      -8566.5
## Luteal phase - Contralateral - E1 - Follicular phase - Ipsilateral - E1 == 0        -8742.2
## Luteal phase - Contralateral - E2 - Follicular phase - Ipsilateral - E1 == 0       -10232.7
## Luteal phase - Ipsilateral - E1 - Follicular phase - Ipsilateral - E1 == 0           1403.7
## Luteal phase - Ipsilateral - E2 - Follicular phase - Ipsilateral - E1 == 0          -7858.8
## Luteal phase - Contralateral - E1 - Follicular phase - Ipsilateral - E2 == 0         -175.7
## Luteal phase - Contralateral - E2 - Follicular phase - Ipsilateral - E2 == 0        -1666.2
## Luteal phase - Ipsilateral - E1 - Follicular phase - Ipsilateral - E2 == 0           9970.1
## Luteal phase - Ipsilateral - E2 - Follicular phase - Ipsilateral - E2 == 0            707.7
## Luteal phase - Contralateral - E2 - Luteal phase - Contralateral - E1 == 0          -1490.5
## Luteal phase - Ipsilateral - E1 - Luteal phase - Contralateral - E1 == 0            10145.9
## Luteal phase - Ipsilateral - E2 - Luteal phase - Contralateral - E1 == 0              883.4
## Luteal phase - Ipsilateral - E1 - Luteal phase - Contralateral - E2 == 0            11636.3
## Luteal phase - Ipsilateral - E2 - Luteal phase - Contralateral - E2 == 0             2373.9
## Luteal phase - Ipsilateral - E2 - Luteal phase - Ipsilateral - E1 == 0              -9262.5
##                                                                                    Std. Error
## Follicular phase - Contralateral - E2 - Follicular phase - Contralateral - E1 == 0     1109.1
## Follicular phase - Ipsilateral - E1 - Follicular phase - Contralateral - E1 == 0       1109.1
## Follicular phase - Ipsilateral - E2 - Follicular phase - Contralateral - E1 == 0       1109.1
## Luteal phase - Contralateral - E1 - Follicular phase - Contralateral - E1 == 0         1271.8
## Luteal phase - Contralateral - E2 - Follicular phase - Contralateral - E1 == 0         1302.9
## Luteal phase - Ipsilateral - E1 - Follicular phase - Contralateral - E1 == 0           1250.9
## Luteal phase - Ipsilateral - E2 - Follicular phase - Contralateral - E1 == 0           1264.6
## Follicular phase - Ipsilateral - E1 - Follicular phase - Contralateral - E2 == 0       1109.1
## Follicular phase - Ipsilateral - E2 - Follicular phase - Contralateral - E2 == 0       1109.1
## Luteal phase - Contralateral - E1 - Follicular phase - Contralateral - E2 == 0         1271.8
## Luteal phase - Contralateral - E2 - Follicular phase - Contralateral - E2 == 0         1302.9
## Luteal phase - Ipsilateral - E1 - Follicular phase - Contralateral - E2 == 0           1250.9
## Luteal phase - Ipsilateral - E2 - Follicular phase - Contralateral - E2 == 0           1264.6
## Follicular phase - Ipsilateral - E2 - Follicular phase - Ipsilateral - E1 == 0         1109.1
## Luteal phase - Contralateral - E1 - Follicular phase - Ipsilateral - E1 == 0           1271.8
## Luteal phase - Contralateral - E2 - Follicular phase - Ipsilateral - E1 == 0           1302.9
## Luteal phase - Ipsilateral - E1 - Follicular phase - Ipsilateral - E1 == 0             1250.9
## Luteal phase - Ipsilateral - E2 - Follicular phase - Ipsilateral - E1 == 0             1264.6
## Luteal phase - Contralateral - E1 - Follicular phase - Ipsilateral - E2 == 0           1271.8
## Luteal phase - Contralateral - E2 - Follicular phase - Ipsilateral - E2 == 0           1302.9
## Luteal phase - Ipsilateral - E1 - Follicular phase - Ipsilateral - E2 == 0             1250.9
## Luteal phase - Ipsilateral - E2 - Follicular phase - Ipsilateral - E2 == 0             1264.6
## Luteal phase - Contralateral - E2 - Luteal phase - Contralateral - E1 == 0             1443.9
## Luteal phase - Ipsilateral - E1 - Luteal phase - Contralateral - E1 == 0               1397.1
## Luteal phase - Ipsilateral - E2 - Luteal phase - Contralateral - E1 == 0               1409.4
## Luteal phase - Ipsilateral - E1 - Luteal phase - Contralateral - E2 == 0               1425.5
## Luteal phase - Ipsilateral - E2 - Luteal phase - Contralateral - E2 == 0               1437.6
## Luteal phase - Ipsilateral - E2 - Luteal phase - Ipsilateral - E1 == 0                 1390.6
##                                                                                    t value
## Follicular phase - Contralateral - E2 - Follicular phase - Contralateral - E1 == 0  -8.105
## Follicular phase - Ipsilateral - E1 - Follicular phase - Contralateral - E1 == 0    -1.558
## Follicular phase - Ipsilateral - E2 - Follicular phase - Contralateral - E1 == 0    -9.281
```

```
## Luteal phase - Contralateral - E1 - Follicular phase - Contralateral - E1 == 0      -8.233
## Luteal phase - Contralateral - E2 - Follicular phase - Contralateral - E1 == 0      -9.180
## Luteal phase - Ipsilateral - E1 - Follicular phase - Contralateral - E1 == 0        -0.259
## Luteal phase - Ipsilateral - E2 - Follicular phase - Contralateral - E1 == 0        -7.581
## Follicular phase - Ipsilateral - E1 - Follicular phase - Contralateral - E2 == 0     6.548
## Follicular phase - Ipsilateral - E2 - Follicular phase - Contralateral - E2 == 0    -1.176
## Luteal phase - Contralateral - E1 - Follicular phase - Contralateral - E2 == 0      -1.164
## Luteal phase - Contralateral - E2 - Follicular phase - Contralateral - E2 == 0      -2.280
## Luteal phase - Ipsilateral - E1 - Follicular phase - Contralateral - E2 == 0         6.928
## Luteal phase - Ipsilateral - E2 - Follicular phase - Contralateral - E2 == 0        -0.472
## Follicular phase - Ipsilateral - E2 - Follicular phase - Ipsilateral - E1 == 0      -7.724
## Luteal phase - Contralateral - E1 - Follicular phase - Ipsilateral - E1 == 0        -6.874
## Luteal phase - Contralateral - E2 - Follicular phase - Ipsilateral - E1 == 0        -7.854
## Luteal phase - Ipsilateral - E1 - Follicular phase - Ipsilateral - E1 == 0           1.122
## Luteal phase - Ipsilateral - E2 - Follicular phase - Ipsilateral - E1 == 0          -6.215
## Luteal phase - Contralateral - E1 - Follicular phase - Ipsilateral - E2 == 0        -0.138
## Luteal phase - Contralateral - E2 - Follicular phase - Ipsilateral - E2 == 0        -1.279
## Luteal phase - Ipsilateral - E1 - Follicular phase - Ipsilateral - E2 == 0           7.971
## Luteal phase - Ipsilateral - E2 - Follicular phase - Ipsilateral - E2 == 0           0.560
## Luteal phase - Contralateral - E2 - Luteal phase - Contralateral - E1 == 0          -1.032
## Luteal phase - Ipsilateral - E1 - Luteal phase - Contralateral - E1 == 0             7.262
## Luteal phase - Ipsilateral - E2 - Luteal phase - Contralateral - E1 == 0             0.627
## Luteal phase - Ipsilateral - E1 - Luteal phase - Contralateral - E2 == 0             8.163
## Luteal phase - Ipsilateral - E2 - Luteal phase - Contralateral - E2 == 0             1.651
## Luteal phase - Ipsilateral - E2 - Luteal phase - Ipsilateral - E1 == 0              -6.661
##                                                                                   Pr(>|t|)
## Follicular phase - Contralateral - E2 - Follicular phase - Contralateral - E1 == 0   <0.001
## Follicular phase - Ipsilateral - E1 - Follicular phase - Contralateral - E1 == 0      0.773
## Follicular phase - Ipsilateral - E2 - Follicular phase - Contralateral - E1 == 0     <0.001
## Luteal phase - Contralateral - E1 - Follicular phase - Contralateral - E1 == 0       <0.001
## Luteal phase - Contralateral - E2 - Follicular phase - Contralateral - E1 == 0       <0.001
## Luteal phase - Ipsilateral - E1 - Follicular phase - Contralateral - E1 == 0          1.000
## Luteal phase - Ipsilateral - E2 - Follicular phase - Contralateral - E1 == 0         <0.001
## Follicular phase - Ipsilateral - E1 - Follicular phase - Contralateral - E2 == 0     <0.001
## Follicular phase - Ipsilateral - E2 - Follicular phase - Contralateral - E2 == 0      0.938
## Luteal phase - Contralateral - E1 - Follicular phase - Contralateral - E2 == 0        0.941
## Luteal phase - Contralateral - E2 - Follicular phase - Contralateral - E2 == 0        0.304
## Luteal phase - Ipsilateral - E1 - Follicular phase - Contralateral - E2 == 0         <0.001
## Luteal phase - Ipsilateral - E2 - Follicular phase - Contralateral - E2 == 0          1.000
## Follicular phase - Ipsilateral - E2 - Follicular phase - Ipsilateral - E1 == 0       <0.001
## Luteal phase - Contralateral - E1 - Follicular phase - Ipsilateral - E1 == 0         <0.001
## Luteal phase - Contralateral - E2 - Follicular phase - Ipsilateral - E1 == 0         <0.001
## Luteal phase - Ipsilateral - E1 - Follicular phase - Ipsilateral - E1 == 0            0.951
## Luteal phase - Ipsilateral - E2 - Follicular phase - Ipsilateral - E1 == 0           <0.001
## Luteal phase - Contralateral - E1 - Follicular phase - Ipsilateral - E2 == 0          1.000
## Luteal phase - Contralateral - E2 - Follicular phase - Ipsilateral - E2 == 0          0.905
## Luteal phase - Ipsilateral - E1 - Follicular phase - Ipsilateral - E2 == 0           <0.001
## Luteal phase - Ipsilateral - E2 - Follicular phase - Ipsilateral - E2 == 0            0.999
## Luteal phase - Contralateral - E2 - Luteal phase - Contralateral - E1 == 0            0.969
## Luteal phase - Ipsilateral - E1 - Luteal phase - Contralateral - E1 == 0             <0.001
## Luteal phase - Ipsilateral - E2 - Luteal phase - Contralateral - E1 == 0              0.998
## Luteal phase - Ipsilateral - E1 - Luteal phase - Contralateral - E2 == 0             <0.001
## Luteal phase - Ipsilateral - E2 - Luteal phase - Contralateral - E2 == 0              0.716
## Luteal phase - Ipsilateral - E2 - Luteal phase - Ipsilateral - E1 == 0               <0.001
```

```
##
## Follicular phase - Contralateral - E2 - Follicular phase - Contralateral - E1 == 0 ***
## Follicular phase - Ipsilateral - E1 - Follicular phase - Contralateral - E1 == 0
## Follicular phase - Ipsilateral - E2 - Follicular phase - Contralateral - E1 == 0   ***
## Luteal phase - Contralateral - E1 - Follicular phase - Contralateral - E1 == 0     ***
## Luteal phase - Contralateral - E2 - Follicular phase - Contralateral - E1 == 0     ***
## Luteal phase - Ipsilateral - E1 - Follicular phase - Contralateral - E1 == 0
## Luteal phase - Ipsilateral - E2 - Follicular phase - Contralateral - E1 == 0       ***
## Follicular phase - Ipsilateral - E1 - Follicular phase - Contralateral - E2 == 0   ***
## Follicular phase - Ipsilateral - E2 - Follicular phase - Contralateral - E2 == 0
## Luteal phase - Contralateral - E1 - Follicular phase - Contralateral - E2 == 0
## Luteal phase - Contralateral - E2 - Follicular phase - Contralateral - E2 == 0
## Luteal phase - Ipsilateral - E1 - Follicular phase - Contralateral - E2 == 0       ***
## Luteal phase - Ipsilateral - E2 - Follicular phase - Contralateral - E2 == 0
## Follicular phase - Ipsilateral - E2 - Follicular phase - Ipsilateral - E1 == 0     ***
## Luteal phase - Contralateral - E1 - Follicular phase - Ipsilateral - E1 == 0       ***
## Luteal phase - Contralateral - E2 - Follicular phase - Ipsilateral - E1 == 0       ***
## Luteal phase - Ipsilateral - E1 - Follicular phase - Ipsilateral - E1 == 0
## Luteal phase - Ipsilateral - E2 - Follicular phase - Ipsilateral - E1 == 0         ***
## Luteal phase - Contralateral - E1 - Follicular phase - Ipsilateral - E2 == 0
## Luteal phase - Contralateral - E2 - Follicular phase - Ipsilateral - E2 == 0
## Luteal phase - Ipsilateral - E1 - Follicular phase - Ipsilateral - E2 == 0         ***
## Luteal phase - Ipsilateral - E2 - Follicular phase - Ipsilateral - E2 == 0
## Luteal phase - Contralateral - E2 - Luteal phase - Contralateral - E1 == 0
## Luteal phase - Ipsilateral - E1 - Luteal phase - Contralateral - E1 == 0           ***
## Luteal phase - Ipsilateral - E2 - Luteal phase - Contralateral - E1 == 0
## Luteal phase - Ipsilateral - E1 - Luteal phase - Contralateral - E2 == 0           ***
## Luteal phase - Ipsilateral - E2 - Luteal phase - Contralateral - E2 == 0
## Luteal phase - Ipsilateral - E2 - Luteal phase - Ipsilateral - E1 == 0             ***
## ---
## Signif. codes:  0 '***' 0.001 '**' 0.01 '*' 0.05 '.' 0.1 ' ' 1
## (Adjusted p values reported -- single-step method)
```

```r
# Medulla
df <- data_medulla

# Create a factor variable for the comparisons
df$Comparison <- factor(paste(df$Estrus_cycle, df$Position, df$Module, sep = " - "))

# Perform pairwise comparisons using Tukey's method
model <- lm(Indentation ~ Comparison, data = df)
comp <- glht(model, linfct = mcp(Comparison = "Tukey"))

# Summarize the results
summary(comp)
```

```
## Warning in RET$pfunction("adjusted", ...): Completion with error > abseps

## Warning in RET$pfunction("adjusted", ...): Completion with error > abseps

##
##   Simultaneous Tests for General Linear Hypotheses
##
## Multiple Comparisons of Means: Tukey Contrasts
##
##
```

```
## Fit: lm(formula = Indentation ~ Comparison, data = df)
##
## Linear Hypotheses:
##                                                                               Estimate
## Follicular phase - Contralateral - E2 - Follicular phase - Contralateral - E1 == 0  -8084.4
## Follicular phase - Ipsilateral - E1 - Follicular phase - Contralateral - E1 == 0    -6729.9
## Follicular phase - Ipsilateral - E2 - Follicular phase - Contralateral - E1 == 0   -11132.0
## Luteal phase - Contralateral - E1 - Follicular phase - Contralateral - E1 == 0      -9904.3
## Luteal phase - Contralateral - E2 - Follicular phase - Contralateral - E1 == 0     -10292.2
## Luteal phase - Ipsilateral - E1 - Follicular phase - Contralateral - E1 == 0        -5071.8
## Luteal phase - Ipsilateral - E2 - Follicular phase - Contralateral - E1 == 0       -10820.2
## Follicular phase - Ipsilateral - E1 - Follicular phase - Contralateral - E2 == 0     1354.5
## Follicular phase - Ipsilateral - E2 - Follicular phase - Contralateral - E2 == 0    -3047.6
## Luteal phase - Contralateral - E1 - Follicular phase - Contralateral - E2 == 0      -1819.9
## Luteal phase - Contralateral - E2 - Follicular phase - Contralateral - E2 == 0      -2207.9
## Luteal phase - Ipsilateral - E1 - Follicular phase - Contralateral - E2 == 0         3012.6
## Luteal phase - Ipsilateral - E2 - Follicular phase - Contralateral - E2 == 0        -2735.9
## Follicular phase - Ipsilateral - E2 - Follicular phase - Ipsilateral - E1 == 0      -4402.1
## Luteal phase - Contralateral - E1 - Follicular phase - Ipsilateral - E1 == 0        -3174.4
## Luteal phase - Contralateral - E2 - Follicular phase - Ipsilateral - E1 == 0        -3562.4
## Luteal phase - Ipsilateral - E1 - Follicular phase - Ipsilateral - E1 == 0           1658.1
## Luteal phase - Ipsilateral - E2 - Follicular phase - Ipsilateral - E1 == 0          -4090.4
## Luteal phase - Contralateral - E1 - Follicular phase - Ipsilateral - E2 == 0         1227.7
## Luteal phase - Contralateral - E2 - Follicular phase - Ipsilateral - E2 == 0          839.8
## Luteal phase - Ipsilateral - E1 - Follicular phase - Ipsilateral - E2 == 0           6060.2
## Luteal phase - Ipsilateral - E2 - Follicular phase - Ipsilateral - E2 == 0            311.8
## Luteal phase - Contralateral - E2 - Luteal phase - Contralateral - E1 == 0           -387.9
## Luteal phase - Ipsilateral - E1 - Luteal phase - Contralateral - E1 == 0             4832.5
## Luteal phase - Ipsilateral - E2 - Luteal phase - Contralateral - E1 == 0             -916.0
## Luteal phase - Ipsilateral - E1 - Luteal phase - Contralateral - E2 == 0             5220.4
## Luteal phase - Ipsilateral - E2 - Luteal phase - Contralateral - E2 == 0             -528.0
## Luteal phase - Ipsilateral - E2 - Luteal phase - Ipsilateral - E1 == 0              -5748.4
##                                                                               Std. Error
## Follicular phase - Contralateral - E2 - Follicular phase - Contralateral - E1 == 0      927.5
## Follicular phase - Ipsilateral - E1 - Follicular phase - Contralateral - E1 == 0       978.8
## Follicular phase - Ipsilateral - E2 - Follicular phase - Contralateral - E1 == 0       978.8
## Luteal phase - Contralateral - E1 - Follicular phase - Contralateral - E1 == 0        1064.9
## Luteal phase - Contralateral - E2 - Follicular phase - Contralateral - E1 == 0        1086.0
## Luteal phase - Ipsilateral - E1 - Follicular phase - Contralateral - E1 == 0          1032.7
## Luteal phase - Ipsilateral - E2 - Follicular phase - Contralateral - E1 == 0          1080.5
## Follicular phase - Ipsilateral - E1 - Follicular phase - Contralateral - E2 == 0       978.8
## Follicular phase - Ipsilateral - E2 - Follicular phase - Contralateral - E2 == 0       978.8
## Luteal phase - Contralateral - E1 - Follicular phase - Contralateral - E2 == 0        1064.9
## Luteal phase - Contralateral - E2 - Follicular phase - Contralateral - E2 == 0        1086.0
## Luteal phase - Ipsilateral - E1 - Follicular phase - Contralateral - E2 == 0          1032.7
## Luteal phase - Ipsilateral - E2 - Follicular phase - Contralateral - E2 == 0          1080.5
## Follicular phase - Ipsilateral - E2 - Follicular phase - Ipsilateral - E1 == 0        1027.5
## Luteal phase - Contralateral - E1 - Follicular phase - Ipsilateral - E1 == 0          1109.9
## Luteal phase - Contralateral - E2 - Follicular phase - Ipsilateral - E1 == 0          1130.1
## Luteal phase - Ipsilateral - E1 - Follicular phase - Ipsilateral - E1 == 0            1079.0
## Luteal phase - Ipsilateral - E2 - Follicular phase - Ipsilateral - E1 == 0            1124.9
## Luteal phase - Contralateral - E1 - Follicular phase - Ipsilateral - E2 == 0          1109.9
## Luteal phase - Contralateral - E2 - Follicular phase - Ipsilateral - E2 == 0          1130.1
## Luteal phase - Ipsilateral - E1 - Follicular phase - Ipsilateral - E2 == 0            1079.0
```

```
## Luteal phase - Ipsilateral - E2 - Follicular phase - Ipsilateral - E2 == 0            1124.9
## Luteal phase - Contralateral - E2 - Luteal phase - Contralateral - E1 == 0            1205.5
## Luteal phase - Ipsilateral - E1 - Luteal phase - Contralateral - E1 == 0              1157.7
## Luteal phase - Ipsilateral - E2 - Luteal phase - Contralateral - E1 == 0              1200.5
## Luteal phase - Ipsilateral - E1 - Luteal phase - Contralateral - E2 == 0              1177.1
## Luteal phase - Ipsilateral - E2 - Luteal phase - Contralateral - E2 == 0              1219.3
## Luteal phase - Ipsilateral - E2 - Luteal phase - Ipsilateral - E1 == 0                1172.1
##                                                                                    t value
## Follicular phase - Contralateral - E2 - Follicular phase - Contralateral - E1 == 0  -8.716
## Follicular phase - Ipsilateral - E1 - Follicular phase - Contralateral - E1 == 0    -6.876
## Follicular phase - Ipsilateral - E2 - Follicular phase - Contralateral - E1 == 0   -11.373
## Luteal phase - Contralateral - E1 - Follicular phase - Contralateral - E1 == 0      -9.301
## Luteal phase - Contralateral - E2 - Follicular phase - Contralateral - E1 == 0      -9.477
## Luteal phase - Ipsilateral - E1 - Follicular phase - Contralateral - E1 == 0        -4.911
## Luteal phase - Ipsilateral - E2 - Follicular phase - Contralateral - E1 == 0       -10.014
## Follicular phase - Ipsilateral - E1 - Follicular phase - Contralateral - E2 == 0     1.384
## Follicular phase - Ipsilateral - E2 - Follicular phase - Contralateral - E2 == 0    -3.114
## Luteal phase - Contralateral - E1 - Follicular phase - Contralateral - E2 == 0      -1.709
## Luteal phase - Contralateral - E2 - Follicular phase - Contralateral - E2 == 0      -2.033
## Luteal phase - Ipsilateral - E1 - Follicular phase - Contralateral - E2 == 0         2.917
## Luteal phase - Ipsilateral - E2 - Follicular phase - Contralateral - E2 == 0        -2.532
## Follicular phase - Ipsilateral - E2 - Follicular phase - Ipsilateral - E1 == 0      -4.284
## Luteal phase - Contralateral - E1 - Follicular phase - Ipsilateral - E1 == 0        -2.860
## Luteal phase - Contralateral - E2 - Follicular phase - Ipsilateral - E1 == 0        -3.152
## Luteal phase - Ipsilateral - E1 - Follicular phase - Ipsilateral - E1 == 0           1.537
## Luteal phase - Ipsilateral - E2 - Follicular phase - Ipsilateral - E1 == 0          -3.636
## Luteal phase - Contralateral - E1 - Follicular phase - Ipsilateral - E2 == 0         1.106
## Luteal phase - Contralateral - E2 - Follicular phase - Ipsilateral - E2 == 0         0.743
## Luteal phase - Ipsilateral - E1 - Follicular phase - Ipsilateral - E2 == 0           5.616
## Luteal phase - Ipsilateral - E2 - Follicular phase - Ipsilateral - E2 == 0           0.277
## Luteal phase - Contralateral - E2 - Luteal phase - Contralateral - E1 == 0          -0.322
## Luteal phase - Ipsilateral - E1 - Luteal phase - Contralateral - E1 == 0             4.174
## Luteal phase - Ipsilateral - E2 - Luteal phase - Contralateral - E1 == 0            -0.763
## Luteal phase - Ipsilateral - E1 - Luteal phase - Contralateral - E2 == 0             4.435
## Luteal phase - Ipsilateral - E2 - Luteal phase - Contralateral - E2 == 0            -0.433
## Luteal phase - Ipsilateral - E2 - Luteal phase - Ipsilateral - E1 == 0              -4.904
##                                                                                   Pr(>|t|)
## Follicular phase - Contralateral - E2 - Follicular phase - Contralateral - E1 == 0    <0.01
## Follicular phase - Ipsilateral - E1 - Follicular phase - Contralateral - E1 == 0      <0.01
## Follicular phase - Ipsilateral - E2 - Follicular phase - Contralateral - E1 == 0      <0.01
## Luteal phase - Contralateral - E1 - Follicular phase - Contralateral - E1 == 0        <0.01
## Luteal phase - Contralateral - E2 - Follicular phase - Contralateral - E1 == 0        <0.01
## Luteal phase - Ipsilateral - E1 - Follicular phase - Contralateral - E1 == 0          <0.01
## Luteal phase - Ipsilateral - E2 - Follicular phase - Contralateral - E1 == 0          <0.01
## Follicular phase - Ipsilateral - E1 - Follicular phase - Contralateral - E2 == 0     0.8632
## Follicular phase - Ipsilateral - E2 - Follicular phase - Contralateral - E2 == 0     0.0398
## Luteal phase - Contralateral - E1 - Follicular phase - Contralateral - E2 == 0       0.6791
## Luteal phase - Contralateral - E2 - Follicular phase - Contralateral - E2 == 0       0.4575
## Luteal phase - Ipsilateral - E1 - Follicular phase - Contralateral - E2 == 0         0.0699
## Luteal phase - Ipsilateral - E2 - Follicular phase - Contralateral - E2 == 0         0.1815
## Follicular phase - Ipsilateral - E2 - Follicular phase - Ipsilateral - E1 == 0        <0.01
## Luteal phase - Contralateral - E1 - Follicular phase - Ipsilateral - E1 == 0         0.0821
## Luteal phase - Contralateral - E2 - Follicular phase - Ipsilateral - E1 == 0         0.0355
## Luteal phase - Ipsilateral - E1 - Follicular phase - Ipsilateral - E1 == 0           0.7853
```

```
## Luteal phase - Ipsilateral - E2 - Follicular phase - Ipsilateral - E1 == 0           <0.01
## Luteal phase - Contralateral - E1 - Follicular phase - Ipsilateral - E2 == 0        0.9549
## Luteal phase - Contralateral - E2 - Follicular phase - Ipsilateral - E2 == 0        0.9956
## Luteal phase - Ipsilateral - E1 - Follicular phase - Ipsilateral - E2 == 0           <0.01
## Luteal phase - Ipsilateral - E2 - Follicular phase - Ipsilateral - E2 == 0          1.0000
## Luteal phase - Contralateral - E2 - Luteal phase - Contralateral - E1 == 0          1.0000
## Luteal phase - Ipsilateral - E1 - Luteal phase - Contralateral - E1 == 0             <0.01
## Luteal phase - Ipsilateral - E2 - Luteal phase - Contralateral - E1 == 0            0.9948
## Luteal phase - Ipsilateral - E1 - Luteal phase - Contralateral - E2 == 0             <0.01
## Luteal phase - Ipsilateral - E2 - Luteal phase - Contralateral - E2 == 0            0.9999
## Luteal phase - Ipsilateral - E2 - Luteal phase - Ipsilateral - E1 == 0               <0.01
##
## Follicular phase - Contralateral - E2 - Follicular phase - Contralateral - E1 == 0 ***
## Follicular phase - Ipsilateral - E1 - Follicular phase - Contralateral - E1 == 0   ***
## Follicular phase - Ipsilateral - E2 - Follicular phase - Contralateral - E1 == 0   ***
## Luteal phase - Contralateral - E1 - Follicular phase - Contralateral - E1 == 0     ***
## Luteal phase - Contralateral - E2 - Follicular phase - Contralateral - E1 == 0     ***
## Luteal phase - Ipsilateral - E1 - Follicular phase - Contralateral - E1 == 0       ***
## Luteal phase - Ipsilateral - E2 - Follicular phase - Contralateral - E1 == 0       ***
## Follicular phase - Ipsilateral - E1 - Follicular phase - Contralateral - E2 == 0
## Follicular phase - Ipsilateral - E2 - Follicular phase - Contralateral - E2 == 0   *
## Luteal phase - Contralateral - E1 - Follicular phase - Contralateral - E2 == 0
## Luteal phase - Contralateral - E2 - Follicular phase - Contralateral - E2 == 0
## Luteal phase - Ipsilateral - E1 - Follicular phase - Contralateral - E2 == 0       .
## Luteal phase - Ipsilateral - E2 - Follicular phase - Contralateral - E2 == 0
## Follicular phase - Ipsilateral - E2 - Follicular phase - Ipsilateral - E1 == 0     ***
## Luteal phase - Contralateral - E1 - Follicular phase - Ipsilateral - E1 == 0       .
## Luteal phase - Contralateral - E2 - Follicular phase - Ipsilateral - E1 == 0       *
## Luteal phase - Ipsilateral - E1 - Follicular phase - Ipsilateral - E1 == 0
## Luteal phase - Ipsilateral - E2 - Follicular phase - Ipsilateral - E1 == 0         **
## Luteal phase - Contralateral - E1 - Follicular phase - Ipsilateral - E2 == 0
## Luteal phase - Contralateral - E2 - Follicular phase - Ipsilateral - E2 == 0
## Luteal phase - Ipsilateral - E1 - Follicular phase - Ipsilateral - E2 == 0         ***
## Luteal phase - Ipsilateral - E2 - Follicular phase - Ipsilateral - E2 == 0
## Luteal phase - Contralateral - E2 - Luteal phase - Contralateral - E1 == 0
## Luteal phase - Ipsilateral - E1 - Luteal phase - Contralateral - E1 == 0           ***
## Luteal phase - Ipsilateral - E2 - Luteal phase - Contralateral - E1 == 0
## Luteal phase - Ipsilateral - E1 - Luteal phase - Contralateral - E2 == 0           ***
## Luteal phase - Ipsilateral - E2 - Luteal phase - Contralateral - E2 == 0
## Luteal phase - Ipsilateral - E2 - Luteal phase - Ipsilateral - E1 == 0             ***
## ---
## Signif. codes:  0 '***' 0.001 '**' 0.01 '*' 0.05 '.' 0.1 ' ' 1
## (Adjusted p values reported -- single-step method)
```

```r
################## YM #######################################

# Read the dataset from a CSV file
df <- read.csv("Ovary_YM.csv")

# Create X_position_group based on X_position values
df$X_position_group <- ifelse(df$X_position <= 750, "Medulla", "Cortex")
df$X_position_group <- factor(df$X_position_group, levels = c("Cortex", "Medulla"))

# Summary data
```

```r
summary_df <- df %>%
  group_by(Cellular, Tissue, Position, Estrus_cycle, X_position_group) %>%
  summarise(mean_YM = mean(YM),
            sd_YM = sd(YM))
```

```
## `summarise()` has grouped output by 'Cellular', 'Tissue', 'Position',
## 'Estrus_cycle'. You can override using the `.groups` argument.
```

```r
# Plot considering distance

png(filename="Ovary_YM_all.png",
    width = 17, height = 10, units = "in",
    res = 600)

par(mgp=c(1.5,0.5,0),
    mar = c(4, #bottom
            4, #left
            3, #top
            2),
    cex.main = 0.8,
    family = "sans" #right
)

# Update the labels for Position and Estrus_cycle
df$Position <- factor(df$Position, labels = c("Contralateral", "Ipsilateral"))
df$Estrus_cycle <- factor(df$Estrus_cycle, labels = c("Follicular phase", "Luteal phase"))

# Create a combined factor variable for Estrus_cycle and Cellular
df$Combined <- factor(paste(df$Estrus_cycle, df$Cellular, sep = " "),
                      levels = c("Follicular phase Native", "Follicular phase Decellularized",
                                 "Luteal phase Native", "Luteal phase Decellularized"))

# Create the plot using ggplot
ggplot(data = df, aes(x = factor(X_position), y = YM, fill = Combined)) +
  geom_boxplot(outlier.shape = NA) +
  geom_jitter(position = position_jitterdodge(), alpha = 1) +
  xlab("Distance from medulla to cortex (um)") +
  ylab("Young's modulus (Pa)") +
  ggtitle("Boxplot and Individual Data Points of YM by X_position, Estrus_cycle, Position, and Estrus_c
  scale_fill_manual(values = c("Follicular phase Native" = viridis_pal(option = "D")(1),
                               "Follicular phase Decellularized" = alpha(viridis_pal(option = "D")(1), 
                               "Luteal phase Native" = "#238A8DFF",
                               "Luteal phase Decellularized" = alpha("#238A8DFF", 0.2)),
                    guide = guide_legend(title = "Combined")) +
  scale_color_viridis_d() +
  theme_bw() +
  facet_grid(Position ~ Estrus_cycle + Cellular, scales = "free_x", space = "free_x") +
  ylim(0, 30000) +
  theme(panel.grid.major = element_blank(), panel.grid.minor = element_blank(),
        axis.text.x = element_text(size = 10, angle = 90, vjust = 0.5, hjust = 0),
        axis.text.y = element_text(size = 12, angle = 0, hjust = 0.5),
        axis.title.x = element_text(size = 16, face = "bold"),
        axis.title.y = element_text(size = 16, face = "bold"),
        legend.position = "right",
```

```
        strip.text = element_text(size = 14, face = "bold"),
        legend.text = element_text(size = 14))  # Increase the size of the legend font

dev.off()
```

```
## pdf
##   2
```

```
# Plot cortex and medulla

png(filename="Ovary_YM_CTX_med.png",
    width = 10, height = 8, units = "in",
    res = 600)

par(mgp=c(1.5,0.5,0),
    mar = c(4, #bottom
            4, #left
            3, #top
            2),
    cex.main = 0.8,
    family = "sans" #right
)



# Update the labels for Position and Estrus_cycle
df$Position <- factor(df$Position, labels = c("Contralateral", "Ipsilateral"))
df$Estrus_cycle <- factor(df$Estrus_cycle, labels = c("Follicular phase", "Luteal phase"))

# Convert Animal_ID to a factor
df$Animal_ID <- as.factor(df$Animal_ID)

# Create a combined factor variable for Estrus_cycle and Cellular
df$Combined <- factor(paste(df$Estrus_cycle, df$Cellular, sep = " "),
                      levels = c("Follicular phase Native", "Follicular phase Decellularized",
                                 "Luteal phase Native", "Luteal phase Decellularized"))

# Create X_position_group based on X_position values
df$X_position_group <- ifelse(df$X_position <= 750, "Medulla", "Cortex")
df$X_position_group <- factor(df$X_position_group, levels = c("Cortex", "Medulla"))

# Define fill colors for each factor level
fill_colors <- c("Follicular phase Native" = viridis_pal(option = "D")(1),
                 "Follicular phase Decellularized" = alpha(viridis_pal(option = "D")(1), 0.2),
                 "Luteal phase Native" = "#238A8DFF",
                 "Luteal phase Decellularized" = alpha("#238A8DFF", 0.2))

# Create the plot using ggplot
ggplot(data = df, aes(x = X_position_group, y = YM, fill = Combined)) +
  geom_boxplot(width = 0.7, outlier.shape = NA) +
  geom_jitter(position = position_jitterdodge(), alpha = 0.8) +
  xlab("") +
  ylab("Young's modulus (Pa)") +
  ggtitle("Boxplot and Individual Data Points of YM by Estrus_cycle, Cellular, Position, and X_position_
```

```
  scale_fill_manual(values = fill_colors, guide = guide_legend(title = "Combined")) +
  theme_bw() +
  ylim(0, 30000) +
  facet_grid(Estrus_cycle ~ Position, space = "free_x", scales = "free_x") +
  scale_x_discrete(labels = c("Cortex", "Medulla"), expand = c(0, 0)) +
  theme(panel.grid.major = element_blank(), panel.grid.minor = element_blank(),
        axis.text.x = element_text(size = 11, angle = 0, hjust = 0.5),
        axis.text.y = element_text(size = 12, angle = 0, hjust = 0.5),
        axis.title.x = element_text(size = 18, face = "bold"),
        axis.title.y = element_text(size = 16, face = "bold"),
        legend.position = "right",
        strip.text = element_text(size = 14, face = "bold"))  # Increase the size of facet labels

dev.off()
```

```
## pdf
##   2
```

```
# Stats

# Read the dataset from a CSV file
df <- read.csv("Ovary_YM.csv")

# Create X_position_group based on X_position values
df$X_position_group <- ifelse(df$X_position <= 750, "Medulla", "Cortex")
# Convert X_position_group to factor
df$X_position_group <- factor(df$X_position_group, labels = c("Cortex", "Medulla"))

# Update the labels for Position, Estrus_cycle, and Cellular
df$Position <- factor(df$Position, labels = c("Contralateral", "Ipsilateral"))
df$Estrus_cycle <- factor(df$Estrus_cycle, labels = c("Follicular phase", "Luteal phase"))
df$Cellular <- factor(df$Cellular, labels = c("Native", "Decellularized"))

# Create a factor variable for the comparisons
df$Comparison <- factor(paste(df$X_position_group, df$Estrus_cycle, df$Position, df$Cellular, sep = " -

# Perform pairwise comparisons using Tukey's method
model <- lm(YM ~ Comparison, data = df)
comp <- glht(model, linfct = mcp(Comparison = "Tukey"))

# Summarize the results
summary(comp)
```

```
## Warning in RET$pfunction("adjusted", ...): Completion with error > abseps

## Warning in RET$pfunction("adjusted", ...): Completion with error > abseps

## Warning in RET$pfunction("adjusted", ...): Completion with error > abseps

## Warning in RET$pfunction("adjusted", ...): Completion with error > abseps

## Warning in RET$pfunction("adjusted", ...): Completion with error > abseps

## Warning in RET$pfunction("adjusted", ...): Completion with error > abseps
```

```
## Warning in RET$pfunction("adjusted", ...): Completion with error > abseps
## Warning in RET$pfunction("adjusted", ...): Completion with error > abseps
## Warning in RET$pfunction("adjusted", ...): Completion with error > abseps
## Warning in RET$pfunction("adjusted", ...): Completion with error > abseps
## Warning in RET$pfunction("adjusted", ...): Completion with error > abseps
## Warning in RET$pfunction("adjusted", ...): Completion with error > abseps
## Warning in RET$pfunction("adjusted", ...): Completion with error > abseps
## Warning in RET$pfunction("adjusted", ...): Completion with error > abseps
## Warning in RET$pfunction("adjusted", ...): Completion with error > abseps
## Warning in RET$pfunction("adjusted", ...): Completion with error > abseps
## Warning in RET$pfunction("adjusted", ...): Completion with error > abseps
## Warning in RET$pfunction("adjusted", ...): Completion with error > abseps
## Warning in RET$pfunction("adjusted", ...): Completion with error > abseps
## Warning in RET$pfunction("adjusted", ...): Completion with error > abseps
## Warning in RET$pfunction("adjusted", ...): Completion with error > abseps
## Warning in RET$pfunction("adjusted", ...): Completion with error > abseps
## Warning in RET$pfunction("adjusted", ...): Completion with error > abseps
## Warning in RET$pfunction("adjusted", ...): Completion with error > abseps
## Warning in RET$pfunction("adjusted", ...): Completion with error > abseps
## Warning in RET$pfunction("adjusted", ...): Completion with error > abseps
## Warning in RET$pfunction("adjusted", ...): Completion with error > abseps
## Warning in RET$pfunction("adjusted", ...): Completion with error > abseps
## Warning in RET$pfunction("adjusted", ...): Completion with error > abseps
## Warning in RET$pfunction("adjusted", ...): Completion with error > abseps
## Warning in RET$pfunction("adjusted", ...): Completion with error > abseps
## Warning in RET$pfunction("adjusted", ...): Completion with error > abseps
## Warning in RET$pfunction("adjusted", ...): Completion with error > abseps
```

```
## Warning in RET$pfunction("adjusted", ...): Completion with error > abseps

## Warning in RET$pfunction("adjusted", ...): Completion with error > abseps

## Warning in RET$pfunction("adjusted", ...): Completion with error > abseps

## Warning in RET$pfunction("adjusted", ...): Completion with error > abseps

## Warning in RET$pfunction("adjusted", ...): Completion with error > abseps

## Warning in RET$pfunction("adjusted", ...): Completion with error > abseps
##
##   Simultaneous Tests for General Linear Hypotheses
##
## Multiple Comparisons of Means: Tukey Contrasts
##
##
## Fit: lm(formula = YM ~ Comparison, data = df)
##
## Linear Hypotheses:
##
## Cortex - Follicular phase - Contralateral - Native - Cortex - Follicular phase - Contralateral - Dec
## Cortex - Follicular phase - Ipsilateral - Decellularized - Cortex - Follicular phase - Contralateral
## Cortex - Follicular phase - Ipsilateral - Native - Cortex - Follicular phase - Contralateral - Decel
## Cortex - Luteal phase - Contralateral - Decellularized - Cortex - Follicular phase - Contralateral -
## Cortex - Luteal phase - Contralateral - Native - Cortex - Follicular phase - Contralateral - Decellu
## Cortex - Luteal phase - Ipsilateral - Decellularized - Cortex - Follicular phase - Contralateral - D
## Cortex - Luteal phase - Ipsilateral - Native - Cortex - Follicular phase - Contralateral - Decellula
## Medulla - Follicular phase - Contralateral - Decellularized - Cortex - Follicular phase - Contralate
## Medulla - Follicular phase - Contralateral - Native - Cortex - Follicular phase - Contralateral - De
## Medulla - Follicular phase - Ipsilateral - Decellularized - Cortex - Follicular phase - Contralatera
## Medulla - Follicular phase - Ipsilateral - Native - Cortex - Follicular phase - Contralateral - Dece
## Medulla - Luteal phase - Contralateral - Decellularized - Cortex - Follicular phase - Contralateral
## Medulla - Luteal phase - Contralateral - Native - Cortex - Follicular phase - Contralateral - Decell
## Medulla - Luteal phase - Ipsilateral - Decellularized - Cortex - Follicular phase - Contralateral -
## Medulla - Luteal phase - Ipsilateral - Native - Cortex - Follicular phase - Contralateral - Decellul
## Cortex - Follicular phase - Ipsilateral - Decellularized - Cortex - Follicular phase - Contralateral
## Cortex - Follicular phase - Ipsilateral - Native - Cortex - Follicular phase - Contralateral - Nativ
## Cortex - Luteal phase - Contralateral - Decellularized - Cortex - Follicular phase - Contralateral -
## Cortex - Luteal phase - Contralateral - Native - Cortex - Follicular phase - Contralateral - Native
## Cortex - Luteal phase - Ipsilateral - Decellularized - Cortex - Follicular phase - Contralateral - N
## Cortex - Luteal phase - Ipsilateral - Native - Cortex - Follicular phase - Contralateral - Native ==
## Medulla - Follicular phase - Contralateral - Decellularized - Cortex - Follicular phase - Contralate
## Medulla - Follicular phase - Contralateral - Native - Cortex - Follicular phase - Contralateral - Na
## Medulla - Follicular phase - Ipsilateral - Decellularized - Cortex - Follicular phase - Contralatera
## Medulla - Follicular phase - Ipsilateral - Native - Cortex - Follicular phase - Contralateral - Nati
## Medulla - Luteal phase - Contralateral - Decellularized - Cortex - Follicular phase - Contralateral
## Medulla - Luteal phase - Contralateral - Native - Cortex - Follicular phase - Contralateral - Native
## Medulla - Luteal phase - Ipsilateral - Decellularized - Cortex - Follicular phase - Contralateral -
## Medulla - Luteal phase - Ipsilateral - Native - Cortex - Follicular phase - Contralateral - Native =
## Cortex - Follicular phase - Ipsilateral - Native - Cortex - Follicular phase - Ipsilateral - Decellu
## Cortex - Luteal phase - Contralateral - Decellularized - Cortex - Follicular phase - Ipsilateral - De
## Cortex - Luteal phase - Contralateral - Native - Cortex - Follicular phase - Ipsilateral - Decellula
```

```
## Cortex - Luteal phase - Ipsilateral - Decellularized - Cortex - Follicular phase - Ipsilateral - Dec
## Cortex - Luteal phase - Ipsilateral - Native - Cortex - Follicular phase - Ipsilateral - Decellulari
## Medulla - Follicular phase - Contralateral - Decellularized - Cortex - Follicular phase - Ipsilateral
## Medulla - Follicular phase - Contralateral - Native - Cortex - Follicular phase - Ipsilateral - Decel
## Medulla - Follicular phase - Ipsilateral - Decellularized - Cortex - Follicular phase - Ipsilateral -
## Medulla - Follicular phase - Ipsilateral - Native - Cortex - Follicular phase - Ipsilateral - Decellu
## Medulla - Luteal phase - Contralateral - Decellularized - Cortex - Follicular phase - Ipsilateral -
## Medulla - Luteal phase - Contralateral - Native - Cortex - Follicular phase - Ipsilateral - Decellula
## Medulla - Luteal phase - Ipsilateral - Decellularized - Cortex - Follicular phase - Ipsilateral - Dec
## Medulla - Luteal phase - Ipsilateral - Native - Cortex - Follicular phase - Ipsilateral - Decellular
## Cortex - Luteal phase - Contralateral - Decellularized - Cortex - Follicular phase - Ipsilateral - N
## Cortex - Luteal phase - Contralateral - Native - Cortex - Follicular phase - Ipsilateral - Native ==
## Cortex - Luteal phase - Ipsilateral - Decellularized - Cortex - Follicular phase - Ipsilateral - Nat
## Cortex - Luteal phase - Ipsilateral - Native - Cortex - Follicular phase - Ipsilateral - Native == 0
## Medulla - Follicular phase - Contralateral - Decellularized - Cortex - Follicular phase - Ipsilateral
## Medulla - Follicular phase - Contralateral - Native - Cortex - Follicular phase - Ipsilateral - Nati
## Medulla - Follicular phase - Ipsilateral - Decellularized - Cortex - Follicular phase - Ipsilateral -
## Medulla - Follicular phase - Ipsilateral - Native - Cortex - Follicular phase - Ipsilateral - Native
## Medulla - Luteal phase - Contralateral - Decellularized - Cortex - Follicular phase - Ipsilateral -
## Medulla - Luteal phase - Contralateral - Native - Cortex - Follicular phase - Ipsilateral - Native =
## Medulla - Luteal phase - Ipsilateral - Decellularized - Cortex - Follicular phase - Ipsilateral - Na
## Medulla - Luteal phase - Ipsilateral - Native - Cortex - Follicular phase - Ipsilateral - Native ==
## Cortex - Luteal phase - Contralateral - Native - Cortex - Luteal phase - Contralateral - Decellulari
## Cortex - Luteal phase - Ipsilateral - Decellularized - Cortex - Luteal phase - Contralateral - Decel
## Cortex - Luteal phase - Ipsilateral - Native - Cortex - Luteal phase - Contralateral - Decellularize
## Medulla - Follicular phase - Contralateral - Decellularized - Cortex - Luteal phase - Contralateral -
## Medulla - Follicular phase - Contralateral - Native - Cortex - Luteal phase - Contralateral - Decellu
## Medulla - Follicular phase - Ipsilateral - Decellularized - Cortex - Luteal phase - Contralateral -
## Medulla - Follicular phase - Ipsilateral - Native - Cortex - Luteal phase - Contralateral - Decellula
## Medulla - Luteal phase - Contralateral - Decellularized - Cortex - Luteal phase - Contralateral - Dec
## Medulla - Luteal phase - Contralateral - Native - Cortex - Luteal phase - Contralateral - Decellular
## Medulla - Luteal phase - Ipsilateral - Decellularized - Cortex - Luteal phase - Contralateral - Decel
## Medulla - Luteal phase - Ipsilateral - Native - Cortex - Luteal phase - Contralateral - Decellulariz
## Cortex - Luteal phase - Ipsilateral - Decellularized - Cortex - Luteal phase - Contralateral - Native
## Cortex - Luteal phase - Ipsilateral - Native - Cortex - Luteal phase - Contralateral - Native == 0
## Medulla - Follicular phase - Contralateral - Decellularized - Cortex - Luteal phase - Contralateral -
## Medulla - Follicular phase - Contralateral - Native - Cortex - Luteal phase - Contralateral - Native
## Medulla - Follicular phase - Ipsilateral - Decellularized - Cortex - Luteal phase - Contralateral -
## Medulla - Follicular phase - Ipsilateral - Native - Cortex - Luteal phase - Contralateral - Native =
## Medulla - Luteal phase - Contralateral - Decellularized - Cortex - Luteal phase - Contralateral - Na
## Medulla - Luteal phase - Contralateral - Native - Cortex - Luteal phase - Contralateral - Native ==
## Medulla - Luteal phase - Ipsilateral - Decellularized - Cortex - Luteal phase - Contralateral - Nati
## Medulla - Luteal phase - Ipsilateral - Native - Cortex - Luteal phase - Contralateral - Native == 0
## Cortex - Luteal phase - Ipsilateral - Native - Cortex - Luteal phase - Ipsilateral - Decellularized =
## Medulla - Follicular phase - Contralateral - Decellularized - Cortex - Luteal phase - Ipsilateral -
## Medulla - Follicular phase - Contralateral - Native - Cortex - Luteal phase - Ipsilateral - Decellula
## Medulla - Follicular phase - Ipsilateral - Decellularized - Cortex - Luteal phase - Ipsilateral - Dec
## Medulla - Follicular phase - Ipsilateral - Native - Cortex - Luteal phase - Ipsilateral - Decellular
## Medulla - Luteal phase - Contralateral - Decellularized - Cortex - Luteal phase - Ipsilateral - Decel
## Medulla - Luteal phase - Contralateral - Native - Cortex - Luteal phase - Ipsilateral - Decellularize
## Medulla - Luteal phase - Ipsilateral - Decellularized - Cortex - Luteal phase - Ipsilateral - Decellu
## Medulla - Luteal phase - Ipsilateral - Native - Cortex - Luteal phase - Ipsilateral - Decellularized
## Medulla - Follicular phase - Contralateral - Decellularized - Cortex - Luteal phase - Ipsilateral -
## Medulla - Follicular phase - Contralateral - Native - Cortex - Luteal phase - Ipsilateral - Native =
```

```
## Medulla - Follicular phase - Ipsilateral - Decellularized - Cortex - Luteal phase - Ipsilateral - Nat
## Medulla - Follicular phase - Ipsilateral - Native - Cortex - Luteal phase - Ipsilateral - Native == 0
## Medulla - Luteal phase - Contralateral - Decellularized - Cortex - Luteal phase - Ipsilateral - Nativ
## Medulla - Luteal phase - Contralateral - Native - Cortex - Luteal phase - Ipsilateral - Native == 0
## Medulla - Luteal phase - Ipsilateral - Decellularized - Cortex - Luteal phase - Ipsilateral - Native
## Medulla - Luteal phase - Ipsilateral - Native - Cortex - Luteal phase - Ipsilateral - Native == 0
## Medulla - Follicular phase - Contralateral - Native - Medulla - Follicular phase - Contralateral - De
## Medulla - Follicular phase - Ipsilateral - Decellularized - Medulla - Follicular phase - Contralatera
## Medulla - Follicular phase - Ipsilateral - Native - Medulla - Follicular phase - Contralateral - Dec
## Medulla - Luteal phase - Contralateral - Decellularized - Medulla - Follicular phase - Contralateral
## Medulla - Luteal phase - Contralateral - Native - Medulla - Follicular phase - Contralateral - Decel
## Medulla - Luteal phase - Ipsilateral - Decellularized - Medulla - Follicular phase - Contralateral -
## Medulla - Luteal phase - Ipsilateral - Native - Medulla - Follicular phase - Contralateral - Decellu
## Medulla - Follicular phase - Ipsilateral - Decellularized - Medulla - Follicular phase - Contralatera
## Medulla - Follicular phase - Ipsilateral - Native - Medulla - Follicular phase - Contralateral - Nat
## Medulla - Luteal phase - Contralateral - Decellularized - Medulla - Follicular phase - Contralateral
## Medulla - Luteal phase - Contralateral - Native - Medulla - Follicular phase - Contralateral - Nativ
## Medulla - Luteal phase - Ipsilateral - Decellularized - Medulla - Follicular phase - Contralateral -
## Medulla - Luteal phase - Ipsilateral - Native - Medulla - Follicular phase - Contralateral - Native
## Medulla - Follicular phase - Ipsilateral - Native - Medulla - Follicular phase - Ipsilateral - Decel
## Medulla - Luteal phase - Contralateral - Decellularized - Medulla - Follicular phase - Ipsilateral -
## Medulla - Luteal phase - Contralateral - Native - Medulla - Follicular phase - Ipsilateral - Decellu
## Medulla - Luteal phase - Ipsilateral - Decellularized - Medulla - Follicular phase - Ipsilateral - De
## Medulla - Luteal phase - Ipsilateral - Native - Medulla - Follicular phase - Ipsilateral - Decellula
## Medulla - Luteal phase - Contralateral - Decellularized - Medulla - Follicular phase - Ipsilateral -
## Medulla - Luteal phase - Contralateral - Native - Medulla - Follicular phase - Ipsilateral - Native
## Medulla - Luteal phase - Ipsilateral - Decellularized - Medulla - Follicular phase - Ipsilateral - Na
## Medulla - Luteal phase - Ipsilateral - Native - Medulla - Follicular phase - Ipsilateral - Native ==
## Medulla - Luteal phase - Contralateral - Native - Medulla - Luteal phase - Contralateral - Decellula
## Medulla - Luteal phase - Ipsilateral - Decellularized - Medulla - Luteal phase - Contralateral - Dec
## Medulla - Luteal phase - Ipsilateral - Native - Medulla - Luteal phase - Contralateral - Decellulari
## Medulla - Luteal phase - Ipsilateral - Decellularized - Medulla - Luteal phase - Contralateral - Nat
## Medulla - Luteal phase - Ipsilateral - Native - Medulla - Luteal phase - Contralateral - Native == 0
## Medulla - Luteal phase - Ipsilateral - Native - Medulla - Luteal phase - Ipsilateral - Decellularize
##
## Cortex - Follicular phase - Contralateral - Native - Cortex - Follicular phase - Contralateral - Dec
## Cortex - Follicular phase - Ipsilateral - Decellularized - Cortex - Follicular phase - Contralateral
## Cortex - Follicular phase - Ipsilateral - Native - Cortex - Follicular phase - Contralateral - Decel
## Cortex - Luteal phase - Contralateral - Decellularized - Cortex - Follicular phase - Contralateral -
## Cortex - Luteal phase - Contralateral - Native - Cortex - Follicular phase - Contralateral - Decellu
## Cortex - Luteal phase - Ipsilateral - Decellularized - Cortex - Follicular phase - Contralateral - De
## Cortex - Luteal phase - Ipsilateral - Native - Cortex - Follicular phase - Contralateral - Decellula
## Medulla - Follicular phase - Contralateral - Decellularized - Cortex - Follicular phase - Contralate
## Medulla - Follicular phase - Contralateral - Native - Cortex - Follicular phase - Contralateral - De
## Medulla - Follicular phase - Ipsilateral - Decellularized - Cortex - Follicular phase - Contralateral
## Medulla - Follicular phase - Ipsilateral - Native - Cortex - Follicular phase - Contralateral - Decel
## Medulla - Luteal phase - Contralateral - Decellularized - Cortex - Follicular phase - Contralateral -
## Medulla - Luteal phase - Contralateral - Native - Cortex - Follicular phase - Contralateral - Decellu
## Medulla - Luteal phase - Ipsilateral - Decellularized - Cortex - Follicular phase - Contralateral - N
## Medulla - Luteal phase - Ipsilateral - Native - Cortex - Follicular phase - Contralateral - Decellula
## Cortex - Follicular phase - Ipsilateral - Decellularized - Cortex - Follicular phase - Contralateral
## Cortex - Follicular phase - Ipsilateral - Native - Cortex - Follicular phase - Contralateral - Nativ
## Cortex - Luteal phase - Contralateral - Decellularized - Cortex - Follicular phase - Contralateral -
## Cortex - Luteal phase - Contralateral - Native - Cortex - Follicular phase - Contralateral - Native
```

```
## Cortex - Luteal phase - Ipsilateral - Decellularized - Cortex - Follicular phase - Contralateral - Na
## Cortex - Luteal phase - Ipsilateral - Native - Cortex - Follicular phase - Contralateral - Native ==
## Medulla - Follicular phase - Contralateral - Decellularized - Cortex - Follicular phase - Contralate
## Medulla - Follicular phase - Contralateral - Native - Cortex - Follicular phase - Contralateral - Na
## Medulla - Follicular phase - Ipsilateral - Decellularized - Cortex - Follicular phase - Contralatera
## Medulla - Follicular phase - Ipsilateral - Native - Cortex - Follicular phase - Contralateral - Nati
## Medulla - Luteal phase - Contralateral - Decellularized - Cortex - Follicular phase - Contralateral
## Medulla - Luteal phase - Contralateral - Native - Cortex - Follicular phase - Contralateral - Native
## Medulla - Luteal phase - Ipsilateral - Decellularized - Cortex - Follicular phase - Contralateral -
## Medulla - Luteal phase - Ipsilateral - Native - Cortex - Follicular phase - Contralateral - Native =
## Cortex - Follicular phase - Ipsilateral - Native - Cortex - Follicular phase - Ipsilateral - Decellu
## Cortex - Luteal phase - Contralateral - Decellularized - Cortex - Follicular phase - Ipsilateral - D
## Cortex - Luteal phase - Contralateral - Native - Cortex - Follicular phase - Ipsilateral - Decellula
## Cortex - Luteal phase - Ipsilateral - Decellularized - Cortex - Follicular phase - Ipsilateral - Dec
## Cortex - Luteal phase - Ipsilateral - Native - Cortex - Follicular phase - Ipsilateral - Decellulari
## Medulla - Follicular phase - Contralateral - Decellularized - Cortex - Follicular phase - Ipsilatera
## Medulla - Follicular phase - Contralateral - Native - Cortex - Follicular phase - Ipsilateral - Dece
## Medulla - Follicular phase - Ipsilateral - Decellularized - Cortex - Follicular phase - Ipsilateral
## Medulla - Follicular phase - Ipsilateral - Native - Cortex - Follicular phase - Ipsilateral - Decell
## Medulla - Luteal phase - Contralateral - Decellularized - Cortex - Follicular phase - Ipsilateral -
## Medulla - Luteal phase - Contralateral - Native - Cortex - Follicular phase - Ipsilateral - Decellul
## Medulla - Luteal phase - Ipsilateral - Decellularized - Cortex - Follicular phase - Ipsilateral - De
## Medulla - Luteal phase - Ipsilateral - Native - Cortex - Follicular phase - Ipsilateral - Decellular
## Cortex - Luteal phase - Contralateral - Decellularized - Cortex - Follicular phase - Ipsilateral - Na
## Cortex - Luteal phase - Contralateral - Native - Cortex - Follicular phase - Ipsilateral - Native ==
## Cortex - Luteal phase - Ipsilateral - Decellularized - Cortex - Follicular phase - Ipsilateral - Nati
## Cortex - Luteal phase - Ipsilateral - Native - Cortex - Follicular phase - Ipsilateral - Native == 0
## Medulla - Follicular phase - Contralateral - Decellularized - Cortex - Follicular phase - Ipsilatera
## Medulla - Follicular phase - Contralateral - Native - Cortex - Follicular phase - Ipsilateral - Nati
## Medulla - Follicular phase - Ipsilateral - Decellularized - Cortex - Follicular phase - Ipsilateral
## Medulla - Follicular phase - Ipsilateral - Native - Cortex - Follicular phase - Ipsilateral - Native
## Medulla - Luteal phase - Contralateral - Decellularized - Cortex - Follicular phase - Ipsilateral -
## Medulla - Luteal phase - Contralateral - Native - Cortex - Follicular phase - Ipsilateral - Native =
## Medulla - Luteal phase - Ipsilateral - Decellularized - Cortex - Follicular phase - Ipsilateral - Na
## Medulla - Luteal phase - Ipsilateral - Native - Cortex - Follicular phase - Ipsilateral - Native ==
## Cortex - Luteal phase - Contralateral - Native - Cortex - Luteal phase - Contralateral - Decellulari
## Cortex - Luteal phase - Ipsilateral - Decellularized - Cortex - Luteal phase - Contralateral - Decel
## Cortex - Luteal phase - Ipsilateral - Native - Cortex - Luteal phase - Contralateral - Decellularize
## Medulla - Follicular phase - Contralateral - Decellularized - Cortex - Luteal phase - Contralateral
## Medulla - Follicular phase - Contralateral - Native - Cortex - Luteal phase - Contralateral - Decellu
## Medulla - Follicular phase - Ipsilateral - Decellularized - Cortex - Luteal phase - Contralateral -
## Medulla - Follicular phase - Ipsilateral - Native - Cortex - Luteal phase - Contralateral - Decellula
## Medulla - Luteal phase - Contralateral - Decellularized - Cortex - Luteal phase - Contralateral - De
## Medulla - Luteal phase - Contralateral - Native - Cortex - Luteal phase - Contralateral - Decellular
## Medulla - Luteal phase - Ipsilateral - Decellularized - Cortex - Luteal phase - Contralateral - Decel
## Medulla - Luteal phase - Ipsilateral - Native - Cortex - Luteal phase - Contralateral - Decellulariz
## Cortex - Luteal phase - Ipsilateral - Decellularized - Cortex - Luteal phase - Contralateral - Nativ
## Cortex - Luteal phase - Ipsilateral - Native - Cortex - Luteal phase - Contralateral - Native == 0
## Medulla - Follicular phase - Contralateral - Decellularized - Cortex - Luteal phase - Contralateral
## Medulla - Follicular phase - Contralateral - Native - Cortex - Luteal phase - Contralateral - Native
## Medulla - Follicular phase - Ipsilateral - Decellularized - Cortex - Luteal phase - Contralateral -
## Medulla - Follicular phase - Ipsilateral - Native - Cortex - Luteal phase - Contralateral - Native =
## Medulla - Luteal phase - Contralateral - Decellularized - Cortex - Luteal phase - Contralateral - Na
## Medulla - Luteal phase - Contralateral - Native - Cortex - Luteal phase - Contralateral - Native ==
```

```
## Medulla - Luteal phase - Ipsilateral - Decellularized - Cortex - Luteal phase - Contralateral - Nativ
## Medulla - Luteal phase - Ipsilateral - Native - Cortex - Luteal phase - Contralateral - Native == 0
## Cortex - Luteal phase - Ipsilateral - Native - Cortex - Luteal phase - Ipsilateral - Decellularized =
## Medulla - Follicular phase - Contralateral - Decellularized - Cortex - Luteal phase - Ipsilateral - D
## Medulla - Follicular phase - Contralateral - Native - Cortex - Luteal phase - Ipsilateral - Decellula
## Medulla - Follicular phase - Ipsilateral - Decellularized - Cortex - Luteal phase - Ipsilateral - Dec
## Medulla - Follicular phase - Ipsilateral - Native - Cortex - Luteal phase - Ipsilateral - Decellulari
## Medulla - Luteal phase - Contralateral - Decellularized - Cortex - Luteal phase - Ipsilateral - Decel
## Medulla - Luteal phase - Contralateral - Native - Cortex - Luteal phase - Ipsilateral - Decellularize
## Medulla - Luteal phase - Ipsilateral - Decellularized - Cortex - Luteal phase - Ipsilateral - Decellu
## Medulla - Luteal phase - Ipsilateral - Native - Cortex - Luteal phase - Ipsilateral - Decellularized
## Medulla - Follicular phase - Contralateral - Decellularized - Cortex - Luteal phase - Ipsilateral - N
## Medulla - Follicular phase - Contralateral - Native - Cortex - Luteal phase - Ipsilateral - Native =
## Medulla - Follicular phase - Ipsilateral - Decellularized - Cortex - Luteal phase - Ipsilateral - Nat
## Medulla - Follicular phase - Ipsilateral - Native - Cortex - Luteal phase - Ipsilateral - Native == 0
## Medulla - Luteal phase - Contralateral - Decellularized - Cortex - Luteal phase - Ipsilateral - Nativ
## Medulla - Luteal phase - Contralateral - Native - Cortex - Luteal phase - Ipsilateral - Native == 0
## Medulla - Luteal phase - Ipsilateral - Decellularized - Cortex - Luteal phase - Ipsilateral - Native
## Medulla - Luteal phase - Ipsilateral - Native - Cortex - Luteal phase - Ipsilateral - Native == 0
## Medulla - Follicular phase - Contralateral - Native - Medulla - Follicular phase - Contralateral - De
## Medulla - Follicular phase - Ipsilateral - Decellularized - Medulla - Follicular phase - Contralatera
## Medulla - Follicular phase - Ipsilateral - Native - Medulla - Follicular phase - Contralateral - Dece
## Medulla - Luteal phase - Contralateral - Decellularized - Medulla - Follicular phase - Contralateral
## Medulla - Luteal phase - Contralateral - Native - Medulla - Follicular phase - Contralateral - Decel
## Medulla - Luteal phase - Ipsilateral - Decellularized - Medulla - Follicular phase - Contralateral -
## Medulla - Luteal phase - Ipsilateral - Native - Medulla - Follicular phase - Contralateral - Decellul
## Medulla - Follicular phase - Ipsilateral - Decellularized - Medulla - Follicular phase - Contralatera
## Medulla - Follicular phase - Ipsilateral - Native - Medulla - Follicular phase - Contralateral - Nati
## Medulla - Luteal phase - Contralateral - Decellularized - Medulla - Follicular phase - Contralateral
## Medulla - Luteal phase - Contralateral - Native - Medulla - Follicular phase - Contralateral - Native
## Medulla - Luteal phase - Ipsilateral - Decellularized - Medulla - Follicular phase - Contralateral -
## Medulla - Luteal phase - Ipsilateral - Native - Medulla - Follicular phase - Contralateral - Native =
## Medulla - Follicular phase - Ipsilateral - Native - Medulla - Follicular phase - Ipsilateral - Decell
## Medulla - Luteal phase - Contralateral - Decellularized - Medulla - Follicular phase - Ipsilateral -
## Medulla - Luteal phase - Contralateral - Native - Medulla - Follicular phase - Ipsilateral - Decellul
## Medulla - Luteal phase - Ipsilateral - Decellularized - Medulla - Follicular phase - Ipsilateral - De
## Medulla - Luteal phase - Ipsilateral - Native - Medulla - Follicular phase - Ipsilateral - Decellula
## Medulla - Luteal phase - Contralateral - Decellularized - Medulla - Follicular phase - Ipsilateral -
## Medulla - Luteal phase - Contralateral - Native - Medulla - Follicular phase - Ipsilateral - Native =
## Medulla - Luteal phase - Ipsilateral - Decellularized - Medulla - Follicular phase - Ipsilateral - Na
## Medulla - Luteal phase - Ipsilateral - Native - Medulla - Follicular phase - Ipsilateral - Native ==
## Medulla - Luteal phase - Contralateral - Native - Medulla - Luteal phase - Contralateral - Decellular
## Medulla - Luteal phase - Ipsilateral - Decellularized - Medulla - Luteal phase - Contralateral - Dece
## Medulla - Luteal phase - Ipsilateral - Native - Medulla - Luteal phase - Contralateral - Decellulari
## Medulla - Luteal phase - Ipsilateral - Decellularized - Medulla - Luteal phase - Contralateral - Nati
## Medulla - Luteal phase - Ipsilateral - Native - Medulla - Luteal phase - Contralateral - Native == 0
## Medulla - Luteal phase - Ipsilateral - Native - Medulla - Luteal phase - Ipsilateral - Decellularized
##
## Cortex - Follicular phase - Contralateral - Native - Cortex - Follicular phase - Contralateral - Dece
## Cortex - Follicular phase - Ipsilateral - Decellularized - Cortex - Follicular phase - Contralateral
## Cortex - Follicular phase - Ipsilateral - Native - Cortex - Follicular phase - Contralateral - Decell
## Cortex - Luteal phase - Contralateral - Decellularized - Cortex - Follicular phase - Contralateral -
## Cortex - Luteal phase - Contralateral - Native - Cortex - Follicular phase - Contralateral - Decellul
## Cortex - Luteal phase - Ipsilateral - Decellularized - Cortex - Follicular phase - Contralateral - De
```

```
## Cortex - Luteal phase - Ipsilateral - Native - Cortex - Follicular phase - Contralateral - Decellula
## Medulla - Follicular phase - Contralateral - Decellularized - Cortex - Follicular phase - Contralate
## Medulla - Follicular phase - Contralateral - Native - Cortex - Follicular phase - Contralateral - De
## Medulla - Follicular phase - Ipsilateral - Decellularized - Cortex - Follicular phase - Contralatera
## Medulla - Follicular phase - Ipsilateral - Native - Cortex - Follicular phase - Contralateral - Dece
## Medulla - Luteal phase - Contralateral - Decellularized - Cortex - Follicular phase - Contralateral
## Medulla - Luteal phase - Contralateral - Native - Cortex - Follicular phase - Contralateral - Decell
## Medulla - Luteal phase - Ipsilateral - Decellularized - Cortex - Follicular phase - Contralateral -
## Medulla - Luteal phase - Ipsilateral - Native - Cortex - Follicular phase - Contralateral - Decellul
## Cortex - Follicular phase - Ipsilateral - Decellularized - Cortex - Follicular phase - Contralateral
## Cortex - Follicular phase - Ipsilateral - Native - Cortex - Follicular phase - Contralateral - Nativ
## Cortex - Luteal phase - Contralateral - Decellularized - Cortex - Follicular phase - Contralateral -
## Cortex - Luteal phase - Contralateral - Native - Cortex - Follicular phase - Contralateral - Native
## Cortex - Luteal phase - Ipsilateral - Decellularized - Cortex - Follicular phase - Contralateral - N
## Cortex - Luteal phase - Ipsilateral - Native - Cortex - Follicular phase - Contralateral - Native ==
## Medulla - Follicular phase - Contralateral - Decellularized - Cortex - Follicular phase - Contralate
## Medulla - Follicular phase - Contralateral - Native - Cortex - Follicular phase - Contralateral - Na
## Medulla - Follicular phase - Ipsilateral - Decellularized - Cortex - Follicular phase - Contralatera
## Medulla - Follicular phase - Ipsilateral - Native - Cortex - Follicular phase - Contralateral - Nati
## Medulla - Luteal phase - Contralateral - Decellularized - Cortex - Follicular phase - Contralateral
## Medulla - Luteal phase - Contralateral - Native - Cortex - Follicular phase - Contralateral - Native
## Medulla - Luteal phase - Ipsilateral - Decellularized - Cortex - Follicular phase - Contralateral -
## Medulla - Luteal phase - Ipsilateral - Native - Cortex - Follicular phase - Contralateral - Native =
## Cortex - Follicular phase - Ipsilateral - Native - Cortex - Follicular phase - Ipsilateral - Decellu
## Cortex - Luteal phase - Contralateral - Decellularized - Cortex - Follicular phase - Ipsilateral - D
## Cortex - Luteal phase - Contralateral - Native - Cortex - Follicular phase - Ipsilateral - Decellula
## Cortex - Luteal phase - Ipsilateral - Decellularized - Cortex - Follicular phase - Ipsilateral - Dec
## Cortex - Luteal phase - Ipsilateral - Native - Cortex - Follicular phase - Ipsilateral - Decellulari
## Medulla - Follicular phase - Contralateral - Decellularized - Cortex - Follicular phase - Ipsilateral
## Medulla - Follicular phase - Contralateral - Native - Cortex - Follicular phase - Ipsilateral - Decel
## Medulla - Follicular phase - Ipsilateral - Decellularized - Cortex - Follicular phase - Ipsilateral
## Medulla - Follicular phase - Ipsilateral - Native - Cortex - Follicular phase - Ipsilateral - Decell
## Medulla - Luteal phase - Contralateral - Decellularized - Cortex - Follicular phase - Ipsilateral -
## Medulla - Luteal phase - Contralateral - Native - Cortex - Follicular phase - Ipsilateral - Decellul
## Medulla - Luteal phase - Ipsilateral - Decellularized - Cortex - Follicular phase - Ipsilateral - De
## Medulla - Luteal phase - Ipsilateral - Native - Cortex - Follicular phase - Ipsilateral - Decellular
## Cortex - Luteal phase - Contralateral - Decellularized - Cortex - Follicular phase - Ipsilateral - N
## Cortex - Luteal phase - Contralateral - Native - Cortex - Follicular phase - Ipsilateral - Native ==
## Cortex - Luteal phase - Ipsilateral - Decellularized - Cortex - Follicular phase - Ipsilateral - Nat
## Cortex - Luteal phase - Ipsilateral - Native - Cortex - Follicular phase - Ipsilateral - Native == 0
## Medulla - Follicular phase - Contralateral - Decellularized - Cortex - Follicular phase - Ipsilatera
## Medulla - Follicular phase - Contralateral - Native - Cortex - Follicular phase - Ipsilateral - Nati
## Medulla - Follicular phase - Ipsilateral - Decellularized - Cortex - Follicular phase - Ipsilateral
## Medulla - Follicular phase - Ipsilateral - Native - Cortex - Follicular phase - Ipsilateral - Native
## Medulla - Luteal phase - Contralateral - Decellularized - Cortex - Follicular phase - Ipsilateral -
## Medulla - Luteal phase - Contralateral - Native - Cortex - Follicular phase - Ipsilateral - Native =
## Medulla - Luteal phase - Ipsilateral - Decellularized - Cortex - Follicular phase - Ipsilateral - Na
## Medulla - Luteal phase - Ipsilateral - Native - Cortex - Follicular phase - Ipsilateral - Native ==
## Cortex - Luteal phase - Contralateral - Native - Cortex - Luteal phase - Contralateral - Decellulari
## Cortex - Luteal phase - Ipsilateral - Decellularized - Cortex - Luteal phase - Contralateral - Decel
## Cortex - Luteal phase - Ipsilateral - Native - Cortex - Luteal phase - Contralateral - Decellularize
## Medulla - Follicular phase - Contralateral - Decellularized - Cortex - Luteal phase - Contralateral
## Medulla - Follicular phase - Contralateral - Native - Cortex - Luteal phase - Contralateral - Decell
## Medulla - Follicular phase - Ipsilateral - Decellularized - Cortex - Luteal phase - Contralateral -
```

```
## Medulla - Follicular phase - Ipsilateral - Native - Cortex - Luteal phase - Contralateral - Decellula
## Medulla - Luteal phase - Contralateral - Decellularized - Cortex - Luteal phase - Contralateral - Dec
## Medulla - Luteal phase - Contralateral - Native - Cortex - Luteal phase - Contralateral - Decellulari
## Medulla - Luteal phase - Ipsilateral - Decellularized - Cortex - Luteal phase - Contralateral - Decel
## Medulla - Luteal phase - Ipsilateral - Native - Cortex - Luteal phase - Contralateral - Decellulariz
## Cortex - Luteal phase - Ipsilateral - Decellularized - Cortex - Luteal phase - Contralateral - Nativ
## Cortex - Luteal phase - Ipsilateral - Native - Cortex - Luteal phase - Contralateral - Native == 0
## Medulla - Follicular phase - Contralateral - Decellularized - Cortex - Luteal phase - Contralateral
## Medulla - Follicular phase - Contralateral - Native - Cortex - Luteal phase - Contralateral - Native
## Medulla - Follicular phase - Ipsilateral - Decellularized - Cortex - Luteal phase - Contralateral -
## Medulla - Follicular phase - Ipsilateral - Native - Cortex - Luteal phase - Contralateral - Native =
## Medulla - Luteal phase - Contralateral - Decellularized - Cortex - Luteal phase - Contralateral - Na
## Medulla - Luteal phase - Contralateral - Native - Cortex - Luteal phase - Contralateral - Native ==
## Medulla - Luteal phase - Ipsilateral - Decellularized - Cortex - Luteal phase - Contralateral - Nati
## Medulla - Luteal phase - Ipsilateral - Native - Cortex - Luteal phase - Contralateral - Native == 0
## Cortex - Luteal phase - Ipsilateral - Native - Cortex - Luteal phase - Ipsilateral - Decellularized
## Medulla - Follicular phase - Contralateral - Decellularized - Cortex - Luteal phase - Ipsilateral -
## Medulla - Follicular phase - Contralateral - Native - Cortex - Luteal phase - Ipsilateral - Decellul
## Medulla - Follicular phase - Ipsilateral - Decellularized - Cortex - Luteal phase - Ipsilateral - De
## Medulla - Follicular phase - Ipsilateral - Native - Cortex - Luteal phase - Ipsilateral - Decellulari
## Medulla - Luteal phase - Contralateral - Decellularized - Cortex - Luteal phase - Ipsilateral - Decel
## Medulla - Luteal phase - Contralateral - Native - Cortex - Luteal phase - Ipsilateral - Decellulariz
## Medulla - Luteal phase - Ipsilateral - Decellularized - Cortex - Luteal phase - Ipsilateral - Decell
## Medulla - Luteal phase - Ipsilateral - Native - Cortex - Luteal phase - Ipsilateral - Decellularized
## Medulla - Follicular phase - Contralateral - Decellularized - Cortex - Luteal phase - Ipsilateral -
## Medulla - Follicular phase - Contralateral - Native - Cortex - Luteal phase - Ipsilateral - Native =
## Medulla - Follicular phase - Ipsilateral - Decellularized - Cortex - Luteal phase - Ipsilateral - Na
## Medulla - Follicular phase - Ipsilateral - Native - Cortex - Luteal phase - Ipsilateral - Native ==
## Medulla - Luteal phase - Contralateral - Decellularized - Cortex - Luteal phase - Ipsilateral - Nati
## Medulla - Luteal phase - Contralateral - Native - Cortex - Luteal phase - Ipsilateral - Native == 0
## Medulla - Luteal phase - Ipsilateral - Decellularized - Cortex - Luteal phase - Ipsilateral - Native
## Medulla - Luteal phase - Ipsilateral - Native - Cortex - Luteal phase - Ipsilateral - Native == 0
## Medulla - Follicular phase - Contralateral - Native - Medulla - Follicular phase - Contralateral - De
## Medulla - Follicular phase - Ipsilateral - Decellularized - Medulla - Follicular phase - Contralatera
## Medulla - Follicular phase - Ipsilateral - Native - Medulla - Follicular phase - Contralateral - Dec
## Medulla - Luteal phase - Contralateral - Decellularized - Medulla - Follicular phase - Contralateral
## Medulla - Luteal phase - Contralateral - Native - Medulla - Follicular phase - Contralateral - Decel
## Medulla - Luteal phase - Ipsilateral - Decellularized - Medulla - Follicular phase - Contralateral -
## Medulla - Luteal phase - Ipsilateral - Native - Medulla - Follicular phase - Contralateral - Decellu
## Medulla - Follicular phase - Ipsilateral - Decellularized - Medulla - Follicular phase - Contralatera
## Medulla - Follicular phase - Ipsilateral - Native - Medulla - Follicular phase - Contralateral - Nati
## Medulla - Luteal phase - Contralateral - Decellularized - Medulla - Follicular phase - Contralateral
## Medulla - Luteal phase - Contralateral - Native - Medulla - Follicular phase - Contralateral - Nativ
## Medulla - Luteal phase - Ipsilateral - Decellularized - Medulla - Follicular phase - Contralateral -
## Medulla - Luteal phase - Ipsilateral - Native - Medulla - Follicular phase - Contralateral - Native
## Medulla - Follicular phase - Ipsilateral - Native - Medulla - Follicular phase - Ipsilateral - Decel
## Medulla - Luteal phase - Contralateral - Decellularized - Medulla - Follicular phase - Ipsilateral -
## Medulla - Luteal phase - Contralateral - Native - Medulla - Follicular phase - Ipsilateral - Decellu
## Medulla - Luteal phase - Ipsilateral - Decellularized - Medulla - Follicular phase - Ipsilateral - De
## Medulla - Luteal phase - Ipsilateral - Native - Medulla - Follicular phase - Ipsilateral - Decellula
## Medulla - Luteal phase - Contralateral - Decellularized - Medulla - Follicular phase - Ipsilateral -
## Medulla - Luteal phase - Contralateral - Native - Medulla - Follicular phase - Ipsilateral - Native =
## Medulla - Luteal phase - Ipsilateral - Decellularized - Medulla - Follicular phase - Ipsilateral - Na
## Medulla - Luteal phase - Ipsilateral - Native - Medulla - Follicular phase - Ipsilateral - Native ==
```

```
## Medulla - Luteal phase - Contralateral - Native - Medulla - Luteal phase - Contralateral - Decellula
## Medulla - Luteal phase - Ipsilateral - Decellularized - Medulla - Luteal phase - Contralateral - Dec
## Medulla - Luteal phase - Ipsilateral - Native - Medulla - Luteal phase - Contralateral - Decellulari
## Medulla - Luteal phase - Ipsilateral - Decellularized - Medulla - Luteal phase - Contralateral - Nat
## Medulla - Luteal phase - Ipsilateral - Native - Medulla - Luteal phase - Contralateral - Native == 0
## Medulla - Luteal phase - Ipsilateral - Native - Medulla - Luteal phase - Ipsilateral - Decellularize
##
## Cortex - Follicular phase - Contralateral - Native - Cortex - Follicular phase - Contralateral - Dec
## Cortex - Follicular phase - Ipsilateral - Decellularized - Cortex - Follicular phase - Contralateral
## Cortex - Follicular phase - Ipsilateral - Native - Cortex - Follicular phase - Contralateral - Decel
## Cortex - Luteal phase - Contralateral - Decellularized - Cortex - Follicular phase - Contralateral -
## Cortex - Luteal phase - Contralateral - Native - Cortex - Follicular phase - Contralateral - Decellu
## Cortex - Luteal phase - Ipsilateral - Decellularized - Cortex - Follicular phase - Contralateral - D
## Cortex - Luteal phase - Ipsilateral - Native - Cortex - Follicular phase - Contralateral - Decellula
## Medulla - Follicular phase - Contralateral - Decellularized - Cortex - Follicular phase - Contralate
## Medulla - Follicular phase - Contralateral - Native - Cortex - Follicular phase - Contralateral - De
## Medulla - Follicular phase - Ipsilateral - Decellularized - Cortex - Follicular phase - Contralatera
## Medulla - Follicular phase - Ipsilateral - Native - Cortex - Follicular phase - Contralateral - Dece
## Medulla - Luteal phase - Contralateral - Decellularized - Cortex - Follicular phase - Contralateral
## Medulla - Luteal phase - Contralateral - Native - Cortex - Follicular phase - Contralateral - Decell
## Medulla - Luteal phase - Ipsilateral - Decellularized - Cortex - Follicular phase - Contralateral -
## Medulla - Luteal phase - Ipsilateral - Native - Cortex - Follicular phase - Contralateral - Decellul
## Cortex - Follicular phase - Ipsilateral - Decellularized - Cortex - Follicular phase - Contralateral
## Cortex - Follicular phase - Ipsilateral - Native - Cortex - Follicular phase - Contralateral - Nativ
## Cortex - Luteal phase - Contralateral - Decellularized - Cortex - Follicular phase - Contralateral -
## Cortex - Luteal phase - Contralateral - Native - Cortex - Follicular phase - Contralateral - Native
## Cortex - Luteal phase - Ipsilateral - Decellularized - Cortex - Follicular phase - Contralateral - N
## Cortex - Luteal phase - Ipsilateral - Native - Cortex - Follicular phase - Contralateral - Native ==
## Medulla - Follicular phase - Contralateral - Decellularized - Cortex - Follicular phase - Contralate
## Medulla - Follicular phase - Contralateral - Native - Cortex - Follicular phase - Contralateral - Na
## Medulla - Follicular phase - Ipsilateral - Decellularized - Cortex - Follicular phase - Contralatera
## Medulla - Follicular phase - Ipsilateral - Native - Cortex - Follicular phase - Contralateral - Nati
## Medulla - Luteal phase - Contralateral - Decellularized - Cortex - Follicular phase - Contralateral
## Medulla - Luteal phase - Contralateral - Native - Cortex - Follicular phase - Contralateral - Native
## Medulla - Luteal phase - Ipsilateral - Decellularized - Cortex - Follicular phase - Contralateral -
## Medulla - Luteal phase - Ipsilateral - Native - Cortex - Follicular phase - Contralateral - Native =
## Cortex - Follicular phase - Ipsilateral - Native - Cortex - Follicular phase - Ipsilateral - Decellu
## Cortex - Luteal phase - Contralateral - Decellularized - Cortex - Follicular phase - Ipsilateral - D
## Cortex - Luteal phase - Contralateral - Native - Cortex - Follicular phase - Ipsilateral - Decellula
## Cortex - Luteal phase - Ipsilateral - Decellularized - Cortex - Follicular phase - Ipsilateral - Dec
## Cortex - Luteal phase - Ipsilateral - Native - Cortex - Follicular phase - Ipsilateral - Decellulari
## Medulla - Follicular phase - Contralateral - Decellularized - Cortex - Follicular phase - Ipsilatera
## Medulla - Follicular phase - Contralateral - Native - Cortex - Follicular phase - Ipsilateral - Dece
## Medulla - Follicular phase - Ipsilateral - Decellularized - Cortex - Follicular phase - Ipsilateral
## Medulla - Follicular phase - Ipsilateral - Native - Cortex - Follicular phase - Ipsilateral - Decell
## Medulla - Luteal phase - Contralateral - Decellularized - Cortex - Follicular phase - Ipsilateral -
## Medulla - Luteal phase - Contralateral - Native - Cortex - Follicular phase - Ipsilateral - Decellul
## Medulla - Luteal phase - Ipsilateral - Decellularized - Cortex - Follicular phase - Ipsilateral - De
## Medulla - Luteal phase - Ipsilateral - Native - Cortex - Follicular phase - Ipsilateral - Decellular
## Cortex - Luteal phase - Contralateral - Decellularized - Cortex - Follicular phase - Ipsilateral - N
## Cortex - Luteal phase - Contralateral - Native - Cortex - Follicular phase - Ipsilateral - Native ==
## Cortex - Luteal phase - Ipsilateral - Decellularized - Cortex - Follicular phase - Ipsilateral - Nat
## Cortex - Luteal phase - Ipsilateral - Native - Cortex - Follicular phase - Ipsilateral - Native == 0
## Medulla - Follicular phase - Contralateral - Decellularized - Cortex - Follicular phase - Ipsilatera
```

```
## Medulla - Follicular phase - Contralateral - Native - Cortex - Follicular phase - Ipsilateral - Nati
## Medulla - Follicular phase - Ipsilateral - Decellularized - Cortex - Follicular phase - Ipsilateral
## Medulla - Follicular phase - Ipsilateral - Native - Cortex - Follicular phase - Ipsilateral - Native
## Medulla - Luteal phase - Contralateral - Decellularized - Cortex - Follicular phase - Ipsilateral -
## Medulla - Luteal phase - Contralateral - Native - Cortex - Follicular phase - Ipsilateral - Native =
## Medulla - Luteal phase - Ipsilateral - Decellularized - Cortex - Follicular phase - Ipsilateral - Na
## Medulla - Luteal phase - Ipsilateral - Native - Cortex - Follicular phase - Ipsilateral - Native ==
## Cortex - Luteal phase - Contralateral - Native - Cortex - Luteal phase - Contralateral - Decellulari
## Cortex - Luteal phase - Ipsilateral - Decellularized - Cortex - Luteal phase - Contralateral - Decel
## Cortex - Luteal phase - Ipsilateral - Native - Cortex - Luteal phase - Contralateral - Decellularize
## Medulla - Follicular phase - Contralateral - Decellularized - Cortex - Luteal phase - Contralateral
## Medulla - Follicular phase - Contralateral - Native - Cortex - Luteal phase - Contralateral - Decell
## Medulla - Follicular phase - Ipsilateral - Decellularized - Cortex - Luteal phase - Contralateral -
## Medulla - Follicular phase - Ipsilateral - Native - Cortex - Luteal phase - Contralateral - Decellul
## Medulla - Luteal phase - Contralateral - Decellularized - Cortex - Luteal phase - Contralateral - De
## Medulla - Luteal phase - Contralateral - Native - Cortex - Luteal phase - Contralateral - Decellular
## Medulla - Luteal phase - Ipsilateral - Decellularized - Cortex - Luteal phase - Contralateral - Dece
## Medulla - Luteal phase - Ipsilateral - Native - Cortex - Luteal phase - Contralateral - Decellulariz
## Cortex - Luteal phase - Ipsilateral - Decellularized - Cortex - Luteal phase - Contralateral - Nativ
## Cortex - Luteal phase - Ipsilateral - Native - Cortex - Luteal phase - Contralateral - Native == 0
## Medulla - Follicular phase - Contralateral - Decellularized - Cortex - Luteal phase - Contralateral
## Medulla - Follicular phase - Contralateral - Native - Cortex - Luteal phase - Contralateral - Native
## Medulla - Follicular phase - Ipsilateral - Decellularized - Cortex - Luteal phase - Contralateral -
## Medulla - Follicular phase - Ipsilateral - Native - Cortex - Luteal phase - Contralateral - Native =
## Medulla - Luteal phase - Contralateral - Decellularized - Cortex - Luteal phase - Contralateral - Na
## Medulla - Luteal phase - Contralateral - Native - Cortex - Luteal phase - Contralateral - Native ==
## Medulla - Luteal phase - Ipsilateral - Decellularized - Cortex - Luteal phase - Contralateral - Nati
## Medulla - Luteal phase - Ipsilateral - Native - Cortex - Luteal phase - Contralateral - Native == 0
## Cortex - Luteal phase - Ipsilateral - Native - Cortex - Luteal phase - Ipsilateral - Decellularized
## Medulla - Follicular phase - Contralateral - Decellularized - Cortex - Luteal phase - Ipsilateral -
## Medulla - Follicular phase - Contralateral - Native - Cortex - Luteal phase - Ipsilateral - Decellul
## Medulla - Follicular phase - Ipsilateral - Decellularized - Cortex - Luteal phase - Ipsilateral - De
## Medulla - Follicular phase - Ipsilateral - Native - Cortex - Luteal phase - Ipsilateral - Decellular
## Medulla - Luteal phase - Contralateral - Decellularized - Cortex - Luteal phase - Ipsilateral - Dece
## Medulla - Luteal phase - Contralateral - Native - Cortex - Luteal phase - Ipsilateral - Decellulariz
## Medulla - Luteal phase - Ipsilateral - Decellularized - Cortex - Luteal phase - Ipsilateral - Decell
## Medulla - Luteal phase - Ipsilateral - Native - Cortex - Luteal phase - Ipsilateral - Decellularized
## Medulla - Follicular phase - Contralateral - Decellularized - Cortex - Luteal phase - Ipsilateral -
## Medulla - Follicular phase - Contralateral - Native - Cortex - Luteal phase - Ipsilateral - Native =
## Medulla - Follicular phase - Ipsilateral - Decellularized - Cortex - Luteal phase - Ipsilateral - Na
## Medulla - Follicular phase - Ipsilateral - Native - Cortex - Luteal phase - Ipsilateral - Native ==
## Medulla - Luteal phase - Contralateral - Decellularized - Cortex - Luteal phase - Ipsilateral - Nati
## Medulla - Luteal phase - Contralateral - Native - Cortex - Luteal phase - Ipsilateral - Native == 0
## Medulla - Luteal phase - Ipsilateral - Decellularized - Cortex - Luteal phase - Ipsilateral - Native
## Medulla - Luteal phase - Ipsilateral - Native - Cortex - Luteal phase - Ipsilateral - Native == 0
## Medulla - Follicular phase - Contralateral - Native - Medulla - Follicular phase - Contralateral - D
## Medulla - Follicular phase - Ipsilateral - Decellularized - Medulla - Follicular phase - Contralater
## Medulla - Follicular phase - Ipsilateral - Native - Medulla - Follicular phase - Contralateral - Dec
## Medulla - Luteal phase - Contralateral - Decellularized - Medulla - Follicular phase - Contralateral
## Medulla - Luteal phase - Contralateral - Native - Medulla - Follicular phase - Contralateral - Decel
## Medulla - Luteal phase - Ipsilateral - Decellularized - Medulla - Follicular phase - Contralateral -
## Medulla - Luteal phase - Ipsilateral - Native - Medulla - Follicular phase - Contralateral - Decellu
## Medulla - Follicular phase - Ipsilateral - Decellularized - Medulla - Follicular phase - Contralater
## Medulla - Follicular phase - Ipsilateral - Native - Medulla - Follicular phase - Contralateral - Nat
```

```
## Medulla - Luteal phase - Contralateral - Decellularized - Medulla - Follicular phase - Contralateral
## Medulla - Luteal phase - Contralateral - Native - Medulla - Follicular phase - Contralateral - Nativ
## Medulla - Luteal phase - Ipsilateral - Decellularized - Medulla - Follicular phase - Contralateral -
## Medulla - Luteal phase - Ipsilateral - Native - Medulla - Follicular phase - Contralateral - Native
## Medulla - Follicular phase - Ipsilateral - Native - Medulla - Follicular phase - Ipsilateral - Decel
## Medulla - Luteal phase - Contralateral - Decellularized - Medulla - Follicular phase - Ipsilateral -
## Medulla - Luteal phase - Contralateral - Native - Medulla - Follicular phase - Ipsilateral - Decellu
## Medulla - Luteal phase - Ipsilateral - Decellularized - Medulla - Follicular phase - Ipsilateral - De
## Medulla - Luteal phase - Ipsilateral - Native - Medulla - Follicular phase - Ipsilateral - Decellula
## Medulla - Luteal phase - Contralateral - Decellularized - Medulla - Follicular phase - Ipsilateral -
## Medulla - Luteal phase - Contralateral - Native - Medulla - Follicular phase - Ipsilateral - Native
## Medulla - Luteal phase - Ipsilateral - Decellularized - Medulla - Follicular phase - Ipsilateral - Na
## Medulla - Luteal phase - Ipsilateral - Native - Medulla - Follicular phase - Ipsilateral - Native ==
## Medulla - Luteal phase - Contralateral - Native - Medulla - Luteal phase - Contralateral - Decellula
## Medulla - Luteal phase - Ipsilateral - Decellularized - Medulla - Luteal phase - Contralateral - Dec
## Medulla - Luteal phase - Ipsilateral - Native - Medulla - Luteal phase - Contralateral - Decellulari
## Medulla - Luteal phase - Ipsilateral - Decellularized - Medulla - Luteal phase - Contralateral - Nat
## Medulla - Luteal phase - Ipsilateral - Native - Medulla - Luteal phase - Contralateral - Native == 0
## Medulla - Luteal phase - Ipsilateral - Native - Medulla - Luteal phase - Ipsilateral - Decellularize
##
## Cortex - Follicular phase - Contralateral - Native - Cortex - Follicular phase - Contralateral - Dec
## Cortex - Follicular phase - Ipsilateral - Decellularized - Cortex - Follicular phase - Contralateral
## Cortex - Follicular phase - Ipsilateral - Native - Cortex - Follicular phase - Contralateral - Decel
## Cortex - Luteal phase - Contralateral - Decellularized - Cortex - Follicular phase - Contralateral -
## Cortex - Luteal phase - Contralateral - Native - Cortex - Follicular phase - Contralateral - Decellu
## Cortex - Luteal phase - Ipsilateral - Decellularized - Cortex - Follicular phase - Contralateral - De
## Cortex - Luteal phase - Ipsilateral - Native - Cortex - Follicular phase - Contralateral - Decellula
## Medulla - Follicular phase - Contralateral - Decellularized - Cortex - Follicular phase - Contralate
## Medulla - Follicular phase - Contralateral - Native - Cortex - Follicular phase - Contralateral - De
## Medulla - Follicular phase - Ipsilateral - Decellularized - Cortex - Follicular phase - Contralatera
## Medulla - Follicular phase - Ipsilateral - Native - Cortex - Follicular phase - Contralateral - Dece
## Medulla - Luteal phase - Contralateral - Decellularized - Cortex - Follicular phase - Contralateral
## Medulla - Luteal phase - Contralateral - Native - Cortex - Follicular phase - Contralateral - Decell
## Medulla - Luteal phase - Ipsilateral - Decellularized - Cortex - Follicular phase - Contralateral -
## Medulla - Luteal phase - Ipsilateral - Native - Cortex - Follicular phase - Contralateral - Decellula
## Cortex - Follicular phase - Ipsilateral - Decellularized - Cortex - Follicular phase - Contralateral
## Cortex - Follicular phase - Ipsilateral - Native - Cortex - Follicular phase - Contralateral - Nativ
## Cortex - Luteal phase - Contralateral - Decellularized - Cortex - Follicular phase - Contralateral -
## Cortex - Luteal phase - Contralateral - Native - Cortex - Follicular phase - Contralateral - Native
## Cortex - Luteal phase - Ipsilateral - Decellularized - Cortex - Follicular phase - Contralateral - Na
## Cortex - Luteal phase - Ipsilateral - Native - Cortex - Follicular phase - Contralateral - Native ==
## Medulla - Follicular phase - Contralateral - Decellularized - Cortex - Follicular phase - Contralate
## Medulla - Follicular phase - Contralateral - Native - Cortex - Follicular phase - Contralateral - Na
## Medulla - Follicular phase - Ipsilateral - Decellularized - Cortex - Follicular phase - Contralatera
## Medulla - Follicular phase - Ipsilateral - Native - Cortex - Follicular phase - Contralateral - Nati
## Medulla - Luteal phase - Contralateral - Decellularized - Cortex - Follicular phase - Contralateral
## Medulla - Luteal phase - Contralateral - Native - Cortex - Follicular phase - Contralateral - Native
## Medulla - Luteal phase - Ipsilateral - Decellularized - Cortex - Follicular phase - Contralateral -
## Medulla - Luteal phase - Ipsilateral - Native - Cortex - Follicular phase - Contralateral - Native =
## Cortex - Follicular phase - Ipsilateral - Native - Cortex - Follicular phase - Ipsilateral - Decellu
## Cortex - Luteal phase - Contralateral - Decellularized - Cortex - Follicular phase - Ipsilateral - De
## Cortex - Luteal phase - Contralateral - Native - Cortex - Follicular phase - Ipsilateral - Decellula
## Cortex - Luteal phase - Ipsilateral - Decellularized - Cortex - Follicular phase - Ipsilateral - Dec
## Cortex - Luteal phase - Ipsilateral - Native - Cortex - Follicular phase - Ipsilateral - Decellulari
```

```
## Medulla - Follicular phase - Contralateral - Decellularized - Cortex - Follicular phase - Ipsilateral
## Medulla - Follicular phase - Contralateral - Native - Cortex - Follicular phase - Ipsilateral - Decel
## Medulla - Follicular phase - Ipsilateral - Decellularized - Cortex - Follicular phase - Ipsilateral
## Medulla - Follicular phase - Ipsilateral - Native - Cortex - Follicular phase - Ipsilateral - Decellu
## Medulla - Luteal phase - Contralateral - Decellularized - Cortex - Follicular phase - Ipsilateral -
## Medulla - Luteal phase - Contralateral - Native - Cortex - Follicular phase - Ipsilateral - Decellula
## Medulla - Luteal phase - Ipsilateral - Decellularized - Cortex - Follicular phase - Ipsilateral - Dec
## Medulla - Luteal phase - Ipsilateral - Native - Cortex - Follicular phase - Ipsilateral - Decellular
## Cortex - Luteal phase - Contralateral - Decellularized - Cortex - Follicular phase - Ipsilateral - Na
## Cortex - Luteal phase - Contralateral - Native - Cortex - Follicular phase - Ipsilateral - Native ==
## Cortex - Luteal phase - Ipsilateral - Decellularized - Cortex - Follicular phase - Ipsilateral - Nati
## Cortex - Luteal phase - Ipsilateral - Native - Cortex - Follicular phase - Ipsilateral - Native == 0
## Medulla - Follicular phase - Contralateral - Decellularized - Cortex - Follicular phase - Ipsilateral
## Medulla - Follicular phase - Contralateral - Native - Cortex - Follicular phase - Ipsilateral - Nativ
## Medulla - Follicular phase - Ipsilateral - Decellularized - Cortex - Follicular phase - Ipsilateral
## Medulla - Follicular phase - Ipsilateral - Native - Cortex - Follicular phase - Ipsilateral - Native
## Medulla - Luteal phase - Contralateral - Decellularized - Cortex - Follicular phase - Ipsilateral -
## Medulla - Luteal phase - Contralateral - Native - Cortex - Follicular phase - Ipsilateral - Native =
## Medulla - Luteal phase - Ipsilateral - Decellularized - Cortex - Follicular phase - Ipsilateral - Na
## Medulla - Luteal phase - Ipsilateral - Native - Cortex - Follicular phase - Ipsilateral - Native ==
## Cortex - Luteal phase - Contralateral - Native - Cortex - Luteal phase - Contralateral - Decellulari
## Cortex - Luteal phase - Ipsilateral - Decellularized - Cortex - Luteal phase - Contralateral - Decel
## Cortex - Luteal phase - Ipsilateral - Native - Cortex - Luteal phase - Contralateral - Decellularize
## Medulla - Follicular phase - Contralateral - Decellularized - Cortex - Luteal phase - Contralateral
## Medulla - Follicular phase - Contralateral - Native - Cortex - Luteal phase - Contralateral - Decellu
## Medulla - Follicular phase - Ipsilateral - Decellularized - Cortex - Luteal phase - Contralateral -
## Medulla - Follicular phase - Ipsilateral - Native - Cortex - Luteal phase - Contralateral - Decellula
## Medulla - Luteal phase - Contralateral - Decellularized - Cortex - Luteal phase - Contralateral - De
## Medulla - Luteal phase - Contralateral - Native - Cortex - Luteal phase - Contralateral - Decellulari
## Medulla - Luteal phase - Ipsilateral - Decellularized - Cortex - Luteal phase - Contralateral - Decel
## Medulla - Luteal phase - Ipsilateral - Native - Cortex - Luteal phase - Contralateral - Decellularize
## Cortex - Luteal phase - Ipsilateral - Decellularized - Cortex - Luteal phase - Contralateral - Native
## Cortex - Luteal phase - Ipsilateral - Native - Cortex - Luteal phase - Contralateral - Native == 0
## Medulla - Follicular phase - Contralateral - Decellularized - Cortex - Luteal phase - Contralateral
## Medulla - Follicular phase - Contralateral - Native - Cortex - Luteal phase - Contralateral - Native
## Medulla - Follicular phase - Ipsilateral - Decellularized - Cortex - Luteal phase - Contralateral -
## Medulla - Follicular phase - Ipsilateral - Native - Cortex - Luteal phase - Contralateral - Native =
## Medulla - Luteal phase - Contralateral - Decellularized - Cortex - Luteal phase - Contralateral - Na
## Medulla - Luteal phase - Contralateral - Native - Cortex - Luteal phase - Contralateral - Native ==
## Medulla - Luteal phase - Ipsilateral - Decellularized - Cortex - Luteal phase - Contralateral - Nati
## Medulla - Luteal phase - Ipsilateral - Native - Cortex - Luteal phase - Contralateral - Native == 0
## Cortex - Luteal phase - Ipsilateral - Native - Cortex - Luteal phase - Ipsilateral - Decellularized
## Medulla - Follicular phase - Contralateral - Decellularized - Cortex - Luteal phase - Ipsilateral -
## Medulla - Follicular phase - Contralateral - Native - Cortex - Luteal phase - Ipsilateral - Decellula
## Medulla - Follicular phase - Ipsilateral - Decellularized - Cortex - Luteal phase - Ipsilateral - De
## Medulla - Follicular phase - Ipsilateral - Native - Cortex - Luteal phase - Ipsilateral - Decellular
## Medulla - Luteal phase - Contralateral - Decellularized - Cortex - Luteal phase - Ipsilateral - Decel
## Medulla - Luteal phase - Contralateral - Native - Cortex - Luteal phase - Ipsilateral - Decellularize
## Medulla - Luteal phase - Ipsilateral - Decellularized - Cortex - Luteal phase - Ipsilateral - Decellu
## Medulla - Luteal phase - Ipsilateral - Native - Cortex - Luteal phase - Ipsilateral - Decellularized
## Medulla - Follicular phase - Contralateral - Decellularized - Cortex - Luteal phase - Ipsilateral -
## Medulla - Follicular phase - Contralateral - Native - Cortex - Luteal phase - Ipsilateral - Native =
## Medulla - Follicular phase - Ipsilateral - Decellularized - Cortex - Luteal phase - Ipsilateral - Na
## Medulla - Follicular phase - Ipsilateral - Native - Cortex - Luteal phase - Ipsilateral - Native ==
```

```
## Medulla - Luteal phase - Contralateral - Decellularized - Cortex - Luteal phase - Ipsilateral - Nati
## Medulla - Luteal phase - Contralateral - Native - Cortex - Luteal phase - Ipsilateral - Native == 0
## Medulla - Luteal phase - Ipsilateral - Decellularized - Cortex - Luteal phase - Ipsilateral - Native
## Medulla - Luteal phase - Ipsilateral - Native - Cortex - Luteal phase - Ipsilateral - Native == 0
## Medulla - Follicular phase - Contralateral - Native - Medulla - Follicular phase - Contralateral - De
## Medulla - Follicular phase - Ipsilateral - Decellularized - Medulla - Follicular phase - Contralatera
## Medulla - Follicular phase - Ipsilateral - Native - Medulla - Follicular phase - Contralateral - Dec
## Medulla - Luteal phase - Contralateral - Decellularized - Medulla - Follicular phase - Contralateral
## Medulla - Luteal phase - Contralateral - Native - Medulla - Follicular phase - Contralateral - Decel
## Medulla - Luteal phase - Ipsilateral - Decellularized - Medulla - Follicular phase - Contralateral -
## Medulla - Luteal phase - Ipsilateral - Native - Medulla - Follicular phase - Contralateral - Decellu
## Medulla - Follicular phase - Ipsilateral - Decellularized - Medulla - Follicular phase - Contralatera
## Medulla - Follicular phase - Ipsilateral - Native - Medulla - Follicular phase - Contralateral - Nati
## Medulla - Luteal phase - Contralateral - Decellularized - Medulla - Follicular phase - Contralateral
## Medulla - Luteal phase - Contralateral - Native - Medulla - Follicular phase - Contralateral - Native
## Medulla - Luteal phase - Ipsilateral - Decellularized - Medulla - Follicular phase - Contralateral -
## Medulla - Luteal phase - Ipsilateral - Native - Medulla - Follicular phase - Contralateral - Native =
## Medulla - Follicular phase - Ipsilateral - Native - Medulla - Follicular phase - Ipsilateral - Decel
## Medulla - Luteal phase - Contralateral - Decellularized - Medulla - Follicular phase - Ipsilateral -
## Medulla - Luteal phase - Contralateral - Native - Medulla - Follicular phase - Ipsilateral - Decellu
## Medulla - Luteal phase - Ipsilateral - Decellularized - Medulla - Follicular phase - Ipsilateral - De
## Medulla - Luteal phase - Ipsilateral - Native - Medulla - Follicular phase - Ipsilateral - Decellula
## Medulla - Luteal phase - Contralateral - Decellularized - Medulla - Follicular phase - Ipsilateral -
## Medulla - Luteal phase - Contralateral - Native - Medulla - Follicular phase - Ipsilateral - Native =
## Medulla - Luteal phase - Ipsilateral - Decellularized - Medulla - Follicular phase - Ipsilateral - Na
## Medulla - Luteal phase - Ipsilateral - Native - Medulla - Follicular phase - Ipsilateral - Native ==
## Medulla - Luteal phase - Contralateral - Native - Medulla - Luteal phase - Contralateral - Decellula
## Medulla - Luteal phase - Ipsilateral - Decellularized - Medulla - Luteal phase - Contralateral - Dec
## Medulla - Luteal phase - Ipsilateral - Native - Medulla - Luteal phase - Contralateral - Decellulari
## Medulla - Luteal phase - Ipsilateral - Decellularized - Medulla - Luteal phase - Contralateral - Nat
## Medulla - Luteal phase - Ipsilateral - Native - Medulla - Luteal phase - Contralateral - Native == 0
## Medulla - Luteal phase - Ipsilateral - Native - Medulla - Luteal phase - Ipsilateral - Decellularize
## ---
## Signif. codes:  0 '***' 0.001 '**' 0.01 '*' 0.05 '.' 0.1 ' ' 1
## (Adjusted p values reported -- single-step method)
```

```
################### Porosity ########################################

# Read the dataset from a CSV file
df <- read.csv("Ovary_porosity.csv")

# Summary data

summary_df <- df %>%
  group_by(Cellular, Tissue, Position, Estrus_cycle) %>%
  summarise(mean_Porosity = mean(Porosity),
            sd_Porosity = sd(Porosity))
```

```
## `summarise()` has grouped output by 'Cellular', 'Tissue', 'Position'. You can
## override using the `.groups` argument.
```

```
# Plot

png(filename="Ovary_porosity.png",
    width = 10, height = 8, units = "in",
```

```r
    res = 600)

par(mgp=c(1.5,0.5,0),
    mar = c(4, #bottom
            4, #left
            3, #top
            2),
    cex.main = 0.8,
    family = "sans" #right
)

# Update the labels for Position and Estrus_cycle
df$Position <- factor(df$Position, labels = c("Contralateral", "Ipsilateral"))
df$Estrus_cycle <- factor(df$Estrus_cycle, labels = c("Follicular phase", "Luteal phase"))

# Convert Animal_ID to a factor
df$Animal_ID <- as.factor(df$Animal_ID)

# Create a combined factor variable for Estrus_cycle and Cellular
df$Combined <- factor(paste(df$Estrus_cycle, df$Cellular, sep = " "),
                      levels = c("Follicular phase Native", "Follicular phase Decellularized",
                                 "Luteal phase Native", "Luteal phase Decellularized"))


# Define fill colors for each factor level
fill_colors <- c("Follicular phase Native" = viridis_pal(option = "D")(1),
                 "Follicular phase Decellularized" = alpha(viridis_pal(option = "D")(1), 0.2),
                 "Luteal phase Native" = "#238A8DFF",
                 "Luteal phase Decellularized" = alpha("#238A8DFF", 0.2))

# Create the plot using ggplot
ggplot(data = df, aes(x = Tissue, y = Porosity, fill = Combined)) +
  geom_boxplot(width = 0.7, outlier.shape = NA) +
  geom_jitter(position = position_jitterdodge(), alpha = 0.8) +
  xlab("") +
  ylab("Estimated Porosity (%)") +
  ggtitle("Boxplot and Individual Data Points of Porosity by Estrus_cycle, Cellular, Position, and X_pos
  scale_fill_manual(values = fill_colors, guide = guide_legend(title = "Combined")) +
  theme_bw() +
  ylim(0, 130) +
  facet_grid(Estrus_cycle ~ Position, space = "free_x", scales = "free_x") +
  scale_x_discrete(labels = c("Cortex", "Medulla"), expand = c(0, 0)) +
  theme(panel.grid.major = element_blank(), panel.grid.minor = element_blank(),
        axis.text.x = element_text(size = 11, angle = 0, hjust = 0.5),
        axis.text.y = element_text(size = 12, angle = 0, hjust = 0.5),
        axis.title.x = element_text(size = 18, face = "bold"),
        axis.title.y = element_text(size = 16, face = "bold"),
        legend.position = "right",
        strip.text = element_text(size = 14, face = "bold"))  # Increase the size of facet labels

dev.off()
```

```
## pdf
##   2
```

```r
# Stats

# Read the dataset from a CSV file
df <- read.csv("Ovary_porosity.csv")

# Update the labels for Position, Estrus_cycle, and Tissue
df$Position <- factor(df$Position, labels = c("Contralateral", "Ipsilateral"))
df$Estrus_cycle <- factor(df$Estrus_cycle, labels = c("Follicular phase", "Luteal phase"))
df$Cellular <- factor(df$Cellular, labels = c("Native", "Decellularized"))

# Update the labels for Position, Estrus_cycle, and Cellular
df$Position <- factor(df$Position, labels = c("Contralateral", "Ipsilateral"))
df$Estrus_cycle <- factor(df$Estrus_cycle, labels = c("Follicular phase", "Luteal phase"))
df$Cellular <- factor(df$Cellular, labels = c("Native", "Decellularized"))

# Create a factor variable for the comparisons
df$Comparison <- factor(paste(df$Tissue, df$Estrus_cycle, df$Position, df$Cellular, sep = " - "))

# Perform pairwise comparisons using Tukey's method
model <- lm(Porosity ~ Comparison, data = df)
comp <- glht(model, linfct = mcp(Comparison = "Tukey"))

# Summarize the results
summary(comp)
```

```
## Warning in RET$pfunction("adjusted", ...): Completion with error > abseps

## Warning in RET$pfunction("adjusted", ...): Completion with error > abseps

## Warning in RET$pfunction("adjusted", ...): Completion with error > abseps

## Warning in RET$pfunction("adjusted", ...): Completion with error > abseps

## Warning in RET$pfunction("adjusted", ...): Completion with error > abseps

## Warning in RET$pfunction("adjusted", ...): Completion with error > abseps

## Warning in RET$pfunction("adjusted", ...): Completion with error > abseps

## Warning in RET$pfunction("adjusted", ...): Completion with error > abseps

## Warning in RET$pfunction("adjusted", ...): Completion with error > abseps

## Warning in RET$pfunction("adjusted", ...): Completion with error > abseps

## Warning in RET$pfunction("adjusted", ...): Completion with error > abseps

## Warning in RET$pfunction("adjusted", ...): Completion with error > abseps

## Warning in RET$pfunction("adjusted", ...): Completion with error > abseps

## Warning in RET$pfunction("adjusted", ...): Completion with error > abseps

## Warning in RET$pfunction("adjusted", ...): Completion with error > abseps
```

```
## Warning in RET$pfunction("adjusted", ...): Completion with error > abseps

## Warning in RET$pfunction("adjusted", ...): Completion with error > abseps

## Warning in RET$pfunction("adjusted", ...): Completion with error > abseps

## Warning in RET$pfunction("adjusted", ...): Completion with error > abseps

## Warning in RET$pfunction("adjusted", ...): Completion with error > abseps

## Warning in RET$pfunction("adjusted", ...): Completion with error > abseps

## Warning in RET$pfunction("adjusted", ...): Completion with error > abseps

## Warning in RET$pfunction("adjusted", ...): Completion with error > abseps

## Warning in RET$pfunction("adjusted", ...): Completion with error > abseps

## Warning in RET$pfunction("adjusted", ...): Completion with error > abseps

## Warning in RET$pfunction("adjusted", ...): Completion with error > abseps

## Warning in RET$pfunction("adjusted", ...): Completion with error > abseps

## Warning in RET$pfunction("adjusted", ...): Completion with error > abseps

## Warning in RET$pfunction("adjusted", ...): Completion with error > abseps

## Warning in RET$pfunction("adjusted", ...): Completion with error > abseps

## Warning in RET$pfunction("adjusted", ...): Completion with error > abseps

## Warning in RET$pfunction("adjusted", ...): Completion with error > abseps

## Warning in RET$pfunction("adjusted", ...): Completion with error > abseps

## Warning in RET$pfunction("adjusted", ...): Completion with error > abseps

## Warning in RET$pfunction("adjusted", ...): Completion with error > abseps

## Warning in RET$pfunction("adjusted", ...): Completion with error > abseps

## Warning in RET$pfunction("adjusted", ...): Completion with error > abseps

## Warning in RET$pfunction("adjusted", ...): Completion with error > abseps

## Warning in RET$pfunction("adjusted", ...): Completion with error > abseps

## Warning in RET$pfunction("adjusted", ...): Completion with error > abseps
##
##   Simultaneous Tests for General Linear Hypotheses
##
```

```
## Multiple Comparisons of Means: Tukey Contrasts
##
##
## Fit: lm(formula = Porosity ~ Comparison, data = df)
##
## Linear Hypotheses:
##
## Cortex - Follicular phase - Contralateral - Native - Cortex - Follicular phase - Contralateral - Dec
## Cortex - Follicular phase - Ipsilateral - Decellularized - Cortex - Follicular phase - Contralateral
## Cortex - Follicular phase - Ipsilateral - Native - Cortex - Follicular phase - Contralateral - Decel
## Cortex - Luteal phase - Contralateral - Decellularized - Cortex - Follicular phase - Contralateral -
## Cortex - Luteal phase - Contralateral - Native - Cortex - Follicular phase - Contralateral - Decellu
## Cortex - Luteal phase - Ipsilateral - Decellularized - Cortex - Follicular phase - Contralateral - D
## Cortex - Luteal phase - Ipsilateral - Native - Cortex - Follicular phase - Contralateral - Decellula
## Medulla - Follicular phase - Contralateral - Decellularized - Cortex - Follicular phase - Contralate
## Medulla - Follicular phase - Contralateral - Native - Cortex - Follicular phase - Contralateral - De
## Medulla - Follicular phase - Ipsilateral - Decellularized - Cortex - Follicular phase - Contralateral
## Medulla - Follicular phase - Ipsilateral - Native - Cortex - Follicular phase - Contralateral - Decel
## Medulla - Luteal phase - Contralateral - Decellularized - Cortex - Follicular phase - Contralateral -
## Medulla - Luteal phase - Contralateral - Native - Cortex - Follicular phase - Contralateral - Decellu
## Medulla - Luteal phase - Ipsilateral - Decellularized - Cortex - Follicular phase - Contralateral - D
## Medulla - Luteal phase - Ipsilateral - Native - Cortex - Follicular phase - Contralateral - Decellula
## Cortex - Follicular phase - Ipsilateral - Decellularized - Cortex - Follicular phase - Contralateral
## Cortex - Follicular phase - Ipsilateral - Native - Cortex - Follicular phase - Contralateral - Nativ
## Cortex - Luteal phase - Contralateral - Decellularized - Cortex - Follicular phase - Contralateral -
## Cortex - Luteal phase - Contralateral - Native - Cortex - Follicular phase - Contralateral - Native =
## Cortex - Luteal phase - Ipsilateral - Decellularized - Cortex - Follicular phase - Contralateral - Na
## Cortex - Luteal phase - Ipsilateral - Native - Cortex - Follicular phase - Contralateral - Native ==
## Medulla - Follicular phase - Contralateral - Decellularized - Cortex - Follicular phase - Contralate
## Medulla - Follicular phase - Contralateral - Native - Cortex - Follicular phase - Contralateral - Na
## Medulla - Follicular phase - Ipsilateral - Decellularized - Cortex - Follicular phase - Contralateral
## Medulla - Follicular phase - Ipsilateral - Native - Cortex - Follicular phase - Contralateral - Nati
## Medulla - Luteal phase - Contralateral - Decellularized - Cortex - Follicular phase - Contralateral -
## Medulla - Luteal phase - Contralateral - Native - Cortex - Follicular phase - Contralateral - Native
## Medulla - Luteal phase - Ipsilateral - Decellularized - Cortex - Follicular phase - Contralateral - N
## Medulla - Luteal phase - Ipsilateral - Native - Cortex - Follicular phase - Contralateral - Native =
## Cortex - Follicular phase - Ipsilateral - Native - Cortex - Follicular phase - Ipsilateral - Decellu
## Cortex - Luteal phase - Contralateral - Decellularized - Cortex - Follicular phase - Ipsilateral - De
## Cortex - Luteal phase - Contralateral - Native - Cortex - Follicular phase - Ipsilateral - Decellula
## Cortex - Luteal phase - Ipsilateral - Decellularized - Cortex - Follicular phase - Ipsilateral - Dec
## Cortex - Luteal phase - Ipsilateral - Native - Cortex - Follicular phase - Ipsilateral - Decellulari
## Medulla - Follicular phase - Contralateral - Decellularized - Cortex - Follicular phase - Ipsilateral
## Medulla - Follicular phase - Contralateral - Native - Cortex - Follicular phase - Ipsilateral - Decel
## Medulla - Follicular phase - Ipsilateral - Decellularized - Cortex - Follicular phase - Ipsilateral -
## Medulla - Follicular phase - Ipsilateral - Native - Cortex - Follicular phase - Ipsilateral - Decellu
## Medulla - Luteal phase - Contralateral - Decellularized - Cortex - Follicular phase - Ipsilateral - D
## Medulla - Luteal phase - Contralateral - Native - Cortex - Follicular phase - Ipsilateral - Decellula
## Medulla - Luteal phase - Ipsilateral - Decellularized - Cortex - Follicular phase - Ipsilateral - Dec
## Medulla - Luteal phase - Ipsilateral - Native - Cortex - Follicular phase - Ipsilateral - Decellular
## Cortex - Luteal phase - Contralateral - Decellularized - Cortex - Follicular phase - Ipsilateral - Na
## Cortex - Luteal phase - Contralateral - Native - Cortex - Follicular phase - Ipsilateral - Native ==
## Cortex - Luteal phase - Ipsilateral - Decellularized - Cortex - Follicular phase - Ipsilateral - Nati
## Cortex - Luteal phase - Ipsilateral - Native - Cortex - Follicular phase - Ipsilateral - Native == 0
## Medulla - Follicular phase - Contralateral - Decellularized - Cortex - Follicular phase - Ipsilateral
```

```
## Medulla - Follicular phase - Contralateral - Native - Cortex - Follicular phase - Ipsilateral - Nativ
## Medulla - Follicular phase - Ipsilateral - Decellularized - Cortex - Follicular phase - Ipsilateral -
## Medulla - Follicular phase - Ipsilateral - Native - Cortex - Follicular phase - Ipsilateral - Native
## Medulla - Luteal phase - Contralateral - Decellularized - Cortex - Follicular phase - Ipsilateral -
## Medulla - Luteal phase - Contralateral - Native - Cortex - Follicular phase - Ipsilateral - Native =
## Medulla - Luteal phase - Ipsilateral - Decellularized - Cortex - Follicular phase - Ipsilateral - Na
## Medulla - Luteal phase - Ipsilateral - Native - Cortex - Follicular phase - Ipsilateral - Native ==
## Cortex - Luteal phase - Contralateral - Native - Cortex - Luteal phase - Contralateral - Decellulari
## Cortex - Luteal phase - Ipsilateral - Decellularized - Cortex - Luteal phase - Contralateral - Decel
## Cortex - Luteal phase - Ipsilateral - Native - Cortex - Luteal phase - Contralateral - Decellularize
## Medulla - Follicular phase - Contralateral - Decellularized - Cortex - Luteal phase - Contralateral -
## Medulla - Follicular phase - Contralateral - Native - Cortex - Luteal phase - Contralateral - Decellu
## Medulla - Follicular phase - Ipsilateral - Decellularized - Cortex - Luteal phase - Contralateral -
## Medulla - Follicular phase - Ipsilateral - Native - Cortex - Luteal phase - Contralateral - Decellula
## Medulla - Luteal phase - Contralateral - Decellularized - Cortex - Luteal phase - Contralateral - Dec
## Medulla - Luteal phase - Contralateral - Native - Cortex - Luteal phase - Contralateral - Decellulari
## Medulla - Luteal phase - Ipsilateral - Decellularized - Cortex - Luteal phase - Contralateral - Decel
## Medulla - Luteal phase - Ipsilateral - Native - Cortex - Luteal phase - Contralateral - Decellularize
## Cortex - Luteal phase - Ipsilateral - Decellularized - Cortex - Luteal phase - Contralateral - Nativ
## Cortex - Luteal phase - Ipsilateral - Native - Cortex - Luteal phase - Contralateral - Native == 0
## Medulla - Follicular phase - Contralateral - Decellularized - Cortex - Luteal phase - Contralateral -
## Medulla - Follicular phase - Contralateral - Native - Cortex - Luteal phase - Contralateral - Native
## Medulla - Follicular phase - Ipsilateral - Decellularized - Cortex - Luteal phase - Contralateral -
## Medulla - Follicular phase - Ipsilateral - Native - Cortex - Luteal phase - Contralateral - Native =
## Medulla - Luteal phase - Contralateral - Decellularized - Cortex - Luteal phase - Contralateral - Na
## Medulla - Luteal phase - Contralateral - Native - Cortex - Luteal phase - Contralateral - Native ==
## Medulla - Luteal phase - Ipsilateral - Decellularized - Cortex - Luteal phase - Contralateral - Nativ
## Medulla - Luteal phase - Ipsilateral - Native - Cortex - Luteal phase - Contralateral - Native == 0
## Cortex - Luteal phase - Ipsilateral - Native - Cortex - Luteal phase - Ipsilateral - Decellularized =
## Medulla - Follicular phase - Contralateral - Decellularized - Cortex - Luteal phase - Ipsilateral -
## Medulla - Follicular phase - Contralateral - Native - Cortex - Luteal phase - Ipsilateral - Decellula
## Medulla - Follicular phase - Ipsilateral - Decellularized - Cortex - Luteal phase - Ipsilateral - Dec
## Medulla - Follicular phase - Ipsilateral - Native - Cortex - Luteal phase - Ipsilateral - Decellulari
## Medulla - Luteal phase - Contralateral - Decellularized - Cortex - Luteal phase - Ipsilateral - Decel
## Medulla - Luteal phase - Contralateral - Native - Cortex - Luteal phase - Ipsilateral - Decellularize
## Medulla - Luteal phase - Ipsilateral - Decellularized - Cortex - Luteal phase - Ipsilateral - Decellu
## Medulla - Luteal phase - Ipsilateral - Native - Cortex - Luteal phase - Ipsilateral - Decellularized
## Medulla - Follicular phase - Contralateral - Decellularized - Cortex - Luteal phase - Ipsilateral -
## Medulla - Follicular phase - Contralateral - Native - Cortex - Luteal phase - Ipsilateral - Native =
## Medulla - Follicular phase - Ipsilateral - Decellularized - Cortex - Luteal phase - Ipsilateral - Na
## Medulla - Follicular phase - Ipsilateral - Native - Cortex - Luteal phase - Ipsilateral - Native ==
## Medulla - Luteal phase - Contralateral - Decellularized - Cortex - Luteal phase - Ipsilateral - Nativ
## Medulla - Luteal phase - Contralateral - Native - Cortex - Luteal phase - Ipsilateral - Native == 0
## Medulla - Luteal phase - Ipsilateral - Decellularized - Cortex - Luteal phase - Ipsilateral - Native
## Medulla - Luteal phase - Ipsilateral - Native - Cortex - Luteal phase - Ipsilateral - Native == 0
## Medulla - Follicular phase - Contralateral - Native - Medulla - Follicular phase - Contralateral - De
## Medulla - Follicular phase - Ipsilateral - Decellularized - Medulla - Follicular phase - Contralatera
## Medulla - Follicular phase - Ipsilateral - Native - Medulla - Follicular phase - Contralateral - Dec
## Medulla - Luteal phase - Contralateral - Decellularized - Medulla - Follicular phase - Contralateral
## Medulla - Luteal phase - Contralateral - Native - Medulla - Follicular phase - Contralateral - Decel
## Medulla - Luteal phase - Ipsilateral - Decellularized - Medulla - Follicular phase - Contralateral -
## Medulla - Luteal phase - Ipsilateral - Native - Medulla - Follicular phase - Contralateral - Decellul
## Medulla - Follicular phase - Ipsilateral - Decellularized - Medulla - Follicular phase - Contralatera
## Medulla - Follicular phase - Ipsilateral - Native - Medulla - Follicular phase - Contralateral - Nati
```

```
## Medulla - Luteal phase - Contralateral - Decellularized - Medulla - Follicular phase - Contralateral
## Medulla - Luteal phase - Contralateral - Native - Medulla - Follicular phase - Contralateral - Nativ
## Medulla - Luteal phase - Ipsilateral - Decellularized - Medulla - Follicular phase - Contralateral -
## Medulla - Luteal phase - Ipsilateral - Native - Medulla - Follicular phase - Contralateral - Native =
## Medulla - Follicular phase - Ipsilateral - Native - Medulla - Follicular phase - Ipsilateral - Decel
## Medulla - Luteal phase - Contralateral - Decellularized - Medulla - Follicular phase - Ipsilateral -
## Medulla - Luteal phase - Contralateral - Native - Medulla - Follicular phase - Ipsilateral - Decellu
## Medulla - Luteal phase - Ipsilateral - Decellularized - Medulla - Follicular phase - Ipsilateral - De
## Medulla - Luteal phase - Ipsilateral - Native - Medulla - Follicular phase - Ipsilateral - Decellula
## Medulla - Luteal phase - Contralateral - Decellularized - Medulla - Follicular phase - Ipsilateral -
## Medulla - Luteal phase - Contralateral - Native - Medulla - Follicular phase - Ipsilateral - Native =
## Medulla - Luteal phase - Ipsilateral - Decellularized - Medulla - Follicular phase - Ipsilateral - Na
## Medulla - Luteal phase - Ipsilateral - Native - Medulla - Follicular phase - Ipsilateral - Native ==
## Medulla - Luteal phase - Contralateral - Native - Medulla - Luteal phase - Contralateral - Decellula
## Medulla - Luteal phase - Ipsilateral - Decellularized - Medulla - Luteal phase - Contralateral - Dec
## Medulla - Luteal phase - Ipsilateral - Native - Medulla - Luteal phase - Contralateral - Decellulari
## Medulla - Luteal phase - Ipsilateral - Decellularized - Medulla - Luteal phase - Contralateral - Nati
## Medulla - Luteal phase - Ipsilateral - Native - Medulla - Luteal phase - Contralateral - Native == 0
## Medulla - Luteal phase - Ipsilateral - Native - Medulla - Luteal phase - Ipsilateral - Decellularize
##
## Cortex - Follicular phase - Contralateral - Native - Cortex - Follicular phase - Contralateral - Dec
## Cortex - Follicular phase - Ipsilateral - Decellularized - Cortex - Follicular phase - Contralateral
## Cortex - Follicular phase - Ipsilateral - Native - Cortex - Follicular phase - Contralateral - Decel
## Cortex - Luteal phase - Contralateral - Decellularized - Cortex - Follicular phase - Contralateral -
## Cortex - Luteal phase - Contralateral - Native - Cortex - Follicular phase - Contralateral - Decellu
## Cortex - Luteal phase - Ipsilateral - Decellularized - Cortex - Follicular phase - Contralateral - De
## Cortex - Luteal phase - Ipsilateral - Native - Cortex - Follicular phase - Contralateral - Decellula
## Medulla - Follicular phase - Contralateral - Decellularized - Cortex - Follicular phase - Contralate
## Medulla - Follicular phase - Contralateral - Native - Cortex - Follicular phase - Contralateral - De
## Medulla - Follicular phase - Ipsilateral - Decellularized - Cortex - Follicular phase - Contralatera
## Medulla - Follicular phase - Ipsilateral - Native - Cortex - Follicular phase - Contralateral - Decel
## Medulla - Luteal phase - Contralateral - Decellularized - Cortex - Follicular phase - Contralateral
## Medulla - Luteal phase - Contralateral - Native - Cortex - Follicular phase - Contralateral - Decellu
## Medulla - Luteal phase - Ipsilateral - Decellularized - Cortex - Follicular phase - Contralateral -
## Medulla - Luteal phase - Ipsilateral - Native - Cortex - Follicular phase - Contralateral - Decellula
## Cortex - Follicular phase - Ipsilateral - Decellularized - Cortex - Follicular phase - Contralateral
## Cortex - Follicular phase - Ipsilateral - Native - Cortex - Follicular phase - Contralateral - Native
## Cortex - Luteal phase - Contralateral - Decellularized - Cortex - Follicular phase - Contralateral -
## Cortex - Luteal phase - Contralateral - Native - Cortex - Follicular phase - Contralateral - Native =
## Cortex - Luteal phase - Ipsilateral - Decellularized - Cortex - Follicular phase - Contralateral - Na
## Cortex - Luteal phase - Ipsilateral - Native - Cortex - Follicular phase - Contralateral - Native ==
## Medulla - Follicular phase - Contralateral - Decellularized - Cortex - Follicular phase - Contralate
## Medulla - Follicular phase - Contralateral - Native - Cortex - Follicular phase - Contralateral - Na
## Medulla - Follicular phase - Ipsilateral - Decellularized - Cortex - Follicular phase - Contralatera
## Medulla - Follicular phase - Ipsilateral - Native - Cortex - Follicular phase - Contralateral - Nati
## Medulla - Luteal phase - Contralateral - Decellularized - Cortex - Follicular phase - Contralateral
## Medulla - Luteal phase - Contralateral - Native - Cortex - Follicular phase - Contralateral - Native
## Medulla - Luteal phase - Ipsilateral - Decellularized - Cortex - Follicular phase - Contralateral -
## Medulla - Luteal phase - Ipsilateral - Native - Cortex - Follicular phase - Contralateral - Native ==
## Cortex - Follicular phase - Ipsilateral - Native - Cortex - Follicular phase - Ipsilateral - Decellu
## Cortex - Luteal phase - Contralateral - Decellularized - Cortex - Follicular phase - Ipsilateral - De
## Cortex - Luteal phase - Contralateral - Native - Cortex - Follicular phase - Ipsilateral - Decellula
## Cortex - Luteal phase - Ipsilateral - Decellularized - Cortex - Follicular phase - Ipsilateral - Dec
## Cortex - Luteal phase - Ipsilateral - Native - Cortex - Follicular phase - Ipsilateral - Decellulari
```

```
## Medulla - Follicular phase - Contralateral - Decellularized - Cortex - Follicular phase - Ipsilateral
## Medulla - Follicular phase - Contralateral - Native - Cortex - Follicular phase - Ipsilateral - Decel
## Medulla - Follicular phase - Ipsilateral - Decellularized - Cortex - Follicular phase - Ipsilateral -
## Medulla - Follicular phase - Ipsilateral - Native - Cortex - Follicular phase - Ipsilateral - Decellu
## Medulla - Luteal phase - Contralateral - Decellularized - Cortex - Follicular phase - Ipsilateral - I
## Medulla - Luteal phase - Contralateral - Native - Cortex - Follicular phase - Ipsilateral - Decellula
## Medulla - Luteal phase - Ipsilateral - Decellularized - Cortex - Follicular phase - Ipsilateral - Dec
## Medulla - Luteal phase - Ipsilateral - Native - Cortex - Follicular phase - Ipsilateral - Decellular
## Cortex - Luteal phase - Contralateral - Decellularized - Cortex - Follicular phase - Ipsilateral - Na
## Cortex - Luteal phase - Contralateral - Native - Cortex - Follicular phase - Ipsilateral - Native ==
## Cortex - Luteal phase - Ipsilateral - Decellularized - Cortex - Follicular phase - Ipsilateral - Nat
## Cortex - Luteal phase - Ipsilateral - Native - Cortex - Follicular phase - Ipsilateral - Native == 0
## Medulla - Follicular phase - Contralateral - Decellularized - Cortex - Follicular phase - Ipsilateral
## Medulla - Follicular phase - Contralateral - Native - Cortex - Follicular phase - Ipsilateral - Nativ
## Medulla - Follicular phase - Ipsilateral - Decellularized - Cortex - Follicular phase - Ipsilateral -
## Medulla - Follicular phase - Ipsilateral - Native - Cortex - Follicular phase - Ipsilateral - Native
## Medulla - Luteal phase - Contralateral - Decellularized - Cortex - Follicular phase - Ipsilateral - I
## Medulla - Luteal phase - Contralateral - Native - Cortex - Follicular phase - Ipsilateral - Native =
## Medulla - Luteal phase - Ipsilateral - Decellularized - Cortex - Follicular phase - Ipsilateral - Na
## Medulla - Luteal phase - Ipsilateral - Native - Cortex - Follicular phase - Ipsilateral - Native == 0
## Cortex - Luteal phase - Contralateral - Native - Cortex - Luteal phase - Contralateral - Decellulari
## Cortex - Luteal phase - Ipsilateral - Decellularized - Cortex - Luteal phase - Contralateral - Decel
## Cortex - Luteal phase - Ipsilateral - Native - Cortex - Luteal phase - Contralateral - Decellularized
## Medulla - Follicular phase - Contralateral - Decellularized - Cortex - Luteal phase - Contralateral -
## Medulla - Follicular phase - Contralateral - Native - Cortex - Luteal phase - Contralateral - Decellu
## Medulla - Follicular phase - Ipsilateral - Decellularized - Cortex - Luteal phase - Contralateral - I
## Medulla - Follicular phase - Ipsilateral - Native - Cortex - Luteal phase - Contralateral - Decellula
## Medulla - Luteal phase - Contralateral - Decellularized - Cortex - Luteal phase - Contralateral - Dec
## Medulla - Luteal phase - Contralateral - Native - Cortex - Luteal phase - Contralateral - Decellulari
## Medulla - Luteal phase - Ipsilateral - Decellularized - Cortex - Luteal phase - Contralateral - Decel
## Medulla - Luteal phase - Ipsilateral - Native - Cortex - Luteal phase - Contralateral - Decellularize
## Cortex - Luteal phase - Ipsilateral - Decellularized - Cortex - Luteal phase - Contralateral - Native
## Cortex - Luteal phase - Ipsilateral - Native - Cortex - Luteal phase - Contralateral - Native == 0
## Medulla - Follicular phase - Contralateral - Decellularized - Cortex - Luteal phase - Contralateral -
## Medulla - Follicular phase - Contralateral - Native - Cortex - Luteal phase - Contralateral - Native
## Medulla - Follicular phase - Ipsilateral - Decellularized - Cortex - Luteal phase - Contralateral - I
## Medulla - Follicular phase - Ipsilateral - Native - Cortex - Luteal phase - Contralateral - Native =
## Medulla - Luteal phase - Contralateral - Decellularized - Cortex - Luteal phase - Contralateral - Na
## Medulla - Luteal phase - Contralateral - Native - Cortex - Luteal phase - Contralateral - Native == 0
## Medulla - Luteal phase - Ipsilateral - Decellularized - Cortex - Luteal phase - Contralateral - Nati
## Medulla - Luteal phase - Ipsilateral - Native - Cortex - Luteal phase - Contralateral - Native == 0
## Cortex - Luteal phase - Ipsilateral - Native - Cortex - Luteal phase - Ipsilateral - Decellularized =
## Medulla - Follicular phase - Contralateral - Decellularized - Cortex - Luteal phase - Ipsilateral - I
## Medulla - Follicular phase - Contralateral - Native - Cortex - Luteal phase - Ipsilateral - Decellula
## Medulla - Follicular phase - Ipsilateral - Decellularized - Cortex - Luteal phase - Ipsilateral - Dec
## Medulla - Follicular phase - Ipsilateral - Native - Cortex - Luteal phase - Ipsilateral - Decellular
## Medulla - Luteal phase - Contralateral - Decellularized - Cortex - Luteal phase - Ipsilateral - Decel
## Medulla - Luteal phase - Contralateral - Native - Cortex - Luteal phase - Ipsilateral - Decellularize
## Medulla - Luteal phase - Ipsilateral - Decellularized - Cortex - Luteal phase - Ipsilateral - Decellu
## Medulla - Luteal phase - Ipsilateral - Native - Cortex - Luteal phase - Ipsilateral - Decellularized
## Medulla - Follicular phase - Contralateral - Decellularized - Cortex - Luteal phase - Ipsilateral - I
## Medulla - Follicular phase - Contralateral - Native - Cortex - Luteal phase - Ipsilateral - Native =
## Medulla - Follicular phase - Ipsilateral - Decellularized - Cortex - Luteal phase - Ipsilateral - Na
## Medulla - Follicular phase - Ipsilateral - Native - Cortex - Luteal phase - Ipsilateral - Native == 0
```

```
## Medulla - Luteal phase - Contralateral - Decellularized - Cortex - Luteal phase - Ipsilateral - Nati
## Medulla - Luteal phase - Contralateral - Native - Cortex - Luteal phase - Ipsilateral - Native == 0
## Medulla - Luteal phase - Ipsilateral - Decellularized - Cortex - Luteal phase - Ipsilateral - Native
## Medulla - Luteal phase - Ipsilateral - Native - Cortex - Luteal phase - Ipsilateral - Native == 0
## Medulla - Follicular phase - Contralateral - Native - Medulla - Follicular phase - Contralateral - De
## Medulla - Follicular phase - Ipsilateral - Decellularized - Medulla - Follicular phase - Contralatera
## Medulla - Follicular phase - Ipsilateral - Native - Medulla - Follicular phase - Contralateral - Dec
## Medulla - Luteal phase - Contralateral - Decellularized - Medulla - Follicular phase - Contralateral
## Medulla - Luteal phase - Contralateral - Native - Medulla - Follicular phase - Contralateral - Decel
## Medulla - Luteal phase - Ipsilateral - Decellularized - Medulla - Follicular phase - Contralateral -
## Medulla - Luteal phase - Ipsilateral - Native - Medulla - Follicular phase - Contralateral - Decellu
## Medulla - Follicular phase - Ipsilateral - Decellularized - Medulla - Follicular phase - Contralatera
## Medulla - Follicular phase - Ipsilateral - Native - Medulla - Follicular phase - Contralateral - Nat
## Medulla - Luteal phase - Contralateral - Decellularized - Medulla - Follicular phase - Contralateral
## Medulla - Luteal phase - Contralateral - Native - Medulla - Follicular phase - Contralateral - Nativ
## Medulla - Luteal phase - Ipsilateral - Decellularized - Medulla - Follicular phase - Contralateral -
## Medulla - Luteal phase - Ipsilateral - Native - Medulla - Follicular phase - Contralateral - Native
## Medulla - Follicular phase - Ipsilateral - Native - Medulla - Follicular phase - Ipsilateral - Decel
## Medulla - Luteal phase - Contralateral - Decellularized - Medulla - Follicular phase - Ipsilateral -
## Medulla - Luteal phase - Contralateral - Native - Medulla - Follicular phase - Ipsilateral - Decellu
## Medulla - Luteal phase - Ipsilateral - Decellularized - Medulla - Follicular phase - Ipsilateral - De
## Medulla - Luteal phase - Ipsilateral - Native - Medulla - Follicular phase - Ipsilateral - Decellula
## Medulla - Luteal phase - Contralateral - Decellularized - Medulla - Follicular phase - Ipsilateral -
## Medulla - Luteal phase - Contralateral - Native - Medulla - Follicular phase - Ipsilateral - Native
## Medulla - Luteal phase - Ipsilateral - Decellularized - Medulla - Follicular phase - Ipsilateral - N
## Medulla - Luteal phase - Ipsilateral - Native - Medulla - Follicular phase - Ipsilateral - Native ==
## Medulla - Luteal phase - Contralateral - Native - Medulla - Luteal phase - Contralateral - Decellula
## Medulla - Luteal phase - Ipsilateral - Decellularized - Medulla - Luteal phase - Contralateral - Dec
## Medulla - Luteal phase - Ipsilateral - Native - Medulla - Luteal phase - Contralateral - Decellulari
## Medulla - Luteal phase - Ipsilateral - Decellularized - Medulla - Luteal phase - Contralateral - Nat
## Medulla - Luteal phase - Ipsilateral - Native - Medulla - Luteal phase - Contralateral - Native == 0
## Medulla - Luteal phase - Ipsilateral - Native - Medulla - Luteal phase - Ipsilateral - Decellularize
##
## Cortex - Follicular phase - Contralateral - Native - Cortex - Follicular phase - Contralateral - Dec
## Cortex - Follicular phase - Ipsilateral - Decellularized - Cortex - Follicular phase - Contralateral
## Cortex - Follicular phase - Ipsilateral - Native - Cortex - Follicular phase - Contralateral - Decel
## Cortex - Luteal phase - Contralateral - Decellularized - Cortex - Follicular phase - Contralateral -
## Cortex - Luteal phase - Contralateral - Native - Cortex - Follicular phase - Contralateral - Decellu
## Cortex - Luteal phase - Ipsilateral - Decellularized - Cortex - Follicular phase - Contralateral - De
## Cortex - Luteal phase - Ipsilateral - Native - Cortex - Follicular phase - Contralateral - Decellula
## Medulla - Follicular phase - Contralateral - Decellularized - Cortex - Follicular phase - Contralater
## Medulla - Follicular phase - Contralateral - Native - Cortex - Follicular phase - Contralateral - Dec
## Medulla - Follicular phase - Ipsilateral - Decellularized - Cortex - Follicular phase - Contralateral
## Medulla - Follicular phase - Ipsilateral - Native - Cortex - Follicular phase - Contralateral - Decel
## Medulla - Luteal phase - Contralateral - Decellularized - Cortex - Follicular phase - Contralateral -
## Medulla - Luteal phase - Contralateral - Native - Cortex - Follicular phase - Contralateral - Decellu
## Medulla - Luteal phase - Ipsilateral - Decellularized - Cortex - Follicular phase - Contralateral - D
## Medulla - Luteal phase - Ipsilateral - Native - Cortex - Follicular phase - Contralateral - Decellula
## Cortex - Follicular phase - Ipsilateral - Decellularized - Cortex - Follicular phase - Contralateral
## Cortex - Follicular phase - Ipsilateral - Native - Cortex - Follicular phase - Contralateral - Nativ
## Cortex - Luteal phase - Contralateral - Decellularized - Cortex - Follicular phase - Contralateral -
## Cortex - Luteal phase - Contralateral - Native - Cortex - Follicular phase - Contralateral - Native
## Cortex - Luteal phase - Ipsilateral - Decellularized - Cortex - Follicular phase - Contralateral - Na
## Cortex - Luteal phase - Ipsilateral - Native - Cortex - Follicular phase - Contralateral - Native ==
```

```
## Medulla - Follicular phase - Contralateral - Decellularized - Cortex - Follicular phase - Contralater
## Medulla - Follicular phase - Contralateral - Native - Cortex - Follicular phase - Contralateral - Nat
## Medulla - Follicular phase - Ipsilateral - Decellularized - Cortex - Follicular phase - Contralateral
## Medulla - Follicular phase - Ipsilateral - Native - Cortex - Follicular phase - Contralateral - Nati
## Medulla - Luteal phase - Contralateral - Decellularized - Cortex - Follicular phase - Contralateral -
## Medulla - Luteal phase - Contralateral - Native - Cortex - Follicular phase - Contralateral - Native
## Medulla - Luteal phase - Ipsilateral - Decellularized - Cortex - Follicular phase - Contralateral - N
## Medulla - Luteal phase - Ipsilateral - Native - Cortex - Follicular phase - Contralateral - Native =
## Cortex - Follicular phase - Ipsilateral - Native - Cortex - Follicular phase - Ipsilateral - Decellu
## Cortex - Luteal phase - Contralateral - Decellularized - Cortex - Follicular phase - Ipsilateral - De
## Cortex - Luteal phase - Contralateral - Native - Cortex - Follicular phase - Ipsilateral - Decellula
## Cortex - Luteal phase - Ipsilateral - Decellularized - Cortex - Follicular phase - Ipsilateral - Dec
## Cortex - Luteal phase - Ipsilateral - Native - Cortex - Follicular phase - Ipsilateral - Decellulari
## Medulla - Follicular phase - Contralateral - Decellularized - Cortex - Follicular phase - Ipsilateral
## Medulla - Follicular phase - Contralateral - Native - Cortex - Follicular phase - Ipsilateral - Decel
## Medulla - Follicular phase - Ipsilateral - Decellularized - Cortex - Follicular phase - Ipsilateral -
## Medulla - Follicular phase - Ipsilateral - Native - Cortex - Follicular phase - Ipsilateral - Decellu
## Medulla - Luteal phase - Contralateral - Decellularized - Cortex - Follicular phase - Ipsilateral - D
## Medulla - Luteal phase - Contralateral - Native - Cortex - Follicular phase - Ipsilateral - Decellula
## Medulla - Luteal phase - Ipsilateral - Decellularized - Cortex - Follicular phase - Ipsilateral - Dec
## Medulla - Luteal phase - Ipsilateral - Native - Cortex - Follicular phase - Ipsilateral - Decellular
## Cortex - Luteal phase - Contralateral - Decellularized - Cortex - Follicular phase - Ipsilateral - Na
## Cortex - Luteal phase - Contralateral - Native - Cortex - Follicular phase - Ipsilateral - Native ==
## Cortex - Luteal phase - Ipsilateral - Decellularized - Cortex - Follicular phase - Ipsilateral - Nati
## Cortex - Luteal phase - Ipsilateral - Native - Cortex - Follicular phase - Ipsilateral - Native == 0
## Medulla - Follicular phase - Contralateral - Decellularized - Cortex - Follicular phase - Ipsilateral
## Medulla - Follicular phase - Contralateral - Native - Cortex - Follicular phase - Ipsilateral - Nativ
## Medulla - Follicular phase - Ipsilateral - Decellularized - Cortex - Follicular phase - Ipsilateral -
## Medulla - Follicular phase - Ipsilateral - Native - Cortex - Follicular phase - Ipsilateral - Native
## Medulla - Luteal phase - Contralateral - Decellularized - Cortex - Follicular phase - Ipsilateral - N
## Medulla - Luteal phase - Contralateral - Native - Cortex - Follicular phase - Ipsilateral - Native =
## Medulla - Luteal phase - Ipsilateral - Decellularized - Cortex - Follicular phase - Ipsilateral - Nat
## Medulla - Luteal phase - Ipsilateral - Native - Cortex - Follicular phase - Ipsilateral - Native == 0
## Cortex - Luteal phase - Contralateral - Native - Cortex - Luteal phase - Contralateral - Decellulari
## Cortex - Luteal phase - Ipsilateral - Decellularized - Cortex - Luteal phase - Contralateral - Decell
## Cortex - Luteal phase - Ipsilateral - Native - Cortex - Luteal phase - Contralateral - Decellularized
## Medulla - Follicular phase - Contralateral - Decellularized - Cortex - Luteal phase - Contralateral -
## Medulla - Follicular phase - Contralateral - Native - Cortex - Luteal phase - Contralateral - Decellu
## Medulla - Follicular phase - Ipsilateral - Decellularized - Cortex - Luteal phase - Contralateral - D
## Medulla - Follicular phase - Ipsilateral - Native - Cortex - Luteal phase - Contralateral - Decellula
## Medulla - Luteal phase - Contralateral - Decellularized - Cortex - Luteal phase - Contralateral - Dec
## Medulla - Luteal phase - Contralateral - Native - Cortex - Luteal phase - Contralateral - Decellular
## Medulla - Luteal phase - Ipsilateral - Decellularized - Cortex - Luteal phase - Contralateral - Decel
## Medulla - Luteal phase - Ipsilateral - Native - Cortex - Luteal phase - Contralateral - Decellularize
## Cortex - Luteal phase - Ipsilateral - Decellularized - Cortex - Luteal phase - Contralateral - Native
## Cortex - Luteal phase - Ipsilateral - Native - Cortex - Luteal phase - Contralateral - Native == 0
## Medulla - Follicular phase - Contralateral - Decellularized - Cortex - Luteal phase - Contralateral -
## Medulla - Follicular phase - Contralateral - Native - Cortex - Luteal phase - Contralateral - Native
## Medulla - Follicular phase - Ipsilateral - Decellularized - Cortex - Luteal phase - Contralateral - N
## Medulla - Follicular phase - Ipsilateral - Native - Cortex - Luteal phase - Contralateral - Native =
## Medulla - Luteal phase - Contralateral - Decellularized - Cortex - Luteal phase - Contralateral - Nat
## Medulla - Luteal phase - Contralateral - Native - Cortex - Luteal phase - Contralateral - Native == 0
## Medulla - Luteal phase - Ipsilateral - Decellularized - Cortex - Luteal phase - Contralateral - Nati
## Medulla - Luteal phase - Ipsilateral - Native - Cortex - Luteal phase - Contralateral - Native == 0
```

```
## Cortex - Luteal phase - Ipsilateral - Native - Cortex - Luteal phase - Ipsilateral - Decellularized =
## Medulla - Follicular phase - Contralateral - Decellularized - Cortex - Luteal phase - Ipsilateral - 
## Medulla - Follicular phase - Contralateral - Native - Cortex - Luteal phase - Ipsilateral - Decellula
## Medulla - Follicular phase - Ipsilateral - Decellularized - Cortex - Luteal phase - Ipsilateral - Dec
## Medulla - Follicular phase - Ipsilateral - Native - Cortex - Luteal phase - Ipsilateral - Decellulari
## Medulla - Luteal phase - Contralateral - Decellularized - Cortex - Luteal phase - Ipsilateral - Decel
## Medulla - Luteal phase - Contralateral - Native - Cortex - Luteal phase - Ipsilateral - Decellulariz
## Medulla - Luteal phase - Ipsilateral - Decellularized - Cortex - Luteal phase - Ipsilateral - Decellu
## Medulla - Luteal phase - Ipsilateral - Native - Cortex - Luteal phase - Ipsilateral - Decellularized
## Medulla - Follicular phase - Contralateral - Decellularized - Cortex - Luteal phase - Ipsilateral - 
## Medulla - Follicular phase - Contralateral - Native - Cortex - Luteal phase - Ipsilateral - Native =
## Medulla - Follicular phase - Ipsilateral - Decellularized - Cortex - Luteal phase - Ipsilateral - Na
## Medulla - Follicular phase - Ipsilateral - Native - Cortex - Luteal phase - Ipsilateral - Native == 
## Medulla - Luteal phase - Contralateral - Decellularized - Cortex - Luteal phase - Ipsilateral - Nati
## Medulla - Luteal phase - Contralateral - Native - Cortex - Luteal phase - Ipsilateral - Native == 0
## Medulla - Luteal phase - Ipsilateral - Decellularized - Cortex - Luteal phase - Ipsilateral - Native
## Medulla - Luteal phase - Ipsilateral - Native - Cortex - Luteal phase - Ipsilateral - Native == 0
## Medulla - Follicular phase - Contralateral - Native - Medulla - Follicular phase - Contralateral - De
## Medulla - Follicular phase - Ipsilateral - Decellularized - Medulla - Follicular phase - Contralatera
## Medulla - Follicular phase - Ipsilateral - Native - Medulla - Follicular phase - Contralateral - Dec
## Medulla - Luteal phase - Contralateral - Decellularized - Medulla - Follicular phase - Contralateral
## Medulla - Luteal phase - Contralateral - Native - Medulla - Follicular phase - Contralateral - Decel
## Medulla - Luteal phase - Ipsilateral - Decellularized - Medulla - Follicular phase - Contralateral -
## Medulla - Luteal phase - Ipsilateral - Native - Medulla - Follicular phase - Contralateral - Decellu
## Medulla - Follicular phase - Ipsilateral - Decellularized - Medulla - Follicular phase - Contralatera
## Medulla - Follicular phase - Ipsilateral - Native - Medulla - Follicular phase - Contralateral - Nati
## Medulla - Luteal phase - Contralateral - Decellularized - Medulla - Follicular phase - Contralateral
## Medulla - Luteal phase - Contralateral - Native - Medulla - Follicular phase - Contralateral - Native
## Medulla - Luteal phase - Ipsilateral - Decellularized - Medulla - Follicular phase - Contralateral -
## Medulla - Luteal phase - Ipsilateral - Native - Medulla - Follicular phase - Contralateral - Native =
## Medulla - Follicular phase - Ipsilateral - Native - Medulla - Follicular phase - Ipsilateral - Decel
## Medulla - Luteal phase - Contralateral - Decellularized - Medulla - Follicular phase - Ipsilateral -
## Medulla - Luteal phase - Contralateral - Native - Medulla - Follicular phase - Ipsilateral - Decellu
## Medulla - Luteal phase - Ipsilateral - Decellularized - Medulla - Follicular phase - Ipsilateral - De
## Medulla - Luteal phase - Ipsilateral - Native - Medulla - Follicular phase - Ipsilateral - Decellula
## Medulla - Luteal phase - Contralateral - Decellularized - Medulla - Follicular phase - Ipsilateral -
## Medulla - Luteal phase - Contralateral - Native - Medulla - Follicular phase - Ipsilateral - Native =
## Medulla - Luteal phase - Ipsilateral - Decellularized - Medulla - Follicular phase - Ipsilateral - Na
## Medulla - Luteal phase - Ipsilateral - Native - Medulla - Follicular phase - Ipsilateral - Native ==
## Medulla - Luteal phase - Contralateral - Native - Medulla - Luteal phase - Contralateral - Decellula
## Medulla - Luteal phase - Ipsilateral - Decellularized - Medulla - Luteal phase - Contralateral - Dec
## Medulla - Luteal phase - Ipsilateral - Native - Medulla - Luteal phase - Contralateral - Decellulari
## Medulla - Luteal phase - Ipsilateral - Decellularized - Medulla - Luteal phase - Contralateral - Nati
## Medulla - Luteal phase - Ipsilateral - Native - Medulla - Luteal phase - Contralateral - Native == 0
## Medulla - Luteal phase - Ipsilateral - Native - Medulla - Luteal phase - Ipsilateral - Decellularize
##
## Cortex - Follicular phase - Contralateral - Native - Cortex - Follicular phase - Contralateral - Dec
## Cortex - Follicular phase - Ipsilateral - Decellularized - Cortex - Follicular phase - Contralateral
## Cortex - Follicular phase - Ipsilateral - Native - Cortex - Follicular phase - Contralateral - Decel
## Cortex - Luteal phase - Contralateral - Decellularized - Cortex - Follicular phase - Contralateral -
## Cortex - Luteal phase - Contralateral - Native - Cortex - Follicular phase - Contralateral - Decellu
## Cortex - Luteal phase - Ipsilateral - Decellularized - Cortex - Follicular phase - Contralateral - De
## Cortex - Luteal phase - Ipsilateral - Native - Cortex - Follicular phase - Contralateral - Decellula
## Medulla - Follicular phase - Contralateral - Decellularized - Cortex - Follicular phase - Contralater
```

```
## Medulla - Follicular phase - Contralateral - Native - Cortex - Follicular phase - Contralateral - Dec
## Medulla - Follicular phase - Ipsilateral - Decellularized - Cortex - Follicular phase - Contralateral
## Medulla - Follicular phase - Ipsilateral - Native - Cortex - Follicular phase - Contralateral - Decel
## Medulla - Luteal phase - Contralateral - Decellularized - Cortex - Follicular phase - Contralateral -
## Medulla - Luteal phase - Contralateral - Native - Cortex - Follicular phase - Contralateral - Decellu
## Medulla - Luteal phase - Ipsilateral - Decellularized - Cortex - Follicular phase - Contralateral -
## Medulla - Luteal phase - Ipsilateral - Native - Cortex - Follicular phase - Contralateral - Decellula
## Cortex - Follicular phase - Ipsilateral - Decellularized - Cortex - Follicular phase - Contralateral
## Cortex - Follicular phase - Ipsilateral - Native - Cortex - Follicular phase - Contralateral - Native
## Cortex - Luteal phase - Contralateral - Decellularized - Cortex - Follicular phase - Contralateral -
## Cortex - Luteal phase - Contralateral - Native - Cortex - Follicular phase - Contralateral - Native =
## Cortex - Luteal phase - Ipsilateral - Decellularized - Cortex - Follicular phase - Contralateral - Na
## Cortex - Luteal phase - Ipsilateral - Native - Cortex - Follicular phase - Contralateral - Native ==
## Medulla - Follicular phase - Contralateral - Decellularized - Cortex - Follicular phase - Contralater
## Medulla - Follicular phase - Contralateral - Native - Cortex - Follicular phase - Contralateral - Na
## Medulla - Follicular phase - Ipsilateral - Decellularized - Cortex - Follicular phase - Contralateral
## Medulla - Follicular phase - Ipsilateral - Native - Cortex - Follicular phase - Contralateral - Nativ
## Medulla - Luteal phase - Contralateral - Decellularized - Cortex - Follicular phase - Contralateral -
## Medulla - Luteal phase - Contralateral - Native - Cortex - Follicular phase - Contralateral - Native
## Medulla - Luteal phase - Ipsilateral - Decellularized - Cortex - Follicular phase - Contralateral -
## Medulla - Luteal phase - Ipsilateral - Native - Cortex - Follicular phase - Contralateral - Native ==
## Cortex - Follicular phase - Ipsilateral - Native - Cortex - Follicular phase - Ipsilateral - Decellu
## Cortex - Luteal phase - Contralateral - Decellularized - Cortex - Follicular phase - Ipsilateral - De
## Cortex - Luteal phase - Contralateral - Native - Cortex - Follicular phase - Ipsilateral - Decellula
## Cortex - Luteal phase - Ipsilateral - Decellularized - Cortex - Follicular phase - Ipsilateral - Dec
## Cortex - Luteal phase - Ipsilateral - Native - Cortex - Follicular phase - Ipsilateral - Decellulari
## Medulla - Follicular phase - Contralateral - Decellularized - Cortex - Follicular phase - Ipsilateral
## Medulla - Follicular phase - Contralateral - Native - Cortex - Follicular phase - Ipsilateral - Decel
## Medulla - Follicular phase - Ipsilateral - Decellularized - Cortex - Follicular phase - Ipsilateral -
## Medulla - Follicular phase - Ipsilateral - Native - Cortex - Follicular phase - Ipsilateral - Decellu
## Medulla - Luteal phase - Contralateral - Decellularized - Cortex - Follicular phase - Ipsilateral -
## Medulla - Luteal phase - Contralateral - Native - Cortex - Follicular phase - Ipsilateral - Decellula
## Medulla - Luteal phase - Ipsilateral - Decellularized - Cortex - Follicular phase - Ipsilateral - Dec
## Medulla - Luteal phase - Ipsilateral - Native - Cortex - Follicular phase - Ipsilateral - Decellular
## Cortex - Luteal phase - Contralateral - Decellularized - Cortex - Follicular phase - Ipsilateral - Na
## Cortex - Luteal phase - Contralateral - Native - Cortex - Follicular phase - Ipsilateral - Native ==
## Cortex - Luteal phase - Ipsilateral - Decellularized - Cortex - Follicular phase - Ipsilateral - Nati
## Cortex - Luteal phase - Ipsilateral - Native - Cortex - Follicular phase - Ipsilateral - Native == 0
## Medulla - Follicular phase - Contralateral - Decellularized - Cortex - Follicular phase - Ipsilateral
## Medulla - Follicular phase - Contralateral - Native - Cortex - Follicular phase - Ipsilateral - Nativ
## Medulla - Follicular phase - Ipsilateral - Decellularized - Cortex - Follicular phase - Ipsilateral -
## Medulla - Follicular phase - Ipsilateral - Native - Cortex - Follicular phase - Ipsilateral - Native
## Medulla - Luteal phase - Contralateral - Decellularized - Cortex - Follicular phase - Ipsilateral -
## Medulla - Luteal phase - Contralateral - Native - Cortex - Follicular phase - Ipsilateral - Native ==
## Medulla - Luteal phase - Ipsilateral - Decellularized - Cortex - Follicular phase - Ipsilateral - Na
## Medulla - Luteal phase - Ipsilateral - Native - Cortex - Follicular phase - Ipsilateral - Native ==
## Cortex - Luteal phase - Contralateral - Native - Cortex - Luteal phase - Contralateral - Decellulari
## Cortex - Luteal phase - Ipsilateral - Decellularized - Cortex - Luteal phase - Contralateral - Decell
## Cortex - Luteal phase - Ipsilateral - Native - Cortex - Luteal phase - Contralateral - Decellularized
## Medulla - Follicular phase - Contralateral - Decellularized - Cortex - Luteal phase - Contralateral -
## Medulla - Follicular phase - Contralateral - Native - Cortex - Luteal phase - Contralateral - Decellu
## Medulla - Follicular phase - Ipsilateral - Decellularized - Cortex - Luteal phase - Contralateral -
## Medulla - Follicular phase - Ipsilateral - Native - Cortex - Luteal phase - Contralateral - Decellula
## Medulla - Luteal phase - Contralateral - Decellularized - Cortex - Luteal phase - Contralateral - Dec
```

```
## Medulla - Luteal phase - Contralateral - Native - Cortex - Luteal phase - Contralateral - Decellulari
## Medulla - Luteal phase - Ipsilateral - Decellularized - Cortex - Luteal phase - Contralateral - Decel
## Medulla - Luteal phase - Ipsilateral - Native - Cortex - Luteal phase - Contralateral - Decellulariz
## Cortex - Luteal phase - Ipsilateral - Decellularized - Cortex - Luteal phase - Contralateral - Nativ
## Cortex - Luteal phase - Ipsilateral - Native - Cortex - Luteal phase - Contralateral - Native == 0
## Medulla - Follicular phase - Contralateral - Decellularized - Cortex - Luteal phase - Contralateral -
## Medulla - Follicular phase - Contralateral - Native - Cortex - Luteal phase - Contralateral - Native
## Medulla - Follicular phase - Ipsilateral - Decellularized - Cortex - Luteal phase - Contralateral -
## Medulla - Follicular phase - Ipsilateral - Native - Cortex - Luteal phase - Contralateral - Native =
## Medulla - Luteal phase - Contralateral - Decellularized - Cortex - Luteal phase - Contralateral - Na
## Medulla - Luteal phase - Contralateral - Native - Cortex - Luteal phase - Contralateral - Native ==
## Medulla - Luteal phase - Ipsilateral - Decellularized - Cortex - Luteal phase - Contralateral - Nati
## Medulla - Luteal phase - Ipsilateral - Native - Cortex - Luteal phase - Contralateral - Native == 0
## Cortex - Luteal phase - Ipsilateral - Native - Cortex - Luteal phase - Ipsilateral - Decellularized
## Medulla - Follicular phase - Contralateral - Decellularized - Cortex - Luteal phase - Ipsilateral -
## Medulla - Follicular phase - Contralateral - Native - Cortex - Luteal phase - Ipsilateral - Decellula
## Medulla - Follicular phase - Ipsilateral - Decellularized - Cortex - Luteal phase - Ipsilateral - De
## Medulla - Follicular phase - Ipsilateral - Native - Cortex - Luteal phase - Ipsilateral - Decellulari
## Medulla - Luteal phase - Contralateral - Decellularized - Cortex - Luteal phase - Ipsilateral - Decel
## Medulla - Luteal phase - Contralateral - Native - Cortex - Luteal phase - Ipsilateral - Decellulariz
## Medulla - Luteal phase - Ipsilateral - Decellularized - Cortex - Luteal phase - Ipsilateral - Decellu
## Medulla - Luteal phase - Ipsilateral - Native - Cortex - Luteal phase - Ipsilateral - Decellularized
## Medulla - Follicular phase - Contralateral - Decellularized - Cortex - Luteal phase - Ipsilateral -
## Medulla - Follicular phase - Contralateral - Native - Cortex - Luteal phase - Ipsilateral - Native =
## Medulla - Follicular phase - Ipsilateral - Decellularized - Cortex - Luteal phase - Ipsilateral - Na
## Medulla - Follicular phase - Ipsilateral - Native - Cortex - Luteal phase - Ipsilateral - Native ==
## Medulla - Luteal phase - Contralateral - Decellularized - Cortex - Luteal phase - Ipsilateral - Nati
## Medulla - Luteal phase - Contralateral - Native - Cortex - Luteal phase - Ipsilateral - Native == 0
## Medulla - Luteal phase - Ipsilateral - Decellularized - Cortex - Luteal phase - Ipsilateral - Native
## Medulla - Luteal phase - Ipsilateral - Native - Cortex - Luteal phase - Ipsilateral - Native == 0
## Medulla - Follicular phase - Contralateral - Native - Medulla - Follicular phase - Contralateral - De
## Medulla - Follicular phase - Ipsilateral - Decellularized - Medulla - Follicular phase - Contralatera
## Medulla - Follicular phase - Ipsilateral - Native - Medulla - Follicular phase - Contralateral - Dec
## Medulla - Luteal phase - Contralateral - Decellularized - Medulla - Follicular phase - Contralateral
## Medulla - Luteal phase - Contralateral - Native - Medulla - Follicular phase - Contralateral - Decel
## Medulla - Luteal phase - Ipsilateral - Decellularized - Medulla - Follicular phase - Contralateral -
## Medulla - Luteal phase - Ipsilateral - Native - Medulla - Follicular phase - Contralateral - Decellu
## Medulla - Follicular phase - Ipsilateral - Decellularized - Medulla - Follicular phase - Contralatera
## Medulla - Follicular phase - Ipsilateral - Native - Medulla - Follicular phase - Contralateral - Nati
## Medulla - Luteal phase - Contralateral - Decellularized - Medulla - Follicular phase - Contralateral
## Medulla - Luteal phase - Contralateral - Native - Medulla - Follicular phase - Contralateral - Native
## Medulla - Luteal phase - Ipsilateral - Decellularized - Medulla - Follicular phase - Contralateral -
## Medulla - Luteal phase - Ipsilateral - Native - Medulla - Follicular phase - Contralateral - Native =
## Medulla - Follicular phase - Ipsilateral - Native - Medulla - Follicular phase - Ipsilateral - Decell
## Medulla - Luteal phase - Contralateral - Decellularized - Medulla - Follicular phase - Ipsilateral -
## Medulla - Luteal phase - Contralateral - Native - Medulla - Follicular phase - Ipsilateral - Decellul
## Medulla - Luteal phase - Ipsilateral - Decellularized - Medulla - Follicular phase - Ipsilateral - De
## Medulla - Luteal phase - Ipsilateral - Native - Medulla - Follicular phase - Ipsilateral - Decellula
## Medulla - Luteal phase - Contralateral - Decellularized - Medulla - Follicular phase - Ipsilateral -
## Medulla - Luteal phase - Contralateral - Native - Medulla - Follicular phase - Ipsilateral - Native =
## Medulla - Luteal phase - Ipsilateral - Decellularized - Medulla - Follicular phase - Ipsilateral - Na
## Medulla - Luteal phase - Ipsilateral - Native - Medulla - Follicular phase - Ipsilateral - Native ==
## Medulla - Luteal phase - Contralateral - Native - Medulla - Luteal phase - Contralateral - Decellula
## Medulla - Luteal phase - Ipsilateral - Decellularized - Medulla - Luteal phase - Contralateral - Dec
```

```
## Medulla - Luteal phase - Ipsilateral - Native - Medulla - Luteal phase - Contralateral - Decellulari
## Medulla - Luteal phase - Ipsilateral - Decellularized - Medulla - Luteal phase - Contralateral - Nati
## Medulla - Luteal phase - Ipsilateral - Native - Medulla - Luteal phase - Contralateral - Native == 0
## Medulla - Luteal phase - Ipsilateral - Native - Medulla - Luteal phase - Ipsilateral - Decellularize
##
## Cortex - Follicular phase - Contralateral - Native - Cortex - Follicular phase - Contralateral - Dec
## Cortex - Follicular phase - Ipsilateral - Decellularized - Cortex - Follicular phase - Contralateral
## Cortex - Follicular phase - Ipsilateral - Native - Cortex - Follicular phase - Contralateral - Decel
## Cortex - Luteal phase - Contralateral - Decellularized - Cortex - Follicular phase - Contralateral -
## Cortex - Luteal phase - Contralateral - Native - Cortex - Follicular phase - Contralateral - Decellu
## Cortex - Luteal phase - Ipsilateral - Decellularized - Cortex - Follicular phase - Contralateral - De
## Cortex - Luteal phase - Ipsilateral - Native - Cortex - Follicular phase - Contralateral - Decellula
## Medulla - Follicular phase - Contralateral - Decellularized - Cortex - Follicular phase - Contralate
## Medulla - Follicular phase - Contralateral - Native - Cortex - Follicular phase - Contralateral - De
## Medulla - Follicular phase - Ipsilateral - Decellularized - Cortex - Follicular phase - Contralatera
## Medulla - Follicular phase - Ipsilateral - Native - Cortex - Follicular phase - Contralateral - Dece
## Medulla - Luteal phase - Contralateral - Decellularized - Cortex - Follicular phase - Contralateral -
## Medulla - Luteal phase - Contralateral - Native - Cortex - Follicular phase - Contralateral - Decell
## Medulla - Luteal phase - Ipsilateral - Decellularized - Cortex - Follicular phase - Contralateral - 
## Medulla - Luteal phase - Ipsilateral - Native - Cortex - Follicular phase - Contralateral - Decellul
## Cortex - Follicular phase - Ipsilateral - Decellularized - Cortex - Follicular phase - Contralateral
## Cortex - Follicular phase - Ipsilateral - Native - Cortex - Follicular phase - Contralateral - Nativ
## Cortex - Luteal phase - Contralateral - Decellularized - Cortex - Follicular phase - Contralateral -
## Cortex - Luteal phase - Contralateral - Native - Cortex - Follicular phase - Contralateral - Native 
## Cortex - Luteal phase - Ipsilateral - Decellularized - Cortex - Follicular phase - Contralateral - N
## Cortex - Luteal phase - Ipsilateral - Native - Cortex - Follicular phase - Contralateral - Native ==
## Medulla - Follicular phase - Contralateral - Decellularized - Cortex - Follicular phase - Contralate
## Medulla - Follicular phase - Contralateral - Native - Cortex - Follicular phase - Contralateral - Na
## Medulla - Follicular phase - Ipsilateral - Decellularized - Cortex - Follicular phase - Contralatera
## Medulla - Follicular phase - Ipsilateral - Native - Cortex - Follicular phase - Contralateral - Nati
## Medulla - Luteal phase - Contralateral - Decellularized - Cortex - Follicular phase - Contralateral -
## Medulla - Luteal phase - Contralateral - Native - Cortex - Follicular phase - Contralateral - Native
## Medulla - Luteal phase - Ipsilateral - Decellularized - Cortex - Follicular phase - Contralateral - 
## Medulla - Luteal phase - Ipsilateral - Native - Cortex - Follicular phase - Contralateral - Native =
## Cortex - Follicular phase - Ipsilateral - Native - Cortex - Follicular phase - Ipsilateral - Decellu
## Cortex - Luteal phase - Contralateral - Decellularized - Cortex - Follicular phase - Ipsilateral - De
## Cortex - Luteal phase - Contralateral - Native - Cortex - Follicular phase - Ipsilateral - Decellula
## Cortex - Luteal phase - Ipsilateral - Decellularized - Cortex - Follicular phase - Ipsilateral - Dec
## Cortex - Luteal phase - Ipsilateral - Native - Cortex - Follicular phase - Ipsilateral - Decellulari
## Medulla - Follicular phase - Contralateral - Decellularized - Cortex - Follicular phase - Ipsilateral
## Medulla - Follicular phase - Contralateral - Native - Cortex - Follicular phase - Ipsilateral - Decel
## Medulla - Follicular phase - Ipsilateral - Decellularized - Cortex - Follicular phase - Ipsilateral -
## Medulla - Follicular phase - Ipsilateral - Native - Cortex - Follicular phase - Ipsilateral - Decellu
## Medulla - Luteal phase - Contralateral - Decellularized - Cortex - Follicular phase - Ipsilateral - 
## Medulla - Luteal phase - Contralateral - Native - Cortex - Follicular phase - Ipsilateral - Decellul
## Medulla - Luteal phase - Ipsilateral - Decellularized - Cortex - Follicular phase - Ipsilateral - Dec
## Medulla - Luteal phase - Ipsilateral - Native - Cortex - Follicular phase - Ipsilateral - Decellular
## Cortex - Luteal phase - Contralateral - Decellularized - Cortex - Follicular phase - Ipsilateral - Na
## Cortex - Luteal phase - Contralateral - Native - Cortex - Follicular phase - Ipsilateral - Native ==
## Cortex - Luteal phase - Ipsilateral - Decellularized - Cortex - Follicular phase - Ipsilateral - Nat
## Cortex - Luteal phase - Ipsilateral - Native - Cortex - Follicular phase - Ipsilateral - Native == 0
## Medulla - Follicular phase - Contralateral - Decellularized - Cortex - Follicular phase - Ipsilateral
## Medulla - Follicular phase - Contralateral - Native - Cortex - Follicular phase - Ipsilateral - Nati
## Medulla - Follicular phase - Ipsilateral - Decellularized - Cortex - Follicular phase - Ipsilateral -
```

```
## Medulla - Follicular phase - Ipsilateral - Native - Cortex - Follicular phase - Ipsilateral - Native
## Medulla - Luteal phase - Contralateral - Decellularized - Cortex - Follicular phase - Ipsilateral -
## Medulla - Luteal phase - Contralateral - Native - Cortex - Follicular phase - Ipsilateral - Native =
## Medulla - Luteal phase - Ipsilateral - Decellularized - Cortex - Follicular phase - Ipsilateral - Na
## Medulla - Luteal phase - Ipsilateral - Native - Cortex - Follicular phase - Ipsilateral - Native ==
## Cortex - Luteal phase - Contralateral - Native - Cortex - Luteal phase - Contralateral - Decellulari
## Cortex - Luteal phase - Ipsilateral - Decellularized - Cortex - Luteal phase - Contralateral - Decell
## Cortex - Luteal phase - Ipsilateral - Native - Cortex - Luteal phase - Contralateral - Decellularize
## Medulla - Follicular phase - Contralateral - Decellularized - Cortex - Luteal phase - Contralateral -
## Medulla - Follicular phase - Contralateral - Native - Cortex - Luteal phase - Contralateral - Decellu
## Medulla - Follicular phase - Ipsilateral - Decellularized - Cortex - Luteal phase - Contralateral -
## Medulla - Follicular phase - Ipsilateral - Native - Cortex - Luteal phase - Contralateral - Decellula
## Medulla - Luteal phase - Contralateral - Decellularized - Cortex - Luteal phase - Contralateral - De
## Medulla - Luteal phase - Contralateral - Native - Cortex - Luteal phase - Contralateral - Decellulari
## Medulla - Luteal phase - Ipsilateral - Decellularized - Cortex - Luteal phase - Contralateral - Decel
## Medulla - Luteal phase - Ipsilateral - Native - Cortex - Luteal phase - Contralateral - Decellulariz
## Cortex - Luteal phase - Ipsilateral - Decellularized - Cortex - Luteal phase - Contralateral - Nativ
## Cortex - Luteal phase - Ipsilateral - Native - Cortex - Luteal phase - Contralateral - Native == 0
## Medulla - Follicular phase - Contralateral - Decellularized - Cortex - Luteal phase - Contralateral -
## Medulla - Follicular phase - Contralateral - Native - Cortex - Luteal phase - Contralateral - Native
## Medulla - Follicular phase - Ipsilateral - Decellularized - Cortex - Luteal phase - Contralateral -
## Medulla - Follicular phase - Ipsilateral - Native - Cortex - Luteal phase - Contralateral - Native =
## Medulla - Luteal phase - Contralateral - Decellularized - Cortex - Luteal phase - Contralateral - Na
## Medulla - Luteal phase - Contralateral - Native - Cortex - Luteal phase - Contralateral - Native ==
## Medulla - Luteal phase - Ipsilateral - Decellularized - Cortex - Luteal phase - Contralateral - Nati
## Medulla - Luteal phase - Ipsilateral - Native - Cortex - Luteal phase - Contralateral - Native == 0
## Cortex - Luteal phase - Ipsilateral - Native - Cortex - Luteal phase - Ipsilateral - Decellularized
## Medulla - Follicular phase - Contralateral - Decellularized - Cortex - Luteal phase - Ipsilateral -
## Medulla - Follicular phase - Contralateral - Native - Cortex - Luteal phase - Ipsilateral - Decellula
## Medulla - Follicular phase - Ipsilateral - Decellularized - Cortex - Luteal phase - Ipsilateral - De
## Medulla - Follicular phase - Ipsilateral - Native - Cortex - Luteal phase - Ipsilateral - Decellulari
## Medulla - Luteal phase - Contralateral - Decellularized - Cortex - Luteal phase - Ipsilateral - Decel
## Medulla - Luteal phase - Contralateral - Native - Cortex - Luteal phase - Ipsilateral - Decellularize
## Medulla - Luteal phase - Ipsilateral - Decellularized - Cortex - Luteal phase - Ipsilateral - Decellu
## Medulla - Luteal phase - Ipsilateral - Native - Cortex - Luteal phase - Ipsilateral - Decellularized
## Medulla - Follicular phase - Contralateral - Decellularized - Cortex - Luteal phase - Ipsilateral -
## Medulla - Follicular phase - Contralateral - Native - Cortex - Luteal phase - Ipsilateral - Native =
## Medulla - Follicular phase - Ipsilateral - Decellularized - Cortex - Luteal phase - Ipsilateral - Na
## Medulla - Follicular phase - Ipsilateral - Native - Cortex - Luteal phase - Ipsilateral - Native ==
## Medulla - Luteal phase - Contralateral - Decellularized - Cortex - Luteal phase - Ipsilateral - Nati
## Medulla - Luteal phase - Contralateral - Native - Cortex - Luteal phase - Ipsilateral - Native == 0
## Medulla - Luteal phase - Ipsilateral - Decellularized - Cortex - Luteal phase - Ipsilateral - Native
## Medulla - Luteal phase - Ipsilateral - Native - Cortex - Luteal phase - Ipsilateral - Native == 0
## Medulla - Follicular phase - Contralateral - Native - Medulla - Follicular phase - Contralateral - De
## Medulla - Follicular phase - Ipsilateral - Decellularized - Medulla - Follicular phase - Contralatera
## Medulla - Follicular phase - Ipsilateral - Native - Medulla - Follicular phase - Contralateral - Dec
## Medulla - Luteal phase - Contralateral - Decellularized - Medulla - Follicular phase - Contralateral
## Medulla - Luteal phase - Contralateral - Native - Medulla - Follicular phase - Contralateral - Decel
## Medulla - Luteal phase - Ipsilateral - Decellularized - Medulla - Follicular phase - Contralateral -
## Medulla - Luteal phase - Ipsilateral - Native - Medulla - Follicular phase - Contralateral - Decellu
## Medulla - Follicular phase - Ipsilateral - Decellularized - Medulla - Follicular phase - Contralatera
## Medulla - Follicular phase - Ipsilateral - Native - Medulla - Follicular phase - Contralateral - Nati
## Medulla - Luteal phase - Contralateral - Decellularized - Medulla - Follicular phase - Contralateral
## Medulla - Luteal phase - Contralateral - Native - Medulla - Follicular phase - Contralateral - Nativ
```

```
## Medulla - Luteal phase - Ipsilateral - Decellularized - Medulla - Follicular phase - Contralateral -
## Medulla - Luteal phase - Ipsilateral - Native - Medulla - Follicular phase - Contralateral - Native =
## Medulla - Follicular phase - Ipsilateral - Native - Medulla - Follicular phase - Ipsilateral - Decell
## Medulla - Luteal phase - Contralateral - Decellularized - Medulla - Follicular phase - Ipsilateral -
## Medulla - Luteal phase - Contralateral - Native - Medulla - Follicular phase - Ipsilateral - Decellul
## Medulla - Luteal phase - Ipsilateral - Decellularized - Medulla - Follicular phase - Ipsilateral - De
## Medulla - Luteal phase - Ipsilateral - Native - Medulla - Follicular phase - Ipsilateral - Decellula
## Medulla - Luteal phase - Contralateral - Decellularized - Medulla - Follicular phase - Ipsilateral -
## Medulla - Luteal phase - Contralateral - Native - Medulla - Follicular phase - Ipsilateral - Native =
## Medulla - Luteal phase - Ipsilateral - Decellularized - Medulla - Follicular phase - Ipsilateral - Na
## Medulla - Luteal phase - Ipsilateral - Native - Medulla - Follicular phase - Ipsilateral - Native ==
## Medulla - Luteal phase - Contralateral - Native - Medulla - Luteal phase - Contralateral - Decellula
## Medulla - Luteal phase - Ipsilateral - Decellularized - Medulla - Luteal phase - Contralateral - Dec
## Medulla - Luteal phase - Ipsilateral - Native - Medulla - Luteal phase - Contralateral - Decellulari
## Medulla - Luteal phase - Ipsilateral - Decellularized - Medulla - Luteal phase - Contralateral - Nat
## Medulla - Luteal phase - Ipsilateral - Native - Medulla - Luteal phase - Contralateral - Native == 0
## Medulla - Luteal phase - Ipsilateral - Native - Medulla - Luteal phase - Ipsilateral - Decellularize
## ---
## Signif. codes:  0 '***' 0.001 '**' 0.01 '*' 0.05 '.' 0.1 ' ' 1
## (Adjusted p values reported -- single-step method)
```

```
################### YM-Porosity correlation #######################################

# Read the dataset from a CSV file
df <- read.csv("Ovary_YM.csv")

# Update the labels for Position and Estrus_cycle
df$Position <- factor(df$Position, labels = c("Contralateral", "Ipsilateral"))
df$Estrus_cycle <- factor(df$Estrus_cycle, labels = c("Follicular phase", "Luteal phase"))

# Convert Animal_ID to a factor
df$Animal_ID <- as.factor(df$Animal_ID)

# Create a combined factor variable for Estrus_cycle and Cellular
df$Combined <- factor(paste(df$Estrus_cycle, df$Cellular, sep = " "),
                      levels = c("Follicular phase Native", "Follicular phase Decellularized",
                                 "Luteal phase Native", "Luteal phase Decellularized"))

# Create X_position_group based on X_position values
df$X_position_group <- ifelse(df$X_position <= 750, "Medulla", "Cortex")
df$X_position_group <- factor(df$X_position_group, levels = c("Cortex", "Medulla"))

# Cortex

png(filename="Ovary_correlation_Cortex.png",
    width = 10, height = 15, units = "in",
    res = 600)

par(mgp=c(1.5,0.5,0),
    mar = c(4, #bottom
            4, #left
            3, #top
            2),
    cex.main = 0.8,
```

```
    family = "sans" #right
)

# Subset the data for Cortex
df_cortex <- subset(df, X_position_group == "Cortex")

# Create scatter plot for Contra-lateral and Decellularized in the Follicular phase
p_contra_decell_follicular <- ggplot(df_cortex, aes(x = Porosity, y = YM, color = Cellular)) +
  geom_point(data = subset(df_cortex, Position == "Contralateral" & Estrus_cycle == "Follicular phase" &
  geom_smooth(data = subset(df_cortex, Position == "Contralateral" & Estrus_cycle == "Follicular phase"
  labs(x = "Porosity (%)", y = "Young's Modulus (Pa)") +
  ggtitle("Contralateral - Decellularized (Follicular phase)") +
  stat_cor(data = subset(df_cortex, Position == "Contralateral" & Estrus_cycle == "Follicular phase" & C
  theme_bw() +
  theme(panel.grid.major = element_blank(),
        panel.grid.minor = element_blank(),
        legend.position = "none",
        axis.title.x = element_text(size = 16),
        axis.title.y = element_text(size = 16))

# Create scatter plot for Contra-lateral and Native in the Follicular phase
p_contra_native_follicular <- ggplot(df_cortex, aes(x = Porosity, y = YM, color = Cellular)) +
  geom_point(data = subset(df_cortex, Position == "Contralateral" & Estrus_cycle == "Follicular phase" &
  geom_smooth(data = subset(df_cortex, Position == "Contralateral" & Estrus_cycle == "Follicular phase"
  labs(x = "Porosity (%)", y = "Young's Modulus (Pa)") +
  ggtitle("Contralateral - Native (Follicular phase)") +
  stat_cor(data = subset(df_cortex, Position == "Contralateral" & Estrus_cycle == "Follicular phase" & C
  theme_bw() +
  theme(panel.grid.major = element_blank(),
        panel.grid.minor = element_blank(),
        legend.position = "none",
        axis.title.x = element_text(size = 16),
        axis.title.y = element_text(size = 16))

# Create scatter plot for Ipsi-lateral and Decellularized in the Follicular phase
p_ipsi_decell_follicular <- ggplot(df_cortex, aes(x = Porosity, y = YM, color = Cellular)) +
  geom_point(data = subset(df_cortex, Position == "Ipsilateral" & Estrus_cycle == "Follicular phase" & C
  geom_smooth(data = subset(df_cortex, Position == "Ipsilateral" & Estrus_cycle == "Follicular phase" &
  labs(x = "Porosity (%)", y = "Young's Modulus (Pa)") +
  ggtitle("Ipsilateral - Decellularized (Follicular phase)") +
  stat_cor(data = subset(df_cortex, Position == "Ipsilateral" & Estrus_cycle == "Follicular phase" & Cel
  theme_bw() +
  theme(panel.grid.major = element_blank(),
        panel.grid.minor = element_blank(),
        legend.position = "none",
        axis.title.x = element_text(size = 16),
        axis.title.y = element_text(size = 16))

# Create scatter plot for Ipsi-lateral and Native in the Follicular phase
p_ipsi_native_follicular <- ggplot(df_cortex, aes(x = Porosity, y = YM, color = Cellular)) +
  geom_point(data = subset(df_cortex, Position == "Ipsilateral" & Estrus_cycle == "Follicular phase" & C
  geom_smooth(data = subset(df_cortex, Position == "Ipsilateral" & Estrus_cycle == "Follicular phase" &
  labs(x = "Porosity (%)", y = "Young's Modulus (Pa)") +
```

```
  ggtitle("Ipsilateral - Native (Follicular phase)") +
  stat_cor(data = subset(df_cortex, Position == "Ipsilateral" & Estrus_cycle == "Follicular phase" & Cel
  theme_bw() +
  theme(panel.grid.major = element_blank(),
        panel.grid.minor = element_blank(),
        legend.position = "none",
        axis.title.x = element_text(size = 16),
        axis.title.y = element_text(size = 16))

# Create scatter plot for Contra-lateral and Decellularized in the Luteal phase
p_contra_decell_luteal <- ggplot(df_cortex, aes(x = Porosity, y = YM, color = Cellular)) +
  geom_point(data = subset(df_cortex, Position == "Contralateral" & Estrus_cycle == "Luteal phase" & Cel
  geom_smooth(data = subset(df_cortex, Position == "Contralateral" & Estrus_cycle == "Luteal phase" & Ce
  labs(x = "Porosity (%)", y = "Young's Modulus (Pa)") +
  ggtitle("Contralateral - Decellularized (Luteal phase)") +
  stat_cor(data = subset(df_cortex, Position == "Contralateral" & Estrus_cycle == "Luteal phase" & Cellu
  theme_bw() +
  theme(panel.grid.major = element_blank(),
        panel.grid.minor = element_blank(),
        legend.position = "none",
        axis.title.x = element_text(size = 16),
        axis.title.y = element_text(size = 16))

# Create scatter plot for Contra-lateral and Native in the Luteal phase
p_contra_native_luteal <- ggplot(df_cortex, aes(x = Porosity, y = YM, color = Cellular)) +
  geom_point(data = subset(df_cortex, Position == "Contralateral" & Estrus_cycle == "Luteal phase" & Cel
  geom_smooth(data = subset(df_cortex, Position == "Contralateral" & Estrus_cycle == "Luteal phase" & Ce
  labs(x = "Porosity (%)", y = "Young's Modulus (Pa)") +
  ggtitle("Contralateral - Native (Luteal phase)") +
  stat_cor(data = subset(df_cortex, Position == "Contralateral" & Estrus_cycle == "Luteal phase" & Cellu
  theme_bw() +
  theme(panel.grid.major = element_blank(),
        panel.grid.minor = element_blank(),
        legend.position = "none",
        axis.title.x = element_text(size = 16),
        axis.title.y = element_text(size = 16))

# Create scatter plot for Ipsi-lateral and Decellularized in the Luteal phase
p_ipsi_decell_luteal <- ggplot(df_cortex, aes(x = Porosity, y = YM, color = Cellular)) +
  geom_point(data = subset(df_cortex, Position == "Ipsilateral" & Estrus_cycle == "Luteal phase" & Cellu
  geom_smooth(data = subset(df_cortex, Position == "Ipsilateral" & Estrus_cycle == "Luteal phase" & Cel
  labs(x = "Porosity (%)", y = "Young's Modulus (Pa)") +
  ggtitle("Ipsilateral - Decellularized (Luteal phase)") +
  stat_cor(data = subset(df_cortex, Position == "Ipsilateral" & Estrus_cycle == "Luteal phase" & Cellula
  theme_bw() +
  theme(panel.grid.major = element_blank(),
        panel.grid.minor = element_blank(),
        legend.position = "none",
        axis.title.x = element_text(size = 16),
        axis.title.y = element_text(size = 16))

# Create scatter plot for Ipsi-lateral and Native in the Luteal phase
p_ipsi_native_luteal <- ggplot(df_cortex, aes(x = Porosity, y = YM, color = Cellular)) +
```

```r
  geom_point(data = subset(df_cortex, Position == "Ipsilateral" & Estrus_cycle == "Luteal phase" & Cellu
  geom_smooth(data = subset(df_cortex, Position == "Ipsilateral" & Estrus_cycle == "Luteal phase" & Cell
  labs(x = "Porosity (%)", y = "Young's Modulus (Pa)") +
  ggtitle("Ipsilateral - Native (Luteal phase)") +
  stat_cor(data = subset(df_cortex, Position == "Ipsilateral" & Estrus_cycle == "Luteal phase" & Cellula
  theme_bw() +
  theme(panel.grid.major = element_blank(),
        panel.grid.minor = element_blank(),
        legend.position = "none",
        axis.title.x = element_text(size = 16),
        axis.title.y = element_text(size = 16))

# Combine the plots using cowplot
p_combined <- cowplot::plot_grid(
  cowplot::plot_grid(p_contra_decell_follicular, p_contra_native_follicular, nrow = 1),
  cowplot::plot_grid(p_ipsi_decell_follicular, p_ipsi_native_follicular, nrow = 1),
  cowplot::plot_grid(p_contra_decell_luteal, p_contra_native_luteal, nrow = 1),
  cowplot::plot_grid(p_ipsi_decell_luteal, p_ipsi_native_luteal, nrow = 1),
  nrow = 4
)
```

```
## `geom_smooth()` using formula = 'y ~ x'
## `geom_smooth()` using formula = 'y ~ x'
## `geom_smooth()` using formula = 'y ~ x'
## `geom_smooth()` using formula = 'y ~ x'
## `geom_smooth()` using formula = 'y ~ x'
## `geom_smooth()` using formula = 'y ~ x'
## `geom_smooth()` using formula = 'y ~ x'
## `geom_smooth()` using formula = 'y ~ x'
```

```r
# Display the combined plot
p_combined


dev.off()
```

```
## pdf
##   2
```

```r
# Medulla

png(filename="Ovary_correlation_Medulla.png",
    width = 10, height = 15, units = "in",
    res = 600)

par(mgp=c(1.5,0.5,0),
    mar = c(4, #bottom
            4, #left
            3, #top
            2),
    cex.main = 0.8,
    family = "sans" #right
)

# Subset the data for Medulla
```

```
df_medulla <- subset(df, X_position_group == "Medulla")

# Create scatter plot for Contra-lateral and Decellularized in the Follicular phase
p_contra_decell_follicular <- ggplot(df_medulla, aes(x = Porosity, y = YM, color = Cellular)) +
  geom_point(data = subset(df_medulla, Position == "Contralateral" & Estrus_cycle == "Follicular phase"
  geom_smooth(data = subset(df_medulla, Position == "Contralateral" & Estrus_cycle == "Follicular phase
  labs(x = "Porosity (%)", y = "Young's Modulus (Pa)") +
  ggtitle("Contralateral - Decellularized (Follicular phase)") +
  stat_cor(data = subset(df_medulla, Position == "Contralateral" & Estrus_cycle == "Follicular phase" &
  theme_bw() +
  theme(panel.grid.major = element_blank(),
        panel.grid.minor = element_blank(),
        legend.position = "none",
        axis.title.x = element_text(size = 16),
        axis.title.y = element_text(size = 16))

# Create scatter plot for Contra-lateral and Native in the Follicular phase
p_contra_native_follicular <- ggplot(df_medulla, aes(x = Porosity, y = YM, color = Cellular)) +
  geom_point(data = subset(df_medulla, Position == "Contralateral" & Estrus_cycle == "Follicular phase"
  geom_smooth(data = subset(df_medulla, Position == "Contralateral" & Estrus_cycle == "Follicular phase
  labs(x = "Porosity (%)", y = "Young's Modulus (Pa)") +
  ggtitle("Contralateral - Native (Follicular phase)") +
  stat_cor(data = subset(df_medulla, Position == "Contralateral" & Estrus_cycle == "Follicular phase" &
  theme_bw() +
  theme(panel.grid.major = element_blank(),
        panel.grid.minor = element_blank(),
        legend.position = "none",
        axis.title.x = element_text(size = 16),
        axis.title.y = element_text(size = 16))

# Create scatter plot for Ipsi-lateral and Decellularized in the Follicular phase
p_ipsi_decell_follicular <- ggplot(df_medulla, aes(x = Porosity, y = YM, color = Cellular)) +
  geom_point(data = subset(df_medulla, Position == "Ipsilateral" & Estrus_cycle == "Follicular phase" &
  geom_smooth(data = subset(df_medulla, Position == "Ipsilateral" & Estrus_cycle == "Follicular phase" &
  labs(x = "Porosity (%)", y = "Young's Modulus (Pa)") +
  ggtitle("Ipsilateral - Decellularized (Follicular phase)") +
  stat_cor(data = subset(df_medulla, Position == "Ipsilateral" & Estrus_cycle == "Follicular phase" & C
  theme_bw() +
  theme(panel.grid.major = element_blank(),
        panel.grid.minor = element_blank(),
        legend.position = "none",
        axis.title.x = element_text(size = 16),
        axis.title.y = element_text(size = 16))

# Create scatter plot for Ipsi-lateral and Native in the Follicular phase
p_ipsi_native_follicular <- ggplot(df_medulla, aes(x = Porosity, y = YM, color = Cellular)) +
  geom_point(data = subset(df_medulla, Position == "Ipsilateral" & Estrus_cycle == "Follicular phase" &
  geom_smooth(data = subset(df_medulla, Position == "Ipsilateral" & Estrus_cycle == "Follicular phase" &
  labs(x = "Porosity (%)", y = "Young's Modulus (Pa)") +
  ggtitle("Ipsilateral - Native (Follicular phase)") +
  stat_cor(data = subset(df_medulla, Position == "Ipsilateral" & Estrus_cycle == "Follicular phase" & C
  theme_bw() +
  theme(panel.grid.major = element_blank(),
```

```
        panel.grid.minor = element_blank(),
        legend.position = "none",
        axis.title.x = element_text(size = 16),
        axis.title.y = element_text(size = 16))

# Create scatter plot for Contra-lateral and Decellularized in the Luteal phase
p_contra_decell_luteal <- ggplot(df_medulla, aes(x = Porosity, y = YM, color = Cellular)) +
  geom_point(data = subset(df_medulla, Position == "Contralateral" & Estrus_cycle == "Luteal phase" & Co
  geom_smooth(data = subset(df_medulla, Position == "Contralateral" & Estrus_cycle == "Luteal phase" & C
  labs(x = "Porosity (%)", y = "Young's Modulus (Pa)") +
  ggtitle("Contralateral - Decellularized (Luteal phase)") +
  stat_cor(data = subset(df_medulla, Position == "Contralateral" & Estrus_cycle == "Luteal phase" & Cel
  theme_bw() +
  theme(panel.grid.major = element_blank(),
        panel.grid.minor = element_blank(),
        legend.position = "none",
        axis.title.x = element_text(size = 16),
        axis.title.y = element_text(size = 16))

# Create scatter plot for Contra-lateral and Native in the Luteal phase
p_contra_native_luteal <- ggplot(df_medulla, aes(x = Porosity, y = YM, color = Cellular)) +
  geom_point(data = subset(df_medulla, Position == "Contralateral" & Estrus_cycle == "Luteal phase" & Co
  geom_smooth(data = subset(df_medulla, Position == "Contralateral" & Estrus_cycle == "Luteal phase" & C
  labs(x = "Porosity (%)", y = "Young's Modulus (Pa)") +
  ggtitle("Contralateral - Native (Luteal phase)") +
  stat_cor(data = subset(df_medulla, Position == "Contralateral" & Estrus_cycle == "Luteal phase" & Cel
  theme_bw() +
  theme(panel.grid.major = element_blank(),
        panel.grid.minor = element_blank(),
        legend.position = "none",
        axis.title.x = element_text(size = 16),
        axis.title.y = element_text(size = 16))

# Create scatter plot for Ipsi-lateral and Decellularized in the Luteal phase
p_ipsi_decell_luteal <- ggplot(df_medulla, aes(x = Porosity, y = YM, color = Cellular)) +
  geom_point(data = subset(df_medulla, Position == "Ipsilateral" & Estrus_cycle == "Luteal phase" & Cel
  geom_smooth(data = subset(df_medulla, Position == "Ipsilateral" & Estrus_cycle == "Luteal phase" & Ce
  labs(x = "Porosity (%)", y = "Young's Modulus (Pa)") +
  ggtitle("Ipsilateral - Decellularized (Luteal phase)") +
  stat_cor(data = subset(df_medulla, Position == "Ipsilateral" & Estrus_cycle == "Luteal phase" & Cellu
  theme_bw() +
  theme(panel.grid.major = element_blank(),
        panel.grid.minor = element_blank(),
        legend.position = "none",
        axis.title.x = element_text(size = 16),
        axis.title.y = element_text(size = 16))

# Create scatter plot for Ipsi-lateral and Native in the Luteal phase
p_ipsi_native_luteal <- ggplot(df_medulla, aes(x = Porosity, y = YM, color = Cellular)) +
  geom_point(data = subset(df_medulla, Position == "Ipsilateral" & Estrus_cycle == "Luteal phase" & Cel
  geom_smooth(data = subset(df_medulla, Position == "Ipsilateral" & Estrus_cycle == "Luteal phase" & Ce
  labs(x = "Porosity (%)", y = "Young's Modulus (Pa)") +
  ggtitle("Ipsilateral - Native (Luteal phase)") +
```

```r
  stat_cor(data = subset(df_medulla, Position == "Ipsilateral" & Estrus_cycle == "Luteal phase" & Cellul
  theme_bw() +
  theme(panel.grid.major = element_blank(),
        panel.grid.minor = element_blank(),
        legend.position = "none",
        axis.title.x = element_text(size = 16),
        axis.title.y = element_text(size = 16))

# Combine the plots using cowplot
p_combined <- cowplot::plot_grid(
  cowplot::plot_grid(p_contra_decell_follicular, p_contra_native_follicular, nrow = 1),
  cowplot::plot_grid(p_ipsi_decell_follicular, p_ipsi_native_follicular, nrow = 1),
  cowplot::plot_grid(p_contra_decell_luteal, p_contra_native_luteal, nrow = 1),
  cowplot::plot_grid(p_ipsi_decell_luteal, p_ipsi_native_luteal, nrow = 1),
  nrow = 4
)
```

```
## `geom_smooth()` using formula = 'y ~ x'
## `geom_smooth()` using formula = 'y ~ x'
## `geom_smooth()` using formula = 'y ~ x'
## `geom_smooth()` using formula = 'y ~ x'
## `geom_smooth()` using formula = 'y ~ x'
## `geom_smooth()` using formula = 'y ~ x'
## `geom_smooth()` using formula = 'y ~ x'
## `geom_smooth()` using formula = 'y ~ x'
```

```r
# Display the combined plot
p_combined

dev.off()
```

```
## pdf
##   2
```

```r
#################################################################
#################### Oviduct Analysis ##############################
#################################################################

################### E' E" #######################################

#data input
data <- read.csv("Oviduct_rheo.csv")

# Subset the data based on X_position_group
data_ampulla <- subset(data, Tissue == "Ampulla")
data_isthmus <- subset(data, Tissue == "Isthmus")

# Plot Ampulla

png(filename="E1_E2_ampulla.png",
    width = 9.86, height = 6.86, units = "in",
    res = 600)

par(mgp=c(1.5,0.5,0),
    mar = c(4, #bottom
```

```r
            4, #left
            3, #top
            2),
    cex.main = 0.8,
    family = "sans" #right
)

# Calculate means and standard deviations
data_summary <- data_ampulla %>%
  group_by(Module, Estrus_cycle, Position, Hz) %>%
  summarise(mean_Indentation = mean(Indentation, na.rm = TRUE),
            sd_Indentation = sd(Indentation, na.rm = TRUE))
```

```
## `summarise()` has grouped output by 'Module', 'Estrus_cycle', 'Position'. You
## can override using the `.groups` argument.
```

```r
# Create a new variable for line type
data_summary <- data_summary %>%
  mutate(LineType = ifelse(Module == "E1", "E1", "E2"))

# Define fill colors for each factor level
fill_colors <- c("Follicular phase Contralateral" = viridis_pal(option = "D")(1),
                 "Follicular phase Ipsilateral" = alpha(viridis_pal(option = "D")(1), 0.2),
                 "Luteal phase Contralateral" = "#238A8DFF",
                 "Luteal phase Ipsilateral" = alpha("#238A8DFF", 0.2))

# Create the line plot with separate lines for each Estrus_cycle and Position
ggplot(data_summary, aes(x = Hz, y = mean_Indentation, group = interaction(Module, Estrus_cycle, Positi
                         color = paste(Estrus_cycle, Position), linetype = LineType)) +
  geom_line(size = 1.5) +  # Adjust line thickness
  geom_point(size = 1) +  # Add dots at each Hz point
  scale_color_manual(values = fill_colors) +  # Use custom fill colors
  scale_linetype_manual(values = c("E1" = "solid", "E2" = "dashed"),
                        labels = c("E1" = "E'", "E2" = "E\"")) +
  scale_x_continuous(breaks = c(1, 5, 10, 20)) +
  scale_y_continuous(labels = scientific_format(), limits = c(0, 16000)) +  # Adjust Y-axis limits
  labs(y = expression(paste("E'/E'' (Pa)")), x = "Frequency (Hz)", color = "Estrus Cycle + Position") +
  theme_minimal() +
  theme(panel.grid.major = element_blank(),
        panel.grid.minor = element_blank(),
        panel.border = element_rect(colour = "black", fill = NA, size = 1),
        axis.text = element_text(size = 12))  # Increase font size of axis labels

dev.off()
```

```
## pdf
##   2
```

```r
# Plot Isthmus

png(filename="E1_E2_isthmus.png",
    width = 9.86, height = 6.86, units = "in",
    res = 600)

par(mgp=c(1.5,0.5,0),
```

```r
    mar = c(4, #bottom
            4, #left
            3, #top
            2),
    cex.main = 0.8,
    family = "sans" #right
)

# Calculate means and standard deviations
data_summary <- data_isthmus %>%
  group_by(Module, Estrus_cycle, Position, Hz) %>%
  summarise(mean_Indentation = mean(Indentation, na.rm = TRUE),
            sd_Indentation = sd(Indentation, na.rm = TRUE))
```

```
## `summarise()` has grouped output by 'Module', 'Estrus_cycle', 'Position'. You
## can override using the `.groups` argument.
```

```r
# Create a new variable for line type
data_summary <- data_summary %>%
  mutate(LineType = ifelse(Module == "E1", "E1", "E2"))

# Define fill colors for each factor level
fill_colors <- c("Follicular phase Contralateral" = viridis_pal(option = "D")(1),
                 "Follicular phase Ipsilateral" = alpha(viridis_pal(option = "D")(1), 0.2),
                 "Luteal phase Contralateral" = "#238A8DFF",
                 "Luteal phase Ipsilateral" = alpha("#238A8DFF", 0.2))

# Create the line plot with separate lines for each Estrus_cycle and Position
ggplot(data_summary, aes(x = Hz, y = mean_Indentation, group = interaction(Module, Estrus_cycle, Positi
                         color = paste(Estrus_cycle, Position), linetype = LineType)) +
  geom_line(size = 1.5) +  # Adjust line thickness
  geom_point(size = 1) +  # Add dots at each Hz point
  scale_color_manual(values = fill_colors) +  # Use custom fill colors
  scale_linetype_manual(values = c("E1" = "solid", "E2" = "dashed"),
                        labels = c("E1" = "E'", "E2" = "E\"")) +
  scale_x_continuous(breaks = c(1, 5, 10, 20)) +
  scale_y_continuous(labels = scientific_format(), limits = c(0, 20000)) +  # Adjust Y-axis limits
  labs(y = expression(paste("E'/E'' (Pa)")), x = "Frequency (Hz)", color = "Estrus Cycle + Position") +
  theme_minimal() +
  theme(panel.grid.major = element_blank(),
        panel.grid.minor = element_blank(),
        panel.border = element_rect(colour = "black", fill = NA, size = 1),
        axis.text = element_text(size = 12))  # Increase font size of axis labels

dev.off()
```

```
## pdf
##   2
```

```r
# Stats

# Ampulla
df <- data_ampulla

# Create a factor variable for the comparisons
```

```r
df$Comparison <- factor(paste(df$Estrus_cycle, df$Position, df$Module, sep = " - "))

# Perform pairwise comparisons using Tukey's method
model <- lm(Indentation ~ Comparison, data = df)
comp <- glht(model, linfct = mcp(Comparison = "Tukey"))

# Summarize the results
summary(comp)
```

```
##
##   Simultaneous Tests for General Linear Hypotheses
##
## Multiple Comparisons of Means: Tukey Contrasts
##
##
## Fit: lm(formula = Indentation ~ Comparison, data = df)
##
## Linear Hypotheses:
##                                                                          Estimate
## Follicular phase - Contralateral - E2 - Follicular phase - Contralateral - E1 == 0  -5779.5
## Follicular phase - Ipsilateral - E1 - Follicular phase - Contralateral - E1 == 0      597.0
## Follicular phase - Ipsilateral - E2 - Follicular phase - Contralateral - E1 == 0    -5576.1
## Luteal phase - Contralateral - E1 - Follicular phase - Contralateral - E1 == 0      -1913.3
## Luteal phase - Contralateral - E2 - Follicular phase - Contralateral - E1 == 0      -3557.2
## Luteal phase - Ipsilateral - E1 - Follicular phase - Contralateral - E1 == 0         4968.9
## Luteal phase - Ipsilateral - E2 - Follicular phase - Contralateral - E1 == 0        -2017.0
## Follicular phase - Ipsilateral - E1 - Follicular phase - Contralateral - E2 == 0     6376.5
## Follicular phase - Ipsilateral - E2 - Follicular phase - Contralateral - E2 == 0      203.4
## Luteal phase - Contralateral - E1 - Follicular phase - Contralateral - E2 == 0       3866.2
## Luteal phase - Contralateral - E2 - Follicular phase - Contralateral - E2 == 0       2222.3
## Luteal phase - Ipsilateral - E1 - Follicular phase - Contralateral - E2 == 0        10748.4
## Luteal phase - Ipsilateral - E2 - Follicular phase - Contralateral - E2 == 0         3762.5
## Follicular phase - Ipsilateral - E2 - Follicular phase - Ipsilateral - E1 == 0      -6173.1
## Luteal phase - Contralateral - E1 - Follicular phase - Ipsilateral - E1 == 0        -2510.3
## Luteal phase - Contralateral - E2 - Follicular phase - Ipsilateral - E1 == 0        -4154.2
## Luteal phase - Ipsilateral - E1 - Follicular phase - Ipsilateral - E1 == 0           4371.9
## Luteal phase - Ipsilateral - E2 - Follicular phase - Ipsilateral - E1 == 0          -2614.0
## Luteal phase - Contralateral - E1 - Follicular phase - Ipsilateral - E2 == 0         3662.8
## Luteal phase - Contralateral - E2 - Follicular phase - Ipsilateral - E2 == 0         2018.9
## Luteal phase - Ipsilateral - E1 - Follicular phase - Ipsilateral - E2 == 0          10545.0
## Luteal phase - Ipsilateral - E2 - Follicular phase - Ipsilateral - E2 == 0           3559.1
## Luteal phase - Contralateral - E2 - Luteal phase - Contralateral - E1 == 0          -1643.9
## Luteal phase - Ipsilateral - E1 - Luteal phase - Contralateral - E1 == 0             6882.2
## Luteal phase - Ipsilateral - E2 - Luteal phase - Contralateral - E1 == 0             -103.7
## Luteal phase - Ipsilateral - E1 - Luteal phase - Contralateral - E2 == 0             8526.1
## Luteal phase - Ipsilateral - E2 - Luteal phase - Contralateral - E2 == 0             1540.2
## Luteal phase - Ipsilateral - E2 - Luteal phase - Ipsilateral - E1 == 0              -6985.9
##                                                                          Std. Error
## Follicular phase - Contralateral - E2 - Follicular phase - Contralateral - E1 == 0     1588.0
## Follicular phase - Ipsilateral - E1 - Follicular phase - Contralateral - E1 == 0       1692.8
## Follicular phase - Ipsilateral - E2 - Follicular phase - Contralateral - E1 == 0       1692.8
## Luteal phase - Contralateral - E1 - Follicular phase - Contralateral - E1 == 0         1497.2
## Luteal phase - Contralateral - E2 - Follicular phase - Contralateral - E1 == 0         1497.2
## Luteal phase - Ipsilateral - E1 - Follicular phase - Contralateral - E1 == 0           1464.0
```

```
## Luteal phase - Ipsilateral - E2 - Follicular phase - Contralateral - E1 == 0          1464.0
## Follicular phase - Ipsilateral - E1 - Follicular phase - Contralateral - E2 == 0      1692.8
## Follicular phase - Ipsilateral - E2 - Follicular phase - Contralateral - E2 == 0      1692.8
## Luteal phase - Contralateral - E1 - Follicular phase - Contralateral - E2 == 0        1497.2
## Luteal phase - Contralateral - E2 - Follicular phase - Contralateral - E2 == 0        1497.2
## Luteal phase - Ipsilateral - E1 - Follicular phase - Contralateral - E2 == 0          1464.0
## Luteal phase - Ipsilateral - E2 - Follicular phase - Contralateral - E2 == 0          1464.0
## Follicular phase - Ipsilateral - E2 - Follicular phase - Ipsilateral - E1 == 0        1791.5
## Luteal phase - Contralateral - E1 - Follicular phase - Ipsilateral - E1 == 0          1607.9
## Luteal phase - Contralateral - E2 - Follicular phase - Ipsilateral - E1 == 0          1607.9
## Luteal phase - Ipsilateral - E1 - Follicular phase - Ipsilateral - E1 == 0            1577.1
## Luteal phase - Ipsilateral - E2 - Follicular phase - Ipsilateral - E1 == 0            1577.1
## Luteal phase - Contralateral - E1 - Follicular phase - Ipsilateral - E2 == 0          1607.9
## Luteal phase - Contralateral - E2 - Follicular phase - Ipsilateral - E2 == 0          1607.9
## Luteal phase - Ipsilateral - E1 - Follicular phase - Ipsilateral - E2 == 0            1577.1
## Luteal phase - Ipsilateral - E2 - Follicular phase - Ipsilateral - E2 == 0            1577.1
## Luteal phase - Contralateral - E2 - Luteal phase - Contralateral - E1 == 0            1400.5
## Luteal phase - Ipsilateral - E1 - Luteal phase - Contralateral - E1 == 0              1365.0
## Luteal phase - Ipsilateral - E2 - Luteal phase - Contralateral - E1 == 0              1365.0
## Luteal phase - Ipsilateral - E1 - Luteal phase - Contralateral - E2 == 0              1365.0
## Luteal phase - Ipsilateral - E2 - Luteal phase - Contralateral - E2 == 0              1365.0
## Luteal phase - Ipsilateral - E2 - Luteal phase - Ipsilateral - E1 == 0                1328.6
##                                                                                    t value
## Follicular phase - Contralateral - E2 - Follicular phase - Contralateral - E1 == 0  -3.640
## Follicular phase - Ipsilateral - E1 - Follicular phase - Contralateral - E1 == 0     0.353
## Follicular phase - Ipsilateral - E2 - Follicular phase - Contralateral - E1 == 0    -3.294
## Luteal phase - Contralateral - E1 - Follicular phase - Contralateral - E1 == 0      -1.278
## Luteal phase - Contralateral - E2 - Follicular phase - Contralateral - E1 == 0      -2.376
## Luteal phase - Ipsilateral - E1 - Follicular phase - Contralateral - E1 == 0         3.394
## Luteal phase - Ipsilateral - E2 - Follicular phase - Contralateral - E1 == 0        -1.378
## Follicular phase - Ipsilateral - E1 - Follicular phase - Contralateral - E2 == 0     3.767
## Follicular phase - Ipsilateral - E2 - Follicular phase - Contralateral - E2 == 0     0.120
## Luteal phase - Contralateral - E1 - Follicular phase - Contralateral - E2 == 0       2.582
## Luteal phase - Contralateral - E2 - Follicular phase - Contralateral - E2 == 0       1.484
## Luteal phase - Ipsilateral - E1 - Follicular phase - Contralateral - E2 == 0         7.342
## Luteal phase - Ipsilateral - E2 - Follicular phase - Contralateral - E2 == 0         2.570
## Follicular phase - Ipsilateral - E2 - Follicular phase - Ipsilateral - E1 == 0      -3.446
## Luteal phase - Contralateral - E1 - Follicular phase - Ipsilateral - E1 == 0        -1.561
## Luteal phase - Contralateral - E2 - Follicular phase - Ipsilateral - E1 == 0        -2.584
## Luteal phase - Ipsilateral - E1 - Follicular phase - Ipsilateral - E1 == 0           2.772
## Luteal phase - Ipsilateral - E2 - Follicular phase - Ipsilateral - E1 == 0          -1.657
## Luteal phase - Contralateral - E1 - Follicular phase - Ipsilateral - E2 == 0         2.278
## Luteal phase - Contralateral - E2 - Follicular phase - Ipsilateral - E2 == 0         1.256
## Luteal phase - Ipsilateral - E1 - Follicular phase - Ipsilateral - E2 == 0           6.686
## Luteal phase - Ipsilateral - E2 - Follicular phase - Ipsilateral - E2 == 0           2.257
## Luteal phase - Contralateral - E2 - Luteal phase - Contralateral - E1 == 0          -1.174
## Luteal phase - Ipsilateral - E1 - Luteal phase - Contralateral - E1 == 0             5.042
## Luteal phase - Ipsilateral - E2 - Luteal phase - Contralateral - E1 == 0            -0.076
## Luteal phase - Ipsilateral - E1 - Luteal phase - Contralateral - E2 == 0             6.246
## Luteal phase - Ipsilateral - E2 - Luteal phase - Contralateral - E2 == 0             1.128
## Luteal phase - Ipsilateral - E2 - Luteal phase - Ipsilateral - E1 == 0              -5.258
##                                                                                   Pr(>|t|)
## Follicular phase - Contralateral - E2 - Follicular phase - Contralateral - E1 == 0  0.00703
## Follicular phase - Ipsilateral - E1 - Follicular phase - Contralateral - E1 == 0    0.99997
```

```
## Follicular phase - Ipsilateral - E2 - Follicular phase - Contralateral - E1 == 0    0.02287
## Luteal phase - Contralateral - E1 - Follicular phase - Contralateral - E1 == 0      0.90555
## Luteal phase - Contralateral - E2 - Follicular phase - Contralateral - E1 == 0      0.25317
## Luteal phase - Ipsilateral - E1 - Follicular phase - Contralateral - E1 == 0        0.01652
## Luteal phase - Ipsilateral - E2 - Follicular phase - Contralateral - E1 == 0        0.86556
## Follicular phase - Ipsilateral - E1 - Follicular phase - Contralateral - E2 == 0    0.00472
## Follicular phase - Ipsilateral - E2 - Follicular phase - Contralateral - E2 == 0    1.00000
## Luteal phase - Contralateral - E1 - Follicular phase - Contralateral - E2 == 0      0.16273
## Luteal phase - Contralateral - E2 - Follicular phase - Contralateral - E2 == 0      0.81368
## Luteal phase - Ipsilateral - E1 - Follicular phase - Contralateral - E2 == 0        < 0.001
## Luteal phase - Ipsilateral - E2 - Follicular phase - Contralateral - E2 == 0        0.16769
## Follicular phase - Ipsilateral - E2 - Follicular phase - Ipsilateral - E1 == 0      0.01393
## Luteal phase - Contralateral - E1 - Follicular phase - Ipsilateral - E1 == 0        0.77093
## Luteal phase - Contralateral - E2 - Follicular phase - Ipsilateral - E1 == 0        0.16230
## Luteal phase - Ipsilateral - E1 - Follicular phase - Ipsilateral - E1 == 0          0.10310
## Luteal phase - Ipsilateral - E2 - Follicular phase - Ipsilateral - E1 == 0          0.71211
## Luteal phase - Contralateral - E1 - Follicular phase - Ipsilateral - E2 == 0        0.30529
## Luteal phase - Contralateral - E2 - Follicular phase - Ipsilateral - E2 == 0        0.91341
## Luteal phase - Ipsilateral - E1 - Follicular phase - Ipsilateral - E2 == 0          < 0.001
## Luteal phase - Ipsilateral - E2 - Follicular phase - Ipsilateral - E2 == 0          0.31730
## Luteal phase - Contralateral - E2 - Luteal phase - Contralateral - E1 == 0          0.93828
## Luteal phase - Ipsilateral - E1 - Luteal phase - Contralateral - E1 == 0            < 0.001
## Luteal phase - Ipsilateral - E2 - Luteal phase - Contralateral - E1 == 0            1.00000
## Luteal phase - Ipsilateral - E1 - Luteal phase - Contralateral - E2 == 0            < 0.001
## Luteal phase - Ipsilateral - E2 - Luteal phase - Contralateral - E2 == 0            0.94971
## Luteal phase - Ipsilateral - E2 - Luteal phase - Ipsilateral - E1 == 0              < 0.001
##
## Follicular phase - Contralateral - E2 - Follicular phase - Contralateral - E1 == 0 **
## Follicular phase - Ipsilateral - E1 - Follicular phase - Contralateral - E1 == 0
## Follicular phase - Ipsilateral - E2 - Follicular phase - Contralateral - E1 == 0   *
## Luteal phase - Contralateral - E1 - Follicular phase - Contralateral - E1 == 0
## Luteal phase - Contralateral - E2 - Follicular phase - Contralateral - E1 == 0
## Luteal phase - Ipsilateral - E1 - Follicular phase - Contralateral - E1 == 0       *
## Luteal phase - Ipsilateral - E2 - Follicular phase - Contralateral - E1 == 0
## Follicular phase - Ipsilateral - E1 - Follicular phase - Contralateral - E2 == 0   **
## Follicular phase - Ipsilateral - E2 - Follicular phase - Contralateral - E2 == 0
## Luteal phase - Contralateral - E1 - Follicular phase - Contralateral - E2 == 0
## Luteal phase - Contralateral - E2 - Follicular phase - Contralateral - E2 == 0
## Luteal phase - Ipsilateral - E1 - Follicular phase - Contralateral - E2 == 0       ***
## Luteal phase - Ipsilateral - E2 - Follicular phase - Contralateral - E2 == 0
## Follicular phase - Ipsilateral - E2 - Follicular phase - Ipsilateral - E1 == 0     *
## Luteal phase - Contralateral - E1 - Follicular phase - Ipsilateral - E1 == 0
## Luteal phase - Contralateral - E2 - Follicular phase - Ipsilateral - E1 == 0
## Luteal phase - Ipsilateral - E1 - Follicular phase - Ipsilateral - E1 == 0
## Luteal phase - Ipsilateral - E2 - Follicular phase - Ipsilateral - E1 == 0
## Luteal phase - Contralateral - E1 - Follicular phase - Ipsilateral - E2 == 0
## Luteal phase - Contralateral - E2 - Follicular phase - Ipsilateral - E2 == 0
## Luteal phase - Ipsilateral - E1 - Follicular phase - Ipsilateral - E2 == 0         ***
## Luteal phase - Ipsilateral - E2 - Follicular phase - Ipsilateral - E2 == 0
## Luteal phase - Contralateral - E2 - Luteal phase - Contralateral - E1 == 0
## Luteal phase - Ipsilateral - E1 - Luteal phase - Contralateral - E1 == 0           ***
## Luteal phase - Ipsilateral - E2 - Luteal phase - Contralateral - E1 == 0
## Luteal phase - Ipsilateral - E1 - Luteal phase - Contralateral - E2 == 0           ***
## Luteal phase - Ipsilateral - E2 - Luteal phase - Contralateral - E2 == 0
```

```
## Luteal phase - Ipsilateral - E2 - Luteal phase - Ipsilateral - E1 == 0             ***
## ---
## Signif. codes:  0 '***' 0.001 '**' 0.01 '*' 0.05 '.' 0.1 ' ' 1
## (Adjusted p values reported -- single-step method)
```

```r
# Isthmus

df <- data_isthmus

# Create a factor variable for the comparisons
df$Comparison <- factor(paste(df$Estrus_cycle, df$Position, df$Module, sep = " - "))

# Perform pairwise comparisons using Tukey's method
model <- lm(Indentation ~ Comparison, data = df)
comp <- glht(model, linfct = mcp(Comparison = "Tukey"))

# Summarize the results
summary(comp)
```

```
##
##   Simultaneous Tests for General Linear Hypotheses
##
## Multiple Comparisons of Means: Tukey Contrasts
##
##
## Fit: lm(formula = Indentation ~ Comparison, data = df)
##
## Linear Hypotheses:
##                                                                            Estimate
## Follicular phase - Contralateral - E2 - Follicular phase - Contralateral - E1 == 0 -8122.789
## Follicular phase - Ipsilateral - E1 - Follicular phase - Contralateral - E1 == 0    784.049
## Follicular phase - Ipsilateral - E2 - Follicular phase - Contralateral - E1 == 0  -7529.365
## Luteal phase - Contralateral - E1 - Follicular phase - Contralateral - E1 == 0     2228.749
## Luteal phase - Contralateral - E2 - Follicular phase - Contralateral - E1 == 0    -6088.611
## Luteal phase - Ipsilateral - E1 - Follicular phase - Contralateral - E1 == 0       2231.245
## Luteal phase - Ipsilateral - E2 - Follicular phase - Contralateral - E1 == 0      -6626.611
## Follicular phase - Ipsilateral - E1 - Follicular phase - Contralateral - E2 == 0   8906.838
## Follicular phase - Ipsilateral - E2 - Follicular phase - Contralateral - E2 == 0    593.425
## Luteal phase - Contralateral - E1 - Follicular phase - Contralateral - E2 == 0    10351.538
## Luteal phase - Contralateral - E2 - Follicular phase - Contralateral - E2 == 0     2034.178
## Luteal phase - Ipsilateral - E1 - Follicular phase - Contralateral - E2 == 0      10354.034
## Luteal phase - Ipsilateral - E2 - Follicular phase - Contralateral - E2 == 0       1496.178
## Follicular phase - Ipsilateral - E2 - Follicular phase - Ipsilateral - E1 == 0    -8313.413
## Luteal phase - Contralateral - E1 - Follicular phase - Ipsilateral - E1 == 0       1444.700
## Luteal phase - Contralateral - E2 - Follicular phase - Ipsilateral - E1 == 0      -6872.660
## Luteal phase - Ipsilateral - E1 - Follicular phase - Ipsilateral - E1 == 0         1447.196
## Luteal phase - Ipsilateral - E2 - Follicular phase - Ipsilateral - E1 == 0        -7410.660
## Luteal phase - Contralateral - E1 - Follicular phase - Ipsilateral - E2 == 0       9758.113
## Luteal phase - Contralateral - E2 - Follicular phase - Ipsilateral - E2 == 0       1440.754
## Luteal phase - Ipsilateral - E1 - Follicular phase - Ipsilateral - E2 == 0         9760.610
## Luteal phase - Ipsilateral - E2 - Follicular phase - Ipsilateral - E2 == 0          902.753
## Luteal phase - Contralateral - E2 - Luteal phase - Contralateral - E1 == 0        -8317.360
## Luteal phase - Ipsilateral - E1 - Luteal phase - Contralateral - E1 == 0              2.496
## Luteal phase - Ipsilateral - E2 - Luteal phase - Contralateral - E1 == 0          -8855.360
## Luteal phase - Ipsilateral - E1 - Luteal phase - Contralateral - E2 == 0           8319.856
```

```
## Luteal phase - Ipsilateral - E2 - Luteal phase - Contralateral - E2 == 0            -538.001
## Luteal phase - Ipsilateral - E2 - Luteal phase - Ipsilateral - E1 == 0             -8857.856
##                                                                                   Std. Error
## Follicular phase - Contralateral - E2 - Follicular phase - Contralateral - E1 == 0   1579.365
## Follicular phase - Ipsilateral - E1 - Follicular phase - Contralateral - E1 == 0     1431.421
## Follicular phase - Ipsilateral - E2 - Follicular phase - Contralateral - E1 == 0     1431.421
## Luteal phase - Contralateral - E1 - Follicular phase - Contralateral - E1 == 0       1412.627
## Luteal phase - Contralateral - E2 - Follicular phase - Contralateral - E1 == 0       1412.627
## Luteal phase - Ipsilateral - E1 - Follicular phase - Contralateral - E1 == 0         1381.114
## Luteal phase - Ipsilateral - E2 - Follicular phase - Contralateral - E1 == 0         1381.114
## Follicular phase - Ipsilateral - E1 - Follicular phase - Contralateral - E2 == 0     1431.421
## Follicular phase - Ipsilateral - E2 - Follicular phase - Contralateral - E2 == 0     1431.421
## Luteal phase - Contralateral - E1 - Follicular phase - Contralateral - E2 == 0       1412.627
## Luteal phase - Contralateral - E2 - Follicular phase - Contralateral - E2 == 0       1412.627
## Luteal phase - Ipsilateral - E1 - Follicular phase - Contralateral - E2 == 0         1381.114
## Luteal phase - Ipsilateral - E2 - Follicular phase - Contralateral - E2 == 0         1381.114
## Follicular phase - Ipsilateral - E2 - Follicular phase - Ipsilateral - E1 == 0       1266.309
## Luteal phase - Contralateral - E1 - Follicular phase - Ipsilateral - E1 == 0         1245.025
## Luteal phase - Contralateral - E2 - Follicular phase - Ipsilateral - E1 == 0         1245.025
## Luteal phase - Ipsilateral - E1 - Follicular phase - Ipsilateral - E1 == 0           1209.152
## Luteal phase - Ipsilateral - E2 - Follicular phase - Ipsilateral - E1 == 0           1209.152
## Luteal phase - Contralateral - E1 - Follicular phase - Ipsilateral - E2 == 0         1245.025
## Luteal phase - Contralateral - E2 - Follicular phase - Ipsilateral - E2 == 0         1245.025
## Luteal phase - Ipsilateral - E1 - Follicular phase - Ipsilateral - E2 == 0           1209.152
## Luteal phase - Ipsilateral - E2 - Follicular phase - Ipsilateral - E2 == 0           1209.152
## Luteal phase - Contralateral - E2 - Luteal phase - Contralateral - E1 == 0           1223.371
## Luteal phase - Ipsilateral - E1 - Luteal phase - Contralateral - E1 == 0             1186.844
## Luteal phase - Ipsilateral - E2 - Luteal phase - Contralateral - E1 == 0             1186.844
## Luteal phase - Ipsilateral - E1 - Luteal phase - Contralateral - E2 == 0             1186.844
## Luteal phase - Ipsilateral - E2 - Luteal phase - Contralateral - E2 == 0             1186.844
## Luteal phase - Ipsilateral - E2 - Luteal phase - Ipsilateral - E1 == 0               1149.157
##                                                                                   t value
## Follicular phase - Contralateral - E2 - Follicular phase - Contralateral - E1 == 0  -5.143
## Follicular phase - Ipsilateral - E1 - Follicular phase - Contralateral - E1 == 0     0.548
## Follicular phase - Ipsilateral - E2 - Follicular phase - Contralateral - E1 == 0    -5.260
## Luteal phase - Contralateral - E1 - Follicular phase - Contralateral - E1 == 0       1.578
## Luteal phase - Contralateral - E2 - Follicular phase - Contralateral - E1 == 0      -4.310
## Luteal phase - Ipsilateral - E1 - Follicular phase - Contralateral - E1 == 0         1.616
## Luteal phase - Ipsilateral - E2 - Follicular phase - Contralateral - E1 == 0        -4.798
## Follicular phase - Ipsilateral - E1 - Follicular phase - Contralateral - E2 == 0     6.222
## Follicular phase - Ipsilateral - E2 - Follicular phase - Contralateral - E2 == 0     0.415
## Luteal phase - Contralateral - E1 - Follicular phase - Contralateral - E2 == 0       7.328
## Luteal phase - Contralateral - E2 - Follicular phase - Contralateral - E2 == 0       1.440
## Luteal phase - Ipsilateral - E1 - Follicular phase - Contralateral - E2 == 0         7.497
## Luteal phase - Ipsilateral - E2 - Follicular phase - Contralateral - E2 == 0         1.083
## Follicular phase - Ipsilateral - E2 - Follicular phase - Ipsilateral - E1 == 0      -6.565
## Luteal phase - Contralateral - E1 - Follicular phase - Ipsilateral - E1 == 0         1.160
## Luteal phase - Contralateral - E2 - Follicular phase - Ipsilateral - E1 == 0        -5.520
## Luteal phase - Ipsilateral - E1 - Follicular phase - Ipsilateral - E1 == 0           1.197
## Luteal phase - Ipsilateral - E2 - Follicular phase - Ipsilateral - E1 == 0          -6.129
## Luteal phase - Contralateral - E1 - Follicular phase - Ipsilateral - E2 == 0         7.838
## Luteal phase - Contralateral - E2 - Follicular phase - Ipsilateral - E2 == 0         1.157
## Luteal phase - Ipsilateral - E1 - Follicular phase - Ipsilateral - E2 == 0           8.072
## Luteal phase - Ipsilateral - E2 - Follicular phase - Ipsilateral - E2 == 0           0.747
```

```
## Luteal phase - Contralateral - E2 - Luteal phase - Contralateral - E1 == 0          -6.799
## Luteal phase - Ipsilateral - E1 - Luteal phase - Contralateral - E1 == 0            0.002
## Luteal phase - Ipsilateral - E2 - Luteal phase - Contralateral - E1 == 0           -7.461
## Luteal phase - Ipsilateral - E1 - Luteal phase - Contralateral - E2 == 0            7.010
## Luteal phase - Ipsilateral - E2 - Luteal phase - Contralateral - E2 == 0           -0.453
## Luteal phase - Ipsilateral - E2 - Luteal phase - Ipsilateral - E1 == 0             -7.708
##                                                                                 Pr(>|t|)
## Follicular phase - Contralateral - E2 - Follicular phase - Contralateral - E1 == 0   <0.001
## Follicular phase - Ipsilateral - E1 - Follicular phase - Contralateral - E1 == 0      0.999
## Follicular phase - Ipsilateral - E2 - Follicular phase - Contralateral - E1 == 0     <0.001
## Luteal phase - Contralateral - E1 - Follicular phase - Contralateral - E1 == 0        0.761
## Luteal phase - Contralateral - E2 - Follicular phase - Contralateral - E1 == 0       <0.001
## Luteal phase - Ipsilateral - E1 - Follicular phase - Contralateral - E1 == 0          0.738
## Luteal phase - Ipsilateral - E2 - Follicular phase - Contralateral - E1 == 0         <0.001
## Follicular phase - Ipsilateral - E1 - Follicular phase - Contralateral - E2 == 0     <0.001
## Follicular phase - Ipsilateral - E2 - Follicular phase - Contralateral - E2 == 0      1.000
## Luteal phase - Contralateral - E1 - Follicular phase - Contralateral - E2 == 0       <0.001
## Luteal phase - Contralateral - E2 - Follicular phase - Contralateral - E2 == 0        0.836
## Luteal phase - Ipsilateral - E1 - Follicular phase - Contralateral - E2 == 0         <0.001
## Luteal phase - Ipsilateral - E2 - Follicular phase - Contralateral - E2 == 0          0.959
## Follicular phase - Ipsilateral - E2 - Follicular phase - Ipsilateral - E1 == 0       <0.001
## Luteal phase - Contralateral - E1 - Follicular phase - Ipsilateral - E1 == 0          0.942
## Luteal phase - Contralateral - E2 - Follicular phase - Ipsilateral - E1 == 0         <0.001
## Luteal phase - Ipsilateral - E1 - Follicular phase - Ipsilateral - E1 == 0            0.932
## Luteal phase - Ipsilateral - E2 - Follicular phase - Ipsilateral - E1 == 0           <0.001
## Luteal phase - Contralateral - E1 - Follicular phase - Ipsilateral - E2 == 0         <0.001
## Luteal phase - Contralateral - E2 - Follicular phase - Ipsilateral - E2 == 0          0.943
## Luteal phase - Ipsilateral - E1 - Follicular phase - Ipsilateral - E2 == 0           <0.001
## Luteal phase - Ipsilateral - E2 - Follicular phase - Ipsilateral - E2 == 0            0.995
## Luteal phase - Contralateral - E2 - Luteal phase - Contralateral - E1 == 0           <0.001
## Luteal phase - Ipsilateral - E1 - Luteal phase - Contralateral - E1 == 0              1.000
## Luteal phase - Ipsilateral - E2 - Luteal phase - Contralateral - E1 == 0             <0.001
## Luteal phase - Ipsilateral - E1 - Luteal phase - Contralateral - E2 == 0             <0.001
## Luteal phase - Ipsilateral - E2 - Luteal phase - Contralateral - E2 == 0              1.000
## Luteal phase - Ipsilateral - E2 - Luteal phase - Ipsilateral - E1 == 0               <0.001
##
## Follicular phase - Contralateral - E2 - Follicular phase - Contralateral - E1 == 0 ***
## Follicular phase - Ipsilateral - E1 - Follicular phase - Contralateral - E1 == 0
## Follicular phase - Ipsilateral - E2 - Follicular phase - Contralateral - E1 == 0   ***
## Luteal phase - Contralateral - E1 - Follicular phase - Contralateral - E1 == 0
## Luteal phase - Contralateral - E2 - Follicular phase - Contralateral - E1 == 0     ***
## Luteal phase - Ipsilateral - E1 - Follicular phase - Contralateral - E1 == 0
## Luteal phase - Ipsilateral - E2 - Follicular phase - Contralateral - E1 == 0       ***
## Follicular phase - Ipsilateral - E1 - Follicular phase - Contralateral - E2 == 0   ***
## Follicular phase - Ipsilateral - E2 - Follicular phase - Contralateral - E2 == 0
## Luteal phase - Contralateral - E1 - Follicular phase - Contralateral - E2 == 0     ***
## Luteal phase - Contralateral - E2 - Follicular phase - Contralateral - E2 == 0
## Luteal phase - Ipsilateral - E1 - Follicular phase - Contralateral - E2 == 0       ***
## Luteal phase - Ipsilateral - E2 - Follicular phase - Contralateral - E2 == 0
## Follicular phase - Ipsilateral - E2 - Follicular phase - Ipsilateral - E1 == 0     ***
## Luteal phase - Contralateral - E1 - Follicular phase - Ipsilateral - E1 == 0
## Luteal phase - Contralateral - E2 - Follicular phase - Ipsilateral - E1 == 0       ***
## Luteal phase - Ipsilateral - E1 - Follicular phase - Ipsilateral - E1 == 0
## Luteal phase - Ipsilateral - E2 - Follicular phase - Ipsilateral - E1 == 0         ***
```

```
## Luteal phase - Contralateral - E1 - Follicular phase - Ipsilateral - E2 == 0        ***
## Luteal phase - Contralateral - E2 - Follicular phase - Ipsilateral - E2 == 0
## Luteal phase - Ipsilateral - E1 - Follicular phase - Ipsilateral - E2 == 0          ***
## Luteal phase - Ipsilateral - E2 - Follicular phase - Ipsilateral - E2 == 0
## Luteal phase - Contralateral - E2 - Luteal phase - Contralateral - E1 == 0          ***
## Luteal phase - Ipsilateral - E1 - Luteal phase - Contralateral - E1 == 0
## Luteal phase - Ipsilateral - E2 - Luteal phase - Contralateral - E1 == 0            ***
## Luteal phase - Ipsilateral - E1 - Luteal phase - Contralateral - E2 == 0            ***
## Luteal phase - Ipsilateral - E2 - Luteal phase - Contralateral - E2 == 0
## Luteal phase - Ipsilateral - E2 - Luteal phase - Ipsilateral - E1 == 0              ***
## ---
## Signif. codes:  0 '***' 0.001 '**' 0.01 '*' 0.05 '.' 0.1 ' ' 1
## (Adjusted p values reported -- single-step method)
```

```
# Plot considering difference msucle-tunica and lumen-stroma####

# Plot Ampulla

#data input
data <- read.csv("Oviduct_rheo.csv")

# Subset the data based on X_position_group
data_ampulla <- subset(data, Tissue == "Ampulla")
data_isthmus <- subset(data, Tissue == "Isthmus")

png(filename="E1_E2_ampulla2.png",
    width = 9.86, height = 6.86, units = "in",
    res = 600)

par(mgp=c(1.5,0.5,0),
    mar = c(4, #bottom
            4, #left
            3, #top
            2),
    cex.main = 0.8,
    family = "sans" #right
)

# Create a new column to define the X_position group
data_ampulla$X_position_group <- ifelse(data_ampulla$X_position <= 400, "Lumen-Stroma surface", "Muscle-

# Calculate means and standard deviations
data_summary <- data_ampulla %>%
  group_by(Module, Estrus_cycle, Position, Hz, X_position_group) %>%
  summarise(mean_Indentation = mean(Indentation, na.rm = TRUE),
            sd_Indentation = sd(Indentation, na.rm = TRUE))
```

```
## `summarise()` has grouped output by 'Module', 'Estrus_cycle', 'Position', 'Hz'.
## You can override using the `.groups` argument.
```

```
# Create a new variable for line type
data_summary <- data_summary %>%
  mutate(LineType = ifelse(Module == "E1", "E1", "E2"))

# Define fill colors for each factor level
```

```r
fill_colors <- c("Follicular phase Contralateral" = viridis_pal(option = "D")(1),
                 "Follicular phase Ipsilateral" = alpha(viridis_pal(option = "D")(1), 0.2),
                 "Luteal phase Contralateral" = "#238A8DFF",
                 "Luteal phase Ipsilateral" = alpha("#238A8DFF", 0.2))

# Create the line plot with facets for X_position_group
ggplot(data_summary, aes(x = Hz, y = mean_Indentation, group = interaction(Module, Estrus_cycle, Positi
                         color = paste(Estrus_cycle, Position), linetype = LineType)) +
  geom_line(size = 1.5) +
  geom_point(size = 1) +
  scale_color_manual(values = fill_colors) +
  scale_linetype_manual(values = c("E1" = "solid", "E2" = "dashed"),
                        labels = c("E1" = "E'", "E2" = "E\"")) +
  scale_x_continuous(breaks = c(1, 5, 10, 20)) +
  scale_y_continuous(labels = scientific_format(), limits = c(0, 16000)) +
  labs(y = expression(paste("E'/E'' (Pa)")), x = "Frequency (Hz)", color = "Estrus Cycle + Position") +
  theme_minimal() +
  theme(panel.grid.major = element_blank(),
        panel.grid.minor = element_blank(),
        panel.border = element_rect(colour = "black", fill = NA, size = 1),
        axis.text = element_text(size = 12)) +
  facet_wrap(~ X_position_group, nrow = 1)
```

```
## Warning: Removed 4 rows containing missing values (`geom_line()`).

## Warning: Removed 4 rows containing missing values (`geom_point()`).
```

```r
dev.off()
```

```
## pdf
##   2
```

```r
# Stats Ampulla

df <- data_ampulla

# Create a factor variable for the comparisons
df$Comparison <- factor(paste(df$Estrus_cycle, df$Position, df$Module, df$X_position_group, sep = " - "

# Perform pairwise comparisons using Tukey's method
model <- lm(Indentation ~ Comparison, data = df)
comp <- glht(model, linfct = mcp(Comparison = "Tukey"))

# Summarize the results
summary(comp)
```

```
## Warning in RET$pfunction("adjusted", ...): Completion with error > abseps

## Warning in RET$pfunction("adjusted", ...): Completion with error > abseps

## Warning in RET$pfunction("adjusted", ...): Completion with error > abseps

## Warning in RET$pfunction("adjusted", ...): Completion with error > abseps

## Warning in RET$pfunction("adjusted", ...): Completion with error > abseps
```

```
## Warning in RET$pfunction("adjusted", ...): Completion with error > abseps

## Warning in RET$pfunction("adjusted", ...): Completion with error > abseps

## Warning in RET$pfunction("adjusted", ...): Completion with error > abseps

## Warning in RET$pfunction("adjusted", ...): Completion with error > abseps

## Warning in RET$pfunction("adjusted", ...): Completion with error > abseps

## Warning in RET$pfunction("adjusted", ...): Completion with error > abseps

## Warning in RET$pfunction("adjusted", ...): Completion with error > abseps

## Warning in RET$pfunction("adjusted", ...): Completion with error > abseps

## Warning in RET$pfunction("adjusted", ...): Completion with error > abseps

## Warning in RET$pfunction("adjusted", ...): Completion with error > abseps

## Warning in RET$pfunction("adjusted", ...): Completion with error > abseps

## Warning in RET$pfunction("adjusted", ...): Completion with error > abseps

## Warning in RET$pfunction("adjusted", ...): Completion with error > abseps

## Warning in RET$pfunction("adjusted", ...): Completion with error > abseps

## Warning in RET$pfunction("adjusted", ...): Completion with error > abseps

## Warning in RET$pfunction("adjusted", ...): Completion with error > abseps

## Warning in RET$pfunction("adjusted", ...): Completion with error > abseps

## Warning in RET$pfunction("adjusted", ...): Completion with error > abseps

## Warning in RET$pfunction("adjusted", ...): Completion with error > abseps

## Warning in RET$pfunction("adjusted", ...): Completion with error > abseps

## Warning in RET$pfunction("adjusted", ...): Completion with error > abseps

## Warning in RET$pfunction("adjusted", ...): Completion with error > abseps

## Warning in RET$pfunction("adjusted", ...): Completion with error > abseps
##
##   Simultaneous Tests for General Linear Hypotheses
##
## Multiple Comparisons of Means: Tukey Contrasts
##
##
## Fit: lm(formula = Indentation ~ Comparison, data = df)
##
```

```
## Linear Hypotheses:
##
## Follicular phase - Contralateral - E1 - Muscle-Tunica surface - Follicular phase - Contralateral - E
## Follicular phase - Contralateral - E2 - Lumen-Stroma surface - Follicular phase - Contralateral - E1
## Follicular phase - Contralateral - E2 - Muscle-Tunica surface - Follicular phase - Contralateral - E
## Follicular phase - Ipsilateral - E1 - Lumen-Stroma surface - Follicular phase - Contralateral - E1 -
## Follicular phase - Ipsilateral - E1 - Muscle-Tunica surface - Follicular phase - Contralateral - E1 -
## Follicular phase - Ipsilateral - E2 - Lumen-Stroma surface - Follicular phase - Contralateral - E1 -
## Follicular phase - Ipsilateral - E2 - Muscle-Tunica surface - Follicular phase - Contralateral - E1 -
## Luteal phase - Contralateral - E1 - Lumen-Stroma surface - Follicular phase - Contralateral - E1 - Lu
## Luteal phase - Contralateral - E1 - Muscle-Tunica surface - Follicular phase - Contralateral - E1 - M
## Luteal phase - Contralateral - E2 - Lumen-Stroma surface - Follicular phase - Contralateral - E1 - Lu
## Luteal phase - Contralateral - E2 - Muscle-Tunica surface - Follicular phase - Contralateral - E1 - M
## Luteal phase - Ipsilateral - E1 - Lumen-Stroma surface - Follicular phase - Contralateral - E1 - Lum
## Luteal phase - Ipsilateral - E1 - Muscle-Tunica surface - Follicular phase - Contralateral - E1 - Lu
## Luteal phase - Ipsilateral - E2 - Lumen-Stroma surface - Follicular phase - Contralateral - E1 - Lum
## Luteal phase - Ipsilateral - E2 - Muscle-Tunica surface - Follicular phase - Contralateral - E1 - Lu
## Follicular phase - Contralateral - E2 - Lumen-Stroma surface - Follicular phase - Contralateral - E1
## Follicular phase - Contralateral - E2 - Muscle-Tunica surface - Follicular phase - Contralateral - E
## Follicular phase - Ipsilateral - E1 - Lumen-Stroma surface - Follicular phase - Contralateral - E1 -
## Follicular phase - Ipsilateral - E1 - Muscle-Tunica surface - Follicular phase - Contralateral - E1 -
## Follicular phase - Ipsilateral - E2 - Lumen-Stroma surface - Follicular phase - Contralateral - E1 -
## Follicular phase - Ipsilateral - E2 - Muscle-Tunica surface - Follicular phase - Contralateral - E1 -
## Luteal phase - Contralateral - E1 - Lumen-Stroma surface - Follicular phase - Contralateral - E1 - Mu
## Luteal phase - Contralateral - E1 - Muscle-Tunica surface - Follicular phase - Contralateral - E1 - M
## Luteal phase - Contralateral - E2 - Lumen-Stroma surface - Follicular phase - Contralateral - E1 - Mu
## Luteal phase - Contralateral - E2 - Muscle-Tunica surface - Follicular phase - Contralateral - E1 - M
## Luteal phase - Ipsilateral - E1 - Lumen-Stroma surface - Follicular phase - Contralateral - E1 - Mus
## Luteal phase - Ipsilateral - E1 - Muscle-Tunica surface - Follicular phase - Contralateral - E1 - Mu
## Luteal phase - Ipsilateral - E2 - Lumen-Stroma surface - Follicular phase - Contralateral - E1 - Mus
## Luteal phase - Ipsilateral - E2 - Muscle-Tunica surface - Follicular phase - Contralateral - E1 - Mu
## Follicular phase - Contralateral - E2 - Muscle-Tunica surface - Follicular phase - Contralateral - E
## Follicular phase - Ipsilateral - E1 - Lumen-Stroma surface - Follicular phase - Contralateral - E2 -
## Follicular phase - Ipsilateral - E1 - Muscle-Tunica surface - Follicular phase - Contralateral - E2 -
## Follicular phase - Ipsilateral - E2 - Lumen-Stroma surface - Follicular phase - Contralateral - E2 -
## Follicular phase - Ipsilateral - E2 - Muscle-Tunica surface - Follicular phase - Contralateral - E2 -
## Luteal phase - Contralateral - E1 - Lumen-Stroma surface - Follicular phase - Contralateral - E2 - Lu
## Luteal phase - Contralateral - E1 - Muscle-Tunica surface - Follicular phase - Contralateral - E2 - M
## Luteal phase - Contralateral - E2 - Lumen-Stroma surface - Follicular phase - Contralateral - E2 - Lu
## Luteal phase - Contralateral - E2 - Muscle-Tunica surface - Follicular phase - Contralateral - E2 - M
## Luteal phase - Ipsilateral - E1 - Lumen-Stroma surface - Follicular phase - Contralateral - E2 - Lum
## Luteal phase - Ipsilateral - E1 - Muscle-Tunica surface - Follicular phase - Contralateral - E2 - Lu
## Luteal phase - Ipsilateral - E2 - Lumen-Stroma surface - Follicular phase - Contralateral - E2 - Lum
## Luteal phase - Ipsilateral - E2 - Muscle-Tunica surface - Follicular phase - Contralateral - E2 - Lu
## Follicular phase - Ipsilateral - E1 - Lumen-Stroma surface - Follicular phase - Contralateral - E2 -
## Follicular phase - Ipsilateral - E1 - Muscle-Tunica surface - Follicular phase - Contralateral - E2 -
## Follicular phase - Ipsilateral - E2 - Lumen-Stroma surface - Follicular phase - Contralateral - E2 -
## Follicular phase - Ipsilateral - E2 - Muscle-Tunica surface - Follicular phase - Contralateral - E2 -
## Luteal phase - Contralateral - E1 - Lumen-Stroma surface - Follicular phase - Contralateral - E2 - Mu
## Luteal phase - Contralateral - E1 - Muscle-Tunica surface - Follicular phase - Contralateral - E2 - M
## Luteal phase - Contralateral - E2 - Lumen-Stroma surface - Follicular phase - Contralateral - E2 - Mu
## Luteal phase - Contralateral - E2 - Muscle-Tunica surface - Follicular phase - Contralateral - E2 - M
## Luteal phase - Ipsilateral - E1 - Lumen-Stroma surface - Follicular phase - Contralateral - E2 - Mus
## Luteal phase - Ipsilateral - E1 - Muscle-Tunica surface - Follicular phase - Contralateral - E2 - Mu
```

```
## Luteal phase - Ipsilateral - E2 - Lumen-Stroma surface - Follicular phase - Contralateral - E2 - Musc
## Luteal phase - Ipsilateral - E2 - Muscle-Tunica surface - Follicular phase - Contralateral - E2 - Mu
## Follicular phase - Ipsilateral - E1 - Muscle-Tunica surface - Follicular phase - Ipsilateral - E1 -
## Follicular phase - Ipsilateral - E2 - Lumen-Stroma surface - Follicular phase - Ipsilateral - E1 - Lu
## Follicular phase - Ipsilateral - E2 - Muscle-Tunica surface - Follicular phase - Ipsilateral - E1 -
## Luteal phase - Contralateral - E1 - Lumen-Stroma surface - Follicular phase - Ipsilateral - E1 - Lum
## Luteal phase - Contralateral - E1 - Muscle-Tunica surface - Follicular phase - Ipsilateral - E1 - Lu
## Luteal phase - Contralateral - E2 - Lumen-Stroma surface - Follicular phase - Ipsilateral - E1 - Lum
## Luteal phase - Contralateral - E2 - Muscle-Tunica surface - Follicular phase - Ipsilateral - E1 - Lu
## Luteal phase - Ipsilateral - E1 - Lumen-Stroma surface - Follicular phase - Ipsilateral - E1 - Lumen-
## Luteal phase - Ipsilateral - E1 - Muscle-Tunica surface - Follicular phase - Ipsilateral - E1 - Lume
## Luteal phase - Ipsilateral - E2 - Lumen-Stroma surface - Follicular phase - Ipsilateral - E1 - Lumen-
## Luteal phase - Ipsilateral - E2 - Muscle-Tunica surface - Follicular phase - Ipsilateral - E1 - Lume
## Follicular phase - Ipsilateral - E2 - Lumen-Stroma surface - Follicular phase - Ipsilateral - E1 - Mu
## Follicular phase - Ipsilateral - E2 - Muscle-Tunica surface - Follicular phase - Ipsilateral - E1 -
## Luteal phase - Contralateral - E1 - Lumen-Stroma surface - Follicular phase - Ipsilateral - E1 - Musc
## Luteal phase - Contralateral - E1 - Muscle-Tunica surface - Follicular phase - Ipsilateral - E1 - Mu
## Luteal phase - Contralateral - E2 - Lumen-Stroma surface - Follicular phase - Ipsilateral - E1 - Musc
## Luteal phase - Contralateral - E2 - Muscle-Tunica surface - Follicular phase - Ipsilateral - E1 - Mu
## Luteal phase - Ipsilateral - E1 - Lumen-Stroma surface - Follicular phase - Ipsilateral - E1 - Muscle
## Luteal phase - Ipsilateral - E1 - Muscle-Tunica surface - Follicular phase - Ipsilateral - E1 - Muscl
## Luteal phase - Ipsilateral - E2 - Lumen-Stroma surface - Follicular phase - Ipsilateral - E1 - Muscle
## Luteal phase - Ipsilateral - E2 - Muscle-Tunica surface - Follicular phase - Ipsilateral - E1 - Muscl
## Follicular phase - Ipsilateral - E2 - Muscle-Tunica surface - Follicular phase - Ipsilateral - E2 -
## Luteal phase - Contralateral - E1 - Lumen-Stroma surface - Follicular phase - Ipsilateral - E2 - Lum
## Luteal phase - Contralateral - E1 - Muscle-Tunica surface - Follicular phase - Ipsilateral - E2 - Lu
## Luteal phase - Contralateral - E2 - Lumen-Stroma surface - Follicular phase - Ipsilateral - E2 - Lum
## Luteal phase - Contralateral - E2 - Muscle-Tunica surface - Follicular phase - Ipsilateral - E2 - Lu
## Luteal phase - Ipsilateral - E1 - Lumen-Stroma surface - Follicular phase - Ipsilateral - E2 - Lumen-
## Luteal phase - Ipsilateral - E1 - Muscle-Tunica surface - Follicular phase - Ipsilateral - E2 - Lume
## Luteal phase - Ipsilateral - E2 - Lumen-Stroma surface - Follicular phase - Ipsilateral - E2 - Lumen-
## Luteal phase - Ipsilateral - E2 - Muscle-Tunica surface - Follicular phase - Ipsilateral - E2 - Lume
## Luteal phase - Contralateral - E1 - Lumen-Stroma surface - Follicular phase - Ipsilateral - E2 - Musc
## Luteal phase - Contralateral - E1 - Muscle-Tunica surface - Follicular phase - Ipsilateral - E2 - Mu
## Luteal phase - Contralateral - E2 - Lumen-Stroma surface - Follicular phase - Ipsilateral - E2 - Musc
## Luteal phase - Contralateral - E2 - Muscle-Tunica surface - Follicular phase - Ipsilateral - E2 - Mu
## Luteal phase - Ipsilateral - E1 - Lumen-Stroma surface - Follicular phase - Ipsilateral - E2 - Muscle
## Luteal phase - Ipsilateral - E1 - Muscle-Tunica surface - Follicular phase - Ipsilateral - E2 - Muscl
## Luteal phase - Ipsilateral - E2 - Lumen-Stroma surface - Follicular phase - Ipsilateral - E2 - Muscle
## Luteal phase - Ipsilateral - E2 - Muscle-Tunica surface - Follicular phase - Ipsilateral - E2 - Muscl
## Luteal phase - Contralateral - E1 - Muscle-Tunica surface - Luteal phase - Contralateral - E1 - Lume
## Luteal phase - Contralateral - E2 - Lumen-Stroma surface - Luteal phase - Contralateral - E1 - Lumen-
## Luteal phase - Contralateral - E2 - Muscle-Tunica surface - Luteal phase - Contralateral - E1 - Lume
## Luteal phase - Ipsilateral - E1 - Lumen-Stroma surface - Luteal phase - Contralateral - E1 - Lumen-S
## Luteal phase - Ipsilateral - E1 - Muscle-Tunica surface - Luteal phase - Contralateral - E1 - Lumen-
## Luteal phase - Ipsilateral - E2 - Lumen-Stroma surface - Luteal phase - Contralateral - E1 - Lumen-S
## Luteal phase - Ipsilateral - E2 - Muscle-Tunica surface - Luteal phase - Contralateral - E1 - Lumen-
## Luteal phase - Contralateral - E2 - Lumen-Stroma surface - Luteal phase - Contralateral - E1 - Muscle
## Luteal phase - Contralateral - E2 - Muscle-Tunica surface - Luteal phase - Contralateral - E1 - Muscl
## Luteal phase - Ipsilateral - E1 - Lumen-Stroma surface - Luteal phase - Contralateral - E1 - Muscle-
## Luteal phase - Ipsilateral - E1 - Muscle-Tunica surface - Luteal phase - Contralateral - E1 - Muscle
## Luteal phase - Ipsilateral - E2 - Lumen-Stroma surface - Luteal phase - Contralateral - E1 - Muscle-
## Luteal phase - Ipsilateral - E2 - Muscle-Tunica surface - Luteal phase - Contralateral - E1 - Muscle
## Luteal phase - Contralateral - E2 - Muscle-Tunica surface - Luteal phase - Contralateral - E2 - Lume
```

```
## Luteal phase - Ipsilateral - E1 - Lumen-Stroma surface - Luteal phase - Contralateral - E2 - Lumen-St
## Luteal phase - Ipsilateral - E1 - Muscle-Tunica surface - Luteal phase - Contralateral - E2 - Lumen-S
## Luteal phase - Ipsilateral - E2 - Lumen-Stroma surface - Luteal phase - Contralateral - E2 - Lumen-St
## Luteal phase - Ipsilateral - E2 - Muscle-Tunica surface - Luteal phase - Contralateral - E2 - Lumen-S
## Luteal phase - Ipsilateral - E1 - Lumen-Stroma surface - Luteal phase - Contralateral - E2 - Muscle-T
## Luteal phase - Ipsilateral - E1 - Muscle-Tunica surface - Luteal phase - Contralateral - E2 - Muscle-
## Luteal phase - Ipsilateral - E2 - Lumen-Stroma surface - Luteal phase - Contralateral - E2 - Muscle-T
## Luteal phase - Ipsilateral - E2 - Muscle-Tunica surface - Luteal phase - Contralateral - E2 - Muscle-
## Luteal phase - Ipsilateral - E1 - Muscle-Tunica surface - Luteal phase - Ipsilateral - E1 - Lumen-Str
## Luteal phase - Ipsilateral - E2 - Lumen-Stroma surface - Luteal phase - Ipsilateral - E1 - Lumen-Stro
## Luteal phase - Ipsilateral - E2 - Muscle-Tunica surface - Luteal phase - Ipsilateral - E1 - Lumen-Str
## Luteal phase - Ipsilateral - E2 - Lumen-Stroma surface - Luteal phase - Ipsilateral - E1 - Muscle-Tun
## Luteal phase - Ipsilateral - E2 - Muscle-Tunica surface - Luteal phase - Ipsilateral - E1 - Muscle-Tu
## Luteal phase - Ipsilateral - E2 - Muscle-Tunica surface - Luteal phase - Ipsilateral - E2 - Lumen-Str
##
## Follicular phase - Contralateral - E1 - Muscle-Tunica surface - Follicular phase - Contralateral - E
## Follicular phase - Contralateral - E2 - Lumen-Stroma surface - Follicular phase - Contralateral - E1
## Follicular phase - Contralateral - E2 - Muscle-Tunica surface - Follicular phase - Contralateral - E
## Follicular phase - Ipsilateral - E1 - Lumen-Stroma surface - Follicular phase - Contralateral - E1 -
## Follicular phase - Ipsilateral - E1 - Muscle-Tunica surface - Follicular phase - Contralateral - E1 -
## Follicular phase - Ipsilateral - E2 - Lumen-Stroma surface - Follicular phase - Contralateral - E1 -
## Follicular phase - Ipsilateral - E2 - Muscle-Tunica surface - Follicular phase - Contralateral - E1 -
## Luteal phase - Contralateral - E1 - Lumen-Stroma surface - Follicular phase - Contralateral - E1 - Lu
## Luteal phase - Contralateral - E1 - Muscle-Tunica surface - Follicular phase - Contralateral - E1 - L
## Luteal phase - Contralateral - E2 - Lumen-Stroma surface - Follicular phase - Contralateral - E1 - Lu
## Luteal phase - Contralateral - E2 - Muscle-Tunica surface - Follicular phase - Contralateral - E1 - L
## Luteal phase - Ipsilateral - E1 - Lumen-Stroma surface - Follicular phase - Contralateral - E1 - Lume
## Luteal phase - Ipsilateral - E1 - Muscle-Tunica surface - Follicular phase - Contralateral - E1 - Lum
## Luteal phase - Ipsilateral - E2 - Lumen-Stroma surface - Follicular phase - Contralateral - E1 - Lume
## Luteal phase - Ipsilateral - E2 - Muscle-Tunica surface - Follicular phase - Contralateral - E1 - Lum
## Follicular phase - Contralateral - E2 - Lumen-Stroma surface - Follicular phase - Contralateral - E1
## Follicular phase - Contralateral - E2 - Muscle-Tunica surface - Follicular phase - Contralateral - E
## Follicular phase - Ipsilateral - E1 - Lumen-Stroma surface - Follicular phase - Contralateral - E1 -
## Follicular phase - Ipsilateral - E1 - Muscle-Tunica surface - Follicular phase - Contralateral - E1 -
## Follicular phase - Ipsilateral - E2 - Lumen-Stroma surface - Follicular phase - Contralateral - E1 -
## Follicular phase - Ipsilateral - E2 - Muscle-Tunica surface - Follicular phase - Contralateral - E1 -
## Luteal phase - Contralateral - E1 - Lumen-Stroma surface - Follicular phase - Contralateral - E1 - Mu
## Luteal phase - Contralateral - E1 - Muscle-Tunica surface - Follicular phase - Contralateral - E1 - M
## Luteal phase - Contralateral - E2 - Lumen-Stroma surface - Follicular phase - Contralateral - E1 - Mu
## Luteal phase - Contralateral - E2 - Muscle-Tunica surface - Follicular phase - Contralateral - E1 - M
## Luteal phase - Ipsilateral - E1 - Lumen-Stroma surface - Follicular phase - Contralateral - E1 - Musc
## Luteal phase - Ipsilateral - E1 - Muscle-Tunica surface - Follicular phase - Contralateral - E1 - Mus
## Luteal phase - Ipsilateral - E2 - Lumen-Stroma surface - Follicular phase - Contralateral - E1 - Musc
## Luteal phase - Ipsilateral - E2 - Muscle-Tunica surface - Follicular phase - Contralateral - E1 - Mus
## Follicular phase - Contralateral - E2 - Muscle-Tunica surface - Follicular phase - Contralateral - E
## Follicular phase - Ipsilateral - E1 - Lumen-Stroma surface - Follicular phase - Contralateral - E2 -
## Follicular phase - Ipsilateral - E1 - Muscle-Tunica surface - Follicular phase - Contralateral - E2 -
## Follicular phase - Ipsilateral - E2 - Lumen-Stroma surface - Follicular phase - Contralateral - E2 -
## Follicular phase - Ipsilateral - E2 - Muscle-Tunica surface - Follicular phase - Contralateral - E2 -
## Luteal phase - Contralateral - E1 - Lumen-Stroma surface - Follicular phase - Contralateral - E2 - Lu
## Luteal phase - Contralateral - E1 - Muscle-Tunica surface - Follicular phase - Contralateral - E2 - L
## Luteal phase - Contralateral - E2 - Lumen-Stroma surface - Follicular phase - Contralateral - E2 - Lu
## Luteal phase - Contralateral - E2 - Muscle-Tunica surface - Follicular phase - Contralateral - E2 - L
## Luteal phase - Ipsilateral - E1 - Lumen-Stroma surface - Follicular phase - Contralateral - E2 - Lume
```

```
## Luteal phase - Ipsilateral - E1 - Muscle-Tunica surface - Follicular phase - Contralateral - E2 - Lu
## Luteal phase - Ipsilateral - E2 - Lumen-Stroma surface - Follicular phase - Contralateral - E2 - Lum
## Luteal phase - Ipsilateral - E2 - Muscle-Tunica surface - Follicular phase - Contralateral - E2 - Lu
## Follicular phase - Ipsilateral - E1 - Lumen-Stroma surface - Follicular phase - Contralateral - E2 -
## Follicular phase - Ipsilateral - E1 - Muscle-Tunica surface - Follicular phase - Contralateral - E2 -
## Follicular phase - Ipsilateral - E2 - Lumen-Stroma surface - Follicular phase - Contralateral - E2 -
## Follicular phase - Ipsilateral - E2 - Muscle-Tunica surface - Follicular phase - Contralateral - E2 -
## Luteal phase - Contralateral - E1 - Lumen-Stroma surface - Follicular phase - Contralateral - E2 - Mu
## Luteal phase - Contralateral - E1 - Muscle-Tunica surface - Follicular phase - Contralateral - E2 -
## Luteal phase - Contralateral - E2 - Lumen-Stroma surface - Follicular phase - Contralateral - E2 - Mu
## Luteal phase - Contralateral - E2 - Muscle-Tunica surface - Follicular phase - Contralateral - E2 -
## Luteal phase - Ipsilateral - E1 - Lumen-Stroma surface - Follicular phase - Contralateral - E2 - Mus
## Luteal phase - Ipsilateral - E1 - Muscle-Tunica surface - Follicular phase - Contralateral - E2 - Mu
## Luteal phase - Ipsilateral - E2 - Lumen-Stroma surface - Follicular phase - Contralateral - E2 - Mus
## Luteal phase - Ipsilateral - E2 - Muscle-Tunica surface - Follicular phase - Contralateral - E2 - Mu
## Follicular phase - Ipsilateral - E1 - Muscle-Tunica surface - Follicular phase - Ipsilateral - E1 -
## Follicular phase - Ipsilateral - E2 - Lumen-Stroma surface - Follicular phase - Ipsilateral - E1 - Lu
## Follicular phase - Ipsilateral - E2 - Muscle-Tunica surface - Follicular phase - Ipsilateral - E1 -
## Luteal phase - Contralateral - E1 - Lumen-Stroma surface - Follicular phase - Ipsilateral - E1 - Lum
## Luteal phase - Contralateral - E1 - Muscle-Tunica surface - Follicular phase - Ipsilateral - E1 - Lu
## Luteal phase - Contralateral - E2 - Lumen-Stroma surface - Follicular phase - Ipsilateral - E1 - Lum
## Luteal phase - Contralateral - E2 - Muscle-Tunica surface - Follicular phase - Ipsilateral - E1 - Lu
## Luteal phase - Ipsilateral - E1 - Lumen-Stroma surface - Follicular phase - Ipsilateral - E1 - Lumen-
## Luteal phase - Ipsilateral - E1 - Muscle-Tunica surface - Follicular phase - Ipsilateral - E1 - Lumen
## Luteal phase - Ipsilateral - E2 - Lumen-Stroma surface - Follicular phase - Ipsilateral - E1 - Lumen-
## Luteal phase - Ipsilateral - E2 - Muscle-Tunica surface - Follicular phase - Ipsilateral - E1 - Lumen
## Follicular phase - Ipsilateral - E2 - Lumen-Stroma surface - Follicular phase - Ipsilateral - E1 - Mu
## Follicular phase - Ipsilateral - E2 - Muscle-Tunica surface - Follicular phase - Ipsilateral - E1 -
## Luteal phase - Contralateral - E1 - Lumen-Stroma surface - Follicular phase - Ipsilateral - E1 - Mus
## Luteal phase - Contralateral - E1 - Muscle-Tunica surface - Follicular phase - Ipsilateral - E1 - Mu
## Luteal phase - Contralateral - E2 - Lumen-Stroma surface - Follicular phase - Ipsilateral - E1 - Mus
## Luteal phase - Contralateral - E2 - Muscle-Tunica surface - Follicular phase - Ipsilateral - E1 - Mu
## Luteal phase - Ipsilateral - E1 - Lumen-Stroma surface - Follicular phase - Ipsilateral - E1 - Muscl
## Luteal phase - Ipsilateral - E1 - Muscle-Tunica surface - Follicular phase - Ipsilateral - E1 - Musc
## Luteal phase - Ipsilateral - E2 - Lumen-Stroma surface - Follicular phase - Ipsilateral - E1 - Muscl
## Luteal phase - Ipsilateral - E2 - Muscle-Tunica surface - Follicular phase - Ipsilateral - E1 - Musc
## Follicular phase - Ipsilateral - E2 - Muscle-Tunica surface - Follicular phase - Ipsilateral - E2 -
## Luteal phase - Contralateral - E1 - Lumen-Stroma surface - Follicular phase - Ipsilateral - E2 - Lum
## Luteal phase - Contralateral - E1 - Muscle-Tunica surface - Follicular phase - Ipsilateral - E2 - Lu
## Luteal phase - Contralateral - E2 - Lumen-Stroma surface - Follicular phase - Ipsilateral - E2 - Lum
## Luteal phase - Contralateral - E2 - Muscle-Tunica surface - Follicular phase - Ipsilateral - E2 - Lu
## Luteal phase - Ipsilateral - E1 - Lumen-Stroma surface - Follicular phase - Ipsilateral - E2 - Lumen-
## Luteal phase - Ipsilateral - E1 - Muscle-Tunica surface - Follicular phase - Ipsilateral - E2 - Lumen
## Luteal phase - Ipsilateral - E2 - Lumen-Stroma surface - Follicular phase - Ipsilateral - E2 - Lumen-
## Luteal phase - Ipsilateral - E2 - Muscle-Tunica surface - Follicular phase - Ipsilateral - E2 - Lumen
## Luteal phase - Contralateral - E1 - Lumen-Stroma surface - Follicular phase - Ipsilateral - E2 - Mus
## Luteal phase - Contralateral - E1 - Muscle-Tunica surface - Follicular phase - Ipsilateral - E2 - Mu
## Luteal phase - Contralateral - E2 - Lumen-Stroma surface - Follicular phase - Ipsilateral - E2 - Mus
## Luteal phase - Contralateral - E2 - Muscle-Tunica surface - Follicular phase - Ipsilateral - E2 - Mu
## Luteal phase - Ipsilateral - E1 - Lumen-Stroma surface - Follicular phase - Ipsilateral - E2 - Muscl
## Luteal phase - Ipsilateral - E1 - Muscle-Tunica surface - Follicular phase - Ipsilateral - E2 - Musc
## Luteal phase - Ipsilateral - E2 - Lumen-Stroma surface - Follicular phase - Ipsilateral - E2 - Muscl
## Luteal phase - Ipsilateral - E2 - Muscle-Tunica surface - Follicular phase - Ipsilateral - E2 - Musc
## Luteal phase - Contralateral - E1 - Muscle-Tunica surface - Luteal phase - Contralateral - E1 - Lumen
```

```
## Luteal phase - Contralateral - E2 - Lumen-Stroma surface - Luteal phase - Contralateral - E1 - Lumen
## Luteal phase - Contralateral - E2 - Muscle-Tunica surface - Luteal phase - Contralateral - E1 - Lumen
## Luteal phase - Ipsilateral - E1 - Lumen-Stroma surface - Luteal phase - Contralateral - E1 - Lumen-St
## Luteal phase - Ipsilateral - E1 - Muscle-Tunica surface - Luteal phase - Contralateral - E1 - Lumen-
## Luteal phase - Ipsilateral - E2 - Lumen-Stroma surface - Luteal phase - Contralateral - E1 - Lumen-St
## Luteal phase - Ipsilateral - E2 - Muscle-Tunica surface - Luteal phase - Contralateral - E1 - Lumen-
## Luteal phase - Contralateral - E2 - Lumen-Stroma surface - Luteal phase - Contralateral - E1 - Muscle
## Luteal phase - Contralateral - E2 - Muscle-Tunica surface - Luteal phase - Contralateral - E1 - Muscl
## Luteal phase - Ipsilateral - E1 - Lumen-Stroma surface - Luteal phase - Contralateral - E1 - Muscle-
## Luteal phase - Ipsilateral - E1 - Muscle-Tunica surface - Luteal phase - Contralateral - E1 - Muscle
## Luteal phase - Ipsilateral - E2 - Lumen-Stroma surface - Luteal phase - Contralateral - E1 - Muscle-
## Luteal phase - Ipsilateral - E2 - Muscle-Tunica surface - Luteal phase - Contralateral - E1 - Muscle
## Luteal phase - Contralateral - E2 - Muscle-Tunica surface - Luteal phase - Contralateral - E2 - Lumen
## Luteal phase - Ipsilateral - E1 - Lumen-Stroma surface - Luteal phase - Contralateral - E2 - Lumen-St
## Luteal phase - Ipsilateral - E1 - Muscle-Tunica surface - Luteal phase - Contralateral - E2 - Lumen-
## Luteal phase - Ipsilateral - E2 - Lumen-Stroma surface - Luteal phase - Contralateral - E2 - Lumen-St
## Luteal phase - Ipsilateral - E2 - Muscle-Tunica surface - Luteal phase - Contralateral - E2 - Lumen-
## Luteal phase - Ipsilateral - E1 - Lumen-Stroma surface - Luteal phase - Contralateral - E2 - Muscle-
## Luteal phase - Ipsilateral - E1 - Muscle-Tunica surface - Luteal phase - Contralateral - E2 - Muscle
## Luteal phase - Ipsilateral - E2 - Lumen-Stroma surface - Luteal phase - Contralateral - E2 - Muscle-
## Luteal phase - Ipsilateral - E2 - Muscle-Tunica surface - Luteal phase - Contralateral - E2 - Muscle
## Luteal phase - Ipsilateral - E1 - Muscle-Tunica surface - Luteal phase - Ipsilateral - E1 - Lumen-St
## Luteal phase - Ipsilateral - E2 - Lumen-Stroma surface - Luteal phase - Ipsilateral - E1 - Lumen-Str
## Luteal phase - Ipsilateral - E2 - Muscle-Tunica surface - Luteal phase - Ipsilateral - E1 - Lumen-St
## Luteal phase - Ipsilateral - E2 - Lumen-Stroma surface - Luteal phase - Ipsilateral - E1 - Muscle-Tu
## Luteal phase - Ipsilateral - E2 - Muscle-Tunica surface - Luteal phase - Ipsilateral - E1 - Muscle-T
## Luteal phase - Ipsilateral - E2 - Muscle-Tunica surface - Luteal phase - Ipsilateral - E2 - Lumen-St
##
## Follicular phase - Contralateral - E1 - Muscle-Tunica surface - Follicular phase - Contralateral - E
## Follicular phase - Contralateral - E2 - Lumen-Stroma surface - Follicular phase - Contralateral - E1
## Follicular phase - Contralateral - E2 - Muscle-Tunica surface - Follicular phase - Contralateral - E
## Follicular phase - Ipsilateral - E1 - Lumen-Stroma surface - Follicular phase - Contralateral - E1 -
## Follicular phase - Ipsilateral - E1 - Muscle-Tunica surface - Follicular phase - Contralateral - E1
## Follicular phase - Ipsilateral - E2 - Lumen-Stroma surface - Follicular phase - Contralateral - E1 -
## Follicular phase - Ipsilateral - E2 - Muscle-Tunica surface - Follicular phase - Contralateral - E1
## Luteal phase - Contralateral - E1 - Lumen-Stroma surface - Follicular phase - Contralateral - E1 - Lu
## Luteal phase - Contralateral - E1 - Muscle-Tunica surface - Follicular phase - Contralateral - E1 -
## Luteal phase - Contralateral - E2 - Lumen-Stroma surface - Follicular phase - Contralateral - E1 - Lu
## Luteal phase - Contralateral - E2 - Muscle-Tunica surface - Follicular phase - Contralateral - E1 -
## Luteal phase - Ipsilateral - E1 - Lumen-Stroma surface - Follicular phase - Contralateral - E1 - Lum
## Luteal phase - Ipsilateral - E1 - Muscle-Tunica surface - Follicular phase - Contralateral - E1 - Lu
## Luteal phase - Ipsilateral - E2 - Lumen-Stroma surface - Follicular phase - Contralateral - E1 - Lum
## Luteal phase - Ipsilateral - E2 - Muscle-Tunica surface - Follicular phase - Contralateral - E1 - Lu
## Follicular phase - Contralateral - E2 - Lumen-Stroma surface - Follicular phase - Contralateral - E1
## Follicular phase - Contralateral - E2 - Muscle-Tunica surface - Follicular phase - Contralateral - E
## Follicular phase - Ipsilateral - E1 - Lumen-Stroma surface - Follicular phase - Contralateral - E1 -
## Follicular phase - Ipsilateral - E1 - Muscle-Tunica surface - Follicular phase - Contralateral - E1
## Follicular phase - Ipsilateral - E2 - Lumen-Stroma surface - Follicular phase - Contralateral - E1 -
## Follicular phase - Ipsilateral - E2 - Muscle-Tunica surface - Follicular phase - Contralateral - E1
## Luteal phase - Contralateral - E1 - Lumen-Stroma surface - Follicular phase - Contralateral - E1 - Mu
## Luteal phase - Contralateral - E1 - Muscle-Tunica surface - Follicular phase - Contralateral - E1 -
## Luteal phase - Contralateral - E2 - Lumen-Stroma surface - Follicular phase - Contralateral - E1 - Mu
## Luteal phase - Contralateral - E2 - Muscle-Tunica surface - Follicular phase - Contralateral - E1 -
## Luteal phase - Ipsilateral - E1 - Lumen-Stroma surface - Follicular phase - Contralateral - E1 - Mus
```

```
## Luteal phase - Ipsilateral - E1 - Muscle-Tunica surface - Follicular phase - Contralateral - E1 - Mu
## Luteal phase - Ipsilateral - E2 - Lumen-Stroma surface - Follicular phase - Contralateral - E1 - Mus
## Luteal phase - Ipsilateral - E2 - Muscle-Tunica surface - Follicular phase - Contralateral - E1 - Mu
## Follicular phase - Contralateral - E2 - Muscle-Tunica surface - Follicular phase - Contralateral - E
## Follicular phase - Ipsilateral - E1 - Lumen-Stroma surface - Follicular phase - Contralateral - E2 -
## Follicular phase - Ipsilateral - E1 - Muscle-Tunica surface - Follicular phase - Contralateral - E2 -
## Follicular phase - Ipsilateral - E2 - Lumen-Stroma surface - Follicular phase - Contralateral - E2 -
## Follicular phase - Ipsilateral - E2 - Muscle-Tunica surface - Follicular phase - Contralateral - E2 -
## Luteal phase - Contralateral - E1 - Lumen-Stroma surface - Follicular phase - Contralateral - E2 - Lu
## Luteal phase - Contralateral - E1 - Muscle-Tunica surface - Follicular phase - Contralateral - E2 -
## Luteal phase - Contralateral - E2 - Lumen-Stroma surface - Follicular phase - Contralateral - E2 - Lu
## Luteal phase - Contralateral - E2 - Muscle-Tunica surface - Follicular phase - Contralateral - E2 -
## Luteal phase - Ipsilateral - E1 - Lumen-Stroma surface - Follicular phase - Contralateral - E2 - Lum
## Luteal phase - Ipsilateral - E1 - Muscle-Tunica surface - Follicular phase - Contralateral - E2 - Lu
## Luteal phase - Ipsilateral - E2 - Lumen-Stroma surface - Follicular phase - Contralateral - E2 - Lum
## Luteal phase - Ipsilateral - E2 - Muscle-Tunica surface - Follicular phase - Contralateral - E2 - Lu
## Follicular phase - Ipsilateral - E1 - Lumen-Stroma surface - Follicular phase - Contralateral - E2 -
## Follicular phase - Ipsilateral - E1 - Muscle-Tunica surface - Follicular phase - Contralateral - E2 -
## Follicular phase - Ipsilateral - E2 - Lumen-Stroma surface - Follicular phase - Contralateral - E2 -
## Follicular phase - Ipsilateral - E2 - Muscle-Tunica surface - Follicular phase - Contralateral - E2 -
## Luteal phase - Contralateral - E1 - Lumen-Stroma surface - Follicular phase - Contralateral - E2 - Mu
## Luteal phase - Contralateral - E1 - Muscle-Tunica surface - Follicular phase - Contralateral - E2 -
## Luteal phase - Contralateral - E2 - Lumen-Stroma surface - Follicular phase - Contralateral - E2 - Mu
## Luteal phase - Contralateral - E2 - Muscle-Tunica surface - Follicular phase - Contralateral - E2 -
## Luteal phase - Ipsilateral - E1 - Lumen-Stroma surface - Follicular phase - Contralateral - E2 - Mus
## Luteal phase - Ipsilateral - E1 - Muscle-Tunica surface - Follicular phase - Contralateral - E2 - Mu
## Luteal phase - Ipsilateral - E2 - Lumen-Stroma surface - Follicular phase - Contralateral - E2 - Mus
## Luteal phase - Ipsilateral - E2 - Muscle-Tunica surface - Follicular phase - Contralateral - E2 - Mu
## Follicular phase - Ipsilateral - E1 - Muscle-Tunica surface - Follicular phase - Ipsilateral - E1 -
## Follicular phase - Ipsilateral - E2 - Lumen-Stroma surface - Follicular phase - Ipsilateral - E1 - Lu
## Follicular phase - Ipsilateral - E2 - Muscle-Tunica surface - Follicular phase - Ipsilateral - E1 -
## Luteal phase - Contralateral - E1 - Lumen-Stroma surface - Follicular phase - Ipsilateral - E1 - Lum
## Luteal phase - Contralateral - E1 - Muscle-Tunica surface - Follicular phase - Ipsilateral - E1 - Lu
## Luteal phase - Contralateral - E2 - Lumen-Stroma surface - Follicular phase - Ipsilateral - E1 - Lum
## Luteal phase - Contralateral - E2 - Muscle-Tunica surface - Follicular phase - Ipsilateral - E1 - Lu
## Luteal phase - Ipsilateral - E1 - Lumen-Stroma surface - Follicular phase - Ipsilateral - E1 - Lumen-
## Luteal phase - Ipsilateral - E1 - Muscle-Tunica surface - Follicular phase - Ipsilateral - E1 - Lumen
## Luteal phase - Ipsilateral - E2 - Lumen-Stroma surface - Follicular phase - Ipsilateral - E1 - Lumen-
## Luteal phase - Ipsilateral - E2 - Muscle-Tunica surface - Follicular phase - Ipsilateral - E1 - Lumen
## Follicular phase - Ipsilateral - E2 - Lumen-Stroma surface - Follicular phase - Ipsilateral - E1 - Mu
## Follicular phase - Ipsilateral - E2 - Muscle-Tunica surface - Follicular phase - Ipsilateral - E1 -
## Luteal phase - Contralateral - E1 - Lumen-Stroma surface - Follicular phase - Ipsilateral - E1 - Mus
## Luteal phase - Contralateral - E1 - Muscle-Tunica surface - Follicular phase - Ipsilateral - E1 - Mu
## Luteal phase - Contralateral - E2 - Lumen-Stroma surface - Follicular phase - Ipsilateral - E1 - Mus
## Luteal phase - Contralateral - E2 - Muscle-Tunica surface - Follicular phase - Ipsilateral - E1 - Mu
## Luteal phase - Ipsilateral - E1 - Lumen-Stroma surface - Follicular phase - Ipsilateral - E1 - Muscle
## Luteal phase - Ipsilateral - E1 - Muscle-Tunica surface - Follicular phase - Ipsilateral - E1 - Muscl
## Luteal phase - Ipsilateral - E2 - Lumen-Stroma surface - Follicular phase - Ipsilateral - E1 - Muscle
## Luteal phase - Ipsilateral - E2 - Muscle-Tunica surface - Follicular phase - Ipsilateral - E1 - Muscl
## Follicular phase - Ipsilateral - E2 - Muscle-Tunica surface - Follicular phase - Ipsilateral - E2 -
## Luteal phase - Contralateral - E1 - Lumen-Stroma surface - Follicular phase - Ipsilateral - E2 - Lum
## Luteal phase - Contralateral - E1 - Muscle-Tunica surface - Follicular phase - Ipsilateral - E2 - Lu
## Luteal phase - Contralateral - E2 - Lumen-Stroma surface - Follicular phase - Ipsilateral - E2 - Lum
## Luteal phase - Contralateral - E2 - Muscle-Tunica surface - Follicular phase - Ipsilateral - E2 - Lu
```

```
## Luteal phase - Ipsilateral - E1 - Lumen-Stroma surface - Follicular phase - Ipsilateral - E2 - Lumen-
## Luteal phase - Ipsilateral - E1 - Muscle-Tunica surface - Follicular phase - Ipsilateral - E2 - Lumen
## Luteal phase - Ipsilateral - E2 - Lumen-Stroma surface - Follicular phase - Ipsilateral - E2 - Lumen-
## Luteal phase - Ipsilateral - E2 - Muscle-Tunica surface - Follicular phase - Ipsilateral - E2 - Lumen
## Luteal phase - Contralateral - E1 - Lumen-Stroma surface - Follicular phase - Ipsilateral - E2 - Musc
## Luteal phase - Contralateral - E1 - Muscle-Tunica surface - Follicular phase - Ipsilateral - E2 - Mus
## Luteal phase - Contralateral - E2 - Lumen-Stroma surface - Follicular phase - Ipsilateral - E2 - Musc
## Luteal phase - Contralateral - E2 - Muscle-Tunica surface - Follicular phase - Ipsilateral - E2 - Mus
## Luteal phase - Ipsilateral - E1 - Lumen-Stroma surface - Follicular phase - Ipsilateral - E2 - Muscle
## Luteal phase - Ipsilateral - E1 - Muscle-Tunica surface - Follicular phase - Ipsilateral - E2 - Muscl
## Luteal phase - Ipsilateral - E2 - Lumen-Stroma surface - Follicular phase - Ipsilateral - E2 - Muscle
## Luteal phase - Ipsilateral - E2 - Muscle-Tunica surface - Follicular phase - Ipsilateral - E2 - Muscl
## Luteal phase - Contralateral - E1 - Muscle-Tunica surface - Luteal phase - Contralateral - E1 - Lumen
## Luteal phase - Contralateral - E2 - Lumen-Stroma surface - Luteal phase - Contralateral - E1 - Lumen-
## Luteal phase - Contralateral - E2 - Muscle-Tunica surface - Luteal phase - Contralateral - E1 - Lumen
## Luteal phase - Ipsilateral - E1 - Lumen-Stroma surface - Luteal phase - Contralateral - E1 - Lumen-St
## Luteal phase - Ipsilateral - E1 - Muscle-Tunica surface - Luteal phase - Contralateral - E1 - Lumen-
## Luteal phase - Ipsilateral - E2 - Lumen-Stroma surface - Luteal phase - Contralateral - E1 - Lumen-St
## Luteal phase - Ipsilateral - E2 - Muscle-Tunica surface - Luteal phase - Contralateral - E1 - Lumen-
## Luteal phase - Contralateral - E2 - Lumen-Stroma surface - Luteal phase - Contralateral - E1 - Muscle
## Luteal phase - Contralateral - E2 - Muscle-Tunica surface - Luteal phase - Contralateral - E1 - Muscl
## Luteal phase - Ipsilateral - E1 - Lumen-Stroma surface - Luteal phase - Contralateral - E1 - Muscle-
## Luteal phase - Ipsilateral - E1 - Muscle-Tunica surface - Luteal phase - Contralateral - E1 - Muscle
## Luteal phase - Ipsilateral - E2 - Lumen-Stroma surface - Luteal phase - Contralateral - E1 - Muscle-
## Luteal phase - Ipsilateral - E2 - Muscle-Tunica surface - Luteal phase - Contralateral - E1 - Muscle
## Luteal phase - Contralateral - E2 - Muscle-Tunica surface - Luteal phase - Contralateral - E2 - Lumen
## Luteal phase - Ipsilateral - E1 - Lumen-Stroma surface - Luteal phase - Contralateral - E2 - Lumen-St
## Luteal phase - Ipsilateral - E1 - Muscle-Tunica surface - Luteal phase - Contralateral - E2 - Lumen-
## Luteal phase - Ipsilateral - E2 - Lumen-Stroma surface - Luteal phase - Contralateral - E2 - Lumen-St
## Luteal phase - Ipsilateral - E2 - Muscle-Tunica surface - Luteal phase - Contralateral - E2 - Lumen-
## Luteal phase - Ipsilateral - E1 - Lumen-Stroma surface - Luteal phase - Contralateral - E2 - Muscle-
## Luteal phase - Ipsilateral - E1 - Muscle-Tunica surface - Luteal phase - Contralateral - E2 - Muscle
## Luteal phase - Ipsilateral - E2 - Lumen-Stroma surface - Luteal phase - Contralateral - E2 - Muscle-
## Luteal phase - Ipsilateral - E2 - Muscle-Tunica surface - Luteal phase - Contralateral - E2 - Muscle
## Luteal phase - Ipsilateral - E1 - Muscle-Tunica surface - Luteal phase - Ipsilateral - E1 - Lumen-St
## Luteal phase - Ipsilateral - E2 - Lumen-Stroma surface - Luteal phase - Ipsilateral - E1 - Lumen-Str
## Luteal phase - Ipsilateral - E2 - Muscle-Tunica surface - Luteal phase - Ipsilateral - E1 - Lumen-St
## Luteal phase - Ipsilateral - E2 - Lumen-Stroma surface - Luteal phase - Ipsilateral - E1 - Muscle-Tu
## Luteal phase - Ipsilateral - E2 - Muscle-Tunica surface - Luteal phase - Ipsilateral - E1 - Muscle-T
## Luteal phase - Ipsilateral - E2 - Muscle-Tunica surface - Luteal phase - Ipsilateral - E2 - Lumen-St
##
## Follicular phase - Contralateral - E1 - Muscle-Tunica surface - Follicular phase - Contralateral - E
## Follicular phase - Contralateral - E2 - Lumen-Stroma surface - Follicular phase - Contralateral - E1
## Follicular phase - Contralateral - E2 - Muscle-Tunica surface - Follicular phase - Contralateral - E
## Follicular phase - Ipsilateral - E1 - Lumen-Stroma surface - Follicular phase - Contralateral - E1 -
## Follicular phase - Ipsilateral - E1 - Muscle-Tunica surface - Follicular phase - Contralateral - E1
## Follicular phase - Ipsilateral - E2 - Lumen-Stroma surface - Follicular phase - Contralateral - E1 -
## Follicular phase - Ipsilateral - E2 - Muscle-Tunica surface - Follicular phase - Contralateral - E1
## Luteal phase - Contralateral - E1 - Lumen-Stroma surface - Follicular phase - Contralateral - E1 - Lu
## Luteal phase - Contralateral - E1 - Muscle-Tunica surface - Follicular phase - Contralateral - E1 -
## Luteal phase - Contralateral - E2 - Lumen-Stroma surface - Follicular phase - Contralateral - E1 - Lu
## Luteal phase - Contralateral - E2 - Muscle-Tunica surface - Follicular phase - Contralateral - E1 -
## Luteal phase - Ipsilateral - E1 - Lumen-Stroma surface - Follicular phase - Contralateral - E1 - Lume
## Luteal phase - Ipsilateral - E1 - Muscle-Tunica surface - Follicular phase - Contralateral - E1 - Lu
```

```
## Luteal phase - Ipsilateral - E2 - Lumen-Stroma surface - Follicular phase - Contralateral - E1 - Lum
## Luteal phase - Ipsilateral - E2 - Muscle-Tunica surface - Follicular phase - Contralateral - E1 - Lu
## Follicular phase - Contralateral - E2 - Lumen-Stroma surface - Follicular phase - Contralateral - E1
## Follicular phase - Contralateral - E2 - Muscle-Tunica surface - Follicular phase - Contralateral - E
## Follicular phase - Ipsilateral - E1 - Lumen-Stroma surface - Follicular phase - Contralateral - E1 -
## Follicular phase - Ipsilateral - E1 - Muscle-Tunica surface - Follicular phase - Contralateral - E1
## Follicular phase - Ipsilateral - E2 - Lumen-Stroma surface - Follicular phase - Contralateral - E1 -
## Follicular phase - Ipsilateral - E2 - Muscle-Tunica surface - Follicular phase - Contralateral - E1
## Luteal phase - Contralateral - E1 - Lumen-Stroma surface - Follicular phase - Contralateral - E1 - Mu
## Luteal phase - Contralateral - E1 - Muscle-Tunica surface - Follicular phase - Contralateral - E1 -
## Luteal phase - Contralateral - E2 - Lumen-Stroma surface - Follicular phase - Contralateral - E1 - Mu
## Luteal phase - Contralateral - E2 - Muscle-Tunica surface - Follicular phase - Contralateral - E1 -
## Luteal phase - Ipsilateral - E1 - Lumen-Stroma surface - Follicular phase - Contralateral - E1 - Mus
## Luteal phase - Ipsilateral - E1 - Muscle-Tunica surface - Follicular phase - Contralateral - E1 - Mu
## Luteal phase - Ipsilateral - E2 - Lumen-Stroma surface - Follicular phase - Contralateral - E1 - Mus
## Luteal phase - Ipsilateral - E2 - Muscle-Tunica surface - Follicular phase - Contralateral - E1 - Mu
## Follicular phase - Contralateral - E2 - Muscle-Tunica surface - Follicular phase - Contralateral - E
## Follicular phase - Ipsilateral - E1 - Lumen-Stroma surface - Follicular phase - Contralateral - E2 -
## Follicular phase - Ipsilateral - E1 - Muscle-Tunica surface - Follicular phase - Contralateral - E2
## Follicular phase - Ipsilateral - E2 - Lumen-Stroma surface - Follicular phase - Contralateral - E2 -
## Follicular phase - Ipsilateral - E2 - Muscle-Tunica surface - Follicular phase - Contralateral - E2
## Luteal phase - Contralateral - E1 - Lumen-Stroma surface - Follicular phase - Contralateral - E2 - Lu
## Luteal phase - Contralateral - E1 - Muscle-Tunica surface - Follicular phase - Contralateral - E2 -
## Luteal phase - Contralateral - E2 - Lumen-Stroma surface - Follicular phase - Contralateral - E2 - Lu
## Luteal phase - Contralateral - E2 - Muscle-Tunica surface - Follicular phase - Contralateral - E2 -
## Luteal phase - Ipsilateral - E1 - Lumen-Stroma surface - Follicular phase - Contralateral - E2 - Lum
## Luteal phase - Ipsilateral - E1 - Muscle-Tunica surface - Follicular phase - Contralateral - E2 - Lu
## Luteal phase - Ipsilateral - E2 - Lumen-Stroma surface - Follicular phase - Contralateral - E2 - Lum
## Luteal phase - Ipsilateral - E2 - Muscle-Tunica surface - Follicular phase - Contralateral - E2 - Lu
## Follicular phase - Ipsilateral - E1 - Lumen-Stroma surface - Follicular phase - Contralateral - E2 -
## Follicular phase - Ipsilateral - E1 - Muscle-Tunica surface - Follicular phase - Contralateral - E2
## Follicular phase - Ipsilateral - E2 - Lumen-Stroma surface - Follicular phase - Contralateral - E2 -
## Follicular phase - Ipsilateral - E2 - Muscle-Tunica surface - Follicular phase - Contralateral - E2
## Luteal phase - Contralateral - E1 - Lumen-Stroma surface - Follicular phase - Contralateral - E2 - Mu
## Luteal phase - Contralateral - E1 - Muscle-Tunica surface - Follicular phase - Contralateral - E2 -
## Luteal phase - Contralateral - E2 - Lumen-Stroma surface - Follicular phase - Contralateral - E2 - Mu
## Luteal phase - Contralateral - E2 - Muscle-Tunica surface - Follicular phase - Contralateral - E2 -
## Luteal phase - Ipsilateral - E1 - Lumen-Stroma surface - Follicular phase - Contralateral - E2 - Mus
## Luteal phase - Ipsilateral - E1 - Muscle-Tunica surface - Follicular phase - Contralateral - E2 - Mu
## Luteal phase - Ipsilateral - E2 - Lumen-Stroma surface - Follicular phase - Contralateral - E2 - Mus
## Luteal phase - Ipsilateral - E2 - Muscle-Tunica surface - Follicular phase - Contralateral - E2 - Mu
## Follicular phase - Ipsilateral - E1 - Muscle-Tunica surface - Follicular phase - Ipsilateral - E1 -
## Follicular phase - Ipsilateral - E2 - Lumen-Stroma surface - Follicular phase - Ipsilateral - E1 - Lu
## Follicular phase - Ipsilateral - E2 - Muscle-Tunica surface - Follicular phase - Ipsilateral - E1 -
## Luteal phase - Contralateral - E1 - Lumen-Stroma surface - Follicular phase - Ipsilateral - E1 - Lum
## Luteal phase - Contralateral - E1 - Muscle-Tunica surface - Follicular phase - Ipsilateral - E1 - Lu
## Luteal phase - Contralateral - E2 - Lumen-Stroma surface - Follicular phase - Ipsilateral - E1 - Lum
## Luteal phase - Contralateral - E2 - Muscle-Tunica surface - Follicular phase - Ipsilateral - E1 - Lu
## Luteal phase - Ipsilateral - E1 - Lumen-Stroma surface - Follicular phase - Ipsilateral - E1 - Lumen
## Luteal phase - Ipsilateral - E1 - Muscle-Tunica surface - Follicular phase - Ipsilateral - E1 - Lume
## Luteal phase - Ipsilateral - E2 - Lumen-Stroma surface - Follicular phase - Ipsilateral - E1 - Lumen
## Luteal phase - Ipsilateral - E2 - Muscle-Tunica surface - Follicular phase - Ipsilateral - E1 - Lume
## Follicular phase - Ipsilateral - E2 - Lumen-Stroma surface - Follicular phase - Ipsilateral - E1 - Mu
## Follicular phase - Ipsilateral - E2 - Muscle-Tunica surface - Follicular phase - Ipsilateral - E1 -
```

```
## Luteal phase - Contralateral - E1 - Lumen-Stroma surface - Follicular phase - Ipsilateral - E1 - Musc
## Luteal phase - Contralateral - E1 - Muscle-Tunica surface - Follicular phase - Ipsilateral - E1 - Mu
## Luteal phase - Contralateral - E2 - Lumen-Stroma surface - Follicular phase - Ipsilateral - E1 - Musc
## Luteal phase - Contralateral - E2 - Muscle-Tunica surface - Follicular phase - Ipsilateral - E1 - Mu
## Luteal phase - Ipsilateral - E1 - Lumen-Stroma surface - Follicular phase - Ipsilateral - E1 - Muscle
## Luteal phase - Ipsilateral - E1 - Muscle-Tunica surface - Follicular phase - Ipsilateral - E1 - Muscl
## Luteal phase - Ipsilateral - E2 - Lumen-Stroma surface - Follicular phase - Ipsilateral - E1 - Muscle
## Luteal phase - Ipsilateral - E2 - Muscle-Tunica surface - Follicular phase - Ipsilateral - E1 - Muscl
## Follicular phase - Ipsilateral - E2 - Muscle-Tunica surface - Follicular phase - Ipsilateral - E2 - 
## Luteal phase - Contralateral - E1 - Lumen-Stroma surface - Follicular phase - Ipsilateral - E2 - Lume
## Luteal phase - Contralateral - E1 - Muscle-Tunica surface - Follicular phase - Ipsilateral - E2 - Lu
## Luteal phase - Contralateral - E2 - Lumen-Stroma surface - Follicular phase - Ipsilateral - E2 - Lume
## Luteal phase - Contralateral - E2 - Muscle-Tunica surface - Follicular phase - Ipsilateral - E2 - Lu
## Luteal phase - Ipsilateral - E1 - Lumen-Stroma surface - Follicular phase - Ipsilateral - E2 - Lumen-
## Luteal phase - Ipsilateral - E1 - Muscle-Tunica surface - Follicular phase - Ipsilateral - E2 - Lumen
## Luteal phase - Ipsilateral - E2 - Lumen-Stroma surface - Follicular phase - Ipsilateral - E2 - Lumen-
## Luteal phase - Ipsilateral - E2 - Muscle-Tunica surface - Follicular phase - Ipsilateral - E2 - Lumen
## Luteal phase - Contralateral - E1 - Lumen-Stroma surface - Follicular phase - Ipsilateral - E2 - Musc
## Luteal phase - Contralateral - E1 - Muscle-Tunica surface - Follicular phase - Ipsilateral - E2 - Mu
## Luteal phase - Contralateral - E2 - Lumen-Stroma surface - Follicular phase - Ipsilateral - E2 - Musc
## Luteal phase - Contralateral - E2 - Muscle-Tunica surface - Follicular phase - Ipsilateral - E2 - Mu
## Luteal phase - Ipsilateral - E1 - Lumen-Stroma surface - Follicular phase - Ipsilateral - E2 - Muscle
## Luteal phase - Ipsilateral - E1 - Muscle-Tunica surface - Follicular phase - Ipsilateral - E2 - Muscl
## Luteal phase - Ipsilateral - E2 - Lumen-Stroma surface - Follicular phase - Ipsilateral - E2 - Muscle
## Luteal phase - Ipsilateral - E2 - Muscle-Tunica surface - Follicular phase - Ipsilateral - E2 - Muscl
## Luteal phase - Contralateral - E1 - Muscle-Tunica surface - Luteal phase - Contralateral - E1 - Lumen
## Luteal phase - Contralateral - E2 - Lumen-Stroma surface - Luteal phase - Contralateral - E1 - Lumen-
## Luteal phase - Contralateral - E2 - Muscle-Tunica surface - Luteal phase - Contralateral - E1 - Lumen
## Luteal phase - Ipsilateral - E1 - Lumen-Stroma surface - Luteal phase - Contralateral - E1 - Lumen-St
## Luteal phase - Ipsilateral - E1 - Muscle-Tunica surface - Luteal phase - Contralateral - E1 - Lumen-S
## Luteal phase - Ipsilateral - E2 - Lumen-Stroma surface - Luteal phase - Contralateral - E1 - Lumen-St
## Luteal phase - Ipsilateral - E2 - Muscle-Tunica surface - Luteal phase - Contralateral - E1 - Lumen-S
## Luteal phase - Contralateral - E2 - Lumen-Stroma surface - Luteal phase - Contralateral - E1 - Muscle
## Luteal phase - Contralateral - E2 - Muscle-Tunica surface - Luteal phase - Contralateral - E1 - Muscl
## Luteal phase - Ipsilateral - E1 - Lumen-Stroma surface - Luteal phase - Contralateral - E1 - Muscle-T
## Luteal phase - Ipsilateral - E1 - Muscle-Tunica surface - Luteal phase - Contralateral - E1 - Muscle-
## Luteal phase - Ipsilateral - E2 - Lumen-Stroma surface - Luteal phase - Contralateral - E1 - Muscle-T
## Luteal phase - Ipsilateral - E2 - Muscle-Tunica surface - Luteal phase - Contralateral - E1 - Muscle-
## Luteal phase - Contralateral - E2 - Muscle-Tunica surface - Luteal phase - Contralateral - E2 - Lumen
## Luteal phase - Ipsilateral - E1 - Lumen-Stroma surface - Luteal phase - Contralateral - E2 - Lumen-St
## Luteal phase - Ipsilateral - E1 - Muscle-Tunica surface - Luteal phase - Contralateral - E2 - Lumen-S
## Luteal phase - Ipsilateral - E2 - Lumen-Stroma surface - Luteal phase - Contralateral - E2 - Lumen-St
## Luteal phase - Ipsilateral - E2 - Muscle-Tunica surface - Luteal phase - Contralateral - E2 - Lumen-S
## Luteal phase - Ipsilateral - E1 - Lumen-Stroma surface - Luteal phase - Contralateral - E2 - Muscle-T
## Luteal phase - Ipsilateral - E1 - Muscle-Tunica surface - Luteal phase - Contralateral - E2 - Muscle-
## Luteal phase - Ipsilateral - E2 - Lumen-Stroma surface - Luteal phase - Contralateral - E2 - Muscle-T
## Luteal phase - Ipsilateral - E2 - Muscle-Tunica surface - Luteal phase - Contralateral - E2 - Muscle-
## Luteal phase - Ipsilateral - E1 - Muscle-Tunica surface - Luteal phase - Ipsilateral - E1 - Lumen-Str
## Luteal phase - Ipsilateral - E2 - Lumen-Stroma surface - Luteal phase - Ipsilateral - E1 - Lumen-Stro
## Luteal phase - Ipsilateral - E2 - Muscle-Tunica surface - Luteal phase - Ipsilateral - E1 - Lumen-Str
## Luteal phase - Ipsilateral - E2 - Lumen-Stroma surface - Luteal phase - Ipsilateral - E1 - Muscle-Tun
## Luteal phase - Ipsilateral - E2 - Muscle-Tunica surface - Luteal phase - Ipsilateral - E1 - Muscle-Tu
## Luteal phase - Ipsilateral - E2 - Muscle-Tunica surface - Luteal phase - Ipsilateral - E2 - Lumen-Str
##
```

```
## Follicular phase - Contralateral - E1 - Muscle-Tunica surface - Follicular phase - Contralateral - E
## Follicular phase - Contralateral - E2 - Lumen-Stroma surface - Follicular phase - Contralateral - E1
## Follicular phase - Contralateral - E2 - Muscle-Tunica surface - Follicular phase - Contralateral - E
## Follicular phase - Ipsilateral - E1 - Lumen-Stroma surface - Follicular phase - Contralateral - E1 -
## Follicular phase - Ipsilateral - E1 - Muscle-Tunica surface - Follicular phase - Contralateral - E1 -
## Follicular phase - Ipsilateral - E2 - Lumen-Stroma surface - Follicular phase - Contralateral - E1 -
## Follicular phase - Ipsilateral - E2 - Muscle-Tunica surface - Follicular phase - Contralateral - E1 -
## Luteal phase - Contralateral - E1 - Lumen-Stroma surface - Follicular phase - Contralateral - E1 - Lu
## Luteal phase - Contralateral - E1 - Muscle-Tunica surface - Follicular phase - Contralateral - E1 -
## Luteal phase - Contralateral - E2 - Lumen-Stroma surface - Follicular phase - Contralateral - E1 - Lu
## Luteal phase - Contralateral - E2 - Muscle-Tunica surface - Follicular phase - Contralateral - E1 -
## Luteal phase - Ipsilateral - E1 - Lumen-Stroma surface - Follicular phase - Contralateral - E1 - Lume
## Luteal phase - Ipsilateral - E1 - Muscle-Tunica surface - Follicular phase - Contralateral - E1 - Lum
## Luteal phase - Ipsilateral - E2 - Lumen-Stroma surface - Follicular phase - Contralateral - E1 - Lume
## Luteal phase - Ipsilateral - E2 - Muscle-Tunica surface - Follicular phase - Contralateral - E1 - Lum
## Follicular phase - Contralateral - E2 - Lumen-Stroma surface - Follicular phase - Contralateral - E1
## Follicular phase - Contralateral - E2 - Muscle-Tunica surface - Follicular phase - Contralateral - E
## Follicular phase - Ipsilateral - E1 - Lumen-Stroma surface - Follicular phase - Contralateral - E1 -
## Follicular phase - Ipsilateral - E1 - Muscle-Tunica surface - Follicular phase - Contralateral - E1 -
## Follicular phase - Ipsilateral - E2 - Lumen-Stroma surface - Follicular phase - Contralateral - E1 -
## Follicular phase - Ipsilateral - E2 - Muscle-Tunica surface - Follicular phase - Contralateral - E1 -
## Luteal phase - Contralateral - E1 - Lumen-Stroma surface - Follicular phase - Contralateral - E1 - Mu
## Luteal phase - Contralateral - E1 - Muscle-Tunica surface - Follicular phase - Contralateral - E1 -
## Luteal phase - Contralateral - E2 - Lumen-Stroma surface - Follicular phase - Contralateral - E1 - Mu
## Luteal phase - Contralateral - E2 - Muscle-Tunica surface - Follicular phase - Contralateral - E1 -
## Luteal phase - Ipsilateral - E1 - Lumen-Stroma surface - Follicular phase - Contralateral - E1 - Musc
## Luteal phase - Ipsilateral - E1 - Muscle-Tunica surface - Follicular phase - Contralateral - E1 - Mus
## Luteal phase - Ipsilateral - E2 - Lumen-Stroma surface - Follicular phase - Contralateral - E1 - Musc
## Luteal phase - Ipsilateral - E2 - Muscle-Tunica surface - Follicular phase - Contralateral - E1 - Mus
## Follicular phase - Contralateral - E2 - Muscle-Tunica surface - Follicular phase - Contralateral - E
## Follicular phase - Ipsilateral - E1 - Lumen-Stroma surface - Follicular phase - Contralateral - E2 -
## Follicular phase - Ipsilateral - E1 - Muscle-Tunica surface - Follicular phase - Contralateral - E2 -
## Follicular phase - Ipsilateral - E2 - Lumen-Stroma surface - Follicular phase - Contralateral - E2 -
## Follicular phase - Ipsilateral - E2 - Muscle-Tunica surface - Follicular phase - Contralateral - E2 -
## Luteal phase - Contralateral - E1 - Lumen-Stroma surface - Follicular phase - Contralateral - E2 - Lu
## Luteal phase - Contralateral - E1 - Muscle-Tunica surface - Follicular phase - Contralateral - E2 -
## Luteal phase - Contralateral - E2 - Lumen-Stroma surface - Follicular phase - Contralateral - E2 - Lu
## Luteal phase - Contralateral - E2 - Muscle-Tunica surface - Follicular phase - Contralateral - E2 -
## Luteal phase - Ipsilateral - E1 - Lumen-Stroma surface - Follicular phase - Contralateral - E2 - Lume
## Luteal phase - Ipsilateral - E1 - Muscle-Tunica surface - Follicular phase - Contralateral - E2 - Lum
## Luteal phase - Ipsilateral - E2 - Lumen-Stroma surface - Follicular phase - Contralateral - E2 - Lume
## Luteal phase - Ipsilateral - E2 - Muscle-Tunica surface - Follicular phase - Contralateral - E2 - Lum
## Follicular phase - Ipsilateral - E1 - Lumen-Stroma surface - Follicular phase - Contralateral - E2 -
## Follicular phase - Ipsilateral - E1 - Muscle-Tunica surface - Follicular phase - Contralateral - E2 -
## Follicular phase - Ipsilateral - E2 - Lumen-Stroma surface - Follicular phase - Contralateral - E2 -
## Follicular phase - Ipsilateral - E2 - Muscle-Tunica surface - Follicular phase - Contralateral - E2 -
## Luteal phase - Contralateral - E1 - Lumen-Stroma surface - Follicular phase - Contralateral - E2 - Mu
## Luteal phase - Contralateral - E1 - Muscle-Tunica surface - Follicular phase - Contralateral - E2 -
## Luteal phase - Contralateral - E2 - Lumen-Stroma surface - Follicular phase - Contralateral - E2 - Mu
## Luteal phase - Contralateral - E2 - Muscle-Tunica surface - Follicular phase - Contralateral - E2 -
## Luteal phase - Ipsilateral - E1 - Lumen-Stroma surface - Follicular phase - Contralateral - E2 - Musc
## Luteal phase - Ipsilateral - E1 - Muscle-Tunica surface - Follicular phase - Contralateral - E2 - Mus
## Luteal phase - Ipsilateral - E2 - Lumen-Stroma surface - Follicular phase - Contralateral - E2 - Musc
## Luteal phase - Ipsilateral - E2 - Muscle-Tunica surface - Follicular phase - Contralateral - E2 - Mus
```

```
## Follicular phase - Ipsilateral - E1 - Muscle-Tunica surface - Follicular phase - Ipsilateral - E1 -
## Follicular phase - Ipsilateral - E2 - Lumen-Stroma surface - Follicular phase - Ipsilateral - E1 - Lu
## Follicular phase - Ipsilateral - E2 - Muscle-Tunica surface - Follicular phase - Ipsilateral - E1 -
## Luteal phase - Contralateral - E1 - Lumen-Stroma surface - Follicular phase - Ipsilateral - E1 - Lume
## Luteal phase - Contralateral - E1 - Muscle-Tunica surface - Follicular phase - Ipsilateral - E1 - Lu
## Luteal phase - Contralateral - E2 - Lumen-Stroma surface - Follicular phase - Ipsilateral - E1 - Lume
## Luteal phase - Contralateral - E2 - Muscle-Tunica surface - Follicular phase - Ipsilateral - E1 - Lu
## Luteal phase - Ipsilateral - E1 - Lumen-Stroma surface - Follicular phase - Ipsilateral - E1 - Lumen
## Luteal phase - Ipsilateral - E1 - Muscle-Tunica surface - Follicular phase - Ipsilateral - E1 - Lume
## Luteal phase - Ipsilateral - E2 - Lumen-Stroma surface - Follicular phase - Ipsilateral - E1 - Lumen
## Luteal phase - Ipsilateral - E2 - Muscle-Tunica surface - Follicular phase - Ipsilateral - E1 - Lume
## Follicular phase - Ipsilateral - E2 - Lumen-Stroma surface - Follicular phase - Ipsilateral - E1 - Mu
## Follicular phase - Ipsilateral - E2 - Muscle-Tunica surface - Follicular phase - Ipsilateral - E1 -
## Luteal phase - Contralateral - E1 - Lumen-Stroma surface - Follicular phase - Ipsilateral - E1 - Musc
## Luteal phase - Contralateral - E1 - Muscle-Tunica surface - Follicular phase - Ipsilateral - E1 - Mu
## Luteal phase - Contralateral - E2 - Lumen-Stroma surface - Follicular phase - Ipsilateral - E1 - Musc
## Luteal phase - Contralateral - E2 - Muscle-Tunica surface - Follicular phase - Ipsilateral - E1 - Mu
## Luteal phase - Ipsilateral - E1 - Lumen-Stroma surface - Follicular phase - Ipsilateral - E1 - Muscle
## Luteal phase - Ipsilateral - E1 - Muscle-Tunica surface - Follicular phase - Ipsilateral - E1 - Muscl
## Luteal phase - Ipsilateral - E2 - Lumen-Stroma surface - Follicular phase - Ipsilateral - E1 - Muscle
## Luteal phase - Ipsilateral - E2 - Muscle-Tunica surface - Follicular phase - Ipsilateral - E1 - Muscl
## Follicular phase - Ipsilateral - E2 - Muscle-Tunica surface - Follicular phase - Ipsilateral - E2 -
## Luteal phase - Contralateral - E1 - Lumen-Stroma surface - Follicular phase - Ipsilateral - E2 - Lume
## Luteal phase - Contralateral - E1 - Muscle-Tunica surface - Follicular phase - Ipsilateral - E2 - Lu
## Luteal phase - Contralateral - E2 - Lumen-Stroma surface - Follicular phase - Ipsilateral - E2 - Lume
## Luteal phase - Contralateral - E2 - Muscle-Tunica surface - Follicular phase - Ipsilateral - E2 - Lu
## Luteal phase - Ipsilateral - E1 - Lumen-Stroma surface - Follicular phase - Ipsilateral - E2 - Lumen
## Luteal phase - Ipsilateral - E1 - Muscle-Tunica surface - Follicular phase - Ipsilateral - E2 - Lume
## Luteal phase - Ipsilateral - E2 - Lumen-Stroma surface - Follicular phase - Ipsilateral - E2 - Lumen
## Luteal phase - Ipsilateral - E2 - Muscle-Tunica surface - Follicular phase - Ipsilateral - E2 - Lume
## Luteal phase - Contralateral - E1 - Lumen-Stroma surface - Follicular phase - Ipsilateral - E2 - Musc
## Luteal phase - Contralateral - E1 - Muscle-Tunica surface - Follicular phase - Ipsilateral - E2 - Mu
## Luteal phase - Contralateral - E2 - Lumen-Stroma surface - Follicular phase - Ipsilateral - E2 - Musc
## Luteal phase - Contralateral - E2 - Muscle-Tunica surface - Follicular phase - Ipsilateral - E2 - Mu
## Luteal phase - Ipsilateral - E1 - Lumen-Stroma surface - Follicular phase - Ipsilateral - E2 - Muscle
## Luteal phase - Ipsilateral - E1 - Muscle-Tunica surface - Follicular phase - Ipsilateral - E2 - Muscl
## Luteal phase - Ipsilateral - E2 - Lumen-Stroma surface - Follicular phase - Ipsilateral - E2 - Muscle
## Luteal phase - Ipsilateral - E2 - Muscle-Tunica surface - Follicular phase - Ipsilateral - E2 - Muscl
## Luteal phase - Contralateral - E1 - Muscle-Tunica surface - Luteal phase - Contralateral - E1 - Lume
## Luteal phase - Contralateral - E2 - Lumen-Stroma surface - Luteal phase - Contralateral - E1 - Lumen
## Luteal phase - Contralateral - E2 - Muscle-Tunica surface - Luteal phase - Contralateral - E1 - Lume
## Luteal phase - Ipsilateral - E1 - Lumen-Stroma surface - Luteal phase - Contralateral - E1 - Lumen-S
## Luteal phase - Ipsilateral - E1 - Muscle-Tunica surface - Luteal phase - Contralateral - E1 - Lumen-
## Luteal phase - Ipsilateral - E2 - Lumen-Stroma surface - Luteal phase - Contralateral - E1 - Lumen-S
## Luteal phase - Ipsilateral - E2 - Muscle-Tunica surface - Luteal phase - Contralateral - E1 - Lumen-
## Luteal phase - Contralateral - E2 - Lumen-Stroma surface - Luteal phase - Contralateral - E1 - Muscle
## Luteal phase - Contralateral - E2 - Muscle-Tunica surface - Luteal phase - Contralateral - E1 - Muscl
## Luteal phase - Ipsilateral - E1 - Lumen-Stroma surface - Luteal phase - Contralateral - E1 - Muscle-
## Luteal phase - Ipsilateral - E1 - Muscle-Tunica surface - Luteal phase - Contralateral - E1 - Muscle
## Luteal phase - Ipsilateral - E2 - Lumen-Stroma surface - Luteal phase - Contralateral - E1 - Muscle-
## Luteal phase - Ipsilateral - E2 - Muscle-Tunica surface - Luteal phase - Contralateral - E1 - Muscle
## Luteal phase - Contralateral - E2 - Muscle-Tunica surface - Luteal phase - Contralateral - E2 - Lume
## Luteal phase - Ipsilateral - E1 - Lumen-Stroma surface - Luteal phase - Contralateral - E2 - Lumen-S
## Luteal phase - Ipsilateral - E1 - Muscle-Tunica surface - Luteal phase - Contralateral - E2 - Lumen-
```

```
## Luteal phase - Ipsilateral - E2 - Lumen-Stroma surface - Luteal phase - Contralateral - E2 - Lumen-St
## Luteal phase - Ipsilateral - E2 - Muscle-Tunica surface - Luteal phase - Contralateral - E2 - Lumen-S
## Luteal phase - Ipsilateral - E1 - Lumen-Stroma surface - Luteal phase - Contralateral - E2 - Muscle-T
## Luteal phase - Ipsilateral - E1 - Muscle-Tunica surface - Luteal phase - Contralateral - E2 - Muscle-
## Luteal phase - Ipsilateral - E2 - Lumen-Stroma surface - Luteal phase - Contralateral - E2 - Muscle-T
## Luteal phase - Ipsilateral - E2 - Muscle-Tunica surface - Luteal phase - Contralateral - E2 - Muscle-
## Luteal phase - Ipsilateral - E1 - Muscle-Tunica surface - Luteal phase - Ipsilateral - E1 - Lumen-Str
## Luteal phase - Ipsilateral - E2 - Lumen-Stroma surface - Luteal phase - Ipsilateral - E1 - Lumen-Stro
## Luteal phase - Ipsilateral - E2 - Muscle-Tunica surface - Luteal phase - Ipsilateral - E1 - Lumen-Str
## Luteal phase - Ipsilateral - E2 - Lumen-Stroma surface - Luteal phase - Ipsilateral - E1 - Muscle-Tun
## Luteal phase - Ipsilateral - E2 - Muscle-Tunica surface - Luteal phase - Ipsilateral - E1 - Muscle-Tu
## Luteal phase - Ipsilateral - E2 - Muscle-Tunica surface - Luteal phase - Ipsilateral - E2 - Lumen-Str
## ---
## Signif. codes:  0 '***' 0.001 '**' 0.01 '*' 0.05 '.' 0.1 ' ' 1
## (Adjusted p values reported -- single-step method)
```

```
# Plot Isthmus

#data input
data <- read.csv("Oviduct_rheo.csv")

# Subset the data based on X_position_group
data_ampulla <- subset(data, Tissue == "Ampulla")
data_isthmus <- subset(data, Tissue == "Isthmus")

png(filename="E1_E2_isthmus2.png",
    width = 9.86, height = 6.86, units = "in",
    res = 600)

par(mgp=c(1.5,0.5,0),
    mar = c(4, #bottom
            4, #left
            3, #top
            2),
    cex.main = 0.8,
    family = "sans" #right
)

# Create a new column to define the X_position group
data_isthmus$X_position_group <- ifelse(data_isthmus$X_position <= 400, "Lumen-Stroma surface", "Muscle-

# Calculate means and standard deviations
data_summary <- data_isthmus %>%
  group_by(Module, Estrus_cycle, Position, Hz, X_position_group) %>%
  summarise(mean_Indentation = mean(Indentation, na.rm = TRUE),
            sd_Indentation = sd(Indentation, na.rm = TRUE))
```

```
## `summarise()` has grouped output by 'Module', 'Estrus_cycle', 'Position', 'Hz'.
## You can override using the `.groups` argument.
```

```
# Create a new variable for line type
data_summary <- data_summary %>%
  mutate(LineType = ifelse(Module == "E1", "E1", "E2"))

# Define fill colors for each factor level
```

```
fill_colors <- c("Follicular phase Contralateral" = viridis_pal(option = "D")(1),
                 "Follicular phase Ipsilateral" = alpha(viridis_pal(option = "D")(1), 0.2),
                 "Luteal phase Contralateral" = "#238A8DFF",
                 "Luteal phase Ipsilateral" = alpha("#238A8DFF", 0.2))

# Create the line plot with facets for X_position_group
ggplot(data_summary, aes(x = Hz, y = mean_Indentation, group = interaction(Module, Estrus_cycle, Positi
                         color = paste(Estrus_cycle, Position), linetype = LineType)) +
  geom_line(size = 1.5) +
  geom_point(size = 1) +
  scale_color_manual(values = fill_colors) +
  scale_linetype_manual(values = c("E1" = "solid", "E2" = "dashed"),
                        labels = c("E1" = "E'", "E2" = "E\"")) +
  scale_x_continuous(breaks = c(1, 5, 10, 20)) +
  scale_y_continuous(labels = scientific_format(), limits = c(0, 30000)) +
  labs(y = expression(paste("E'/E'' (Pa)")), x = "Frequency (Hz)", color = "Estrus Cycle + Position") +
  theme_minimal() +
  theme(panel.grid.major = element_blank(),
        panel.grid.minor = element_blank(),
        panel.border = element_rect(colour = "black", fill = NA, size = 1),
        axis.text = element_text(size = 12)) +
  facet_wrap(~ X_position_group, nrow = 1)

dev.off()
```

```
## pdf
##   2
```

```
# Stats Isthmus

df <- data_isthmus

# Create a factor variable for the comparisons
df$Comparison <- factor(paste(df$Estrus_cycle, df$Position, df$Module, df$X_position_group, sep = " - "

# Perform pairwise comparisons using Tukey's method
model <- lm(Indentation ~ Comparison, data = df)
comp <- glht(model, linfct = mcp(Comparison = "Tukey"))

# Summarize the results
summary(comp)
```

```
## Warning in RET$pfunction("adjusted", ...): Completion with error > abseps

## Warning in RET$pfunction("adjusted", ...): Completion with error > abseps

## Warning in RET$pfunction("adjusted", ...): Completion with error > abseps

## Warning in RET$pfunction("adjusted", ...): Completion with error > abseps

## Warning in RET$pfunction("adjusted", ...): Completion with error > abseps

## Warning in RET$pfunction("adjusted", ...): Completion with error > abseps

## Warning in RET$pfunction("adjusted", ...): Completion with error > abseps
```

```
## Warning in RET$pfunction("adjusted", ...): Completion with error > abseps

## Warning in RET$pfunction("adjusted", ...): Completion with error > abseps

## Warning in RET$pfunction("adjusted", ...): Completion with error > abseps

## Warning in RET$pfunction("adjusted", ...): Completion with error > abseps

## Warning in RET$pfunction("adjusted", ...): Completion with error > abseps

## Warning in RET$pfunction("adjusted", ...): Completion with error > abseps

## Warning in RET$pfunction("adjusted", ...): Completion with error > abseps

## Warning in RET$pfunction("adjusted", ...): Completion with error > abseps

## Warning in RET$pfunction("adjusted", ...): Completion with error > abseps

## Warning in RET$pfunction("adjusted", ...): Completion with error > abseps

## Warning in RET$pfunction("adjusted", ...): Completion with error > abseps

## Warning in RET$pfunction("adjusted", ...): Completion with error > abseps

## Warning in RET$pfunction("adjusted", ...): Completion with error > abseps

## Warning in RET$pfunction("adjusted", ...): Completion with error > abseps

## Warning in RET$pfunction("adjusted", ...): Completion with error > abseps

## Warning in RET$pfunction("adjusted", ...): Completion with error > abseps

## Warning in RET$pfunction("adjusted", ...): Completion with error > abseps

## Warning in RET$pfunction("adjusted", ...): Completion with error > abseps

## Warning in RET$pfunction("adjusted", ...): Completion with error > abseps

## Warning in RET$pfunction("adjusted", ...): Completion with error > abseps

## Warning in RET$pfunction("adjusted", ...): Completion with error > abseps

## Warning in RET$pfunction("adjusted", ...): Completion with error > abseps

## Warning in RET$pfunction("adjusted", ...): Completion with error > abseps

## Warning in RET$pfunction("adjusted", ...): Completion with error > abseps

## Warning in RET$pfunction("adjusted", ...): Completion with error > abseps

## Warning in RET$pfunction("adjusted", ...): Completion with error > abseps

## Warning in RET$pfunction("adjusted", ...): Completion with error > abseps
```

```
## Warning in RET$pfunction("adjusted", ...): Completion with error > abseps

## Warning in RET$pfunction("adjusted", ...): Completion with error > abseps
##
##   Simultaneous Tests for General Linear Hypotheses
##
## Multiple Comparisons of Means: Tukey Contrasts
##
##
## Fit: lm(formula = Indentation ~ Comparison, data = df)
##
## Linear Hypotheses:
##
## Follicular phase - Contralateral - E1 - Muscle-Tunica surface - Follicular phase - Contralateral - E
## Follicular phase - Contralateral - E2 - Lumen-Stroma surface - Follicular phase - Contralateral - E1
## Follicular phase - Contralateral - E2 - Muscle-Tunica surface - Follicular phase - Contralateral - E
## Follicular phase - Ipsilateral - E1 - Lumen-Stroma surface - Follicular phase - Contralateral - E1 -
## Follicular phase - Ipsilateral - E1 - Muscle-Tunica surface - Follicular phase - Contralateral - E1
## Follicular phase - Ipsilateral - E2 - Lumen-Stroma surface - Follicular phase - Contralateral - E1 -
## Follicular phase - Ipsilateral - E2 - Muscle-Tunica surface - Follicular phase - Contralateral - E1
## Luteal phase - Contralateral - E1 - Lumen-Stroma surface - Follicular phase - Contralateral - E1 - Lu
## Luteal phase - Contralateral - E1 - Muscle-Tunica surface - Follicular phase - Contralateral - E1 -
## Luteal phase - Contralateral - E2 - Lumen-Stroma surface - Follicular phase - Contralateral - E1 - Lu
## Luteal phase - Contralateral - E2 - Muscle-Tunica surface - Follicular phase - Contralateral - E1 -
## Luteal phase - Ipsilateral - E1 - Lumen-Stroma surface - Follicular phase - Contralateral - E1 - Lum
## Luteal phase - Ipsilateral - E1 - Muscle-Tunica surface - Follicular phase - Contralateral - E1 - Lu
## Luteal phase - Ipsilateral - E2 - Lumen-Stroma surface - Follicular phase - Contralateral - E1 - Lum
## Luteal phase - Ipsilateral - E2 - Muscle-Tunica surface - Follicular phase - Contralateral - E1 - Lu
## Follicular phase - Contralateral - E2 - Lumen-Stroma surface - Follicular phase - Contralateral - E1
## Follicular phase - Contralateral - E2 - Muscle-Tunica surface - Follicular phase - Contralateral - E
## Follicular phase - Ipsilateral - E1 - Lumen-Stroma surface - Follicular phase - Contralateral - E1 -
## Follicular phase - Ipsilateral - E1 - Muscle-Tunica surface - Follicular phase - Contralateral - E1
## Follicular phase - Ipsilateral - E2 - Lumen-Stroma surface - Follicular phase - Contralateral - E1 -
## Follicular phase - Ipsilateral - E2 - Muscle-Tunica surface - Follicular phase - Contralateral - E1
## Luteal phase - Contralateral - E1 - Lumen-Stroma surface - Follicular phase - Contralateral - E1 - Mu
## Luteal phase - Contralateral - E1 - Muscle-Tunica surface - Follicular phase - Contralateral - E1 -
## Luteal phase - Contralateral - E2 - Lumen-Stroma surface - Follicular phase - Contralateral - E1 - Mu
## Luteal phase - Contralateral - E2 - Muscle-Tunica surface - Follicular phase - Contralateral - E1 -
## Luteal phase - Ipsilateral - E1 - Lumen-Stroma surface - Follicular phase - Contralateral - E1 - Mus
## Luteal phase - Ipsilateral - E1 - Muscle-Tunica surface - Follicular phase - Contralateral - E1 - Mu
## Luteal phase - Ipsilateral - E2 - Lumen-Stroma surface - Follicular phase - Contralateral - E1 - Mus
## Luteal phase - Ipsilateral - E2 - Muscle-Tunica surface - Follicular phase - Contralateral - E1 - Mu
## Follicular phase - Contralateral - E2 - Muscle-Tunica surface - Follicular phase - Contralateral - E
## Follicular phase - Ipsilateral - E1 - Lumen-Stroma surface - Follicular phase - Contralateral - E2 -
## Follicular phase - Ipsilateral - E1 - Muscle-Tunica surface - Follicular phase - Contralateral - E2
## Follicular phase - Ipsilateral - E2 - Lumen-Stroma surface - Follicular phase - Contralateral - E2 -
## Follicular phase - Ipsilateral - E2 - Muscle-Tunica surface - Follicular phase - Contralateral - E2
## Luteal phase - Contralateral - E1 - Lumen-Stroma surface - Follicular phase - Contralateral - E2 - Lu
## Luteal phase - Contralateral - E1 - Muscle-Tunica surface - Follicular phase - Contralateral - E2 -
## Luteal phase - Contralateral - E2 - Lumen-Stroma surface - Follicular phase - Contralateral - E2 - Lu
## Luteal phase - Contralateral - E2 - Muscle-Tunica surface - Follicular phase - Contralateral - E2 -
## Luteal phase - Ipsilateral - E1 - Lumen-Stroma surface - Follicular phase - Contralateral - E2 - Lum
```

```
## Luteal phase - Ipsilateral - E1 - Muscle-Tunica surface - Follicular phase - Contralateral - E2 - Lur
## Luteal phase - Ipsilateral - E2 - Lumen-Stroma surface - Follicular phase - Contralateral - E2 - Lume
## Luteal phase - Ipsilateral - E2 - Muscle-Tunica surface - Follicular phase - Contralateral - E2 - Lur
## Follicular phase - Ipsilateral - E1 - Lumen-Stroma surface - Follicular phase - Contralateral - E2 -
## Follicular phase - Ipsilateral - E1 - Muscle-Tunica surface - Follicular phase - Contralateral - E2 -
## Follicular phase - Ipsilateral - E2 - Lumen-Stroma surface - Follicular phase - Contralateral - E2 -
## Follicular phase - Ipsilateral - E2 - Muscle-Tunica surface - Follicular phase - Contralateral - E2 -
## Luteal phase - Contralateral - E1 - Lumen-Stroma surface - Follicular phase - Contralateral - E2 - Mu
## Luteal phase - Contralateral - E1 - Muscle-Tunica surface - Follicular phase - Contralateral - E2 -
## Luteal phase - Contralateral - E2 - Lumen-Stroma surface - Follicular phase - Contralateral - E2 - Mu
## Luteal phase - Contralateral - E2 - Muscle-Tunica surface - Follicular phase - Contralateral - E2 -
## Luteal phase - Ipsilateral - E1 - Lumen-Stroma surface - Follicular phase - Contralateral - E2 - Musc
## Luteal phase - Ipsilateral - E1 - Muscle-Tunica surface - Follicular phase - Contralateral - E2 - Mus
## Luteal phase - Ipsilateral - E2 - Lumen-Stroma surface - Follicular phase - Contralateral - E2 - Musc
## Luteal phase - Ipsilateral - E2 - Muscle-Tunica surface - Follicular phase - Contralateral - E2 - Mus
## Follicular phase - Ipsilateral - E1 - Muscle-Tunica surface - Follicular phase - Ipsilateral - E1 -
## Follicular phase - Ipsilateral - E2 - Lumen-Stroma surface - Follicular phase - Ipsilateral - E1 - Lu
## Follicular phase - Ipsilateral - E2 - Muscle-Tunica surface - Follicular phase - Ipsilateral - E1 -
## Luteal phase - Contralateral - E1 - Lumen-Stroma surface - Follicular phase - Ipsilateral - E1 - Lume
## Luteal phase - Contralateral - E1 - Muscle-Tunica surface - Follicular phase - Ipsilateral - E1 - Lur
## Luteal phase - Contralateral - E2 - Lumen-Stroma surface - Follicular phase - Ipsilateral - E1 - Lume
## Luteal phase - Contralateral - E2 - Muscle-Tunica surface - Follicular phase - Ipsilateral - E1 - Lur
## Luteal phase - Ipsilateral - E1 - Lumen-Stroma surface - Follicular phase - Ipsilateral - E1 - Lumen-
## Luteal phase - Ipsilateral - E1 - Muscle-Tunica surface - Follicular phase - Ipsilateral - E1 - Lumen
## Luteal phase - Ipsilateral - E2 - Lumen-Stroma surface - Follicular phase - Ipsilateral - E1 - Lumen-
## Luteal phase - Ipsilateral - E2 - Muscle-Tunica surface - Follicular phase - Ipsilateral - E1 - Lumen
## Follicular phase - Ipsilateral - E2 - Lumen-Stroma surface - Follicular phase - Ipsilateral - E1 - Mu
## Follicular phase - Ipsilateral - E2 - Muscle-Tunica surface - Follicular phase - Ipsilateral - E1 -
## Luteal phase - Contralateral - E1 - Lumen-Stroma surface - Follicular phase - Ipsilateral - E1 - Musc
## Luteal phase - Contralateral - E1 - Muscle-Tunica surface - Follicular phase - Ipsilateral - E1 - Mus
## Luteal phase - Contralateral - E2 - Lumen-Stroma surface - Follicular phase - Ipsilateral - E1 - Musc
## Luteal phase - Contralateral - E2 - Muscle-Tunica surface - Follicular phase - Ipsilateral - E1 - Mus
## Luteal phase - Ipsilateral - E1 - Lumen-Stroma surface - Follicular phase - Ipsilateral - E1 - Muscle
## Luteal phase - Ipsilateral - E1 - Muscle-Tunica surface - Follicular phase - Ipsilateral - E1 - Muscl
## Luteal phase - Ipsilateral - E2 - Lumen-Stroma surface - Follicular phase - Ipsilateral - E1 - Muscle
## Luteal phase - Ipsilateral - E2 - Muscle-Tunica surface - Follicular phase - Ipsilateral - E1 - Muscl
## Follicular phase - Ipsilateral - E2 - Muscle-Tunica surface - Follicular phase - Ipsilateral - E2 -
## Luteal phase - Contralateral - E1 - Lumen-Stroma surface - Follicular phase - Ipsilateral - E2 - Lume
## Luteal phase - Contralateral - E1 - Muscle-Tunica surface - Follicular phase - Ipsilateral - E2 - Lur
## Luteal phase - Contralateral - E2 - Lumen-Stroma surface - Follicular phase - Ipsilateral - E2 - Lume
## Luteal phase - Contralateral - E2 - Muscle-Tunica surface - Follicular phase - Ipsilateral - E2 - Lur
## Luteal phase - Ipsilateral - E1 - Lumen-Stroma surface - Follicular phase - Ipsilateral - E2 - Lumen-
## Luteal phase - Ipsilateral - E1 - Muscle-Tunica surface - Follicular phase - Ipsilateral - E2 - Lumen
## Luteal phase - Ipsilateral - E2 - Lumen-Stroma surface - Follicular phase - Ipsilateral - E2 - Lumen-
## Luteal phase - Ipsilateral - E2 - Muscle-Tunica surface - Follicular phase - Ipsilateral - E2 - Lumen
## Luteal phase - Contralateral - E1 - Lumen-Stroma surface - Follicular phase - Ipsilateral - E2 - Musc
## Luteal phase - Contralateral - E1 - Muscle-Tunica surface - Follicular phase - Ipsilateral - E2 - Mus
## Luteal phase - Contralateral - E2 - Lumen-Stroma surface - Follicular phase - Ipsilateral - E2 - Musc
## Luteal phase - Contralateral - E2 - Muscle-Tunica surface - Follicular phase - Ipsilateral - E2 - Mus
## Luteal phase - Ipsilateral - E1 - Lumen-Stroma surface - Follicular phase - Ipsilateral - E2 - Muscle
## Luteal phase - Ipsilateral - E1 - Muscle-Tunica surface - Follicular phase - Ipsilateral - E2 - Muscl
## Luteal phase - Ipsilateral - E2 - Lumen-Stroma surface - Follicular phase - Ipsilateral - E2 - Muscle
## Luteal phase - Ipsilateral - E2 - Muscle-Tunica surface - Follicular phase - Ipsilateral - E2 - Muscl
## Luteal phase - Contralateral - E1 - Muscle-Tunica surface - Luteal phase - Contralateral - E1 - Lumen
```

```
## Luteal phase - Contralateral - E2 - Lumen-Stroma surface - Luteal phase - Contralateral - E1 - Lumen-
## Luteal phase - Contralateral - E2 - Muscle-Tunica surface - Luteal phase - Contralateral - E1 - Lumen
## Luteal phase - Ipsilateral - E1 - Lumen-Stroma surface - Luteal phase - Contralateral - E1 - Lumen-St
## Luteal phase - Ipsilateral - E1 - Muscle-Tunica surface - Luteal phase - Contralateral - E1 - Lumen-S
## Luteal phase - Ipsilateral - E2 - Lumen-Stroma surface - Luteal phase - Contralateral - E1 - Lumen-St
## Luteal phase - Ipsilateral - E2 - Muscle-Tunica surface - Luteal phase - Contralateral - E1 - Lumen-S
## Luteal phase - Contralateral - E2 - Lumen-Stroma surface - Luteal phase - Contralateral - E1 - Muscle
## Luteal phase - Contralateral - E2 - Muscle-Tunica surface - Luteal phase - Contralateral - E1 - Muscl
## Luteal phase - Ipsilateral - E1 - Lumen-Stroma surface - Luteal phase - Contralateral - E1 - Muscle-T
## Luteal phase - Ipsilateral - E1 - Muscle-Tunica surface - Luteal phase - Contralateral - E1 - Muscle-
## Luteal phase - Ipsilateral - E2 - Lumen-Stroma surface - Luteal phase - Contralateral - E1 - Muscle-T
## Luteal phase - Ipsilateral - E2 - Muscle-Tunica surface - Luteal phase - Contralateral - E1 - Muscle-
## Luteal phase - Contralateral - E2 - Muscle-Tunica surface - Luteal phase - Contralateral - E2 - Lumen
## Luteal phase - Ipsilateral - E1 - Lumen-Stroma surface - Luteal phase - Contralateral - E2 - Lumen-St
## Luteal phase - Ipsilateral - E1 - Muscle-Tunica surface - Luteal phase - Contralateral - E2 - Lumen-S
## Luteal phase - Ipsilateral - E2 - Lumen-Stroma surface - Luteal phase - Contralateral - E2 - Lumen-St
## Luteal phase - Ipsilateral - E2 - Muscle-Tunica surface - Luteal phase - Contralateral - E2 - Lumen-S
## Luteal phase - Ipsilateral - E1 - Lumen-Stroma surface - Luteal phase - Contralateral - E2 - Muscle-T
## Luteal phase - Ipsilateral - E1 - Muscle-Tunica surface - Luteal phase - Contralateral - E2 - Muscle-
## Luteal phase - Ipsilateral - E2 - Lumen-Stroma surface - Luteal phase - Contralateral - E2 - Muscle-T
## Luteal phase - Ipsilateral - E2 - Muscle-Tunica surface - Luteal phase - Contralateral - E2 - Muscle-
## Luteal phase - Ipsilateral - E1 - Muscle-Tunica surface - Luteal phase - Ipsilateral - E1 - Lumen-Str
## Luteal phase - Ipsilateral - E2 - Lumen-Stroma surface - Luteal phase - Ipsilateral - E1 - Lumen-Stro
## Luteal phase - Ipsilateral - E2 - Muscle-Tunica surface - Luteal phase - Ipsilateral - E1 - Lumen-Str
## Luteal phase - Ipsilateral - E2 - Lumen-Stroma surface - Luteal phase - Ipsilateral - E1 - Muscle-Tun
## Luteal phase - Ipsilateral - E2 - Muscle-Tunica surface - Luteal phase - Ipsilateral - E1 - Muscle-Tu
## Luteal phase - Ipsilateral - E2 - Muscle-Tunica surface - Luteal phase - Ipsilateral - E2 - Lumen-Str
##
## Follicular phase - Contralateral - E1 - Muscle-Tunica surface - Follicular phase - Contralateral - E
## Follicular phase - Contralateral - E2 - Lumen-Stroma surface - Follicular phase - Contralateral - E1
## Follicular phase - Contralateral - E2 - Muscle-Tunica surface - Follicular phase - Contralateral - E
## Follicular phase - Ipsilateral - E1 - Lumen-Stroma surface - Follicular phase - Contralateral - E1 -
## Follicular phase - Ipsilateral - E1 - Muscle-Tunica surface - Follicular phase - Contralateral - E1 -
## Follicular phase - Ipsilateral - E2 - Lumen-Stroma surface - Follicular phase - Contralateral - E1 -
## Follicular phase - Ipsilateral - E2 - Muscle-Tunica surface - Follicular phase - Contralateral - E1 -
## Luteal phase - Contralateral - E1 - Lumen-Stroma surface - Follicular phase - Contralateral - E1 - Lu
## Luteal phase - Contralateral - E1 - Muscle-Tunica surface - Follicular phase - Contralateral - E1 - 
## Luteal phase - Contralateral - E2 - Lumen-Stroma surface - Follicular phase - Contralateral - E1 - Lu
## Luteal phase - Contralateral - E2 - Muscle-Tunica surface - Follicular phase - Contralateral - E1 - 
## Luteal phase - Ipsilateral - E1 - Lumen-Stroma surface - Follicular phase - Contralateral - E1 - Lume
## Luteal phase - Ipsilateral - E1 - Muscle-Tunica surface - Follicular phase - Contralateral - E1 - Lum
## Luteal phase - Ipsilateral - E2 - Lumen-Stroma surface - Follicular phase - Contralateral - E1 - Lume
## Luteal phase - Ipsilateral - E2 - Muscle-Tunica surface - Follicular phase - Contralateral - E1 - Lum
## Follicular phase - Contralateral - E2 - Lumen-Stroma surface - Follicular phase - Contralateral - E1
## Follicular phase - Contralateral - E2 - Muscle-Tunica surface - Follicular phase - Contralateral - E
## Follicular phase - Ipsilateral - E1 - Lumen-Stroma surface - Follicular phase - Contralateral - E1 -
## Follicular phase - Ipsilateral - E1 - Muscle-Tunica surface - Follicular phase - Contralateral - E1 -
## Follicular phase - Ipsilateral - E2 - Lumen-Stroma surface - Follicular phase - Contralateral - E1 -
## Follicular phase - Ipsilateral - E2 - Muscle-Tunica surface - Follicular phase - Contralateral - E1 -
## Luteal phase - Contralateral - E1 - Lumen-Stroma surface - Follicular phase - Contralateral - E1 - Mu
## Luteal phase - Contralateral - E1 - Muscle-Tunica surface - Follicular phase - Contralateral - E1 - 
## Luteal phase - Contralateral - E2 - Lumen-Stroma surface - Follicular phase - Contralateral - E1 - Mu
## Luteal phase - Contralateral - E2 - Muscle-Tunica surface - Follicular phase - Contralateral - E1 - 
## Luteal phase - Ipsilateral - E1 - Lumen-Stroma surface - Follicular phase - Contralateral - E1 - Musc
```

```
## Luteal phase - Ipsilateral - E1 - Muscle-Tunica surface - Follicular phase - Contralateral - E1 - Mus
## Luteal phase - Ipsilateral - E2 - Lumen-Stroma surface - Follicular phase - Contralateral - E1 - Musc
## Luteal phase - Ipsilateral - E2 - Muscle-Tunica surface - Follicular phase - Contralateral - E1 - Mus
## Follicular phase - Contralateral - E2 - Muscle-Tunica surface - Follicular phase - Contralateral - E
## Follicular phase - Ipsilateral - E1 - Lumen-Stroma surface - Follicular phase - Contralateral - E2 -
## Follicular phase - Ipsilateral - E1 - Muscle-Tunica surface - Follicular phase - Contralateral - E2 -
## Follicular phase - Ipsilateral - E2 - Lumen-Stroma surface - Follicular phase - Contralateral - E2 -
## Follicular phase - Ipsilateral - E2 - Muscle-Tunica surface - Follicular phase - Contralateral - E2 -
## Luteal phase - Contralateral - E1 - Lumen-Stroma surface - Follicular phase - Contralateral - E2 - Lu
## Luteal phase - Contralateral - E1 - Muscle-Tunica surface - Follicular phase - Contralateral - E2 -
## Luteal phase - Contralateral - E2 - Lumen-Stroma surface - Follicular phase - Contralateral - E2 - Lu
## Luteal phase - Contralateral - E2 - Muscle-Tunica surface - Follicular phase - Contralateral - E2 -
## Luteal phase - Ipsilateral - E1 - Lumen-Stroma surface - Follicular phase - Contralateral - E2 - Lum
## Luteal phase - Ipsilateral - E1 - Muscle-Tunica surface - Follicular phase - Contralateral - E2 - Lu
## Luteal phase - Ipsilateral - E2 - Lumen-Stroma surface - Follicular phase - Contralateral - E2 - Lum
## Luteal phase - Ipsilateral - E2 - Muscle-Tunica surface - Follicular phase - Contralateral - E2 - Lu
## Follicular phase - Ipsilateral - E1 - Lumen-Stroma surface - Follicular phase - Contralateral - E2 -
## Follicular phase - Ipsilateral - E1 - Muscle-Tunica surface - Follicular phase - Contralateral - E2 -
## Follicular phase - Ipsilateral - E2 - Lumen-Stroma surface - Follicular phase - Contralateral - E2 -
## Follicular phase - Ipsilateral - E2 - Muscle-Tunica surface - Follicular phase - Contralateral - E2 -
## Luteal phase - Contralateral - E1 - Lumen-Stroma surface - Follicular phase - Contralateral - E2 - Mu
## Luteal phase - Contralateral - E1 - Muscle-Tunica surface - Follicular phase - Contralateral - E2 -
## Luteal phase - Contralateral - E2 - Lumen-Stroma surface - Follicular phase - Contralateral - E2 - Mu
## Luteal phase - Contralateral - E2 - Muscle-Tunica surface - Follicular phase - Contralateral - E2 -
## Luteal phase - Ipsilateral - E1 - Lumen-Stroma surface - Follicular phase - Contralateral - E2 - Musc
## Luteal phase - Ipsilateral - E1 - Muscle-Tunica surface - Follicular phase - Contralateral - E2 - Mus
## Luteal phase - Ipsilateral - E2 - Lumen-Stroma surface - Follicular phase - Contralateral - E2 - Musc
## Luteal phase - Ipsilateral - E2 - Muscle-Tunica surface - Follicular phase - Contralateral - E2 - Mus
## Follicular phase - Ipsilateral - E1 - Muscle-Tunica surface - Follicular phase - Ipsilateral - E1 -
## Follicular phase - Ipsilateral - E2 - Lumen-Stroma surface - Follicular phase - Ipsilateral - E1 - Lu
## Follicular phase - Ipsilateral - E2 - Muscle-Tunica surface - Follicular phase - Ipsilateral - E1 -
## Luteal phase - Contralateral - E1 - Lumen-Stroma surface - Follicular phase - Ipsilateral - E1 - Lum
## Luteal phase - Contralateral - E1 - Muscle-Tunica surface - Follicular phase - Ipsilateral - E1 - Lu
## Luteal phase - Contralateral - E2 - Lumen-Stroma surface - Follicular phase - Ipsilateral - E1 - Lum
## Luteal phase - Contralateral - E2 - Muscle-Tunica surface - Follicular phase - Ipsilateral - E1 - Lu
## Luteal phase - Ipsilateral - E1 - Lumen-Stroma surface - Follicular phase - Ipsilateral - E1 - Lumen-
## Luteal phase - Ipsilateral - E1 - Muscle-Tunica surface - Follicular phase - Ipsilateral - E1 - Lumen
## Luteal phase - Ipsilateral - E2 - Lumen-Stroma surface - Follicular phase - Ipsilateral - E1 - Lumen-
## Luteal phase - Ipsilateral - E2 - Muscle-Tunica surface - Follicular phase - Ipsilateral - E1 - Lumen
## Follicular phase - Ipsilateral - E2 - Lumen-Stroma surface - Follicular phase - Ipsilateral - E1 - Mu
## Follicular phase - Ipsilateral - E2 - Muscle-Tunica surface - Follicular phase - Ipsilateral - E1 -
## Luteal phase - Contralateral - E1 - Lumen-Stroma surface - Follicular phase - Ipsilateral - E1 - Musc
## Luteal phase - Contralateral - E1 - Muscle-Tunica surface - Follicular phase - Ipsilateral - E1 - Mus
## Luteal phase - Contralateral - E2 - Lumen-Stroma surface - Follicular phase - Ipsilateral - E1 - Musc
## Luteal phase - Contralateral - E2 - Muscle-Tunica surface - Follicular phase - Ipsilateral - E1 - Mus
## Luteal phase - Ipsilateral - E1 - Lumen-Stroma surface - Follicular phase - Ipsilateral - E1 - Muscle
## Luteal phase - Ipsilateral - E1 - Muscle-Tunica surface - Follicular phase - Ipsilateral - E1 - Muscl
## Luteal phase - Ipsilateral - E2 - Lumen-Stroma surface - Follicular phase - Ipsilateral - E1 - Muscle
## Luteal phase - Ipsilateral - E2 - Muscle-Tunica surface - Follicular phase - Ipsilateral - E1 - Muscl
## Follicular phase - Ipsilateral - E2 - Muscle-Tunica surface - Follicular phase - Ipsilateral - E2 -
## Luteal phase - Contralateral - E1 - Lumen-Stroma surface - Follicular phase - Ipsilateral - E2 - Lum
## Luteal phase - Contralateral - E1 - Muscle-Tunica surface - Follicular phase - Ipsilateral - E2 - Lu
## Luteal phase - Contralateral - E2 - Lumen-Stroma surface - Follicular phase - Ipsilateral - E2 - Lum
## Luteal phase - Contralateral - E2 - Muscle-Tunica surface - Follicular phase - Ipsilateral - E2 - Lu
```

```
## Luteal phase - Ipsilateral - E1 - Lumen-Stroma surface - Follicular phase - Ipsilateral - E2 - Lumen-
## Luteal phase - Ipsilateral - E1 - Muscle-Tunica surface - Follicular phase - Ipsilateral - E2 - Lumen
## Luteal phase - Ipsilateral - E2 - Lumen-Stroma surface - Follicular phase - Ipsilateral - E2 - Lumen-
## Luteal phase - Ipsilateral - E2 - Muscle-Tunica surface - Follicular phase - Ipsilateral - E2 - Lumen
## Luteal phase - Contralateral - E1 - Lumen-Stroma surface - Follicular phase - Ipsilateral - E2 - Musc
## Luteal phase - Contralateral - E1 - Muscle-Tunica surface - Follicular phase - Ipsilateral - E2 - Mus
## Luteal phase - Contralateral - E2 - Lumen-Stroma surface - Follicular phase - Ipsilateral - E2 - Musc
## Luteal phase - Contralateral - E2 - Muscle-Tunica surface - Follicular phase - Ipsilateral - E2 - Mus
## Luteal phase - Ipsilateral - E1 - Lumen-Stroma surface - Follicular phase - Ipsilateral - E2 - Muscle
## Luteal phase - Ipsilateral - E1 - Muscle-Tunica surface - Follicular phase - Ipsilateral - E2 - Muscl
## Luteal phase - Ipsilateral - E2 - Lumen-Stroma surface - Follicular phase - Ipsilateral - E2 - Muscle
## Luteal phase - Ipsilateral - E2 - Muscle-Tunica surface - Follicular phase - Ipsilateral - E2 - Muscl
## Luteal phase - Contralateral - E1 - Muscle-Tunica surface - Luteal phase - Contralateral - E1 - Lumen
## Luteal phase - Contralateral - E2 - Lumen-Stroma surface - Luteal phase - Contralateral - E1 - Lumen-
## Luteal phase - Contralateral - E2 - Muscle-Tunica surface - Luteal phase - Contralateral - E1 - Lumen
## Luteal phase - Ipsilateral - E1 - Lumen-Stroma surface - Luteal phase - Contralateral - E1 - Lumen-S
## Luteal phase - Ipsilateral - E1 - Muscle-Tunica surface - Luteal phase - Contralateral - E1 - Lumen-
## Luteal phase - Ipsilateral - E2 - Lumen-Stroma surface - Luteal phase - Contralateral - E1 - Lumen-S
## Luteal phase - Ipsilateral - E2 - Muscle-Tunica surface - Luteal phase - Contralateral - E1 - Lumen-
## Luteal phase - Contralateral - E2 - Lumen-Stroma surface - Luteal phase - Contralateral - E1 - Muscle
## Luteal phase - Contralateral - E2 - Muscle-Tunica surface - Luteal phase - Contralateral - E1 - Muscl
## Luteal phase - Ipsilateral - E1 - Lumen-Stroma surface - Luteal phase - Contralateral - E1 - Muscle-
## Luteal phase - Ipsilateral - E1 - Muscle-Tunica surface - Luteal phase - Contralateral - E1 - Muscle
## Luteal phase - Ipsilateral - E2 - Lumen-Stroma surface - Luteal phase - Contralateral - E1 - Muscle-
## Luteal phase - Ipsilateral - E2 - Muscle-Tunica surface - Luteal phase - Contralateral - E1 - Muscle
## Luteal phase - Contralateral - E2 - Muscle-Tunica surface - Luteal phase - Contralateral - E2 - Lumen
## Luteal phase - Ipsilateral - E1 - Lumen-Stroma surface - Luteal phase - Contralateral - E2 - Lumen-S
## Luteal phase - Ipsilateral - E1 - Muscle-Tunica surface - Luteal phase - Contralateral - E2 - Lumen-
## Luteal phase - Ipsilateral - E2 - Lumen-Stroma surface - Luteal phase - Contralateral - E2 - Lumen-S
## Luteal phase - Ipsilateral - E2 - Muscle-Tunica surface - Luteal phase - Contralateral - E2 - Lumen-
## Luteal phase - Ipsilateral - E1 - Lumen-Stroma surface - Luteal phase - Contralateral - E2 - Muscle-
## Luteal phase - Ipsilateral - E1 - Muscle-Tunica surface - Luteal phase - Contralateral - E2 - Muscle
## Luteal phase - Ipsilateral - E2 - Lumen-Stroma surface - Luteal phase - Contralateral - E2 - Muscle-
## Luteal phase - Ipsilateral - E2 - Muscle-Tunica surface - Luteal phase - Contralateral - E2 - Muscle
## Luteal phase - Ipsilateral - E1 - Muscle-Tunica surface - Luteal phase - Ipsilateral - E1 - Lumen-St
## Luteal phase - Ipsilateral - E2 - Lumen-Stroma surface - Luteal phase - Ipsilateral - E1 - Lumen-Str
## Luteal phase - Ipsilateral - E2 - Muscle-Tunica surface - Luteal phase - Ipsilateral - E1 - Lumen-St
## Luteal phase - Ipsilateral - E2 - Lumen-Stroma surface - Luteal phase - Ipsilateral - E1 - Muscle-Tu
## Luteal phase - Ipsilateral - E2 - Muscle-Tunica surface - Luteal phase - Ipsilateral - E1 - Muscle-T
## Luteal phase - Ipsilateral - E2 - Muscle-Tunica surface - Luteal phase - Ipsilateral - E2 - Lumen-St
##
## Follicular phase - Contralateral - E1 - Muscle-Tunica surface - Follicular phase - Contralateral - E
## Follicular phase - Contralateral - E2 - Lumen-Stroma surface - Follicular phase - Contralateral - E1
## Follicular phase - Contralateral - E2 - Muscle-Tunica surface - Follicular phase - Contralateral - E
## Follicular phase - Ipsilateral - E1 - Lumen-Stroma surface - Follicular phase - Contralateral - E1 -
## Follicular phase - Ipsilateral - E1 - Muscle-Tunica surface - Follicular phase - Contralateral - E1
## Follicular phase - Ipsilateral - E2 - Lumen-Stroma surface - Follicular phase - Contralateral - E1 -
## Follicular phase - Ipsilateral - E2 - Muscle-Tunica surface - Follicular phase - Contralateral - E1
## Luteal phase - Contralateral - E1 - Lumen-Stroma surface - Follicular phase - Contralateral - E1 - Lu
## Luteal phase - Contralateral - E1 - Muscle-Tunica surface - Follicular phase - Contralateral - E1 -
## Luteal phase - Contralateral - E2 - Lumen-Stroma surface - Follicular phase - Contralateral - E1 - Lu
## Luteal phase - Contralateral - E2 - Muscle-Tunica surface - Follicular phase - Contralateral - E1 -
## Luteal phase - Ipsilateral - E1 - Lumen-Stroma surface - Follicular phase - Contralateral - E1 - Lume
## Luteal phase - Ipsilateral - E1 - Muscle-Tunica surface - Follicular phase - Contralateral - E1 - Lu
```

```
## Luteal phase - Ipsilateral - E2 - Lumen-Stroma surface - Follicular phase - Contralateral - E1 - Lume
## Luteal phase - Ipsilateral - E2 - Muscle-Tunica surface - Follicular phase - Contralateral - E1 - Lu
## Follicular phase - Contralateral - E2 - Lumen-Stroma surface - Follicular phase - Contralateral - E1
## Follicular phase - Contralateral - E2 - Muscle-Tunica surface - Follicular phase - Contralateral - E
## Follicular phase - Ipsilateral - E1 - Lumen-Stroma surface - Follicular phase - Contralateral - E1 -
## Follicular phase - Ipsilateral - E1 - Muscle-Tunica surface - Follicular phase - Contralateral - E1 -
## Follicular phase - Ipsilateral - E2 - Lumen-Stroma surface - Follicular phase - Contralateral - E1 -
## Follicular phase - Ipsilateral - E2 - Muscle-Tunica surface - Follicular phase - Contralateral - E1 -
## Luteal phase - Contralateral - E1 - Lumen-Stroma surface - Follicular phase - Contralateral - E1 - Mu
## Luteal phase - Contralateral - E1 - Muscle-Tunica surface - Follicular phase - Contralateral - E1 - M
## Luteal phase - Contralateral - E2 - Lumen-Stroma surface - Follicular phase - Contralateral - E1 - Mu
## Luteal phase - Contralateral - E2 - Muscle-Tunica surface - Follicular phase - Contralateral - E1 - M
## Luteal phase - Ipsilateral - E1 - Lumen-Stroma surface - Follicular phase - Contralateral - E1 - Musc
## Luteal phase - Ipsilateral - E1 - Muscle-Tunica surface - Follicular phase - Contralateral - E1 - Mus
## Luteal phase - Ipsilateral - E2 - Lumen-Stroma surface - Follicular phase - Contralateral - E1 - Musc
## Luteal phase - Ipsilateral - E2 - Muscle-Tunica surface - Follicular phase - Contralateral - E1 - Mus
## Follicular phase - Contralateral - E2 - Muscle-Tunica surface - Follicular phase - Contralateral - E
## Follicular phase - Ipsilateral - E1 - Lumen-Stroma surface - Follicular phase - Contralateral - E2 -
## Follicular phase - Ipsilateral - E1 - Muscle-Tunica surface - Follicular phase - Contralateral - E2 -
## Follicular phase - Ipsilateral - E2 - Lumen-Stroma surface - Follicular phase - Contralateral - E2 -
## Follicular phase - Ipsilateral - E2 - Muscle-Tunica surface - Follicular phase - Contralateral - E2 -
## Luteal phase - Contralateral - E1 - Lumen-Stroma surface - Follicular phase - Contralateral - E2 - Lu
## Luteal phase - Contralateral - E1 - Muscle-Tunica surface - Follicular phase - Contralateral - E2 - M
## Luteal phase - Contralateral - E2 - Lumen-Stroma surface - Follicular phase - Contralateral - E2 - Lu
## Luteal phase - Contralateral - E2 - Muscle-Tunica surface - Follicular phase - Contralateral - E2 - M
## Luteal phase - Ipsilateral - E1 - Lumen-Stroma surface - Follicular phase - Contralateral - E2 - Lume
## Luteal phase - Ipsilateral - E1 - Muscle-Tunica surface - Follicular phase - Contralateral - E2 - Lu
## Luteal phase - Ipsilateral - E2 - Lumen-Stroma surface - Follicular phase - Contralateral - E2 - Lume
## Luteal phase - Ipsilateral - E2 - Muscle-Tunica surface - Follicular phase - Contralateral - E2 - Lu
## Follicular phase - Ipsilateral - E1 - Lumen-Stroma surface - Follicular phase - Contralateral - E2 -
## Follicular phase - Ipsilateral - E1 - Muscle-Tunica surface - Follicular phase - Contralateral - E2 -
## Follicular phase - Ipsilateral - E2 - Lumen-Stroma surface - Follicular phase - Contralateral - E2 -
## Follicular phase - Ipsilateral - E2 - Muscle-Tunica surface - Follicular phase - Contralateral - E2 -
## Luteal phase - Contralateral - E1 - Lumen-Stroma surface - Follicular phase - Contralateral - E2 - Mu
## Luteal phase - Contralateral - E1 - Muscle-Tunica surface - Follicular phase - Contralateral - E2 - M
## Luteal phase - Contralateral - E2 - Lumen-Stroma surface - Follicular phase - Contralateral - E2 - Mu
## Luteal phase - Contralateral - E2 - Muscle-Tunica surface - Follicular phase - Contralateral - E2 - M
## Luteal phase - Ipsilateral - E1 - Lumen-Stroma surface - Follicular phase - Contralateral - E2 - Musc
## Luteal phase - Ipsilateral - E1 - Muscle-Tunica surface - Follicular phase - Contralateral - E2 - Mus
## Luteal phase - Ipsilateral - E2 - Lumen-Stroma surface - Follicular phase - Contralateral - E2 - Musc
## Luteal phase - Ipsilateral - E2 - Muscle-Tunica surface - Follicular phase - Contralateral - E2 - Mus
## Follicular phase - Ipsilateral - E1 - Muscle-Tunica surface - Follicular phase - Ipsilateral - E1 - M
## Follicular phase - Ipsilateral - E2 - Lumen-Stroma surface - Follicular phase - Ipsilateral - E1 - Lu
## Follicular phase - Ipsilateral - E2 - Muscle-Tunica surface - Follicular phase - Ipsilateral - E1 - M
## Luteal phase - Contralateral - E1 - Lumen-Stroma surface - Follicular phase - Ipsilateral - E1 - Lume
## Luteal phase - Contralateral - E1 - Muscle-Tunica surface - Follicular phase - Ipsilateral - E1 - Lu
## Luteal phase - Contralateral - E2 - Lumen-Stroma surface - Follicular phase - Ipsilateral - E1 - Lume
## Luteal phase - Contralateral - E2 - Muscle-Tunica surface - Follicular phase - Ipsilateral - E1 - Lu
## Luteal phase - Ipsilateral - E1 - Lumen-Stroma surface - Follicular phase - Ipsilateral - E1 - Lumen-
## Luteal phase - Ipsilateral - E1 - Muscle-Tunica surface - Follicular phase - Ipsilateral - E1 - Lumen
## Luteal phase - Ipsilateral - E2 - Lumen-Stroma surface - Follicular phase - Ipsilateral - E1 - Lumen-
## Luteal phase - Ipsilateral - E2 - Muscle-Tunica surface - Follicular phase - Ipsilateral - E1 - Lumen
## Follicular phase - Ipsilateral - E2 - Lumen-Stroma surface - Follicular phase - Ipsilateral - E1 - Mu
## Follicular phase - Ipsilateral - E2 - Muscle-Tunica surface - Follicular phase - Ipsilateral - E1 - M
```

```
## Luteal phase - Contralateral - E1 - Lumen-Stroma surface - Follicular phase - Ipsilateral - E1 - Musc
## Luteal phase - Contralateral - E1 - Muscle-Tunica surface - Follicular phase - Ipsilateral - E1 - Mu
## Luteal phase - Contralateral - E2 - Lumen-Stroma surface - Follicular phase - Ipsilateral - E1 - Musc
## Luteal phase - Contralateral - E2 - Muscle-Tunica surface - Follicular phase - Ipsilateral - E1 - Mu
## Luteal phase - Ipsilateral - E1 - Lumen-Stroma surface - Follicular phase - Ipsilateral - E1 - Muscle
## Luteal phase - Ipsilateral - E1 - Muscle-Tunica surface - Follicular phase - Ipsilateral - E1 - Muscl
## Luteal phase - Ipsilateral - E2 - Lumen-Stroma surface - Follicular phase - Ipsilateral - E1 - Muscle
## Luteal phase - Ipsilateral - E2 - Muscle-Tunica surface - Follicular phase - Ipsilateral - E1 - Muscl
## Follicular phase - Ipsilateral - E2 - Muscle-Tunica surface - Follicular phase - Ipsilateral - E2 - 
## Luteal phase - Contralateral - E1 - Lumen-Stroma surface - Follicular phase - Ipsilateral - E2 - Lume
## Luteal phase - Contralateral - E1 - Muscle-Tunica surface - Follicular phase - Ipsilateral - E2 - Lu
## Luteal phase - Contralateral - E2 - Lumen-Stroma surface - Follicular phase - Ipsilateral - E2 - Lume
## Luteal phase - Contralateral - E2 - Muscle-Tunica surface - Follicular phase - Ipsilateral - E2 - Lu
## Luteal phase - Ipsilateral - E1 - Lumen-Stroma surface - Follicular phase - Ipsilateral - E2 - Lumen-
## Luteal phase - Ipsilateral - E1 - Muscle-Tunica surface - Follicular phase - Ipsilateral - E2 - Lumen
## Luteal phase - Ipsilateral - E2 - Lumen-Stroma surface - Follicular phase - Ipsilateral - E2 - Lumen-
## Luteal phase - Ipsilateral - E2 - Muscle-Tunica surface - Follicular phase - Ipsilateral - E2 - Lumen
## Luteal phase - Contralateral - E1 - Lumen-Stroma surface - Follicular phase - Ipsilateral - E2 - Musc
## Luteal phase - Contralateral - E1 - Muscle-Tunica surface - Follicular phase - Ipsilateral - E2 - Mu
## Luteal phase - Contralateral - E2 - Lumen-Stroma surface - Follicular phase - Ipsilateral - E2 - Musc
## Luteal phase - Contralateral - E2 - Muscle-Tunica surface - Follicular phase - Ipsilateral - E2 - Mu
## Luteal phase - Ipsilateral - E1 - Lumen-Stroma surface - Follicular phase - Ipsilateral - E2 - Muscle
## Luteal phase - Ipsilateral - E1 - Muscle-Tunica surface - Follicular phase - Ipsilateral - E2 - Muscl
## Luteal phase - Ipsilateral - E2 - Lumen-Stroma surface - Follicular phase - Ipsilateral - E2 - Muscle
## Luteal phase - Ipsilateral - E2 - Muscle-Tunica surface - Follicular phase - Ipsilateral - E2 - Muscl
## Luteal phase - Contralateral - E1 - Muscle-Tunica surface - Luteal phase - Contralateral - E1 - Lumen
## Luteal phase - Contralateral - E2 - Lumen-Stroma surface - Luteal phase - Contralateral - E1 - Lumen-
## Luteal phase - Contralateral - E2 - Muscle-Tunica surface - Luteal phase - Contralateral - E1 - Lumen
## Luteal phase - Ipsilateral - E1 - Lumen-Stroma surface - Luteal phase - Contralateral - E1 - Lumen-St
## Luteal phase - Ipsilateral - E1 - Muscle-Tunica surface - Luteal phase - Contralateral - E1 - Lumen-S
## Luteal phase - Ipsilateral - E2 - Lumen-Stroma surface - Luteal phase - Contralateral - E1 - Lumen-St
## Luteal phase - Ipsilateral - E2 - Muscle-Tunica surface - Luteal phase - Contralateral - E1 - Lumen-S
## Luteal phase - Contralateral - E2 - Lumen-Stroma surface - Luteal phase - Contralateral - E1 - Muscle
## Luteal phase - Contralateral - E2 - Muscle-Tunica surface - Luteal phase - Contralateral - E1 - Muscl
## Luteal phase - Ipsilateral - E1 - Lumen-Stroma surface - Luteal phase - Contralateral - E1 - Muscle-T
## Luteal phase - Ipsilateral - E1 - Muscle-Tunica surface - Luteal phase - Contralateral - E1 - Muscle-
## Luteal phase - Ipsilateral - E2 - Lumen-Stroma surface - Luteal phase - Contralateral - E1 - Muscle-T
## Luteal phase - Ipsilateral - E2 - Muscle-Tunica surface - Luteal phase - Contralateral - E1 - Muscle-
## Luteal phase - Contralateral - E2 - Muscle-Tunica surface - Luteal phase - Contralateral - E2 - Lumen
## Luteal phase - Ipsilateral - E1 - Lumen-Stroma surface - Luteal phase - Contralateral - E2 - Lumen-St
## Luteal phase - Ipsilateral - E1 - Muscle-Tunica surface - Luteal phase - Contralateral - E2 - Lumen-S
## Luteal phase - Ipsilateral - E2 - Lumen-Stroma surface - Luteal phase - Contralateral - E2 - Lumen-St
## Luteal phase - Ipsilateral - E2 - Muscle-Tunica surface - Luteal phase - Contralateral - E2 - Lumen-S
## Luteal phase - Ipsilateral - E1 - Lumen-Stroma surface - Luteal phase - Contralateral - E2 - Muscle-T
## Luteal phase - Ipsilateral - E1 - Muscle-Tunica surface - Luteal phase - Contralateral - E2 - Muscle-
## Luteal phase - Ipsilateral - E2 - Lumen-Stroma surface - Luteal phase - Contralateral - E2 - Muscle-T
## Luteal phase - Ipsilateral - E2 - Muscle-Tunica surface - Luteal phase - Contralateral - E2 - Muscle-
## Luteal phase - Ipsilateral - E1 - Muscle-Tunica surface - Luteal phase - Ipsilateral - E1 - Lumen-Str
## Luteal phase - Ipsilateral - E2 - Lumen-Stroma surface - Luteal phase - Ipsilateral - E1 - Lumen-Stro
## Luteal phase - Ipsilateral - E2 - Muscle-Tunica surface - Luteal phase - Ipsilateral - E1 - Lumen-Str
## Luteal phase - Ipsilateral - E2 - Lumen-Stroma surface - Luteal phase - Ipsilateral - E1 - Muscle-Tun
## Luteal phase - Ipsilateral - E2 - Muscle-Tunica surface - Luteal phase - Ipsilateral - E1 - Muscle-Tu
## Luteal phase - Ipsilateral - E2 - Muscle-Tunica surface - Luteal phase - Ipsilateral - E2 - Lumen-Str
##
```

```
## Follicular phase - Contralateral - E1 - Muscle-Tunica surface - Follicular phase - Contralateral - E
## Follicular phase - Contralateral - E2 - Lumen-Stroma surface - Follicular phase - Contralateral - E1
## Follicular phase - Contralateral - E2 - Muscle-Tunica surface - Follicular phase - Contralateral - E
## Follicular phase - Ipsilateral - E1 - Lumen-Stroma surface - Follicular phase - Contralateral - E1 -
## Follicular phase - Ipsilateral - E1 - Muscle-Tunica surface - Follicular phase - Contralateral - E1
## Follicular phase - Ipsilateral - E2 - Lumen-Stroma surface - Follicular phase - Contralateral - E1 -
## Follicular phase - Ipsilateral - E2 - Muscle-Tunica surface - Follicular phase - Contralateral - E1
## Luteal phase - Contralateral - E1 - Lumen-Stroma surface - Follicular phase - Contralateral - E1 - Lu
## Luteal phase - Contralateral - E1 - Muscle-Tunica surface - Follicular phase - Contralateral - E1 -
## Luteal phase - Contralateral - E2 - Lumen-Stroma surface - Follicular phase - Contralateral - E1 - Lu
## Luteal phase - Contralateral - E2 - Muscle-Tunica surface - Follicular phase - Contralateral - E1 -
## Luteal phase - Ipsilateral - E1 - Lumen-Stroma surface - Follicular phase - Contralateral - E1 - Lume
## Luteal phase - Ipsilateral - E1 - Muscle-Tunica surface - Follicular phase - Contralateral - E1 - Lum
## Luteal phase - Ipsilateral - E2 - Lumen-Stroma surface - Follicular phase - Contralateral - E1 - Lume
## Luteal phase - Ipsilateral - E2 - Muscle-Tunica surface - Follicular phase - Contralateral - E1 - Lum
## Follicular phase - Contralateral - E2 - Lumen-Stroma surface - Follicular phase - Contralateral - E1
## Follicular phase - Contralateral - E2 - Muscle-Tunica surface - Follicular phase - Contralateral - E
## Follicular phase - Ipsilateral - E1 - Lumen-Stroma surface - Follicular phase - Contralateral - E1 -
## Follicular phase - Ipsilateral - E1 - Muscle-Tunica surface - Follicular phase - Contralateral - E1
## Follicular phase - Ipsilateral - E2 - Lumen-Stroma surface - Follicular phase - Contralateral - E1 -
## Follicular phase - Ipsilateral - E2 - Muscle-Tunica surface - Follicular phase - Contralateral - E1
## Luteal phase - Contralateral - E1 - Lumen-Stroma surface - Follicular phase - Contralateral - E1 - Mu
## Luteal phase - Contralateral - E1 - Muscle-Tunica surface - Follicular phase - Contralateral - E1 -
## Luteal phase - Contralateral - E2 - Lumen-Stroma surface - Follicular phase - Contralateral - E1 - Mu
## Luteal phase - Contralateral - E2 - Muscle-Tunica surface - Follicular phase - Contralateral - E1 -
## Luteal phase - Ipsilateral - E1 - Lumen-Stroma surface - Follicular phase - Contralateral - E1 - Musc
## Luteal phase - Ipsilateral - E1 - Muscle-Tunica surface - Follicular phase - Contralateral - E1 - Mus
## Luteal phase - Ipsilateral - E2 - Lumen-Stroma surface - Follicular phase - Contralateral - E1 - Musc
## Luteal phase - Ipsilateral - E2 - Muscle-Tunica surface - Follicular phase - Contralateral - E1 - Mus
## Follicular phase - Contralateral - E2 - Muscle-Tunica surface - Follicular phase - Contralateral - E
## Follicular phase - Ipsilateral - E1 - Lumen-Stroma surface - Follicular phase - Contralateral - E2 -
## Follicular phase - Ipsilateral - E1 - Muscle-Tunica surface - Follicular phase - Contralateral - E2
## Follicular phase - Ipsilateral - E2 - Lumen-Stroma surface - Follicular phase - Contralateral - E2 -
## Follicular phase - Ipsilateral - E2 - Muscle-Tunica surface - Follicular phase - Contralateral - E2
## Luteal phase - Contralateral - E1 - Lumen-Stroma surface - Follicular phase - Contralateral - E2 - Lu
## Luteal phase - Contralateral - E1 - Muscle-Tunica surface - Follicular phase - Contralateral - E2 -
## Luteal phase - Contralateral - E2 - Lumen-Stroma surface - Follicular phase - Contralateral - E2 - Lu
## Luteal phase - Contralateral - E2 - Muscle-Tunica surface - Follicular phase - Contralateral - E2 -
## Luteal phase - Ipsilateral - E1 - Lumen-Stroma surface - Follicular phase - Contralateral - E2 - Lume
## Luteal phase - Ipsilateral - E1 - Muscle-Tunica surface - Follicular phase - Contralateral - E2 - Lum
## Luteal phase - Ipsilateral - E2 - Lumen-Stroma surface - Follicular phase - Contralateral - E2 - Lume
## Luteal phase - Ipsilateral - E2 - Muscle-Tunica surface - Follicular phase - Contralateral - E2 - Lum
## Follicular phase - Ipsilateral - E1 - Lumen-Stroma surface - Follicular phase - Contralateral - E2 -
## Follicular phase - Ipsilateral - E1 - Muscle-Tunica surface - Follicular phase - Contralateral - E2
## Follicular phase - Ipsilateral - E2 - Lumen-Stroma surface - Follicular phase - Contralateral - E2 -
## Follicular phase - Ipsilateral - E2 - Muscle-Tunica surface - Follicular phase - Contralateral - E2
## Luteal phase - Contralateral - E1 - Lumen-Stroma surface - Follicular phase - Contralateral - E2 - Mu
## Luteal phase - Contralateral - E1 - Muscle-Tunica surface - Follicular phase - Contralateral - E2 -
## Luteal phase - Contralateral - E2 - Lumen-Stroma surface - Follicular phase - Contralateral - E2 - Mu
## Luteal phase - Contralateral - E2 - Muscle-Tunica surface - Follicular phase - Contralateral - E2 -
## Luteal phase - Ipsilateral - E1 - Lumen-Stroma surface - Follicular phase - Contralateral - E2 - Musc
## Luteal phase - Ipsilateral - E1 - Muscle-Tunica surface - Follicular phase - Contralateral - E2 - Mus
## Luteal phase - Ipsilateral - E2 - Lumen-Stroma surface - Follicular phase - Contralateral - E2 - Musc
## Luteal phase - Ipsilateral - E2 - Muscle-Tunica surface - Follicular phase - Contralateral - E2 - Mus
```

```
## Follicular phase - Ipsilateral - E1 - Muscle-Tunica surface - Follicular phase - Ipsilateral - E1 -
## Follicular phase - Ipsilateral - E2 - Lumen-Stroma surface - Follicular phase - Ipsilateral - E1 - Lu
## Follicular phase - Ipsilateral - E2 - Muscle-Tunica surface - Follicular phase - Ipsilateral - E1 -
## Luteal phase - Contralateral - E1 - Lumen-Stroma surface - Follicular phase - Ipsilateral - E1 - Lume
## Luteal phase - Contralateral - E1 - Muscle-Tunica surface - Follicular phase - Ipsilateral - E1 - Lu
## Luteal phase - Contralateral - E2 - Lumen-Stroma surface - Follicular phase - Ipsilateral - E1 - Lume
## Luteal phase - Contralateral - E2 - Muscle-Tunica surface - Follicular phase - Ipsilateral - E1 - Lu
## Luteal phase - Ipsilateral - E1 - Lumen-Stroma surface - Follicular phase - Ipsilateral - E1 - Lumen
## Luteal phase - Ipsilateral - E1 - Muscle-Tunica surface - Follicular phase - Ipsilateral - E1 - Lume
## Luteal phase - Ipsilateral - E2 - Lumen-Stroma surface - Follicular phase - Ipsilateral - E1 - Lumen
## Luteal phase - Ipsilateral - E2 - Muscle-Tunica surface - Follicular phase - Ipsilateral - E1 - Lume
## Follicular phase - Ipsilateral - E2 - Lumen-Stroma surface - Follicular phase - Ipsilateral - E1 - Mu
## Follicular phase - Ipsilateral - E2 - Muscle-Tunica surface - Follicular phase - Ipsilateral - E1 -
## Luteal phase - Contralateral - E1 - Lumen-Stroma surface - Follicular phase - Ipsilateral - E1 - Mus
## Luteal phase - Contralateral - E1 - Muscle-Tunica surface - Follicular phase - Ipsilateral - E1 - Mu
## Luteal phase - Contralateral - E2 - Lumen-Stroma surface - Follicular phase - Ipsilateral - E1 - Mus
## Luteal phase - Contralateral - E2 - Muscle-Tunica surface - Follicular phase - Ipsilateral - E1 - Mu
## Luteal phase - Ipsilateral - E1 - Lumen-Stroma surface - Follicular phase - Ipsilateral - E1 - Muscl
## Luteal phase - Ipsilateral - E1 - Muscle-Tunica surface - Follicular phase - Ipsilateral - E1 - Musc
## Luteal phase - Ipsilateral - E2 - Lumen-Stroma surface - Follicular phase - Ipsilateral - E1 - Muscl
## Luteal phase - Ipsilateral - E2 - Muscle-Tunica surface - Follicular phase - Ipsilateral - E1 - Musc
## Follicular phase - Ipsilateral - E2 - Muscle-Tunica surface - Follicular phase - Ipsilateral - E2 -
## Luteal phase - Contralateral - E1 - Lumen-Stroma surface - Follicular phase - Ipsilateral - E2 - Lum
## Luteal phase - Contralateral - E1 - Muscle-Tunica surface - Follicular phase - Ipsilateral - E2 - Lu
## Luteal phase - Contralateral - E2 - Lumen-Stroma surface - Follicular phase - Ipsilateral - E2 - Lum
## Luteal phase - Contralateral - E2 - Muscle-Tunica surface - Follicular phase - Ipsilateral - E2 - Lu
## Luteal phase - Ipsilateral - E1 - Lumen-Stroma surface - Follicular phase - Ipsilateral - E2 - Lumen
## Luteal phase - Ipsilateral - E1 - Muscle-Tunica surface - Follicular phase - Ipsilateral - E2 - Lume
## Luteal phase - Ipsilateral - E2 - Lumen-Stroma surface - Follicular phase - Ipsilateral - E2 - Lumen
## Luteal phase - Ipsilateral - E2 - Muscle-Tunica surface - Follicular phase - Ipsilateral - E2 - Lume
## Luteal phase - Contralateral - E1 - Lumen-Stroma surface - Follicular phase - Ipsilateral - E2 - Mus
## Luteal phase - Contralateral - E1 - Muscle-Tunica surface - Follicular phase - Ipsilateral - E2 - Mu
## Luteal phase - Contralateral - E2 - Lumen-Stroma surface - Follicular phase - Ipsilateral - E2 - Mus
## Luteal phase - Contralateral - E2 - Muscle-Tunica surface - Follicular phase - Ipsilateral - E2 - Mu
## Luteal phase - Ipsilateral - E1 - Lumen-Stroma surface - Follicular phase - Ipsilateral - E2 - Muscl
## Luteal phase - Ipsilateral - E1 - Muscle-Tunica surface - Follicular phase - Ipsilateral - E2 - Musc
## Luteal phase - Ipsilateral - E2 - Lumen-Stroma surface - Follicular phase - Ipsilateral - E2 - Muscl
## Luteal phase - Ipsilateral - E2 - Muscle-Tunica surface - Follicular phase - Ipsilateral - E2 - Musc
## Luteal phase - Contralateral - E1 - Muscle-Tunica surface - Luteal phase - Contralateral - E1 - Lume
## Luteal phase - Contralateral - E2 - Lumen-Stroma surface - Luteal phase - Contralateral - E1 - Lumen
## Luteal phase - Contralateral - E2 - Muscle-Tunica surface - Luteal phase - Contralateral - E1 - Lume
## Luteal phase - Ipsilateral - E1 - Lumen-Stroma surface - Luteal phase - Contralateral - E1 - Lumen-S
## Luteal phase - Ipsilateral - E1 - Muscle-Tunica surface - Luteal phase - Contralateral - E1 - Lumen-
## Luteal phase - Ipsilateral - E2 - Lumen-Stroma surface - Luteal phase - Contralateral - E1 - Lumen-S
## Luteal phase - Ipsilateral - E2 - Muscle-Tunica surface - Luteal phase - Contralateral - E1 - Lumen-
## Luteal phase - Contralateral - E2 - Lumen-Stroma surface - Luteal phase - Contralateral - E1 - Muscl
## Luteal phase - Contralateral - E2 - Muscle-Tunica surface - Luteal phase - Contralateral - E1 - Musc
## Luteal phase - Ipsilateral - E1 - Lumen-Stroma surface - Luteal phase - Contralateral - E1 - Muscle-
## Luteal phase - Ipsilateral - E1 - Muscle-Tunica surface - Luteal phase - Contralateral - E1 - Muscle
## Luteal phase - Ipsilateral - E2 - Lumen-Stroma surface - Luteal phase - Contralateral - E1 - Muscle-
## Luteal phase - Ipsilateral - E2 - Muscle-Tunica surface - Luteal phase - Contralateral - E1 - Muscle
## Luteal phase - Contralateral - E2 - Muscle-Tunica surface - Luteal phase - Contralateral - E2 - Lume
## Luteal phase - Ipsilateral - E1 - Lumen-Stroma surface - Luteal phase - Contralateral - E2 - Lumen-S
## Luteal phase - Ipsilateral - E1 - Muscle-Tunica surface - Luteal phase - Contralateral - E2 - Lumen-
```

```
## Luteal phase - Ipsilateral - E2 - Lumen-Stroma surface - Luteal phase - Contralateral - E2 - Lumen-St
## Luteal phase - Ipsilateral - E2 - Muscle-Tunica surface - Luteal phase - Contralateral - E2 - Lumen-S
## Luteal phase - Ipsilateral - E1 - Lumen-Stroma surface - Luteal phase - Contralateral - E2 - Muscle-T
## Luteal phase - Ipsilateral - E1 - Muscle-Tunica surface - Luteal phase - Contralateral - E2 - Muscle-
## Luteal phase - Ipsilateral - E2 - Lumen-Stroma surface - Luteal phase - Contralateral - E2 - Muscle-T
## Luteal phase - Ipsilateral - E2 - Muscle-Tunica surface - Luteal phase - Contralateral - E2 - Muscle-
## Luteal phase - Ipsilateral - E1 - Muscle-Tunica surface - Luteal phase - Ipsilateral - E1 - Lumen-St
## Luteal phase - Ipsilateral - E2 - Lumen-Stroma surface - Luteal phase - Ipsilateral - E1 - Lumen-Stro
## Luteal phase - Ipsilateral - E2 - Muscle-Tunica surface - Luteal phase - Ipsilateral - E1 - Lumen-St
## Luteal phase - Ipsilateral - E2 - Lumen-Stroma surface - Luteal phase - Ipsilateral - E1 - Muscle-Tun
## Luteal phase - Ipsilateral - E2 - Muscle-Tunica surface - Luteal phase - Ipsilateral - E1 - Muscle-Tu
## Luteal phase - Ipsilateral - E2 - Muscle-Tunica surface - Luteal phase - Ipsilateral - E2 - Lumen-St
##
## Follicular phase - Contralateral - E1 - Muscle-Tunica surface - Follicular phase - Contralateral - E
## Follicular phase - Contralateral - E2 - Lumen-Stroma surface - Follicular phase - Contralateral - E1
## Follicular phase - Contralateral - E2 - Muscle-Tunica surface - Follicular phase - Contralateral - E
## Follicular phase - Ipsilateral - E1 - Lumen-Stroma surface - Follicular phase - Contralateral - E1 -
## Follicular phase - Ipsilateral - E1 - Muscle-Tunica surface - Follicular phase - Contralateral - E1
## Follicular phase - Ipsilateral - E2 - Lumen-Stroma surface - Follicular phase - Contralateral - E1 -
## Follicular phase - Ipsilateral - E2 - Muscle-Tunica surface - Follicular phase - Contralateral - E1
## Luteal phase - Contralateral - E1 - Lumen-Stroma surface - Follicular phase - Contralateral - E1 - Lu
## Luteal phase - Contralateral - E1 - Muscle-Tunica surface - Follicular phase - Contralateral - E1 -
## Luteal phase - Contralateral - E2 - Lumen-Stroma surface - Follicular phase - Contralateral - E1 - Lu
## Luteal phase - Contralateral - E2 - Muscle-Tunica surface - Follicular phase - Contralateral - E1 -
## Luteal phase - Ipsilateral - E1 - Lumen-Stroma surface - Follicular phase - Contralateral - E1 - Lume
## Luteal phase - Ipsilateral - E1 - Muscle-Tunica surface - Follicular phase - Contralateral - E1 - Lum
## Luteal phase - Ipsilateral - E2 - Lumen-Stroma surface - Follicular phase - Contralateral - E1 - Lume
## Luteal phase - Ipsilateral - E2 - Muscle-Tunica surface - Follicular phase - Contralateral - E1 - Lum
## Follicular phase - Contralateral - E2 - Lumen-Stroma surface - Follicular phase - Contralateral - E1
## Follicular phase - Contralateral - E2 - Muscle-Tunica surface - Follicular phase - Contralateral - E
## Follicular phase - Ipsilateral - E1 - Lumen-Stroma surface - Follicular phase - Contralateral - E1 -
## Follicular phase - Ipsilateral - E1 - Muscle-Tunica surface - Follicular phase - Contralateral - E1
## Follicular phase - Ipsilateral - E2 - Lumen-Stroma surface - Follicular phase - Contralateral - E1 -
## Follicular phase - Ipsilateral - E2 - Muscle-Tunica surface - Follicular phase - Contralateral - E1
## Luteal phase - Contralateral - E1 - Lumen-Stroma surface - Follicular phase - Contralateral - E1 - Mu
## Luteal phase - Contralateral - E1 - Muscle-Tunica surface - Follicular phase - Contralateral - E1 -
## Luteal phase - Contralateral - E2 - Lumen-Stroma surface - Follicular phase - Contralateral - E1 - Mu
## Luteal phase - Contralateral - E2 - Muscle-Tunica surface - Follicular phase - Contralateral - E1 -
## Luteal phase - Ipsilateral - E1 - Lumen-Stroma surface - Follicular phase - Contralateral - E1 - Musc
## Luteal phase - Ipsilateral - E1 - Muscle-Tunica surface - Follicular phase - Contralateral - E1 - Mus
## Luteal phase - Ipsilateral - E2 - Lumen-Stroma surface - Follicular phase - Contralateral - E1 - Musc
## Luteal phase - Ipsilateral - E2 - Muscle-Tunica surface - Follicular phase - Contralateral - E1 - Mus
## Follicular phase - Contralateral - E2 - Muscle-Tunica surface - Follicular phase - Contralateral - E
## Follicular phase - Ipsilateral - E1 - Lumen-Stroma surface - Follicular phase - Contralateral - E2 -
## Follicular phase - Ipsilateral - E1 - Muscle-Tunica surface - Follicular phase - Contralateral - E2
## Follicular phase - Ipsilateral - E2 - Lumen-Stroma surface - Follicular phase - Contralateral - E2 -
## Follicular phase - Ipsilateral - E2 - Muscle-Tunica surface - Follicular phase - Contralateral - E2
## Luteal phase - Contralateral - E1 - Lumen-Stroma surface - Follicular phase - Contralateral - E2 - Lu
## Luteal phase - Contralateral - E1 - Muscle-Tunica surface - Follicular phase - Contralateral - E2 -
## Luteal phase - Contralateral - E2 - Lumen-Stroma surface - Follicular phase - Contralateral - E2 - Lu
## Luteal phase - Contralateral - E2 - Muscle-Tunica surface - Follicular phase - Contralateral - E2 -
## Luteal phase - Ipsilateral - E1 - Lumen-Stroma surface - Follicular phase - Contralateral - E2 - Lume
## Luteal phase - Ipsilateral - E1 - Muscle-Tunica surface - Follicular phase - Contralateral - E2 - Lum
## Luteal phase - Ipsilateral - E2 - Lumen-Stroma surface - Follicular phase - Contralateral - E2 - Lume
```

```
## Luteal phase - Ipsilateral - E2 - Muscle-Tunica surface - Follicular phase - Contralateral - E2 - Lu
## Follicular phase - Ipsilateral - E1 - Lumen-Stroma surface - Follicular phase - Contralateral - E2 -
## Follicular phase - Ipsilateral - E1 - Muscle-Tunica surface - Follicular phase - Contralateral - E2 -
## Follicular phase - Ipsilateral - E2 - Lumen-Stroma surface - Follicular phase - Contralateral - E2 -
## Follicular phase - Ipsilateral - E2 - Muscle-Tunica surface - Follicular phase - Contralateral - E2 -
## Luteal phase - Contralateral - E1 - Lumen-Stroma surface - Follicular phase - Contralateral - E2 - Mu
## Luteal phase - Contralateral - E1 - Muscle-Tunica surface - Follicular phase - Contralateral - E2 -
## Luteal phase - Contralateral - E2 - Lumen-Stroma surface - Follicular phase - Contralateral - E2 - Mu
## Luteal phase - Contralateral - E2 - Muscle-Tunica surface - Follicular phase - Contralateral - E2 -
## Luteal phase - Ipsilateral - E1 - Lumen-Stroma surface - Follicular phase - Contralateral - E2 - Musc
## Luteal phase - Ipsilateral - E1 - Muscle-Tunica surface - Follicular phase - Contralateral - E2 - Mus
## Luteal phase - Ipsilateral - E2 - Lumen-Stroma surface - Follicular phase - Contralateral - E2 - Musc
## Luteal phase - Ipsilateral - E2 - Muscle-Tunica surface - Follicular phase - Contralateral - E2 - Mus
## Follicular phase - Ipsilateral - E1 - Muscle-Tunica surface - Follicular phase - Ipsilateral - E1 -
## Follicular phase - Ipsilateral - E2 - Lumen-Stroma surface - Follicular phase - Ipsilateral - E1 - Lu
## Follicular phase - Ipsilateral - E2 - Muscle-Tunica surface - Follicular phase - Ipsilateral - E1 -
## Luteal phase - Contralateral - E1 - Lumen-Stroma surface - Follicular phase - Ipsilateral - E1 - Lum
## Luteal phase - Contralateral - E1 - Muscle-Tunica surface - Follicular phase - Ipsilateral - E1 - Lu
## Luteal phase - Contralateral - E2 - Lumen-Stroma surface - Follicular phase - Ipsilateral - E1 - Lum
## Luteal phase - Contralateral - E2 - Muscle-Tunica surface - Follicular phase - Ipsilateral - E1 - Lu
## Luteal phase - Ipsilateral - E1 - Lumen-Stroma surface - Follicular phase - Ipsilateral - E1 - Lumen
## Luteal phase - Ipsilateral - E1 - Muscle-Tunica surface - Follicular phase - Ipsilateral - E1 - Lume
## Luteal phase - Ipsilateral - E2 - Lumen-Stroma surface - Follicular phase - Ipsilateral - E1 - Lumen
## Luteal phase - Ipsilateral - E2 - Muscle-Tunica surface - Follicular phase - Ipsilateral - E1 - Lume
## Follicular phase - Ipsilateral - E2 - Lumen-Stroma surface - Follicular phase - Ipsilateral - E1 - Mu
## Follicular phase - Ipsilateral - E2 - Muscle-Tunica surface - Follicular phase - Ipsilateral - E1 -
## Luteal phase - Contralateral - E1 - Lumen-Stroma surface - Follicular phase - Ipsilateral - E1 - Musc
## Luteal phase - Contralateral - E1 - Muscle-Tunica surface - Follicular phase - Ipsilateral - E1 - Mus
## Luteal phase - Contralateral - E2 - Lumen-Stroma surface - Follicular phase - Ipsilateral - E1 - Musc
## Luteal phase - Contralateral - E2 - Muscle-Tunica surface - Follicular phase - Ipsilateral - E1 - Mus
## Luteal phase - Ipsilateral - E1 - Lumen-Stroma surface - Follicular phase - Ipsilateral - E1 - Muscle
## Luteal phase - Ipsilateral - E1 - Muscle-Tunica surface - Follicular phase - Ipsilateral - E1 - Muscl
## Luteal phase - Ipsilateral - E2 - Lumen-Stroma surface - Follicular phase - Ipsilateral - E1 - Muscle
## Luteal phase - Ipsilateral - E2 - Muscle-Tunica surface - Follicular phase - Ipsilateral - E1 - Muscl
## Follicular phase - Ipsilateral - E2 - Muscle-Tunica surface - Follicular phase - Ipsilateral - E2 -
## Luteal phase - Contralateral - E1 - Lumen-Stroma surface - Follicular phase - Ipsilateral - E2 - Lum
## Luteal phase - Contralateral - E1 - Muscle-Tunica surface - Follicular phase - Ipsilateral - E2 - Lu
## Luteal phase - Contralateral - E2 - Lumen-Stroma surface - Follicular phase - Ipsilateral - E2 - Lum
## Luteal phase - Contralateral - E2 - Muscle-Tunica surface - Follicular phase - Ipsilateral - E2 - Lu
## Luteal phase - Ipsilateral - E1 - Lumen-Stroma surface - Follicular phase - Ipsilateral - E2 - Lumen
## Luteal phase - Ipsilateral - E1 - Muscle-Tunica surface - Follicular phase - Ipsilateral - E2 - Lume
## Luteal phase - Ipsilateral - E2 - Lumen-Stroma surface - Follicular phase - Ipsilateral - E2 - Lumen
## Luteal phase - Ipsilateral - E2 - Muscle-Tunica surface - Follicular phase - Ipsilateral - E2 - Lume
## Luteal phase - Contralateral - E1 - Lumen-Stroma surface - Follicular phase - Ipsilateral - E2 - Musc
## Luteal phase - Contralateral - E1 - Muscle-Tunica surface - Follicular phase - Ipsilateral - E2 - Mus
## Luteal phase - Contralateral - E2 - Lumen-Stroma surface - Follicular phase - Ipsilateral - E2 - Musc
## Luteal phase - Contralateral - E2 - Muscle-Tunica surface - Follicular phase - Ipsilateral - E2 - Mus
## Luteal phase - Ipsilateral - E1 - Lumen-Stroma surface - Follicular phase - Ipsilateral - E2 - Muscle
## Luteal phase - Ipsilateral - E1 - Muscle-Tunica surface - Follicular phase - Ipsilateral - E2 - Muscl
## Luteal phase - Ipsilateral - E2 - Lumen-Stroma surface - Follicular phase - Ipsilateral - E2 - Muscle
## Luteal phase - Ipsilateral - E2 - Muscle-Tunica surface - Follicular phase - Ipsilateral - E2 - Muscl
## Luteal phase - Contralateral - E1 - Muscle-Tunica surface - Luteal phase - Contralateral - E1 - Lume
## Luteal phase - Contralateral - E2 - Lumen-Stroma surface - Luteal phase - Contralateral - E1 - Lumen
## Luteal phase - Contralateral - E2 - Muscle-Tunica surface - Luteal phase - Contralateral - E1 - Lume
```

```
## Luteal phase - Ipsilateral - E1 - Lumen-Stroma surface - Luteal phase - Contralateral - E1 - Lumen-St
## Luteal phase - Ipsilateral - E1 - Muscle-Tunica surface - Luteal phase - Contralateral - E1 - Lumen-S
## Luteal phase - Ipsilateral - E2 - Lumen-Stroma surface - Luteal phase - Contralateral - E1 - Lumen-St
## Luteal phase - Ipsilateral - E2 - Muscle-Tunica surface - Luteal phase - Contralateral - E1 - Lumen-S
## Luteal phase - Contralateral - E2 - Lumen-Stroma surface - Luteal phase - Contralateral - E1 - Muscle
## Luteal phase - Contralateral - E2 - Muscle-Tunica surface - Luteal phase - Contralateral - E1 - Muscl
## Luteal phase - Ipsilateral - E1 - Lumen-Stroma surface - Luteal phase - Contralateral - E1 - Muscle-T
## Luteal phase - Ipsilateral - E1 - Muscle-Tunica surface - Luteal phase - Contralateral - E1 - Muscle-
## Luteal phase - Ipsilateral - E2 - Lumen-Stroma surface - Luteal phase - Contralateral - E1 - Muscle-T
## Luteal phase - Ipsilateral - E2 - Muscle-Tunica surface - Luteal phase - Contralateral - E1 - Muscle-
## Luteal phase - Contralateral - E2 - Muscle-Tunica surface - Luteal phase - Contralateral - E2 - Lumen
## Luteal phase - Ipsilateral - E1 - Lumen-Stroma surface - Luteal phase - Contralateral - E2 - Lumen-St
## Luteal phase - Ipsilateral - E1 - Muscle-Tunica surface - Luteal phase - Contralateral - E2 - Lumen-S
## Luteal phase - Ipsilateral - E2 - Lumen-Stroma surface - Luteal phase - Contralateral - E2 - Lumen-St
## Luteal phase - Ipsilateral - E2 - Muscle-Tunica surface - Luteal phase - Contralateral - E2 - Lumen-S
## Luteal phase - Ipsilateral - E1 - Lumen-Stroma surface - Luteal phase - Contralateral - E2 - Muscle-T
## Luteal phase - Ipsilateral - E1 - Muscle-Tunica surface - Luteal phase - Contralateral - E2 - Muscle-
## Luteal phase - Ipsilateral - E2 - Lumen-Stroma surface - Luteal phase - Contralateral - E2 - Muscle-T
## Luteal phase - Ipsilateral - E2 - Muscle-Tunica surface - Luteal phase - Contralateral - E2 - Muscle-
## Luteal phase - Ipsilateral - E1 - Muscle-Tunica surface - Luteal phase - Ipsilateral - E1 - Lumen-Str
## Luteal phase - Ipsilateral - E2 - Lumen-Stroma surface - Luteal phase - Ipsilateral - E1 - Lumen-Stro
## Luteal phase - Ipsilateral - E2 - Muscle-Tunica surface - Luteal phase - Ipsilateral - E1 - Lumen-Str
## Luteal phase - Ipsilateral - E2 - Lumen-Stroma surface - Luteal phase - Ipsilateral - E1 - Muscle-Tun
## Luteal phase - Ipsilateral - E2 - Muscle-Tunica surface - Luteal phase - Ipsilateral - E1 - Muscle-Tu
## Luteal phase - Ipsilateral - E2 - Muscle-Tunica surface - Luteal phase - Ipsilateral - E2 - Lumen-Str
## ---
## Signif. codes:  0 '***' 0.001 '**' 0.01 '*' 0.05 '.' 0.1 ' ' 1
## (Adjusted p values reported -- single-step method)
```

```r
################### YM ########################################

# Read the dataset from a CSV file
df <- read.csv("Oviduct_YM.csv")

# Summary data

summary_df <- df %>%
  group_by(Cellular, Tissue, Position, Estrus_cycle) %>%
  summarise(mean_YM = mean(YM),
            sd_YM = sd(YM))
```

```
## `summarise()` has grouped output by 'Cellular', 'Tissue', 'Position'. You can
## override using the `.groups` argument.
```

```r
# Summary data distance division

# Read the dataset from a CSV file
df <- read.csv("Oviduct_YM.csv")

# Create a new column to define the X_position group
df$X_position_group <- ifelse(df$X_position <= 400, "Lumen-Stroma surface", "Muscle-Tunica surface")

# Update the labels for Position, Estrus_cycle, and Tissue
df$Position <- factor(df$Position, labels = c("Contralateral", "Ipsilateral"))
df$Estrus_cycle <- factor(df$Estrus_cycle, labels = c("Follicular phase", "Luteal phase"))
```

```
df$Tissue <- factor(df$Tissue, labels = c("Ampulla", "Isthmus"))

summary_df <- df %>%
  group_by(Cellular, Tissue, Position, Estrus_cycle, X_position_group) %>%
  summarise(mean_YM = mean(YM),
            sd_YM = sd(YM))
```

```
## `summarise()` has grouped output by 'Cellular', 'Tissue', 'Position',
## 'Estrus_cycle'. You can override using the `.groups` argument.
```

```
# Plot

#Plot Isthmus vs Ampula


# Read the dataset from a CSV file
df <- read.csv("Oviduct_YM.csv")

png(filename="Oiduct_YM_muscle vs stroma2.png",
    width = 15, height = 10, units = "in",
    res = 600)

par(mgp=c(1.5,0.5,0),
    mar = c(4, #bottom
            4, #left
            3, #top
            2),
    cex.main = 0.8,
    family = "sans" #right
)

# Update the labels for Position, Estrus_cycle, and Tissue
df$Position <- factor(df$Position, labels = c("Contralateral", "Ipsilateral"))
df$Estrus_cycle <- factor(df$Estrus_cycle, labels = c("Follicular phase", "Luteal phase"))
df$Tissue <- factor(df$Tissue, labels = c("Ampulla", "Isthmus"))

# Create a combined factor variable for Estrus_cycle, Cellular, Position, and Tissue
df$Combined <- factor(paste(df$Estrus_cycle, df$Cellular, df$Position, df$Tissue, sep = " "),
                      levels = c("Follicular phase Native Contralateral Ampulla",
                                 "Follicular phase Decellularized Contralateral Ampulla",
                                 "Luteal phase Native Contralateral Ampulla",
                                 "Luteal phase Decellularized Contralateral Ampulla",
                                 "Follicular phase Native Ipsilateral Ampulla",
                                 "Follicular phase Decellularized Ipsilateral Ampulla",
                                 "Luteal phase Native Ipsilateral Ampulla",
                                 "Luteal phase Decellularized Ipsilateral Ampulla",
                                 "Follicular phase Native Contralateral Isthmus",
                                 "Follicular phase Decellularized Contralateral Isthmus",
                                 "Luteal phase Native Contralateral Isthmus",
                                 "Luteal phase Decellularized Contralateral Isthmus",
                                 "Follicular phase Native Ipsilateral Isthmus",
                                 "Follicular phase Decellularized Ipsilateral Isthmus",
                                 "Luteal phase Native Ipsilateral Isthmus",
                                 "Luteal phase Decellularized Ipsilateral Isthmus"))
```

```r
# Define fill colors for each factor level
fill_colors <- c("Follicular phase Native Contralateral Ampulla" = viridis_pal(option = "D")(1),
                 "Follicular phase Decellularized Contralateral Ampulla" = alpha(viridis_pal(option = "
                 "Luteal phase Native Contralateral Ampulla" = "#238A8DFF",
                 "Luteal phase Decellularized Contralateral Ampulla" = alpha("#238A8DFF", 0.2),
                 "Follicular phase Native Ipsilateral Ampulla" = viridis_pal(option = "D")(1),
                 "Follicular phase Decellularized Ipsilateral Ampulla" = alpha(viridis_pal(option = "D"
                 "Luteal phase Native Ipsilateral Ampulla" = "#238A8DFF",
                 "Luteal phase Decellularized Ipsilateral Ampulla" = alpha("#238A8DFF", 0.2),
                 "Follicular phase Native Contralateral Isthmus" = viridis_pal(option = "D")(1),
                 "Follicular phase Decellularized Contralateral Isthmus" = alpha(viridis_pal(option = "
                 "Luteal phase Native Contralateral Isthmus" = "#238A8DFF",
                 "Luteal phase Decellularized Contralateral Isthmus" = alpha("#238A8DFF", 0.2),
                 "Follicular phase Native Ipsilateral Isthmus" = viridis_pal(option = "D")(1),
                 "Follicular phase Decellularized Ipsilateral Isthmus" = alpha(viridis_pal(option = "D"
                 "Luteal phase Native Ipsilateral Isthmus" = "#238A8DFF",
                 "Luteal phase Decellularized Ipsilateral Isthmus" = alpha("#238A8DFF", 0.2))

# Create the plot using ggplot
ggplot(data = df, aes(x = Tissue, y = YM, fill = Combined)) +
  geom_boxplot(width = 0.7, outlier.shape = NA) +
  geom_jitter(position = position_jitterdodge(), alpha = 0.8) +
  xlab("Tissue") +
  ylab("Young's modulus (Pa)") +
  ggtitle("Boxplot and Individual Data Points of YM by Estrus_cycle, Cellular, Position, Tissue") +
  scale_fill_manual(values = fill_colors, guide = guide_legend(title = "Combined")) +
  theme_bw() +
  ylim(0, 30000) +
  facet_grid(Estrus_cycle ~ Position, space = "free_x", scales = "free_x") +
  theme(panel.grid.major = element_blank(), panel.grid.minor = element_blank(),
        axis.text.x = element_text(size = 11, angle = 0, hjust = 0.5),
        axis.text.y = element_text(size = 12, angle = 0, hjust = 0.5),
        axis.title.x = element_text(size = 18, face = "bold"),
        axis.title.y = element_text(size = 16, face = "bold"),
        legend.position = "right",
        strip.text = element_text(size = 14, face = "bold"))
```

```
## Warning: Removed 8 rows containing non-finite values (`stat_boxplot()`).

## Warning: Removed 8 rows containing missing values (`geom_point()`).
```

```r
dev.off()
```

```
## pdf
##   2
```

```r
# Plot considering distance

# Read the dataset from a CSV file
df <- read.csv("Oviduct_YM.csv")


png(filename="Oviduct_YM_distance.png",
    width = 20, height = 10, units = "in",
    res = 600)
```

```r
par(mgp=c(1.5,0.5,0),
    mar = c(4, #bottom
            4, #left
            3, #top
            2),
    cex.main = 0.8,
    family = "sans" #right
)

# Update the labels for Position and Estrus_cycle
df$Position <- factor(df$Position, labels = c("Contralateral", "Ipsilateral"))
df$Estrus_cycle <- factor(df$Estrus_cycle, labels = c("Follicular phase", "Luteal phase"))

# Create a combined factor variable for Estrus_cycle and Cellular
df$Combined <- factor(paste(df$Estrus_cycle, df$Cellular, sep = " "),
                      levels = c("Follicular phase Native", "Follicular phase Decellularized",
                                 "Luteal phase Native", "Luteal phase Decellularized"))

# Create the plot using ggplot
ggplot(data = df, aes(x = factor(X_position), y = YM, fill = Combined)) +
  geom_boxplot(outlier.shape = NA) +
  geom_jitter(position = position_jitterdodge(), alpha = 1) +
  xlab("Distance from lumen to outer wall (um)") +
  ylab("Young's modulus (Pa)") +
  ggtitle("Boxplot and Individual Data Points of YM by X_position, Estrus_cycle, Position, and Estrus_c
  scale_fill_manual(values = c("Follicular phase Native" = viridis_pal(option = "D")(1),
                               "Follicular phase Decellularized" = alpha(viridis_pal(option = "D")(1), 
                               "Luteal phase Native" = "#238A8DFF",
                               "Luteal phase Decellularized" = alpha("#238A8DFF", 0.2)),
                    guide = guide_legend(title = "Combined")) +
  scale_color_viridis_d() +
  theme_bw() +
  facet_grid(Tissue ~ Position + Estrus_cycle + Cellular, scales = "free_x", space = "free_x") +
  ylim(0, 30000) +
  theme(panel.grid.major = element_blank(), panel.grid.minor = element_blank(),
        axis.text.x = element_text(size = 10, angle = 0, vjust = 0.5, hjust = 0),
        axis.text.y = element_text(size = 12, angle = 0, hjust = 0.5),
        axis.title.x = element_text(size = 16, face = "bold"),
        axis.title.y = element_text(size = 16, face = "bold"),
        legend.position = "right",
        strip.text = element_text(size = 14, face = "bold"),
        legend.text = element_text(size = 14))  # Increase the size of the legend font
```

```
## Warning: Removed 8 rows containing non-finite values (`stat_boxplot()`).
## Removed 8 rows containing missing values (`geom_point()`).
```

```r
dev.off()
```

```
## pdf
##   2
```

```r
# Plot considering lumen-stroma and muscle-tunica


# Read the dataset from a CSV file
```

```r
df <- read.csv("Oviduct_YM.csv")


png(filename="Oiduct_YM_muscle vs stroma.png",
    width = 20, height = 10, units = "in",
    res = 600)

par(mgp=c(1.5,0.5,0),
    mar = c(4, #bottom
            4, #left
            3, #top
            2),
    cex.main = 0.8,
    family = "sans" #right
)


# Create a new column to define the X_position group
df$X_position_group <- ifelse(df$X_position <= 400, "Lumen-Stroma surface", "Muscle-Tunica surface")


# Update the labels for Position, Estrus_cycle, and Tissue
df$Position <- factor(df$Position, labels = c("Contralateral", "Ipsilateral"))
df$Estrus_cycle <- factor(df$Estrus_cycle, labels = c("Follicular phase", "Luteal phase"))
df$Tissue <- factor(df$Tissue, labels = c("Ampulla", "Isthmus"))

# Create a combined factor variable for Estrus_cycle, Cellular, Position, and Tissue
df$Combined <- factor(paste(df$Estrus_cycle, df$Cellular, df$Position, df$Tissue, sep = " "),
                      levels = c("Follicular phase Native Contralateral Ampulla",
                                 "Follicular phase Decellularized Contralateral Ampulla",
                                 "Luteal phase Native Contralateral Ampulla",
                                 "Luteal phase Decellularized Contralateral Ampulla",
                                 "Follicular phase Native Ipsilateral Ampulla",
                                 "Follicular phase Decellularized Ipsilateral Ampulla",
                                 "Luteal phase Native Ipsilateral Ampulla",
                                 "Luteal phase Decellularized Ipsilateral Ampulla",
                                 "Follicular phase Native Contralateral Isthmus",
                                 "Follicular phase Decellularized Contralateral Isthmus",
                                 "Luteal phase Native Contra-lateral Isthmus",
                                 "Luteal phase Decellularized Contralateral Isthmus",
                                 "Follicular phase Native Ipsilateral Isthmus",
                                 "Follicular phase Decellularized Ipsilateral Isthmus",
                                 "Luteal phase Native Ipsilateral Isthmus",
                                 "Luteal phase Decellularized Ipsilateral Isthmus"))

# Define fill colors for each factor level
fill_colors <- c("Follicular phase Native Contralateral Ampulla" = viridis_pal(option = "D")(1),
                 "Follicular phase Decellularized Contralateral Ampulla" = alpha(viridis_pal(option = "
                 "Luteal phase Native Contralateral Ampulla" = "#238A8DFF",
                 "Luteal phase Decellularized Contralateral Ampulla" = alpha("#238A8DFF", 0.2),
                 "Follicular phase Native Ipsilateral Ampulla" = viridis_pal(option = "D")(1),
                 "Follicular phase Decellularized Ipsilateral Ampulla" = alpha(viridis_pal(option = "D"
                 "Luteal phase Native Ipsilateral Ampulla" = "#238A8DFF",
```

```
                 "Luteal phase Decellularized Ipsilateral Ampulla" = alpha("#238A8DFF", 0.2),
                 "Follicular phase Native Contralateral Isthmus" = viridis_pal(option = "D")(1),
                 "Follicular phase Decellularized Contralateral Isthmus" = alpha(viridis_pal(option = "
                 "Luteal phase Native Contralateral Isthmus" = "#238A8DFF",
                 "Luteal phase Decellularized Contralateral Isthmus" = alpha("#238A8DFF", 0.2),
                 "Follicular phase Native Ipsilateral Isthmus" = viridis_pal(option = "D")(1),
                 "Follicular phase Decellularized Ipsilateral Isthmus" = alpha(viridis_pal(option = "D"
                 "Luteal phase Native Ipsilateral Isthmus" = "#238A8DFF",
                 "Luteal phase Decellularized Ipsilateral Isthmus" = alpha("#238A8DFF", 0.2))

# Create the plot using ggplot
ggplot(data = df, aes(x = X_position_group, y = YM, fill = Combined)) +
  geom_boxplot(width = 0.7, outlier.shape = NA) +
  geom_jitter(position = position_jitterdodge(), alpha = 0.8) +
  xlab("") +
  ylab("Young's modulus (Pa)") +
  ggtitle("Boxplot and Individual Data Points of YM by Estrus_cycle, Cellular, Position, Tissue, and X_
  scale_fill_manual(values = fill_colors, guide = guide_legend(title = "Combined")) +
  theme_bw() +
  ylim(0, 30000) +
  facet_grid(Estrus_cycle ~ Position + Tissue, space = "free_x", scales = "free_x") +
  scale_x_discrete(labels = c("Lumen-Stroma\n surface", "Muscle-Tunica\n surface"), expand = c(0, 0)) +
  theme(panel.grid.major = element_blank(), panel.grid.minor = element_blank(),
        axis.text.x = element_text(size = 11, angle = 0, hjust = 0.5),
        axis.text.y = element_text(size = 12, angle = 0, hjust = 0.5),
        axis.title.x = element_text(size = 18, face = "bold"),
        axis.title.y = element_text(size = 16, face = "bold"),
        legend.position = "right",
        strip.text = element_text(size = 14, face = "bold"))
```

```
## Warning: Removed 8 rows containing non-finite values (`stat_boxplot()`).
## Removed 8 rows containing missing values (`geom_point()`).
```

```
dev.off()
```

```
## pdf
##   2
```

```
# Stat

# Read the dataset from a CSV file
df <- read.csv("Oviduct_YM.csv")

# Update the labels for Position, Estrus_cycle, and Tissue
df$Position <- factor(df$Position, labels = c("Contralateral", "Ipsilateral"))
df$Estrus_cycle <- factor(df$Estrus_cycle, labels = c("Follicular phase", "Luteal phase"))
df$Cellular <- factor(df$Cellular, labels = c("Native", "Decellularized"))

# Create a factor variable for the comparisons
df$Comparison <- factor(paste(df$Tissue, df$Estrus_cycle, df$Position, df$Cellular, sep = " - "))

# Perform pairwise comparisons using Tukey's method
model <- lm(YM ~ Comparison, data = df)
comp <- glht(model, linfct = mcp(Comparison = "Tukey"))
```

```r
# Summarize the results
summary(comp)
```

```
## Warning in RET$pfunction("adjusted", ...): Completion with error > abseps
## Warning in RET$pfunction("adjusted", ...): Completion with error > abseps

## Warning in RET$pfunction("adjusted", ...): Completion with error > abseps

## Warning in RET$pfunction("adjusted", ...): Completion with error > abseps

## Warning in RET$pfunction("adjusted", ...): Completion with error > abseps

## Warning in RET$pfunction("adjusted", ...): Completion with error > abseps

## Warning in RET$pfunction("adjusted", ...): Completion with error > abseps

## Warning in RET$pfunction("adjusted", ...): Completion with error > abseps

## Warning in RET$pfunction("adjusted", ...): Completion with error > abseps

## Warning in RET$pfunction("adjusted", ...): Completion with error > abseps

## Warning in RET$pfunction("adjusted", ...): Completion with error > abseps

## Warning in RET$pfunction("adjusted", ...): Completion with error > abseps

## Warning in RET$pfunction("adjusted", ...): Completion with error > abseps

## Warning in RET$pfunction("adjusted", ...): Completion with error > abseps

## Warning in RET$pfunction("adjusted", ...): Completion with error > abseps

## Warning in RET$pfunction("adjusted", ...): Completion with error > abseps

## Warning in RET$pfunction("adjusted", ...): Completion with error > abseps

## Warning in RET$pfunction("adjusted", ...): Completion with error > abseps

## Warning in RET$pfunction("adjusted", ...): Completion with error > abseps

## Warning in RET$pfunction("adjusted", ...): Completion with error > abseps

## Warning in RET$pfunction("adjusted", ...): Completion with error > abseps

## Warning in RET$pfunction("adjusted", ...): Completion with error > abseps

## Warning in RET$pfunction("adjusted", ...): Completion with error > abseps

## Warning in RET$pfunction("adjusted", ...): Completion with error > abseps

## Warning in RET$pfunction("adjusted", ...): Completion with error > abseps

## Warning in RET$pfunction("adjusted", ...): Completion with error > abseps
```

```
## Warning in RET$pfunction("adjusted", ...): Completion with error > abseps

## Warning in RET$pfunction("adjusted", ...): Completion with error > abseps

## Warning in RET$pfunction("adjusted", ...): Completion with error > abseps

## Warning in RET$pfunction("adjusted", ...): Completion with error > abseps

## Warning in RET$pfunction("adjusted", ...): Completion with error > abseps

## Warning in RET$pfunction("adjusted", ...): Completion with error > abseps

## Warning in RET$pfunction("adjusted", ...): Completion with error > abseps

## Warning in RET$pfunction("adjusted", ...): Completion with error > abseps

## Warning in RET$pfunction("adjusted", ...): Completion with error > abseps

## Warning in RET$pfunction("adjusted", ...): Completion with error > abseps

## Warning in RET$pfunction("adjusted", ...): Completion with error > abseps

## Warning in RET$pfunction("adjusted", ...): Completion with error > abseps

## Warning in RET$pfunction("adjusted", ...): Completion with error > abseps

## Warning in RET$pfunction("adjusted", ...): Completion with error > abseps

## Warning in RET$pfunction("adjusted", ...): Completion with error > abseps

## Warning in RET$pfunction("adjusted", ...): Completion with error > abseps

## Warning in RET$pfunction("adjusted", ...): Completion with error > abseps
##
##   Simultaneous Tests for General Linear Hypotheses
##
## Multiple Comparisons of Means: Tukey Contrasts
##
##
## Fit: lm(formula = YM ~ Comparison, data = df)
##
## Linear Hypotheses:
##
## Ampulla - Follicular phase - Contralateral - Native - Ampulla - Follicular phase - Contralateral - De
## Ampulla - Follicular phase - Ipsilateral - Decellularized - Ampulla - Follicular phase - Contralatera
## Ampulla - Follicular phase - Ipsilateral - Native - Ampulla - Follicular phase - Contralateral - Dec
## Ampulla - Luteal phase - Contralateral - Decellularized - Ampulla - Follicular phase - Contralateral
## Ampulla - Luteal phase - Contralateral - Native - Ampulla - Follicular phase - Contralateral - Decel
## Ampulla - Luteal phase - Ipsilateral - Decellularized - Ampulla - Follicular phase - Contralateral -
## Ampulla - Luteal phase - Ipsilateral - Native - Ampulla - Follicular phase - Contralateral - Decellul
## Isthmus - Follicular phase - Contralateral - Decellularized - Ampulla - Follicular phase - Contralate
## Isthmus - Follicular phase - Contralateral - Native - Ampulla - Follicular phase - Contralateral - De
```

```
## Isthmus - Follicular phase - Ipsilateral - Decellularized - Ampulla - Follicular phase - Contralatera
## Isthmus - Follicular phase - Ipsilateral - Native - Ampulla - Follicular phase - Contralateral - Dec
## Isthmus - Luteal phase - Contralateral - Decellularized - Ampulla - Follicular phase - Contralateral
## Isthmus - Luteal phase - Contralateral - Native - Ampulla - Follicular phase - Contralateral - Decel
## Isthmus - Luteal phase - Ipsilateral - Decellularized - Ampulla - Follicular phase - Contralateral -
## Isthmus - Luteal phase - Ipsilateral - Native - Ampulla - Follicular phase - Contralateral - Decellu
## Ampulla - Follicular phase - Ipsilateral - Decellularized - Ampulla - Follicular phase - Contralatera
## Ampulla - Follicular phase - Ipsilateral - Native - Ampulla - Follicular phase - Contralateral - Nat
## Ampulla - Luteal phase - Contralateral - Decellularized - Ampulla - Follicular phase - Contralateral
## Ampulla - Luteal phase - Contralateral - Native - Ampulla - Follicular phase - Contralateral - Nativ
## Ampulla - Luteal phase - Ipsilateral - Decellularized - Ampulla - Follicular phase - Contralateral -
## Ampulla - Luteal phase - Ipsilateral - Native - Ampulla - Follicular phase - Contralateral - Native =
## Isthmus - Follicular phase - Contralateral - Decellularized - Ampulla - Follicular phase - Contralat
## Isthmus - Follicular phase - Contralateral - Native - Ampulla - Follicular phase - Contralateral - N
## Isthmus - Follicular phase - Ipsilateral - Decellularized - Ampulla - Follicular phase - Contralatera
## Isthmus - Follicular phase - Ipsilateral - Native - Ampulla - Follicular phase - Contralateral - Nat
## Isthmus - Luteal phase - Contralateral - Decellularized - Ampulla - Follicular phase - Contralateral
## Isthmus - Luteal phase - Contralateral - Native - Ampulla - Follicular phase - Contralateral - Nativ
## Isthmus - Luteal phase - Ipsilateral - Decellularized - Ampulla - Follicular phase - Contralateral -
## Isthmus - Luteal phase - Ipsilateral - Native - Ampulla - Follicular phase - Contralateral - Native =
## Ampulla - Follicular phase - Ipsilateral - Native - Ampulla - Follicular phase - Ipsilateral - Decel
## Ampulla - Luteal phase - Contralateral - Decellularized - Ampulla - Follicular phase - Ipsilateral -
## Ampulla - Luteal phase - Contralateral - Native - Ampulla - Follicular phase - Ipsilateral - Decellu
## Ampulla - Luteal phase - Ipsilateral - Decellularized - Ampulla - Follicular phase - Ipsilateral - De
## Ampulla - Luteal phase - Ipsilateral - Native - Ampulla - Follicular phase - Ipsilateral - Decellula
## Isthmus - Follicular phase - Contralateral - Decellularized - Ampulla - Follicular phase - Ipsilatera
## Isthmus - Follicular phase - Contralateral - Native - Ampulla - Follicular phase - Ipsilateral - Dec
## Isthmus - Follicular phase - Ipsilateral - Decellularized - Ampulla - Follicular phase - Ipsilateral
## Isthmus - Follicular phase - Ipsilateral - Native - Ampulla - Follicular phase - Ipsilateral - Decel
## Isthmus - Luteal phase - Contralateral - Decellularized - Ampulla - Follicular phase - Ipsilateral -
## Isthmus - Luteal phase - Contralateral - Native - Ampulla - Follicular phase - Ipsilateral - Decellu
## Isthmus - Luteal phase - Ipsilateral - Decellularized - Ampulla - Follicular phase - Ipsilateral - De
## Isthmus - Luteal phase - Ipsilateral - Native - Ampulla - Follicular phase - Ipsilateral - Decellula
## Ampulla - Luteal phase - Contralateral - Decellularized - Ampulla - Follicular phase - Ipsilateral -
## Ampulla - Luteal phase - Contralateral - Native - Ampulla - Follicular phase - Ipsilateral - Native =
## Ampulla - Luteal phase - Ipsilateral - Decellularized - Ampulla - Follicular phase - Ipsilateral - Na
## Ampulla - Luteal phase - Ipsilateral - Native - Ampulla - Follicular phase - Ipsilateral - Native ==
## Isthmus - Follicular phase - Contralateral - Decellularized - Ampulla - Follicular phase - Ipsilatera
## Isthmus - Follicular phase - Contralateral - Native - Ampulla - Follicular phase - Ipsilateral - Nat
## Isthmus - Follicular phase - Ipsilateral - Decellularized - Ampulla - Follicular phase - Ipsilateral
## Isthmus - Follicular phase - Ipsilateral - Native - Ampulla - Follicular phase - Ipsilateral - Nativ
## Isthmus - Luteal phase - Contralateral - Decellularized - Ampulla - Follicular phase - Ipsilateral -
## Isthmus - Luteal phase - Contralateral - Native - Ampulla - Follicular phase - Ipsilateral - Native =
## Isthmus - Luteal phase - Ipsilateral - Decellularized - Ampulla - Follicular phase - Ipsilateral - Na
## Isthmus - Luteal phase - Ipsilateral - Native - Ampulla - Follicular phase - Ipsilateral - Native ==
## Ampulla - Luteal phase - Contralateral - Native - Ampulla - Luteal phase - Contralateral - Decellula
## Ampulla - Luteal phase - Ipsilateral - Decellularized - Ampulla - Luteal phase - Contralateral - Dec
## Ampulla - Luteal phase - Ipsilateral - Native - Ampulla - Luteal phase - Contralateral - Decellulari
## Isthmus - Follicular phase - Contralateral - Decellularized - Ampulla - Luteal phase - Contralateral
## Isthmus - Follicular phase - Contralateral - Native - Ampulla - Luteal phase - Contralateral - Decel
## Isthmus - Follicular phase - Ipsilateral - Decellularized - Ampulla - Luteal phase - Contralateral -
## Isthmus - Follicular phase - Ipsilateral - Native - Ampulla - Luteal phase - Contralateral - Decellu
## Isthmus - Luteal phase - Contralateral - Decellularized - Ampulla - Luteal phase - Contralateral - De
## Isthmus - Luteal phase - Contralateral - Native - Ampulla - Luteal phase - Contralateral - Decellula
```

```
## Isthmus - Luteal phase - Ipsilateral - Decellularized - Ampulla - Luteal phase - Contralateral - Dec
## Isthmus - Luteal phase - Ipsilateral - Native - Ampulla - Luteal phase - Contralateral - Decellulari
## Ampulla - Luteal phase - Ipsilateral - Decellularized - Ampulla - Luteal phase - Contralateral - Nat
## Ampulla - Luteal phase - Ipsilateral - Native - Ampulla - Luteal phase - Contralateral - Native == 0
## Isthmus - Follicular phase - Contralateral - Decellularized - Ampulla - Luteal phase - Contralateral
## Isthmus - Follicular phase - Contralateral - Native - Ampulla - Luteal phase - Contralateral - Nativ
## Isthmus - Follicular phase - Ipsilateral - Decellularized - Ampulla - Luteal phase - Contralateral -
## Isthmus - Follicular phase - Ipsilateral - Native - Ampulla - Luteal phase - Contralateral - Native =
## Isthmus - Luteal phase - Contralateral - Decellularized - Ampulla - Luteal phase - Contralateral - Na
## Isthmus - Luteal phase - Contralateral - Native - Ampulla - Luteal phase - Contralateral - Native ==
## Isthmus - Luteal phase - Ipsilateral - Decellularized - Ampulla - Luteal phase - Contralateral - Nat
## Isthmus - Luteal phase - Ipsilateral - Native - Ampulla - Luteal phase - Contralateral - Native == 0
## Ampulla - Luteal phase - Ipsilateral - Native - Ampulla - Luteal phase - Ipsilateral - Decellularize
## Isthmus - Follicular phase - Contralateral - Decellularized - Ampulla - Luteal phase - Ipsilateral -
## Isthmus - Follicular phase - Contralateral - Native - Ampulla - Luteal phase - Ipsilateral - Decellu
## Isthmus - Follicular phase - Ipsilateral - Decellularized - Ampulla - Luteal phase - Ipsilateral - D
## Isthmus - Follicular phase - Ipsilateral - Native - Ampulla - Luteal phase - Ipsilateral - Decellula
## Isthmus - Luteal phase - Contralateral - Decellularized - Ampulla - Luteal phase - Ipsilateral - Dec
## Isthmus - Luteal phase - Contralateral - Native - Ampulla - Luteal phase - Ipsilateral - Decellulari
## Isthmus - Luteal phase - Ipsilateral - Decellularized - Ampulla - Luteal phase - Ipsilateral - Decel
## Isthmus - Luteal phase - Ipsilateral - Native - Ampulla - Luteal phase - Ipsilateral - Decellularize
## Isthmus - Follicular phase - Contralateral - Decellularized - Ampulla - Luteal phase - Ipsilateral -
## Isthmus - Follicular phase - Contralateral - Native - Ampulla - Luteal phase - Ipsilateral - Native =
## Isthmus - Follicular phase - Ipsilateral - Decellularized - Ampulla - Luteal phase - Ipsilateral - Na
## Isthmus - Follicular phase - Ipsilateral - Native - Ampulla - Luteal phase - Ipsilateral - Native ==
## Isthmus - Luteal phase - Contralateral - Decellularized - Ampulla - Luteal phase - Ipsilateral - Nat
## Isthmus - Luteal phase - Contralateral - Native - Ampulla - Luteal phase - Ipsilateral - Native == 0
## Isthmus - Luteal phase - Ipsilateral - Decellularized - Ampulla - Luteal phase - Ipsilateral - Nativ
## Isthmus - Luteal phase - Ipsilateral - Native - Ampulla - Luteal phase - Ipsilateral - Native == 0
## Isthmus - Follicular phase - Contralateral - Native - Isthmus - Follicular phase - Contralateral - D
## Isthmus - Follicular phase - Ipsilateral - Decellularized - Isthmus - Follicular phase - Contralater
## Isthmus - Follicular phase - Ipsilateral - Native - Isthmus - Follicular phase - Contralateral - Dec
## Isthmus - Luteal phase - Contralateral - Decellularized - Isthmus - Follicular phase - Contralateral
## Isthmus - Luteal phase - Contralateral - Native - Isthmus - Follicular phase - Contralateral - Decel
## Isthmus - Luteal phase - Ipsilateral - Decellularized - Isthmus - Follicular phase - Contralateral -
## Isthmus - Luteal phase - Ipsilateral - Native - Isthmus - Follicular phase - Contralateral - Decellu
## Isthmus - Follicular phase - Ipsilateral - Decellularized - Isthmus - Follicular phase - Contralater
## Isthmus - Follicular phase - Ipsilateral - Native - Isthmus - Follicular phase - Contralateral - Nat
## Isthmus - Luteal phase - Contralateral - Decellularized - Isthmus - Follicular phase - Contralateral
## Isthmus - Luteal phase - Contralateral - Native - Isthmus - Follicular phase - Contralateral - Nativ
## Isthmus - Luteal phase - Ipsilateral - Decellularized - Isthmus - Follicular phase - Contralateral -
## Isthmus - Luteal phase - Ipsilateral - Native - Isthmus - Follicular phase - Contralateral - Native =
## Isthmus - Follicular phase - Ipsilateral - Native - Isthmus - Follicular phase - Ipsilateral - Decel
## Isthmus - Luteal phase - Contralateral - Decellularized - Isthmus - Follicular phase - Ipsilateral -
## Isthmus - Luteal phase - Contralateral - Native - Isthmus - Follicular phase - Ipsilateral - Decellu
## Isthmus - Luteal phase - Ipsilateral - Decellularized - Isthmus - Follicular phase - Ipsilateral - D
## Isthmus - Luteal phase - Ipsilateral - Native - Isthmus - Follicular phase - Ipsilateral - Decellula
## Isthmus - Luteal phase - Contralateral - Decellularized - Isthmus - Follicular phase - Ipsilateral -
## Isthmus - Luteal phase - Contralateral - Native - Isthmus - Follicular phase - Ipsilateral - Native =
## Isthmus - Luteal phase - Ipsilateral - Decellularized - Isthmus - Follicular phase - Ipsilateral - Na
## Isthmus - Luteal phase - Ipsilateral - Native - Isthmus - Follicular phase - Ipsilateral - Native ==
## Isthmus - Luteal phase - Contralateral - Native - Isthmus - Luteal phase - Contralateral - Decellula
## Isthmus - Luteal phase - Ipsilateral - Decellularized - Isthmus - Luteal phase - Contralateral - Dec
## Isthmus - Luteal phase - Ipsilateral - Native - Isthmus - Luteal phase - Contralateral - Decellulari
```

```
## Isthmus - Luteal phase - Ipsilateral - Decellularized - Isthmus - Luteal phase - Contralateral - Nati
## Isthmus - Luteal phase - Ipsilateral - Native - Isthmus - Luteal phase - Contralateral - Native == 0
## Isthmus - Luteal phase - Ipsilateral - Native - Isthmus - Luteal phase - Ipsilateral - Decellularized
##
## Ampulla - Follicular phase - Contralateral - Native - Ampulla - Follicular phase - Contralateral - De
## Ampulla - Follicular phase - Ipsilateral - Decellularized - Ampulla - Follicular phase - Contralatera
## Ampulla - Follicular phase - Ipsilateral - Native - Ampulla - Follicular phase - Contralateral - Dec
## Ampulla - Luteal phase - Contralateral - Decellularized - Ampulla - Follicular phase - Contralateral
## Ampulla - Luteal phase - Contralateral - Native - Ampulla - Follicular phase - Contralateral - Decell
## Ampulla - Luteal phase - Ipsilateral - Decellularized - Ampulla - Follicular phase - Contralateral -
## Ampulla - Luteal phase - Ipsilateral - Native - Ampulla - Follicular phase - Contralateral - Decellul
## Isthmus - Follicular phase - Contralateral - Decellularized - Ampulla - Follicular phase - Contralate
## Isthmus - Follicular phase - Contralateral - Native - Ampulla - Follicular phase - Contralateral - De
## Isthmus - Follicular phase - Ipsilateral - Decellularized - Ampulla - Follicular phase - Contralatera
## Isthmus - Follicular phase - Ipsilateral - Native - Ampulla - Follicular phase - Contralateral - Dec
## Isthmus - Luteal phase - Contralateral - Decellularized - Ampulla - Follicular phase - Contralateral
## Isthmus - Luteal phase - Contralateral - Native - Ampulla - Follicular phase - Contralateral - Decell
## Isthmus - Luteal phase - Ipsilateral - Decellularized - Ampulla - Follicular phase - Contralateral -
## Isthmus - Luteal phase - Ipsilateral - Native - Ampulla - Follicular phase - Contralateral - Decellul
## Ampulla - Follicular phase - Ipsilateral - Decellularized - Ampulla - Follicular phase - Contralatera
## Ampulla - Follicular phase - Ipsilateral - Native - Ampulla - Follicular phase - Contralateral - Nati
## Ampulla - Luteal phase - Contralateral - Decellularized - Ampulla - Follicular phase - Contralateral
## Ampulla - Luteal phase - Contralateral - Native - Ampulla - Follicular phase - Contralateral - Native
## Ampulla - Luteal phase - Ipsilateral - Decellularized - Ampulla - Follicular phase - Contralateral -
## Ampulla - Luteal phase - Ipsilateral - Native - Ampulla - Follicular phase - Contralateral - Native =
## Isthmus - Follicular phase - Contralateral - Decellularized - Ampulla - Follicular phase - Contralate
## Isthmus - Follicular phase - Contralateral - Native - Ampulla - Follicular phase - Contralateral - Na
## Isthmus - Follicular phase - Ipsilateral - Decellularized - Ampulla - Follicular phase - Contralatera
## Isthmus - Follicular phase - Ipsilateral - Native - Ampulla - Follicular phase - Contralateral - Nati
## Isthmus - Luteal phase - Contralateral - Decellularized - Ampulla - Follicular phase - Contralateral
## Isthmus - Luteal phase - Contralateral - Native - Ampulla - Follicular phase - Contralateral - Native
## Isthmus - Luteal phase - Ipsilateral - Decellularized - Ampulla - Follicular phase - Contralateral -
## Isthmus - Luteal phase - Ipsilateral - Native - Ampulla - Follicular phase - Contralateral - Native =
## Ampulla - Follicular phase - Ipsilateral - Native - Ampulla - Follicular phase - Ipsilateral - Decell
## Ampulla - Luteal phase - Contralateral - Decellularized - Ampulla - Follicular phase - Ipsilateral -
## Ampulla - Luteal phase - Contralateral - Native - Ampulla - Follicular phase - Ipsilateral - Decellul
## Ampulla - Luteal phase - Ipsilateral - Decellularized - Ampulla - Follicular phase - Ipsilateral - De
## Ampulla - Luteal phase - Ipsilateral - Native - Ampulla - Follicular phase - Ipsilateral - Decellular
## Isthmus - Follicular phase - Contralateral - Decellularized - Ampulla - Follicular phase - Ipsilatera
## Isthmus - Follicular phase - Contralateral - Native - Ampulla - Follicular phase - Ipsilateral - Dec
## Isthmus - Follicular phase - Ipsilateral - Decellularized - Ampulla - Follicular phase - Ipsilateral
## Isthmus - Follicular phase - Ipsilateral - Native - Ampulla - Follicular phase - Ipsilateral - Decell
## Isthmus - Luteal phase - Contralateral - Decellularized - Ampulla - Follicular phase - Ipsilateral -
## Isthmus - Luteal phase - Contralateral - Native - Ampulla - Follicular phase - Ipsilateral - Decellul
## Isthmus - Luteal phase - Ipsilateral - Decellularized - Ampulla - Follicular phase - Ipsilateral - De
## Isthmus - Luteal phase - Ipsilateral - Native - Ampulla - Follicular phase - Ipsilateral - Decellular
## Ampulla - Luteal phase - Contralateral - Decellularized - Ampulla - Follicular phase - Ipsilateral -
## Ampulla - Luteal phase - Contralateral - Native - Ampulla - Follicular phase - Ipsilateral - Native =
## Ampulla - Luteal phase - Ipsilateral - Decellularized - Ampulla - Follicular phase - Ipsilateral - Na
## Ampulla - Luteal phase - Ipsilateral - Native - Ampulla - Follicular phase - Ipsilateral - Native ==
## Isthmus - Follicular phase - Contralateral - Decellularized - Ampulla - Follicular phase - Ipsilatera
## Isthmus - Follicular phase - Contralateral - Native - Ampulla - Follicular phase - Ipsilateral - Nati
## Isthmus - Follicular phase - Ipsilateral - Decellularized - Ampulla - Follicular phase - Ipsilateral
## Isthmus - Follicular phase - Ipsilateral - Native - Ampulla - Follicular phase - Ipsilateral - Native
```

```
## Isthmus - Luteal phase - Contralateral - Decellularized - Ampulla - Follicular phase - Ipsilateral -
## Isthmus - Luteal phase - Contralateral - Native - Ampulla - Follicular phase - Ipsilateral - Native =
## Isthmus - Luteal phase - Ipsilateral - Decellularized - Ampulla - Follicular phase - Ipsilateral - Na
## Isthmus - Luteal phase - Ipsilateral - Native - Ampulla - Follicular phase - Ipsilateral - Native ==
## Ampulla - Luteal phase - Contralateral - Native - Ampulla - Luteal phase - Contralateral - Decellula
## Ampulla - Luteal phase - Ipsilateral - Decellularized - Ampulla - Luteal phase - Contralateral - Dec
## Ampulla - Luteal phase - Ipsilateral - Native - Ampulla - Luteal phase - Contralateral - Decellulari
## Isthmus - Follicular phase - Contralateral - Decellularized - Ampulla - Luteal phase - Contralateral
## Isthmus - Follicular phase - Contralateral - Native - Ampulla - Luteal phase - Contralateral - Decel
## Isthmus - Follicular phase - Ipsilateral - Decellularized - Ampulla - Luteal phase - Contralateral -
## Isthmus - Follicular phase - Ipsilateral - Native - Ampulla - Luteal phase - Contralateral - Decellu
## Isthmus - Luteal phase - Contralateral - Decellularized - Ampulla - Luteal phase - Contralateral - De
## Isthmus - Luteal phase - Contralateral - Native - Ampulla - Luteal phase - Contralateral - Decellula
## Isthmus - Luteal phase - Ipsilateral - Decellularized - Ampulla - Luteal phase - Contralateral - Dec
## Isthmus - Luteal phase - Ipsilateral - Native - Ampulla - Luteal phase - Contralateral - Decellulari
## Ampulla - Luteal phase - Ipsilateral - Decellularized - Ampulla - Luteal phase - Contralateral - Nat
## Ampulla - Luteal phase - Ipsilateral - Native - Ampulla - Luteal phase - Contralateral - Native == 0
## Isthmus - Follicular phase - Contralateral - Decellularized - Ampulla - Luteal phase - Contralateral
## Isthmus - Follicular phase - Contralateral - Native - Ampulla - Luteal phase - Contralateral - Nativ
## Isthmus - Follicular phase - Ipsilateral - Decellularized - Ampulla - Luteal phase - Contralateral -
## Isthmus - Follicular phase - Ipsilateral - Native - Ampulla - Luteal phase - Contralateral - Native =
## Isthmus - Luteal phase - Contralateral - Decellularized - Ampulla - Luteal phase - Contralateral - Na
## Isthmus - Luteal phase - Contralateral - Native - Ampulla - Luteal phase - Contralateral - Native ==
## Isthmus - Luteal phase - Ipsilateral - Decellularized - Ampulla - Luteal phase - Contralateral - Nat
## Isthmus - Luteal phase - Ipsilateral - Native - Ampulla - Luteal phase - Contralateral - Native == 0
## Ampulla - Luteal phase - Ipsilateral - Native - Ampulla - Luteal phase - Ipsilateral - Decellularized
## Isthmus - Follicular phase - Contralateral - Decellularized - Ampulla - Luteal phase - Ipsilateral -
## Isthmus - Follicular phase - Contralateral - Native - Ampulla - Luteal phase - Ipsilateral - Decellu
## Isthmus - Follicular phase - Ipsilateral - Decellularized - Ampulla - Luteal phase - Ipsilateral - De
## Isthmus - Follicular phase - Ipsilateral - Native - Ampulla - Luteal phase - Ipsilateral - Decellula
## Isthmus - Luteal phase - Contralateral - Decellularized - Ampulla - Luteal phase - Ipsilateral - Dec
## Isthmus - Luteal phase - Contralateral - Native - Ampulla - Luteal phase - Ipsilateral - Decellulari
## Isthmus - Luteal phase - Ipsilateral - Decellularized - Ampulla - Luteal phase - Ipsilateral - Decel
## Isthmus - Luteal phase - Ipsilateral - Native - Ampulla - Luteal phase - Ipsilateral - Decellularized
## Isthmus - Follicular phase - Contralateral - Decellularized - Ampulla - Luteal phase - Ipsilateral -
## Isthmus - Follicular phase - Contralateral - Native - Ampulla - Luteal phase - Ipsilateral - Native =
## Isthmus - Follicular phase - Ipsilateral - Decellularized - Ampulla - Luteal phase - Ipsilateral - Na
## Isthmus - Follicular phase - Ipsilateral - Native - Ampulla - Luteal phase - Ipsilateral - Native ==
## Isthmus - Luteal phase - Contralateral - Decellularized - Ampulla - Luteal phase - Ipsilateral - Nat
## Isthmus - Luteal phase - Contralateral - Native - Ampulla - Luteal phase - Ipsilateral - Native == 0
## Isthmus - Luteal phase - Ipsilateral - Decellularized - Ampulla - Luteal phase - Ipsilateral - Native
## Isthmus - Luteal phase - Ipsilateral - Native - Ampulla - Luteal phase - Ipsilateral - Native == 0
## Isthmus - Follicular phase - Contralateral - Native - Isthmus - Follicular phase - Contralateral - De
## Isthmus - Follicular phase - Ipsilateral - Decellularized - Isthmus - Follicular phase - Contralatera
## Isthmus - Follicular phase - Ipsilateral - Native - Isthmus - Follicular phase - Contralateral - Dec
## Isthmus - Luteal phase - Contralateral - Decellularized - Isthmus - Follicular phase - Contralateral
## Isthmus - Luteal phase - Contralateral - Native - Isthmus - Follicular phase - Contralateral - Decel
## Isthmus - Luteal phase - Ipsilateral - Decellularized - Isthmus - Follicular phase - Contralateral -
## Isthmus - Luteal phase - Ipsilateral - Native - Isthmus - Follicular phase - Contralateral - Decellu
## Isthmus - Follicular phase - Ipsilateral - Decellularized - Isthmus - Follicular phase - Contralatera
## Isthmus - Follicular phase - Ipsilateral - Native - Isthmus - Follicular phase - Contralateral - Nat
## Isthmus - Luteal phase - Contralateral - Decellularized - Isthmus - Follicular phase - Contralateral
## Isthmus - Luteal phase - Contralateral - Native - Isthmus - Follicular phase - Contralateral - Nativ
## Isthmus - Luteal phase - Ipsilateral - Decellularized - Isthmus - Follicular phase - Contralateral -
```

```
## Isthmus - Luteal phase - Ipsilateral - Native - Isthmus - Follicular phase - Contralateral - Native =
## Isthmus - Follicular phase - Ipsilateral - Native - Isthmus - Follicular phase - Ipsilateral - Decell
## Isthmus - Luteal phase - Contralateral - Decellularized - Isthmus - Follicular phase - Ipsilateral -
## Isthmus - Luteal phase - Contralateral - Native - Isthmus - Follicular phase - Ipsilateral - Decellul
## Isthmus - Luteal phase - Ipsilateral - Decellularized - Isthmus - Follicular phase - Ipsilateral - De
## Isthmus - Luteal phase - Ipsilateral - Native - Isthmus - Follicular phase - Ipsilateral - Decellula
## Isthmus - Luteal phase - Contralateral - Decellularized - Isthmus - Follicular phase - Ipsilateral -
## Isthmus - Luteal phase - Contralateral - Native - Isthmus - Follicular phase - Ipsilateral - Native =
## Isthmus - Luteal phase - Ipsilateral - Decellularized - Isthmus - Follicular phase - Ipsilateral - Na
## Isthmus - Luteal phase - Ipsilateral - Native - Isthmus - Follicular phase - Ipsilateral - Native ==
## Isthmus - Luteal phase - Contralateral - Native - Isthmus - Luteal phase - Contralateral - Decellula
## Isthmus - Luteal phase - Ipsilateral - Decellularized - Isthmus - Luteal phase - Contralateral - Dec
## Isthmus - Luteal phase - Ipsilateral - Native - Isthmus - Luteal phase - Contralateral - Decellulari
## Isthmus - Luteal phase - Ipsilateral - Decellularized - Isthmus - Luteal phase - Contralateral - Nat
## Isthmus - Luteal phase - Ipsilateral - Native - Isthmus - Luteal phase - Contralateral - Native == 0
## Isthmus - Luteal phase - Ipsilateral - Native - Isthmus - Luteal phase - Ipsilateral - Decellularize
##
## Ampulla - Follicular phase - Contralateral - Native - Ampulla - Follicular phase - Contralateral - De
## Ampulla - Follicular phase - Ipsilateral - Decellularized - Ampulla - Follicular phase - Contralatera
## Ampulla - Follicular phase - Ipsilateral - Native - Ampulla - Follicular phase - Contralateral - Dec
## Ampulla - Luteal phase - Contralateral - Decellularized - Ampulla - Follicular phase - Contralateral
## Ampulla - Luteal phase - Contralateral - Native - Ampulla - Follicular phase - Contralateral - Decel
## Ampulla - Luteal phase - Ipsilateral - Decellularized - Ampulla - Follicular phase - Contralateral -
## Ampulla - Luteal phase - Ipsilateral - Native - Ampulla - Follicular phase - Contralateral - Decellu
## Isthmus - Follicular phase - Contralateral - Decellularized - Ampulla - Follicular phase - Contralate
## Isthmus - Follicular phase - Contralateral - Native - Ampulla - Follicular phase - Contralateral - De
## Isthmus - Follicular phase - Ipsilateral - Decellularized - Ampulla - Follicular phase - Contralatera
## Isthmus - Follicular phase - Ipsilateral - Native - Ampulla - Follicular phase - Contralateral - Dec
## Isthmus - Luteal phase - Contralateral - Decellularized - Ampulla - Follicular phase - Contralateral
## Isthmus - Luteal phase - Contralateral - Native - Ampulla - Follicular phase - Contralateral - Decel
## Isthmus - Luteal phase - Ipsilateral - Decellularized - Ampulla - Follicular phase - Contralateral -
## Isthmus - Luteal phase - Ipsilateral - Native - Ampulla - Follicular phase - Contralateral - Decellu
## Ampulla - Follicular phase - Ipsilateral - Decellularized - Ampulla - Follicular phase - Contralatera
## Ampulla - Follicular phase - Ipsilateral - Native - Ampulla - Follicular phase - Contralateral - Nati
## Ampulla - Luteal phase - Contralateral - Decellularized - Ampulla - Follicular phase - Contralateral
## Ampulla - Luteal phase - Contralateral - Native - Ampulla - Follicular phase - Contralateral - Native
## Ampulla - Luteal phase - Ipsilateral - Decellularized - Ampulla - Follicular phase - Contralateral -
## Ampulla - Luteal phase - Ipsilateral - Native - Ampulla - Follicular phase - Contralateral - Native =
## Isthmus - Follicular phase - Contralateral - Decellularized - Ampulla - Follicular phase - Contralate
## Isthmus - Follicular phase - Contralateral - Native - Ampulla - Follicular phase - Contralateral - Na
## Isthmus - Follicular phase - Ipsilateral - Decellularized - Ampulla - Follicular phase - Contralatera
## Isthmus - Follicular phase - Ipsilateral - Native - Ampulla - Follicular phase - Contralateral - Nati
## Isthmus - Luteal phase - Contralateral - Decellularized - Ampulla - Follicular phase - Contralateral
## Isthmus - Luteal phase - Contralateral - Native - Ampulla - Follicular phase - Contralateral - Native
## Isthmus - Luteal phase - Ipsilateral - Decellularized - Ampulla - Follicular phase - Contralateral -
## Isthmus - Luteal phase - Ipsilateral - Native - Ampulla - Follicular phase - Contralateral - Native =
## Ampulla - Follicular phase - Ipsilateral - Native - Ampulla - Follicular phase - Ipsilateral - Decell
## Ampulla - Luteal phase - Contralateral - Decellularized - Ampulla - Follicular phase - Ipsilateral -
## Ampulla - Luteal phase - Contralateral - Native - Ampulla - Follicular phase - Ipsilateral - Decellul
## Ampulla - Luteal phase - Ipsilateral - Decellularized - Ampulla - Follicular phase - Ipsilateral - De
## Ampulla - Luteal phase - Ipsilateral - Native - Ampulla - Follicular phase - Ipsilateral - Decellula
## Isthmus - Follicular phase - Contralateral - Decellularized - Ampulla - Follicular phase - Ipsilatera
## Isthmus - Follicular phase - Contralateral - Native - Ampulla - Follicular phase - Ipsilateral - Dec
## Isthmus - Follicular phase - Ipsilateral - Decellularized - Ampulla - Follicular phase - Ipsilateral
```

```
## Isthmus - Follicular phase - Ipsilateral - Native - Ampulla - Follicular phase - Ipsilateral - Decell
## Isthmus - Luteal phase - Contralateral - Decellularized - Ampulla - Follicular phase - Ipsilateral -
## Isthmus - Luteal phase - Contralateral - Native - Ampulla - Follicular phase - Ipsilateral - Decellul
## Isthmus - Luteal phase - Ipsilateral - Decellularized - Ampulla - Follicular phase - Ipsilateral - De
## Isthmus - Luteal phase - Ipsilateral - Native - Ampulla - Follicular phase - Ipsilateral - Decellula
## Ampulla - Luteal phase - Contralateral - Decellularized - Ampulla - Follicular phase - Ipsilateral -
## Ampulla - Luteal phase - Contralateral - Native - Ampulla - Follicular phase - Ipsilateral - Native =
## Ampulla - Luteal phase - Ipsilateral - Decellularized - Ampulla - Follicular phase - Ipsilateral - Na
## Ampulla - Luteal phase - Ipsilateral - Native - Ampulla - Follicular phase - Ipsilateral - Native ==
## Isthmus - Follicular phase - Contralateral - Decellularized - Ampulla - Follicular phase - Ipsilatera
## Isthmus - Follicular phase - Contralateral - Native - Ampulla - Follicular phase - Ipsilateral - Nati
## Isthmus - Follicular phase - Ipsilateral - Decellularized - Ampulla - Follicular phase - Ipsilateral
## Isthmus - Follicular phase - Ipsilateral - Native - Ampulla - Follicular phase - Ipsilateral - Native
## Isthmus - Luteal phase - Contralateral - Decellularized - Ampulla - Follicular phase - Ipsilateral -
## Isthmus - Luteal phase - Contralateral - Native - Ampulla - Follicular phase - Ipsilateral - Native =
## Isthmus - Luteal phase - Ipsilateral - Decellularized - Ampulla - Follicular phase - Ipsilateral - Na
## Isthmus - Luteal phase - Ipsilateral - Native - Ampulla - Follicular phase - Ipsilateral - Native ==
## Ampulla - Luteal phase - Contralateral - Native - Ampulla - Luteal phase - Contralateral - Decellula
## Ampulla - Luteal phase - Ipsilateral - Decellularized - Ampulla - Luteal phase - Contralateral - Dec
## Ampulla - Luteal phase - Ipsilateral - Native - Ampulla - Luteal phase - Contralateral - Decellulari
## Isthmus - Follicular phase - Contralateral - Decellularized - Ampulla - Luteal phase - Contralateral
## Isthmus - Follicular phase - Contralateral - Native - Ampulla - Luteal phase - Contralateral - Decell
## Isthmus - Follicular phase - Ipsilateral - Decellularized - Ampulla - Luteal phase - Contralateral -
## Isthmus - Follicular phase - Ipsilateral - Native - Ampulla - Luteal phase - Contralateral - Decellul
## Isthmus - Luteal phase - Contralateral - Decellularized - Ampulla - Luteal phase - Contralateral - De
## Isthmus - Luteal phase - Contralateral - Native - Ampulla - Luteal phase - Contralateral - Decellula
## Isthmus - Luteal phase - Ipsilateral - Decellularized - Ampulla - Luteal phase - Contralateral - Dec
## Isthmus - Luteal phase - Ipsilateral - Native - Ampulla - Luteal phase - Contralateral - Decellulari
## Ampulla - Luteal phase - Ipsilateral - Decellularized - Ampulla - Luteal phase - Contralateral - Nati
## Ampulla - Luteal phase - Ipsilateral - Native - Ampulla - Luteal phase - Contralateral - Native == 0
## Isthmus - Follicular phase - Contralateral - Decellularized - Ampulla - Luteal phase - Contralateral
## Isthmus - Follicular phase - Contralateral - Native - Ampulla - Luteal phase - Contralateral - Native
## Isthmus - Follicular phase - Ipsilateral - Decellularized - Ampulla - Luteal phase - Contralateral -
## Isthmus - Follicular phase - Ipsilateral - Native - Ampulla - Luteal phase - Contralateral - Native =
## Isthmus - Luteal phase - Contralateral - Decellularized - Ampulla - Luteal phase - Contralateral - Na
## Isthmus - Luteal phase - Contralateral - Native - Ampulla - Luteal phase - Contralateral - Native ==
## Isthmus - Luteal phase - Ipsilateral - Decellularized - Ampulla - Luteal phase - Contralateral - Nati
## Isthmus - Luteal phase - Ipsilateral - Native - Ampulla - Luteal phase - Contralateral - Native == 0
## Ampulla - Luteal phase - Ipsilateral - Native - Ampulla - Luteal phase - Ipsilateral - Decellularize
## Isthmus - Follicular phase - Contralateral - Decellularized - Ampulla - Luteal phase - Ipsilateral -
## Isthmus - Follicular phase - Contralateral - Native - Ampulla - Luteal phase - Ipsilateral - Decellul
## Isthmus - Follicular phase - Ipsilateral - Decellularized - Ampulla - Luteal phase - Ipsilateral - De
## Isthmus - Follicular phase - Ipsilateral - Native - Ampulla - Luteal phase - Ipsilateral - Decellula
## Isthmus - Luteal phase - Contralateral - Decellularized - Ampulla - Luteal phase - Ipsilateral - Dec
## Isthmus - Luteal phase - Contralateral - Native - Ampulla - Luteal phase - Ipsilateral - Decellulari
## Isthmus - Luteal phase - Ipsilateral - Decellularized - Ampulla - Luteal phase - Ipsilateral - Decell
## Isthmus - Luteal phase - Ipsilateral - Native - Ampulla - Luteal phase - Ipsilateral - Decellularize
## Isthmus - Follicular phase - Contralateral - Decellularized - Ampulla - Luteal phase - Ipsilateral -
## Isthmus - Follicular phase - Contralateral - Native - Ampulla - Luteal phase - Ipsilateral - Native =
## Isthmus - Follicular phase - Ipsilateral - Decellularized - Ampulla - Luteal phase - Ipsilateral - Na
## Isthmus - Follicular phase - Ipsilateral - Native - Ampulla - Luteal phase - Ipsilateral - Native ==
## Isthmus - Luteal phase - Contralateral - Decellularized - Ampulla - Luteal phase - Ipsilateral - Nati
## Isthmus - Luteal phase - Contralateral - Native - Ampulla - Luteal phase - Ipsilateral - Native == 0
## Isthmus - Luteal phase - Ipsilateral - Decellularized - Ampulla - Luteal phase - Ipsilateral - Native
```

```
## Isthmus - Luteal phase - Ipsilateral - Native - Ampulla - Luteal phase - Ipsilateral - Native == 0
## Isthmus - Follicular phase - Contralateral - Native - Isthmus - Follicular phase - Contralateral - De
## Isthmus - Follicular phase - Ipsilateral - Decellularized - Isthmus - Follicular phase - Contralatera
## Isthmus - Follicular phase - Ipsilateral - Native - Isthmus - Follicular phase - Contralateral - Dec
## Isthmus - Luteal phase - Contralateral - Decellularized - Isthmus - Follicular phase - Contralateral
## Isthmus - Luteal phase - Contralateral - Native - Isthmus - Follicular phase - Contralateral - Decell
## Isthmus - Luteal phase - Ipsilateral - Decellularized - Isthmus - Follicular phase - Contralateral -
## Isthmus - Luteal phase - Ipsilateral - Native - Isthmus - Follicular phase - Contralateral - Decellu
## Isthmus - Follicular phase - Ipsilateral - Decellularized - Isthmus - Follicular phase - Contralatera
## Isthmus - Follicular phase - Ipsilateral - Native - Isthmus - Follicular phase - Contralateral - Nat
## Isthmus - Luteal phase - Contralateral - Decellularized - Isthmus - Follicular phase - Contralateral
## Isthmus - Luteal phase - Contralateral - Native - Isthmus - Follicular phase - Contralateral - Native
## Isthmus - Luteal phase - Ipsilateral - Decellularized - Isthmus - Follicular phase - Contralateral -
## Isthmus - Luteal phase - Ipsilateral - Native - Isthmus - Follicular phase - Contralateral - Native =
## Isthmus - Follicular phase - Ipsilateral - Native - Isthmus - Follicular phase - Ipsilateral - Decell
## Isthmus - Luteal phase - Contralateral - Decellularized - Isthmus - Follicular phase - Ipsilateral -
## Isthmus - Luteal phase - Contralateral - Native - Isthmus - Follicular phase - Ipsilateral - Decellu
## Isthmus - Luteal phase - Ipsilateral - Decellularized - Isthmus - Follicular phase - Ipsilateral - De
## Isthmus - Luteal phase - Ipsilateral - Native - Isthmus - Follicular phase - Ipsilateral - Decellula
## Isthmus - Luteal phase - Contralateral - Decellularized - Isthmus - Follicular phase - Ipsilateral -
## Isthmus - Luteal phase - Contralateral - Native - Isthmus - Follicular phase - Ipsilateral - Native =
## Isthmus - Luteal phase - Ipsilateral - Decellularized - Isthmus - Follicular phase - Ipsilateral - Na
## Isthmus - Luteal phase - Ipsilateral - Native - Isthmus - Follicular phase - Ipsilateral - Native ==
## Isthmus - Luteal phase - Contralateral - Native - Isthmus - Luteal phase - Contralateral - Decellula
## Isthmus - Luteal phase - Ipsilateral - Decellularized - Isthmus - Luteal phase - Contralateral - Dece
## Isthmus - Luteal phase - Ipsilateral - Native - Isthmus - Luteal phase - Contralateral - Decellulari
## Isthmus - Luteal phase - Ipsilateral - Decellularized - Isthmus - Luteal phase - Contralateral - Nati
## Isthmus - Luteal phase - Ipsilateral - Native - Isthmus - Luteal phase - Contralateral - Native == 0
## Isthmus - Luteal phase - Ipsilateral - Native - Isthmus - Luteal phase - Ipsilateral - Decellularize
##
## Ampulla - Follicular phase - Contralateral - Native - Ampulla - Follicular phase - Contralateral - De
## Ampulla - Follicular phase - Ipsilateral - Decellularized - Ampulla - Follicular phase - Contralatera
## Ampulla - Follicular phase - Ipsilateral - Native - Ampulla - Follicular phase - Contralateral - Dec
## Ampulla - Luteal phase - Contralateral - Decellularized - Ampulla - Follicular phase - Contralateral
## Ampulla - Luteal phase - Contralateral - Native - Ampulla - Follicular phase - Contralateral - Decell
## Ampulla - Luteal phase - Ipsilateral - Decellularized - Ampulla - Follicular phase - Contralateral -
## Ampulla - Luteal phase - Ipsilateral - Native - Ampulla - Follicular phase - Contralateral - Decellu
## Isthmus - Follicular phase - Contralateral - Decellularized - Ampulla - Follicular phase - Contralate
## Isthmus - Follicular phase - Contralateral - Native - Ampulla - Follicular phase - Contralateral - De
## Isthmus - Follicular phase - Ipsilateral - Decellularized - Ampulla - Follicular phase - Contralatera
## Isthmus - Follicular phase - Ipsilateral - Native - Ampulla - Follicular phase - Contralateral - Dec
## Isthmus - Luteal phase - Contralateral - Decellularized - Ampulla - Follicular phase - Contralateral
## Isthmus - Luteal phase - Contralateral - Native - Ampulla - Follicular phase - Contralateral - Decell
## Isthmus - Luteal phase - Ipsilateral - Decellularized - Ampulla - Follicular phase - Contralateral -
## Isthmus - Luteal phase - Ipsilateral - Native - Ampulla - Follicular phase - Contralateral - Decellu
## Ampulla - Follicular phase - Ipsilateral - Decellularized - Ampulla - Follicular phase - Contralatera
## Ampulla - Follicular phase - Ipsilateral - Native - Ampulla - Follicular phase - Contralateral - Nati
## Ampulla - Luteal phase - Contralateral - Decellularized - Ampulla - Follicular phase - Contralateral
## Ampulla - Luteal phase - Contralateral - Native - Ampulla - Follicular phase - Contralateral - Native
## Ampulla - Luteal phase - Ipsilateral - Decellularized - Ampulla - Follicular phase - Contralateral -
## Ampulla - Luteal phase - Ipsilateral - Native - Ampulla - Follicular phase - Contralateral - Native =
## Isthmus - Follicular phase - Contralateral - Decellularized - Ampulla - Follicular phase - Contralate
## Isthmus - Follicular phase - Contralateral - Native - Ampulla - Follicular phase - Contralateral - Na
## Isthmus - Follicular phase - Ipsilateral - Decellularized - Ampulla - Follicular phase - Contralatera
```

```
## Isthmus - Follicular phase - Ipsilateral - Native - Ampulla - Follicular phase - Contralateral - Nati
## Isthmus - Luteal phase - Contralateral - Decellularized - Ampulla - Follicular phase - Contralateral
## Isthmus - Luteal phase - Contralateral - Native - Ampulla - Follicular phase - Contralateral - Nativ
## Isthmus - Luteal phase - Ipsilateral - Decellularized - Ampulla - Follicular phase - Contralateral -
## Isthmus - Luteal phase - Ipsilateral - Native - Ampulla - Follicular phase - Contralateral - Native =
## Ampulla - Follicular phase - Ipsilateral - Native - Ampulla - Follicular phase - Ipsilateral - Decel
## Ampulla - Luteal phase - Contralateral - Decellularized - Ampulla - Follicular phase - Ipsilateral -
## Ampulla - Luteal phase - Contralateral - Native - Ampulla - Follicular phase - Ipsilateral - Decellu
## Ampulla - Luteal phase - Ipsilateral - Decellularized - Ampulla - Follicular phase - Ipsilateral - De
## Ampulla - Luteal phase - Ipsilateral - Native - Ampulla - Follicular phase - Ipsilateral - Decellula
## Isthmus - Follicular phase - Contralateral - Decellularized - Ampulla - Follicular phase - Ipsilatera
## Isthmus - Follicular phase - Contralateral - Native - Ampulla - Follicular phase - Ipsilateral - Dec
## Isthmus - Follicular phase - Ipsilateral - Decellularized - Ampulla - Follicular phase - Ipsilateral
## Isthmus - Follicular phase - Ipsilateral - Native - Ampulla - Follicular phase - Ipsilateral - Decel
## Isthmus - Luteal phase - Contralateral - Decellularized - Ampulla - Follicular phase - Ipsilateral -
## Isthmus - Luteal phase - Contralateral - Native - Ampulla - Follicular phase - Ipsilateral - Decellu
## Isthmus - Luteal phase - Ipsilateral - Decellularized - Ampulla - Follicular phase - Ipsilateral - De
## Isthmus - Luteal phase - Ipsilateral - Native - Ampulla - Follicular phase - Ipsilateral - Decellula
## Ampulla - Luteal phase - Contralateral - Decellularized - Ampulla - Follicular phase - Ipsilateral -
## Ampulla - Luteal phase - Contralateral - Native - Ampulla - Follicular phase - Ipsilateral - Native =
## Ampulla - Luteal phase - Ipsilateral - Decellularized - Ampulla - Follicular phase - Ipsilateral - Na
## Ampulla - Luteal phase - Ipsilateral - Native - Ampulla - Follicular phase - Ipsilateral - Native ==
## Isthmus - Follicular phase - Contralateral - Decellularized - Ampulla - Follicular phase - Ipsilatera
## Isthmus - Follicular phase - Contralateral - Native - Ampulla - Follicular phase - Ipsilateral - Nati
## Isthmus - Follicular phase - Ipsilateral - Decellularized - Ampulla - Follicular phase - Ipsilateral
## Isthmus - Follicular phase - Ipsilateral - Native - Ampulla - Follicular phase - Ipsilateral - Nativ
## Isthmus - Luteal phase - Contralateral - Decellularized - Ampulla - Follicular phase - Ipsilateral -
## Isthmus - Luteal phase - Contralateral - Native - Ampulla - Follicular phase - Ipsilateral - Native =
## Isthmus - Luteal phase - Ipsilateral - Decellularized - Ampulla - Follicular phase - Ipsilateral - Na
## Isthmus - Luteal phase - Ipsilateral - Native - Ampulla - Follicular phase - Ipsilateral - Native ==
## Ampulla - Luteal phase - Contralateral - Native - Ampulla - Luteal phase - Contralateral - Decellula
## Ampulla - Luteal phase - Ipsilateral - Decellularized - Ampulla - Luteal phase - Contralateral - Dec
## Ampulla - Luteal phase - Ipsilateral - Native - Ampulla - Luteal phase - Contralateral - Decellulari
## Isthmus - Follicular phase - Contralateral - Decellularized - Ampulla - Luteal phase - Contralateral
## Isthmus - Follicular phase - Contralateral - Native - Ampulla - Luteal phase - Contralateral - Decel
## Isthmus - Follicular phase - Ipsilateral - Decellularized - Ampulla - Luteal phase - Contralateral -
## Isthmus - Follicular phase - Ipsilateral - Native - Ampulla - Luteal phase - Contralateral - Decellu
## Isthmus - Luteal phase - Contralateral - Decellularized - Ampulla - Luteal phase - Contralateral - De
## Isthmus - Luteal phase - Contralateral - Native - Ampulla - Luteal phase - Contralateral - Decellula
## Isthmus - Luteal phase - Ipsilateral - Decellularized - Ampulla - Luteal phase - Contralateral - Dec
## Isthmus - Luteal phase - Ipsilateral - Native - Ampulla - Luteal phase - Contralateral - Decellulari
## Ampulla - Luteal phase - Ipsilateral - Decellularized - Ampulla - Luteal phase - Contralateral - Nati
## Ampulla - Luteal phase - Ipsilateral - Native - Ampulla - Luteal phase - Contralateral - Native == 0
## Isthmus - Follicular phase - Contralateral - Decellularized - Ampulla - Luteal phase - Contralateral
## Isthmus - Follicular phase - Contralateral - Native - Ampulla - Luteal phase - Contralateral - Native
## Isthmus - Follicular phase - Ipsilateral - Decellularized - Ampulla - Luteal phase - Contralateral -
## Isthmus - Follicular phase - Ipsilateral - Native - Ampulla - Luteal phase - Contralateral - Native =
## Isthmus - Luteal phase - Contralateral - Decellularized - Ampulla - Luteal phase - Contralateral - Na
## Isthmus - Luteal phase - Contralateral - Native - Ampulla - Luteal phase - Contralateral - Native ==
## Isthmus - Luteal phase - Ipsilateral - Decellularized - Ampulla - Luteal phase - Contralateral - Nati
## Isthmus - Luteal phase - Ipsilateral - Native - Ampulla - Luteal phase - Contralateral - Native == 0
## Ampulla - Luteal phase - Ipsilateral - Native - Ampulla - Luteal phase - Ipsilateral - Decellularized
## Isthmus - Follicular phase - Contralateral - Decellularized - Ampulla - Luteal phase - Ipsilateral -
## Isthmus - Follicular phase - Contralateral - Native - Ampulla - Luteal phase - Ipsilateral - Decellul
```

```
## Isthmus - Follicular phase - Ipsilateral - Decellularized - Ampulla - Luteal phase - Ipsilateral - De
## Isthmus - Follicular phase - Ipsilateral - Native - Ampulla - Luteal phase - Ipsilateral - Decellula
## Isthmus - Luteal phase - Contralateral - Decellularized - Ampulla - Luteal phase - Ipsilateral - Dec
## Isthmus - Luteal phase - Contralateral - Native - Ampulla - Luteal phase - Ipsilateral - Decellulari
## Isthmus - Luteal phase - Ipsilateral - Decellularized - Ampulla - Luteal phase - Ipsilateral - Decel
## Isthmus - Luteal phase - Ipsilateral - Native - Ampulla - Luteal phase - Ipsilateral - Decellularize
## Isthmus - Follicular phase - Contralateral - Decellularized - Ampulla - Luteal phase - Ipsilateral -
## Isthmus - Follicular phase - Contralateral - Native - Ampulla - Luteal phase - Ipsilateral - Native =
## Isthmus - Follicular phase - Ipsilateral - Decellularized - Ampulla - Luteal phase - Ipsilateral - Na
## Isthmus - Follicular phase - Ipsilateral - Native - Ampulla - Luteal phase - Ipsilateral - Native ==
## Isthmus - Luteal phase - Contralateral - Decellularized - Ampulla - Luteal phase - Ipsilateral - Nati
## Isthmus - Luteal phase - Contralateral - Native - Ampulla - Luteal phase - Ipsilateral - Native == 0
## Isthmus - Luteal phase - Ipsilateral - Decellularized - Ampulla - Luteal phase - Ipsilateral - Native
## Isthmus - Luteal phase - Ipsilateral - Native - Ampulla - Luteal phase - Ipsilateral - Native == 0
## Isthmus - Follicular phase - Contralateral - Native - Isthmus - Follicular phase - Contralateral - De
## Isthmus - Follicular phase - Ipsilateral - Decellularized - Isthmus - Follicular phase - Contralatera
## Isthmus - Follicular phase - Ipsilateral - Native - Isthmus - Follicular phase - Contralateral - Dec
## Isthmus - Luteal phase - Contralateral - Decellularized - Isthmus - Follicular phase - Contralateral
## Isthmus - Luteal phase - Contralateral - Native - Isthmus - Follicular phase - Contralateral - Decel
## Isthmus - Luteal phase - Ipsilateral - Decellularized - Isthmus - Follicular phase - Contralateral -
## Isthmus - Luteal phase - Ipsilateral - Native - Isthmus - Follicular phase - Contralateral - Decellu
## Isthmus - Follicular phase - Ipsilateral - Decellularized - Isthmus - Follicular phase - Contralatera
## Isthmus - Follicular phase - Ipsilateral - Native - Isthmus - Follicular phase - Contralateral - Nati
## Isthmus - Luteal phase - Contralateral - Decellularized - Isthmus - Follicular phase - Contralateral
## Isthmus - Luteal phase - Contralateral - Native - Isthmus - Follicular phase - Contralateral - Native
## Isthmus - Luteal phase - Ipsilateral - Decellularized - Isthmus - Follicular phase - Contralateral -
## Isthmus - Luteal phase - Ipsilateral - Native - Isthmus - Follicular phase - Contralateral - Native =
## Isthmus - Follicular phase - Ipsilateral - Native - Isthmus - Follicular phase - Ipsilateral - Decel
## Isthmus - Luteal phase - Contralateral - Decellularized - Isthmus - Follicular phase - Ipsilateral -
## Isthmus - Luteal phase - Contralateral - Native - Isthmus - Follicular phase - Ipsilateral - Decellu
## Isthmus - Luteal phase - Ipsilateral - Decellularized - Isthmus - Follicular phase - Ipsilateral - De
## Isthmus - Luteal phase - Ipsilateral - Native - Isthmus - Follicular phase - Ipsilateral - Decellula
## Isthmus - Luteal phase - Contralateral - Decellularized - Isthmus - Follicular phase - Ipsilateral -
## Isthmus - Luteal phase - Contralateral - Native - Isthmus - Follicular phase - Ipsilateral - Native =
## Isthmus - Luteal phase - Ipsilateral - Decellularized - Isthmus - Follicular phase - Ipsilateral - Na
## Isthmus - Luteal phase - Ipsilateral - Native - Isthmus - Follicular phase - Ipsilateral - Native ==
## Isthmus - Luteal phase - Contralateral - Native - Isthmus - Luteal phase - Contralateral - Decellula
## Isthmus - Luteal phase - Ipsilateral - Decellularized - Isthmus - Luteal phase - Contralateral - Dec
## Isthmus - Luteal phase - Ipsilateral - Native - Isthmus - Luteal phase - Contralateral - Decellulari
## Isthmus - Luteal phase - Ipsilateral - Decellularized - Isthmus - Luteal phase - Contralateral - Nati
## Isthmus - Luteal phase - Ipsilateral - Native - Isthmus - Luteal phase - Contralateral - Native == 0
## Isthmus - Luteal phase - Ipsilateral - Native - Isthmus - Luteal phase - Ipsilateral - Decellularize
##
## Ampulla - Follicular phase - Contralateral - Native - Ampulla - Follicular phase - Contralateral - De
## Ampulla - Follicular phase - Ipsilateral - Decellularized - Ampulla - Follicular phase - Contralatera
## Ampulla - Follicular phase - Ipsilateral - Native - Ampulla - Follicular phase - Contralateral - Dec
## Ampulla - Luteal phase - Contralateral - Decellularized - Ampulla - Follicular phase - Contralateral
## Ampulla - Luteal phase - Contralateral - Native - Ampulla - Follicular phase - Contralateral - Decel
## Ampulla - Luteal phase - Ipsilateral - Decellularized - Ampulla - Follicular phase - Contralateral -
## Ampulla - Luteal phase - Ipsilateral - Native - Ampulla - Follicular phase - Contralateral - Decellu
## Isthmus - Follicular phase - Contralateral - Decellularized - Ampulla - Follicular phase - Contralate
## Isthmus - Follicular phase - Contralateral - Native - Ampulla - Follicular phase - Contralateral - De
## Isthmus - Follicular phase - Ipsilateral - Decellularized - Ampulla - Follicular phase - Contralatera
## Isthmus - Follicular phase - Ipsilateral - Native - Ampulla - Follicular phase - Contralateral - Dec
```

```
## Isthmus - Luteal phase - Contralateral - Decellularized - Ampulla - Follicular phase - Contralateral
## Isthmus - Luteal phase - Contralateral - Native - Ampulla - Follicular phase - Contralateral - Decell
## Isthmus - Luteal phase - Ipsilateral - Decellularized - Ampulla - Follicular phase - Contralateral -
## Isthmus - Luteal phase - Ipsilateral - Native - Ampulla - Follicular phase - Contralateral - Decellu
## Ampulla - Follicular phase - Ipsilateral - Decellularized - Ampulla - Follicular phase - Contralatera
## Ampulla - Follicular phase - Ipsilateral - Native - Ampulla - Follicular phase - Contralateral - Nat
## Ampulla - Luteal phase - Contralateral - Decellularized - Ampulla - Follicular phase - Contralateral
## Ampulla - Luteal phase - Contralateral - Native - Ampulla - Follicular phase - Contralateral - Nativ
## Ampulla - Luteal phase - Ipsilateral - Decellularized - Ampulla - Follicular phase - Contralateral -
## Ampulla - Luteal phase - Ipsilateral - Native - Ampulla - Follicular phase - Contralateral - Native =
## Isthmus - Follicular phase - Contralateral - Decellularized - Ampulla - Follicular phase - Contralate
## Isthmus - Follicular phase - Contralateral - Native - Ampulla - Follicular phase - Contralateral - Na
## Isthmus - Follicular phase - Ipsilateral - Decellularized - Ampulla - Follicular phase - Contralatera
## Isthmus - Follicular phase - Ipsilateral - Native - Ampulla - Follicular phase - Contralateral - Nat
## Isthmus - Luteal phase - Contralateral - Decellularized - Ampulla - Follicular phase - Contralateral
## Isthmus - Luteal phase - Contralateral - Native - Ampulla - Follicular phase - Contralateral - Nativ
## Isthmus - Luteal phase - Ipsilateral - Decellularized - Ampulla - Follicular phase - Contralateral -
## Isthmus - Luteal phase - Ipsilateral - Native - Ampulla - Follicular phase - Contralateral - Native =
## Ampulla - Follicular phase - Ipsilateral - Native - Ampulla - Follicular phase - Ipsilateral - Decell
## Ampulla - Luteal phase - Contralateral - Decellularized - Ampulla - Follicular phase - Ipsilateral -
## Ampulla - Luteal phase - Contralateral - Native - Ampulla - Follicular phase - Ipsilateral - Decellu
## Ampulla - Luteal phase - Ipsilateral - Decellularized - Ampulla - Follicular phase - Ipsilateral - D
## Ampulla - Luteal phase - Ipsilateral - Native - Ampulla - Follicular phase - Ipsilateral - Decellula
## Isthmus - Follicular phase - Contralateral - Decellularized - Ampulla - Follicular phase - Ipsilatera
## Isthmus - Follicular phase - Contralateral - Native - Ampulla - Follicular phase - Ipsilateral - Dec
## Isthmus - Follicular phase - Ipsilateral - Decellularized - Ampulla - Follicular phase - Ipsilateral
## Isthmus - Follicular phase - Ipsilateral - Native - Ampulla - Follicular phase - Ipsilateral - Decell
## Isthmus - Luteal phase - Contralateral - Decellularized - Ampulla - Follicular phase - Ipsilateral -
## Isthmus - Luteal phase - Contralateral - Native - Ampulla - Follicular phase - Ipsilateral - Decellu
## Isthmus - Luteal phase - Ipsilateral - Decellularized - Ampulla - Follicular phase - Ipsilateral - D
## Isthmus - Luteal phase - Ipsilateral - Native - Ampulla - Follicular phase - Ipsilateral - Decellula
## Ampulla - Luteal phase - Contralateral - Decellularized - Ampulla - Follicular phase - Ipsilateral -
## Ampulla - Luteal phase - Contralateral - Native - Ampulla - Follicular phase - Ipsilateral - Native =
## Ampulla - Luteal phase - Ipsilateral - Decellularized - Ampulla - Follicular phase - Ipsilateral - Na
## Ampulla - Luteal phase - Ipsilateral - Native - Ampulla - Follicular phase - Ipsilateral - Native ==
## Isthmus - Follicular phase - Contralateral - Decellularized - Ampulla - Follicular phase - Ipsilatera
## Isthmus - Follicular phase - Contralateral - Native - Ampulla - Follicular phase - Ipsilateral - Nat
## Isthmus - Follicular phase - Ipsilateral - Decellularized - Ampulla - Follicular phase - Ipsilateral
## Isthmus - Follicular phase - Ipsilateral - Native - Ampulla - Follicular phase - Ipsilateral - Nativ
## Isthmus - Luteal phase - Contralateral - Decellularized - Ampulla - Follicular phase - Ipsilateral -
## Isthmus - Luteal phase - Contralateral - Native - Ampulla - Follicular phase - Ipsilateral - Native =
## Isthmus - Luteal phase - Ipsilateral - Decellularized - Ampulla - Follicular phase - Ipsilateral - Na
## Isthmus - Luteal phase - Ipsilateral - Native - Ampulla - Follicular phase - Ipsilateral - Native ==
## Ampulla - Luteal phase - Contralateral - Native - Ampulla - Luteal phase - Contralateral - Decellula
## Ampulla - Luteal phase - Ipsilateral - Decellularized - Ampulla - Luteal phase - Contralateral - Dec
## Ampulla - Luteal phase - Ipsilateral - Native - Ampulla - Luteal phase - Contralateral - Decellulari
## Isthmus - Follicular phase - Contralateral - Decellularized - Ampulla - Luteal phase - Contralateral
## Isthmus - Follicular phase - Contralateral - Native - Ampulla - Luteal phase - Contralateral - Decel
## Isthmus - Follicular phase - Ipsilateral - Decellularized - Ampulla - Luteal phase - Contralateral -
## Isthmus - Follicular phase - Ipsilateral - Native - Ampulla - Luteal phase - Contralateral - Decellu
## Isthmus - Luteal phase - Contralateral - Decellularized - Ampulla - Luteal phase - Contralateral - De
## Isthmus - Luteal phase - Contralateral - Native - Ampulla - Luteal phase - Contralateral - Decellula
## Isthmus - Luteal phase - Ipsilateral - Decellularized - Ampulla - Luteal phase - Contralateral - Dec
## Isthmus - Luteal phase - Ipsilateral - Native - Ampulla - Luteal phase - Contralateral - Decellulari
```

```
## Ampulla - Luteal phase - Ipsilateral - Decellularized - Ampulla - Luteal phase - Contralateral - Nati
## Ampulla - Luteal phase - Ipsilateral - Native - Ampulla - Luteal phase - Contralateral - Native == 0
## Isthmus - Follicular phase - Contralateral - Decellularized - Ampulla - Luteal phase - Contralateral
## Isthmus - Follicular phase - Contralateral - Native - Ampulla - Luteal phase - Contralateral - Native
## Isthmus - Follicular phase - Ipsilateral - Decellularized - Ampulla - Luteal phase - Contralateral -
## Isthmus - Follicular phase - Ipsilateral - Native - Ampulla - Luteal phase - Contralateral - Native =
## Isthmus - Luteal phase - Contralateral - Decellularized - Ampulla - Luteal phase - Contralateral - Na
## Isthmus - Luteal phase - Contralateral - Native - Ampulla - Luteal phase - Contralateral - Native ==
## Isthmus - Luteal phase - Ipsilateral - Decellularized - Ampulla - Luteal phase - Contralateral - Nati
## Isthmus - Luteal phase - Ipsilateral - Native - Ampulla - Luteal phase - Contralateral - Native == 0
## Ampulla - Luteal phase - Ipsilateral - Native - Ampulla - Luteal phase - Ipsilateral - Decellularized
## Isthmus - Follicular phase - Contralateral - Decellularized - Ampulla - Luteal phase - Ipsilateral -
## Isthmus - Follicular phase - Contralateral - Native - Ampulla - Luteal phase - Ipsilateral - Decellu
## Isthmus - Follicular phase - Ipsilateral - Decellularized - Ampulla - Luteal phase - Ipsilateral - De
## Isthmus - Follicular phase - Ipsilateral - Native - Ampulla - Luteal phase - Ipsilateral - Decellula
## Isthmus - Luteal phase - Contralateral - Decellularized - Ampulla - Luteal phase - Ipsilateral - Dec
## Isthmus - Luteal phase - Contralateral - Native - Ampulla - Luteal phase - Ipsilateral - Decellulari
## Isthmus - Luteal phase - Ipsilateral - Decellularized - Ampulla - Luteal phase - Ipsilateral - Decel
## Isthmus - Luteal phase - Ipsilateral - Native - Ampulla - Luteal phase - Ipsilateral - Decellularized
## Isthmus - Follicular phase - Contralateral - Decellularized - Ampulla - Luteal phase - Ipsilateral -
## Isthmus - Follicular phase - Contralateral - Native - Ampulla - Luteal phase - Ipsilateral - Native =
## Isthmus - Follicular phase - Ipsilateral - Decellularized - Ampulla - Luteal phase - Ipsilateral - Na
## Isthmus - Follicular phase - Ipsilateral - Native - Ampulla - Luteal phase - Ipsilateral - Native ==
## Isthmus - Luteal phase - Contralateral - Decellularized - Ampulla - Luteal phase - Ipsilateral - Nati
## Isthmus - Luteal phase - Contralateral - Native - Ampulla - Luteal phase - Ipsilateral - Native == 0
## Isthmus - Luteal phase - Ipsilateral - Decellularized - Ampulla - Luteal phase - Ipsilateral - Native
## Isthmus - Luteal phase - Ipsilateral - Native - Ampulla - Luteal phase - Ipsilateral - Native == 0
## Isthmus - Follicular phase - Contralateral - Native - Isthmus - Follicular phase - Contralateral - De
## Isthmus - Follicular phase - Ipsilateral - Decellularized - Isthmus - Follicular phase - Contralatera
## Isthmus - Follicular phase - Ipsilateral - Native - Isthmus - Follicular phase - Contralateral - Dec
## Isthmus - Luteal phase - Contralateral - Decellularized - Isthmus - Follicular phase - Contralateral
## Isthmus - Luteal phase - Contralateral - Native - Isthmus - Follicular phase - Contralateral - Decel
## Isthmus - Luteal phase - Ipsilateral - Decellularized - Isthmus - Follicular phase - Contralateral -
## Isthmus - Luteal phase - Ipsilateral - Native - Isthmus - Follicular phase - Contralateral - Decellu
## Isthmus - Follicular phase - Ipsilateral - Decellularized - Isthmus - Follicular phase - Contralatera
## Isthmus - Follicular phase - Ipsilateral - Native - Isthmus - Follicular phase - Contralateral - Nati
## Isthmus - Luteal phase - Contralateral - Decellularized - Isthmus - Follicular phase - Contralateral
## Isthmus - Luteal phase - Contralateral - Native - Isthmus - Follicular phase - Contralateral - Native
## Isthmus - Luteal phase - Ipsilateral - Decellularized - Isthmus - Follicular phase - Contralateral -
## Isthmus - Luteal phase - Ipsilateral - Native - Isthmus - Follicular phase - Contralateral - Native =
## Isthmus - Follicular phase - Ipsilateral - Native - Isthmus - Follicular phase - Ipsilateral - Decell
## Isthmus - Luteal phase - Contralateral - Decellularized - Isthmus - Follicular phase - Ipsilateral -
## Isthmus - Luteal phase - Contralateral - Native - Isthmus - Follicular phase - Ipsilateral - Decellul
## Isthmus - Luteal phase - Ipsilateral - Decellularized - Isthmus - Follicular phase - Ipsilateral - De
## Isthmus - Luteal phase - Ipsilateral - Native - Isthmus - Follicular phase - Ipsilateral - Decellula
## Isthmus - Luteal phase - Contralateral - Decellularized - Isthmus - Follicular phase - Ipsilateral -
## Isthmus - Luteal phase - Contralateral - Native - Isthmus - Follicular phase - Ipsilateral - Native =
## Isthmus - Luteal phase - Ipsilateral - Decellularized - Isthmus - Follicular phase - Ipsilateral - Na
## Isthmus - Luteal phase - Ipsilateral - Native - Isthmus - Follicular phase - Ipsilateral - Native ==
## Isthmus - Luteal phase - Contralateral - Native - Isthmus - Luteal phase - Contralateral - Decellula
## Isthmus - Luteal phase - Ipsilateral - Decellularized - Isthmus - Luteal phase - Contralateral - Dec
## Isthmus - Luteal phase - Ipsilateral - Native - Isthmus - Luteal phase - Contralateral - Decellulari
## Isthmus - Luteal phase - Ipsilateral - Decellularized - Isthmus - Luteal phase - Contralateral - Nati
## Isthmus - Luteal phase - Ipsilateral - Native - Isthmus - Luteal phase - Contralateral - Native == 0
```

```
## Isthmus - Luteal phase - Ipsilateral - Native - Isthmus - Luteal phase - Ipsilateral - Decellularized
## ---
## Signif. codes:  0 '***' 0.001 '**' 0.01 '*' 0.05 '.' 0.1 ' ' 1
## (Adjusted p values reported -- single-step method)
```

```
################### Porosity ######################################

# Read the dataset from a CSV file
df <- read.csv("Oviduct_porosity.csv")

# Summary data

summary_df <- df %>%
  group_by(Cellular, Tissue, Position, Estrus_cycle) %>%
  summarise(mean_Porosity = mean(Porosity),
            sd_Porosity = sd(Porosity))
```

```
## `summarise()` has grouped output by 'Cellular', 'Tissue', 'Position'. You can
## override using the `.groups` argument.
```

```
# Plot

# Read the dataset from a CSV file
df <- read.csv("Oviduct_porosity.csv")

png(filename="Oiduct_porosity2.png",
    width = 15, height = 10, units = "in",
    res = 600)

par(mgp=c(1.5,0.5,0),
    mar = c(4, #bottom
            4, #left
            3, #top
            2),
    cex.main = 0.8,
    family = "sans" #right
)

# Update the labels for Position, Estrus_cycle, and Tissue
df$Position <- factor(df$Position, labels = c("Contralateral", "Ipsilateral"))
df$Estrus_cycle <- factor(df$Estrus_cycle, labels = c("Follicular phase", "Luteal phase"))
df$Tissue <- factor(df$Tissue, labels = c("Ampulla", "Isthmus"))

# Create a combined factor variable for Estrus_cycle, Cellular, Position, and Tissue
df$Combined <- factor(paste(df$Estrus_cycle, df$Cellular, df$Position, df$Tissue, sep = " "),
                      levels = c("Follicular phase Native Contralateral Ampulla",
                                 "Follicular phase Decellularized Contralateral Ampulla",
                                 "Luteal phase Native Contralateral Ampulla",
                                 "Luteal phase Decellularized Contralateral Ampulla",
                                 "Follicular phase Native Ipsilateral Ampulla",
                                 "Follicular phase Decellularized Ipsilateral Ampulla",
                                 "Luteal phase Native Ipsilateral Ampulla",
                                 "Luteal phase Decellularized Ipsilateral Ampulla",
                                 "Follicular phase Native Contralateral Isthmus",
                                 "Follicular phase Decellularized Contralateral Isthmus",
```

```r
                                  "Luteal phase Native Contralateral Isthmus",
                                  "Luteal phase Decellularized Contralateral Isthmus",
                                  "Follicular phase Native Ipsilateral Isthmus",
                                  "Follicular phase Decellularized Ipsilateral Isthmus",
                                  "Luteal phase Native Ipsilateral Isthmus",
                                  "Luteal phase Decellularized Ipsilateral Isthmus"))

# Define fill colors for each factor level
fill_colors <- c("Follicular phase Native Contralateral Ampulla" = viridis_pal(option = "D")(1),
                 "Follicular phase Decellularized Contralateral Ampulla" = alpha(viridis_pal(option = "
                 "Luteal phase Native Contralateral Ampulla" = "#238A8DFF",
                 "Luteal phase Decellularized Contralateral Ampulla" = alpha("#238A8DFF", 0.2),
                 "Follicular phase Native Ipsilateral Ampulla" = viridis_pal(option = "D")(1),
                 "Follicular phase Decellularized Ipsilateral Ampulla" = alpha(viridis_pal(option = "D"
                 "Luteal phase Native Ipsilateral Ampulla" = "#238A8DFF",
                 "Luteal phase Decellularized Ipsilateral Ampulla" = alpha("#238A8DFF", 0.2),
                 "Follicular phase Native Contralateral Isthmus" = viridis_pal(option = "D")(1),
                 "Follicular phase Decellularized Contralateral Isthmus" = alpha(viridis_pal(option = "
                 "Luteal phase Native Contralateral Isthmus" = "#238A8DFF",
                 "Luteal phase Decellularized Contralateral Isthmus" = alpha("#238A8DFF", 0.2),
                 "Follicular phase Native Ipsilateral Isthmus" = viridis_pal(option = "D")(1),
                 "Follicular phase Decellularized Ipsilateral Isthmus" = alpha(viridis_pal(option = "D"
                 "Luteal phase Native Ipsilateral Isthmus" = "#238A8DFF",
                 "Luteal phase Decellularized Ipsilateral Isthmus" = alpha("#238A8DFF", 0.2))

# Create the plot using ggplot
ggplot(data = df, aes(x = Tissue, y = Porosity, fill = Combined)) +
  geom_boxplot(width = 0.7, outlier.shape = NA) +
  geom_jitter(position = position_jitterdodge(), alpha = 0.8) +
  xlab("Tissue") +
  ylab("Estimated Porosity (%)") +
  ggtitle("Boxplot and Individual Data Points of Porosity by Estrus_cycle, Cellular, Position, Tissue")
  scale_fill_manual(values = fill_colors, guide = guide_legend(title = "Combined")) +
  theme_bw() +
  ylim(0, 140) +
  facet_grid(Estrus_cycle ~ Position, space = "free_x", scales = "free_x") +
  theme(panel.grid.major = element_blank(), panel.grid.minor = element_blank(),
        axis.text.x = element_text(size = 11, angle = 0, hjust = 0.5),
        axis.text.y = element_text(size = 12, angle = 0, hjust = 0.5),
        axis.title.x = element_text(size = 18, face = "bold"),
        axis.title.y = element_text(size = 16, face = "bold"),
        legend.position = "right",
        strip.text = element_text(size = 14, face = "bold"))

dev.off()
```

```
## pdf
##   2
```

```r
# Stat

# Read the dataset from a CSV file
df <- read.csv("Oviduct_porosity.csv")
```

```r
# Update the labels for Position, Estrus_cycle, and Tissue
df$Position <- factor(df$Position, labels = c("Contralateral", "Ipsilateral"))
df$Estrus_cycle <- factor(df$Estrus_cycle, labels = c("Follicular phase", "Luteal phase"))
df$Cellular <- factor(df$Cellular, labels = c("Native", "Decellularized"))

# Create a factor variable for the comparisons
df$Comparison <- factor(paste(df$Tissue, df$Estrus_cycle, df$Position, df$Cellular, sep = " - "))

# Perform pairwise comparisons using Tukey's method
model <- lm(Porosity ~ Comparison, data = df)
comp <- glht(model, linfct = mcp(Comparison = "Tukey"))

# Summarize the results
summary(comp)
```

```
## Warning in RET$pfunction("adjusted", ...): Completion with error > abseps

## Warning in RET$pfunction("adjusted", ...): Completion with error > abseps

## Warning in RET$pfunction("adjusted", ...): Completion with error > abseps

## Warning in RET$pfunction("adjusted", ...): Completion with error > abseps

## Warning in RET$pfunction("adjusted", ...): Completion with error > abseps

## Warning in RET$pfunction("adjusted", ...): Completion with error > abseps

## Warning in RET$pfunction("adjusted", ...): Completion with error > abseps
```

```
##
##   Simultaneous Tests for General Linear Hypotheses
##
## Multiple Comparisons of Means: Tukey Contrasts
##
##
## Fit: lm(formula = Porosity ~ Comparison, data = df)
##
## Linear Hypotheses:
##
## Ampulla - Follicular phase - Contralateral - Native - Ampulla - Follicular phase - Contralateral - De
## Ampulla - Follicular phase - Ipsilateral - Decellularized - Ampulla - Follicular phase - Contralatera
## Ampulla - Follicular phase - Ipsilateral - Native - Ampulla - Follicular phase - Contralateral - Dece
## Ampulla - Luteal phase - Contralateral - Decellularized - Ampulla - Follicular phase - Contralateral
## Ampulla - Luteal phase - Contralateral - Native - Ampulla - Follicular phase - Contralateral - Decel
## Ampulla - Luteal phase - Ipsilateral - Decellularized - Ampulla - Follicular phase - Contralateral -
## Ampulla - Luteal phase - Ipsilateral - Native - Ampulla - Follicular phase - Contralateral - Decellu
## Isthmus - Follicular phase - Contralateral - Decellularized - Ampulla - Follicular phase - Contralate
## Isthmus - Follicular phase - Contralateral - Native - Ampulla - Follicular phase - Contralateral - De
## Isthmus - Follicular phase - Ipsilateral - Decellularized - Ampulla - Follicular phase - Contralatera
## Isthmus - Follicular phase - Ipsilateral - Native - Ampulla - Follicular phase - Contralateral - Dece
## Isthmus - Luteal phase - Contralateral - Decellularized - Ampulla - Follicular phase - Contralateral
## Isthmus - Luteal phase - Contralateral - Native - Ampulla - Follicular phase - Contralateral - Decel
## Isthmus - Luteal phase - Ipsilateral - Decellularized - Ampulla - Follicular phase - Contralateral -
## Isthmus - Luteal phase - Ipsilateral - Native - Ampulla - Follicular phase - Contralateral - Decellu
```

```
## Ampulla - Follicular phase - Ipsilateral - Decellularized - Ampulla - Follicular phase - Contralatera
## Ampulla - Follicular phase - Ipsilateral - Native - Ampulla - Follicular phase - Contralateral - Nati
## Ampulla - Luteal phase - Contralateral - Decellularized - Ampulla - Follicular phase - Contralateral
## Ampulla - Luteal phase - Contralateral - Native - Ampulla - Follicular phase - Contralateral - Nativ
## Ampulla - Luteal phase - Ipsilateral - Decellularized - Ampulla - Follicular phase - Contralateral -
## Ampulla - Luteal phase - Ipsilateral - Native - Ampulla - Follicular phase - Contralateral - Native
## Isthmus - Follicular phase - Contralateral - Decellularized - Ampulla - Follicular phase - Contralat
## Isthmus - Follicular phase - Contralateral - Native - Ampulla - Follicular phase - Contralateral - Na
## Isthmus - Follicular phase - Ipsilateral - Decellularized - Ampulla - Follicular phase - Contralatera
## Isthmus - Follicular phase - Ipsilateral - Native - Ampulla - Follicular phase - Contralateral - Nati
## Isthmus - Luteal phase - Contralateral - Decellularized - Ampulla - Follicular phase - Contralateral
## Isthmus - Luteal phase - Contralateral - Native - Ampulla - Follicular phase - Contralateral - Nativ
## Isthmus - Luteal phase - Ipsilateral - Decellularized - Ampulla - Follicular phase - Contralateral -
## Isthmus - Luteal phase - Ipsilateral - Native - Ampulla - Follicular phase - Contralateral - Native
## Ampulla - Follicular phase - Ipsilateral - Native - Ampulla - Follicular phase - Ipsilateral - Decel
## Ampulla - Luteal phase - Contralateral - Decellularized - Ampulla - Follicular phase - Ipsilateral -
## Ampulla - Luteal phase - Contralateral - Native - Ampulla - Follicular phase - Ipsilateral - Decellu
## Ampulla - Luteal phase - Ipsilateral - Decellularized - Ampulla - Follicular phase - Ipsilateral - De
## Ampulla - Luteal phase - Ipsilateral - Native - Ampulla - Follicular phase - Ipsilateral - Decellula
## Isthmus - Follicular phase - Contralateral - Decellularized - Ampulla - Follicular phase - Ipsilatera
## Isthmus - Follicular phase - Contralateral - Native - Ampulla - Follicular phase - Ipsilateral - Dec
## Isthmus - Follicular phase - Ipsilateral - Decellularized - Ampulla - Follicular phase - Ipsilateral
## Isthmus - Follicular phase - Ipsilateral - Native - Ampulla - Follicular phase - Ipsilateral - Decel
## Isthmus - Luteal phase - Contralateral - Decellularized - Ampulla - Follicular phase - Ipsilateral -
## Isthmus - Luteal phase - Contralateral - Native - Ampulla - Follicular phase - Ipsilateral - Decellu
## Isthmus - Luteal phase - Ipsilateral - Decellularized - Ampulla - Follicular phase - Ipsilateral - De
## Isthmus - Luteal phase - Ipsilateral - Native - Ampulla - Follicular phase - Ipsilateral - Decellula
## Ampulla - Luteal phase - Contralateral - Decellularized - Ampulla - Follicular phase - Ipsilateral -
## Ampulla - Luteal phase - Contralateral - Native - Ampulla - Follicular phase - Ipsilateral - Native
## Ampulla - Luteal phase - Ipsilateral - Decellularized - Ampulla - Follicular phase - Ipsilateral - Na
## Ampulla - Luteal phase - Ipsilateral - Native - Ampulla - Follicular phase - Ipsilateral - Native ==
## Isthmus - Follicular phase - Contralateral - Decellularized - Ampulla - Follicular phase - Ipsilatera
## Isthmus - Follicular phase - Contralateral - Native - Ampulla - Follicular phase - Ipsilateral - Nati
## Isthmus - Follicular phase - Ipsilateral - Decellularized - Ampulla - Follicular phase - Ipsilateral
## Isthmus - Follicular phase - Ipsilateral - Native - Ampulla - Follicular phase - Ipsilateral - Nativ
## Isthmus - Luteal phase - Contralateral - Decellularized - Ampulla - Follicular phase - Ipsilateral -
## Isthmus - Luteal phase - Contralateral - Native - Ampulla - Follicular phase - Ipsilateral - Native
## Isthmus - Luteal phase - Ipsilateral - Decellularized - Ampulla - Follicular phase - Ipsilateral - Na
## Isthmus - Luteal phase - Ipsilateral - Native - Ampulla - Follicular phase - Ipsilateral - Native ==
## Ampulla - Luteal phase - Contralateral - Native - Ampulla - Luteal phase - Contralateral - Decellula
## Ampulla - Luteal phase - Ipsilateral - Decellularized - Ampulla - Luteal phase - Contralateral - Dec
## Ampulla - Luteal phase - Ipsilateral - Native - Ampulla - Luteal phase - Contralateral - Decellulari
## Isthmus - Follicular phase - Contralateral - Decellularized - Ampulla - Luteal phase - Contralateral
## Isthmus - Follicular phase - Contralateral - Native - Ampulla - Luteal phase - Contralateral - Decel
## Isthmus - Follicular phase - Ipsilateral - Decellularized - Ampulla - Luteal phase - Contralateral -
## Isthmus - Follicular phase - Ipsilateral - Native - Ampulla - Luteal phase - Contralateral - Decellu
## Isthmus - Luteal phase - Contralateral - Decellularized - Ampulla - Luteal phase - Contralateral - De
## Isthmus - Luteal phase - Contralateral - Native - Ampulla - Luteal phase - Contralateral - Decellula
## Isthmus - Luteal phase - Ipsilateral - Decellularized - Ampulla - Luteal phase - Contralateral - Dec
## Isthmus - Luteal phase - Ipsilateral - Native - Ampulla - Luteal phase - Contralateral - Decellulari
## Ampulla - Luteal phase - Ipsilateral - Decellularized - Ampulla - Luteal phase - Contralateral - Nati
## Ampulla - Luteal phase - Ipsilateral - Native - Ampulla - Luteal phase - Contralateral - Native == 0
## Isthmus - Follicular phase - Contralateral - Decellularized - Ampulla - Luteal phase - Contralateral
## Isthmus - Follicular phase - Contralateral - Native - Ampulla - Luteal phase - Contralateral - Nativ
```

```
## Isthmus - Follicular phase - Ipsilateral - Decellularized - Ampulla - Luteal phase - Contralateral -
## Isthmus - Follicular phase - Ipsilateral - Native - Ampulla - Luteal phase - Contralateral - Native =
## Isthmus - Luteal phase - Contralateral - Decellularized - Ampulla - Luteal phase - Contralateral - Na
## Isthmus - Luteal phase - Contralateral - Native - Ampulla - Luteal phase - Contralateral - Native ==
## Isthmus - Luteal phase - Ipsilateral - Decellularized - Ampulla - Luteal phase - Contralateral - Nati
## Isthmus - Luteal phase - Ipsilateral - Native - Ampulla - Luteal phase - Contralateral - Native == 0
## Ampulla - Luteal phase - Ipsilateral - Native - Ampulla - Luteal phase - Ipsilateral - Decellularized
## Isthmus - Follicular phase - Contralateral - Decellularized - Ampulla - Luteal phase - Ipsilateral -
## Isthmus - Follicular phase - Contralateral - Native - Ampulla - Luteal phase - Ipsilateral - Decellul
## Isthmus - Follicular phase - Ipsilateral - Decellularized - Ampulla - Luteal phase - Ipsilateral - De
## Isthmus - Follicular phase - Ipsilateral - Native - Ampulla - Luteal phase - Ipsilateral - Decellula
## Isthmus - Luteal phase - Contralateral - Decellularized - Ampulla - Luteal phase - Ipsilateral - Dece
## Isthmus - Luteal phase - Contralateral - Native - Ampulla - Luteal phase - Ipsilateral - Decellulari
## Isthmus - Luteal phase - Ipsilateral - Decellularized - Ampulla - Luteal phase - Ipsilateral - Decel
## Isthmus - Luteal phase - Ipsilateral - Native - Ampulla - Luteal phase - Ipsilateral - Decellularized
## Isthmus - Follicular phase - Contralateral - Decellularized - Ampulla - Luteal phase - Ipsilateral -
## Isthmus - Follicular phase - Contralateral - Native - Ampulla - Luteal phase - Ipsilateral - Native =
## Isthmus - Follicular phase - Ipsilateral - Decellularized - Ampulla - Luteal phase - Ipsilateral - Na
## Isthmus - Follicular phase - Ipsilateral - Native - Ampulla - Luteal phase - Ipsilateral - Native ==
## Isthmus - Luteal phase - Contralateral - Decellularized - Ampulla - Luteal phase - Ipsilateral - Nati
## Isthmus - Luteal phase - Contralateral - Native - Ampulla - Luteal phase - Ipsilateral - Native == 0
## Isthmus - Luteal phase - Ipsilateral - Decellularized - Ampulla - Luteal phase - Ipsilateral - Native
## Isthmus - Luteal phase - Ipsilateral - Native - Ampulla - Luteal phase - Ipsilateral - Native == 0
## Isthmus - Follicular phase - Contralateral - Native - Isthmus - Follicular phase - Contralateral - De
## Isthmus - Follicular phase - Ipsilateral - Decellularized - Isthmus - Follicular phase - Contralatera
## Isthmus - Follicular phase - Ipsilateral - Native - Isthmus - Follicular phase - Contralateral - Dece
## Isthmus - Luteal phase - Contralateral - Decellularized - Isthmus - Follicular phase - Contralateral
## Isthmus - Luteal phase - Contralateral - Native - Isthmus - Follicular phase - Contralateral - Decell
## Isthmus - Luteal phase - Ipsilateral - Decellularized - Isthmus - Follicular phase - Contralateral -
## Isthmus - Luteal phase - Ipsilateral - Native - Isthmus - Follicular phase - Contralateral - Decellul
## Isthmus - Follicular phase - Ipsilateral - Decellularized - Isthmus - Follicular phase - Contralatera
## Isthmus - Follicular phase - Ipsilateral - Native - Isthmus - Follicular phase - Contralateral - Nati
## Isthmus - Luteal phase - Contralateral - Decellularized - Isthmus - Follicular phase - Contralateral
## Isthmus - Luteal phase - Contralateral - Native - Isthmus - Follicular phase - Contralateral - Native
## Isthmus - Luteal phase - Ipsilateral - Decellularized - Isthmus - Follicular phase - Contralateral -
## Isthmus - Luteal phase - Ipsilateral - Native - Isthmus - Follicular phase - Contralateral - Native =
## Isthmus - Follicular phase - Ipsilateral - Native - Isthmus - Follicular phase - Ipsilateral - Decell
## Isthmus - Luteal phase - Contralateral - Decellularized - Isthmus - Follicular phase - Ipsilateral -
## Isthmus - Luteal phase - Contralateral - Native - Isthmus - Follicular phase - Ipsilateral - Decellul
## Isthmus - Luteal phase - Ipsilateral - Decellularized - Isthmus - Follicular phase - Ipsilateral - De
## Isthmus - Luteal phase - Ipsilateral - Native - Isthmus - Follicular phase - Ipsilateral - Decellula
## Isthmus - Luteal phase - Contralateral - Decellularized - Isthmus - Follicular phase - Ipsilateral -
## Isthmus - Luteal phase - Contralateral - Native - Isthmus - Follicular phase - Ipsilateral - Native =
## Isthmus - Luteal phase - Ipsilateral - Decellularized - Isthmus - Follicular phase - Ipsilateral - Na
## Isthmus - Luteal phase - Ipsilateral - Native - Isthmus - Follicular phase - Ipsilateral - Native ==
## Isthmus - Luteal phase - Contralateral - Native - Isthmus - Luteal phase - Contralateral - Decellula
## Isthmus - Luteal phase - Ipsilateral - Decellularized - Isthmus - Luteal phase - Contralateral - Dece
## Isthmus - Luteal phase - Ipsilateral - Native - Isthmus - Luteal phase - Contralateral - Decellulari
## Isthmus - Luteal phase - Ipsilateral - Decellularized - Isthmus - Luteal phase - Contralateral - Nati
## Isthmus - Luteal phase - Ipsilateral - Native - Isthmus - Luteal phase - Contralateral - Native == 0
## Isthmus - Luteal phase - Ipsilateral - Native - Isthmus - Luteal phase - Ipsilateral - Decellularized
##
## Ampulla - Follicular phase - Contralateral - Native - Ampulla - Follicular phase - Contralateral - De
## Ampulla - Follicular phase - Ipsilateral - Decellularized - Ampulla - Follicular phase - Contralatera
```

```
## Ampulla - Follicular phase - Ipsilateral - Native - Ampulla - Follicular phase - Contralateral - Dec
## Ampulla - Luteal phase - Contralateral - Decellularized - Ampulla - Follicular phase - Contralateral
## Ampulla - Luteal phase - Contralateral - Native - Ampulla - Follicular phase - Contralateral - Decell
## Ampulla - Luteal phase - Ipsilateral - Decellularized - Ampulla - Follicular phase - Contralateral -
## Ampulla - Luteal phase - Ipsilateral - Native - Ampulla - Follicular phase - Contralateral - Decellul
## Isthmus - Follicular phase - Contralateral - Decellularized - Ampulla - Follicular phase - Contralat
## Isthmus - Follicular phase - Contralateral - Native - Ampulla - Follicular phase - Contralateral - De
## Isthmus - Follicular phase - Ipsilateral - Decellularized - Ampulla - Follicular phase - Contralatera
## Isthmus - Follicular phase - Ipsilateral - Native - Ampulla - Follicular phase - Contralateral - Dec
## Isthmus - Luteal phase - Contralateral - Decellularized - Ampulla - Follicular phase - Contralateral
## Isthmus - Luteal phase - Contralateral - Native - Ampulla - Follicular phase - Contralateral - Decell
## Isthmus - Luteal phase - Ipsilateral - Decellularized - Ampulla - Follicular phase - Contralateral -
## Isthmus - Luteal phase - Ipsilateral - Native - Ampulla - Follicular phase - Contralateral - Decellul
## Ampulla - Follicular phase - Ipsilateral - Decellularized - Ampulla - Follicular phase - Contralatera
## Ampulla - Follicular phase - Ipsilateral - Native - Ampulla - Follicular phase - Contralateral - Nati
## Ampulla - Luteal phase - Contralateral - Decellularized - Ampulla - Follicular phase - Contralateral
## Ampulla - Luteal phase - Contralateral - Native - Ampulla - Follicular phase - Contralateral - Native
## Ampulla - Luteal phase - Ipsilateral - Decellularized - Ampulla - Follicular phase - Contralateral -
## Ampulla - Luteal phase - Ipsilateral - Native - Ampulla - Follicular phase - Contralateral - Native =
## Isthmus - Follicular phase - Contralateral - Decellularized - Ampulla - Follicular phase - Contralat
## Isthmus - Follicular phase - Contralateral - Native - Ampulla - Follicular phase - Contralateral - Na
## Isthmus - Follicular phase - Ipsilateral - Decellularized - Ampulla - Follicular phase - Contralatera
## Isthmus - Follicular phase - Ipsilateral - Native - Ampulla - Follicular phase - Contralateral - Nati
## Isthmus - Luteal phase - Contralateral - Decellularized - Ampulla - Follicular phase - Contralateral
## Isthmus - Luteal phase - Contralateral - Native - Ampulla - Follicular phase - Contralateral - Native
## Isthmus - Luteal phase - Ipsilateral - Decellularized - Ampulla - Follicular phase - Contralateral -
## Isthmus - Luteal phase - Ipsilateral - Native - Ampulla - Follicular phase - Contralateral - Native =
## Ampulla - Follicular phase - Ipsilateral - Native - Ampulla - Follicular phase - Ipsilateral - Decell
## Ampulla - Luteal phase - Contralateral - Decellularized - Ampulla - Follicular phase - Ipsilateral -
## Ampulla - Luteal phase - Contralateral - Native - Ampulla - Follicular phase - Ipsilateral - Decellul
## Ampulla - Luteal phase - Ipsilateral - Decellularized - Ampulla - Follicular phase - Ipsilateral - De
## Ampulla - Luteal phase - Ipsilateral - Native - Ampulla - Follicular phase - Ipsilateral - Decellula
## Isthmus - Follicular phase - Contralateral - Decellularized - Ampulla - Follicular phase - Ipsilatera
## Isthmus - Follicular phase - Contralateral - Native - Ampulla - Follicular phase - Ipsilateral - Dec
## Isthmus - Follicular phase - Ipsilateral - Decellularized - Ampulla - Follicular phase - Ipsilateral
## Isthmus - Follicular phase - Ipsilateral - Native - Ampulla - Follicular phase - Ipsilateral - Decell
## Isthmus - Luteal phase - Contralateral - Decellularized - Ampulla - Follicular phase - Ipsilateral -
## Isthmus - Luteal phase - Contralateral - Native - Ampulla - Follicular phase - Ipsilateral - Decellul
## Isthmus - Luteal phase - Ipsilateral - Decellularized - Ampulla - Follicular phase - Ipsilateral - De
## Isthmus - Luteal phase - Ipsilateral - Native - Ampulla - Follicular phase - Ipsilateral - Decellula
## Ampulla - Luteal phase - Contralateral - Decellularized - Ampulla - Follicular phase - Ipsilateral -
## Ampulla - Luteal phase - Contralateral - Native - Ampulla - Follicular phase - Ipsilateral - Native =
## Ampulla - Luteal phase - Ipsilateral - Decellularized - Ampulla - Follicular phase - Ipsilateral - Na
## Ampulla - Luteal phase - Ipsilateral - Native - Ampulla - Follicular phase - Ipsilateral - Native ==
## Isthmus - Follicular phase - Contralateral - Decellularized - Ampulla - Follicular phase - Ipsilatera
## Isthmus - Follicular phase - Contralateral - Native - Ampulla - Follicular phase - Ipsilateral - Nati
## Isthmus - Follicular phase - Ipsilateral - Decellularized - Ampulla - Follicular phase - Ipsilateral
## Isthmus - Follicular phase - Ipsilateral - Native - Ampulla - Follicular phase - Ipsilateral - Native
## Isthmus - Luteal phase - Contralateral - Decellularized - Ampulla - Follicular phase - Ipsilateral -
## Isthmus - Luteal phase - Contralateral - Native - Ampulla - Follicular phase - Ipsilateral - Native =
## Isthmus - Luteal phase - Ipsilateral - Decellularized - Ampulla - Follicular phase - Ipsilateral - Na
## Isthmus - Luteal phase - Ipsilateral - Native - Ampulla - Follicular phase - Ipsilateral - Native ==
## Ampulla - Luteal phase - Contralateral - Native - Ampulla - Luteal phase - Contralateral - Decellula
## Ampulla - Luteal phase - Ipsilateral - Decellularized - Ampulla - Luteal phase - Contralateral - Dec
```

```
## Ampulla - Luteal phase - Ipsilateral - Native - Ampulla - Luteal phase - Contralateral - Decellulari
## Isthmus - Follicular phase - Contralateral - Decellularized - Ampulla - Luteal phase - Contralateral
## Isthmus - Follicular phase - Contralateral - Native - Ampulla - Luteal phase - Contralateral - Decel
## Isthmus - Follicular phase - Ipsilateral - Decellularized - Ampulla - Luteal phase - Contralateral -
## Isthmus - Follicular phase - Ipsilateral - Native - Ampulla - Luteal phase - Contralateral - Decellu
## Isthmus - Luteal phase - Contralateral - Decellularized - Ampulla - Luteal phase - Contralateral - De
## Isthmus - Luteal phase - Contralateral - Native - Ampulla - Luteal phase - Contralateral - Decellula
## Isthmus - Luteal phase - Ipsilateral - Decellularized - Ampulla - Luteal phase - Contralateral - Dec
## Isthmus - Luteal phase - Ipsilateral - Native - Ampulla - Luteal phase - Contralateral - Decellulari
## Ampulla - Luteal phase - Ipsilateral - Decellularized - Ampulla - Luteal phase - Contralateral - Nati
## Ampulla - Luteal phase - Ipsilateral - Native - Ampulla - Luteal phase - Contralateral - Native == 0
## Isthmus - Follicular phase - Contralateral - Decellularized - Ampulla - Luteal phase - Contralateral
## Isthmus - Follicular phase - Contralateral - Native - Ampulla - Luteal phase - Contralateral - Native
## Isthmus - Follicular phase - Ipsilateral - Decellularized - Ampulla - Luteal phase - Contralateral -
## Isthmus - Follicular phase - Ipsilateral - Native - Ampulla - Luteal phase - Contralateral - Native =
## Isthmus - Luteal phase - Contralateral - Decellularized - Ampulla - Luteal phase - Contralateral - Na
## Isthmus - Luteal phase - Contralateral - Native - Ampulla - Luteal phase - Contralateral - Native ==
## Isthmus - Luteal phase - Ipsilateral - Decellularized - Ampulla - Luteal phase - Contralateral - Nati
## Isthmus - Luteal phase - Ipsilateral - Native - Ampulla - Luteal phase - Contralateral - Native == 0
## Ampulla - Luteal phase - Ipsilateral - Native - Ampulla - Luteal phase - Ipsilateral - Decellularized
## Isthmus - Follicular phase - Contralateral - Decellularized - Ampulla - Luteal phase - Ipsilateral -
## Isthmus - Follicular phase - Contralateral - Native - Ampulla - Luteal phase - Ipsilateral - Decellu
## Isthmus - Follicular phase - Ipsilateral - Decellularized - Ampulla - Luteal phase - Ipsilateral - De
## Isthmus - Follicular phase - Ipsilateral - Native - Ampulla - Luteal phase - Ipsilateral - Decellula
## Isthmus - Luteal phase - Contralateral - Decellularized - Ampulla - Luteal phase - Ipsilateral - Dec
## Isthmus - Luteal phase - Contralateral - Native - Ampulla - Luteal phase - Ipsilateral - Decellulari
## Isthmus - Luteal phase - Ipsilateral - Decellularized - Ampulla - Luteal phase - Ipsilateral - Decel
## Isthmus - Luteal phase - Ipsilateral - Native - Ampulla - Luteal phase - Ipsilateral - Decellularized
## Isthmus - Follicular phase - Contralateral - Decellularized - Ampulla - Luteal phase - Ipsilateral -
## Isthmus - Follicular phase - Contralateral - Native - Ampulla - Luteal phase - Ipsilateral - Native =
## Isthmus - Follicular phase - Ipsilateral - Decellularized - Ampulla - Luteal phase - Ipsilateral - Na
## Isthmus - Follicular phase - Ipsilateral - Native - Ampulla - Luteal phase - Ipsilateral - Native ==
## Isthmus - Luteal phase - Contralateral - Decellularized - Ampulla - Luteal phase - Ipsilateral - Nati
## Isthmus - Luteal phase - Contralateral - Native - Ampulla - Luteal phase - Ipsilateral - Native == 0
## Isthmus - Luteal phase - Ipsilateral - Decellularized - Ampulla - Luteal phase - Ipsilateral - Native
## Isthmus - Luteal phase - Ipsilateral - Native - Ampulla - Luteal phase - Ipsilateral - Native == 0
## Isthmus - Follicular phase - Contralateral - Native - Isthmus - Follicular phase - Contralateral - De
## Isthmus - Follicular phase - Ipsilateral - Decellularized - Isthmus - Follicular phase - Contralatera
## Isthmus - Follicular phase - Ipsilateral - Native - Isthmus - Follicular phase - Contralateral - Dec
## Isthmus - Luteal phase - Contralateral - Decellularized - Isthmus - Follicular phase - Contralateral
## Isthmus - Luteal phase - Contralateral - Native - Isthmus - Follicular phase - Contralateral - Decel
## Isthmus - Luteal phase - Ipsilateral - Decellularized - Isthmus - Follicular phase - Contralateral -
## Isthmus - Luteal phase - Ipsilateral - Native - Isthmus - Follicular phase - Contralateral - Decellu
## Isthmus - Follicular phase - Ipsilateral - Decellularized - Isthmus - Follicular phase - Contralatera
## Isthmus - Follicular phase - Ipsilateral - Native - Isthmus - Follicular phase - Contralateral - Nati
## Isthmus - Luteal phase - Contralateral - Decellularized - Isthmus - Follicular phase - Contralateral
## Isthmus - Luteal phase - Contralateral - Native - Isthmus - Follicular phase - Contralateral - Native
## Isthmus - Luteal phase - Ipsilateral - Decellularized - Isthmus - Follicular phase - Contralateral -
## Isthmus - Luteal phase - Ipsilateral - Native - Isthmus - Follicular phase - Contralateral - Native =
## Isthmus - Follicular phase - Ipsilateral - Native - Isthmus - Follicular phase - Ipsilateral - Decel
## Isthmus - Luteal phase - Contralateral - Decellularized - Isthmus - Follicular phase - Ipsilateral -
## Isthmus - Luteal phase - Contralateral - Native - Isthmus - Follicular phase - Ipsilateral - Decellu
## Isthmus - Luteal phase - Ipsilateral - Decellularized - Isthmus - Follicular phase - Ipsilateral - De
## Isthmus - Luteal phase - Ipsilateral - Native - Isthmus - Follicular phase - Ipsilateral - Decellula
```

```
## Isthmus - Luteal phase - Contralateral - Decellularized - Isthmus - Follicular phase - Ipsilateral -
## Isthmus - Luteal phase - Contralateral - Native - Isthmus - Follicular phase - Ipsilateral - Native =
## Isthmus - Luteal phase - Ipsilateral - Decellularized - Isthmus - Follicular phase - Ipsilateral - Na
## Isthmus - Luteal phase - Ipsilateral - Native - Isthmus - Follicular phase - Ipsilateral - Native ==
## Isthmus - Luteal phase - Contralateral - Native - Isthmus - Luteal phase - Contralateral - Decellula
## Isthmus - Luteal phase - Ipsilateral - Decellularized - Isthmus - Luteal phase - Contralateral - Dec
## Isthmus - Luteal phase - Ipsilateral - Native - Isthmus - Luteal phase - Contralateral - Decellulari
## Isthmus - Luteal phase - Ipsilateral - Decellularized - Isthmus - Luteal phase - Contralateral - Nati
## Isthmus - Luteal phase - Ipsilateral - Native - Isthmus - Luteal phase - Contralateral - Native == 0
## Isthmus - Luteal phase - Ipsilateral - Native - Isthmus - Luteal phase - Ipsilateral - Decellularized
##
## Ampulla - Follicular phase - Contralateral - Native - Ampulla - Follicular phase - Contralateral - De
## Ampulla - Follicular phase - Ipsilateral - Decellularized - Ampulla - Follicular phase - Contralatera
## Ampulla - Follicular phase - Ipsilateral - Native - Ampulla - Follicular phase - Contralateral - Dec
## Ampulla - Luteal phase - Contralateral - Decellularized - Ampulla - Follicular phase - Contralateral
## Ampulla - Luteal phase - Contralateral - Native - Ampulla - Follicular phase - Contralateral - Decel
## Ampulla - Luteal phase - Ipsilateral - Decellularized - Ampulla - Follicular phase - Contralateral -
## Ampulla - Luteal phase - Ipsilateral - Native - Ampulla - Follicular phase - Contralateral - Decellu
## Isthmus - Follicular phase - Contralateral - Decellularized - Ampulla - Follicular phase - Contralat
## Isthmus - Follicular phase - Contralateral - Native - Ampulla - Follicular phase - Contralateral - De
## Isthmus - Follicular phase - Ipsilateral - Decellularized - Ampulla - Follicular phase - Contralatera
## Isthmus - Follicular phase - Ipsilateral - Native - Ampulla - Follicular phase - Contralateral - Dec
## Isthmus - Luteal phase - Contralateral - Decellularized - Ampulla - Follicular phase - Contralateral
## Isthmus - Luteal phase - Contralateral - Native - Ampulla - Follicular phase - Contralateral - Decel
## Isthmus - Luteal phase - Ipsilateral - Decellularized - Ampulla - Follicular phase - Contralateral -
## Isthmus - Luteal phase - Ipsilateral - Native - Ampulla - Follicular phase - Contralateral - Decellu
## Ampulla - Follicular phase - Ipsilateral - Decellularized - Ampulla - Follicular phase - Contralatera
## Ampulla - Follicular phase - Ipsilateral - Native - Ampulla - Follicular phase - Contralateral - Nati
## Ampulla - Luteal phase - Contralateral - Decellularized - Ampulla - Follicular phase - Contralateral
## Ampulla - Luteal phase - Contralateral - Native - Ampulla - Follicular phase - Contralateral - Native
## Ampulla - Luteal phase - Ipsilateral - Decellularized - Ampulla - Follicular phase - Contralateral -
## Ampulla - Luteal phase - Ipsilateral - Native - Ampulla - Follicular phase - Contralateral - Native =
## Isthmus - Follicular phase - Contralateral - Decellularized - Ampulla - Follicular phase - Contralat
## Isthmus - Follicular phase - Contralateral - Native - Ampulla - Follicular phase - Contralateral - Na
## Isthmus - Follicular phase - Ipsilateral - Decellularized - Ampulla - Follicular phase - Contralatera
## Isthmus - Follicular phase - Ipsilateral - Native - Ampulla - Follicular phase - Contralateral - Nati
## Isthmus - Luteal phase - Contralateral - Decellularized - Ampulla - Follicular phase - Contralateral
## Isthmus - Luteal phase - Contralateral - Native - Ampulla - Follicular phase - Contralateral - Native
## Isthmus - Luteal phase - Ipsilateral - Decellularized - Ampulla - Follicular phase - Contralateral -
## Isthmus - Luteal phase - Ipsilateral - Native - Ampulla - Follicular phase - Contralateral - Native =
## Ampulla - Follicular phase - Ipsilateral - Native - Ampulla - Follicular phase - Ipsilateral - Decell
## Ampulla - Luteal phase - Contralateral - Decellularized - Ampulla - Follicular phase - Ipsilateral -
## Ampulla - Luteal phase - Contralateral - Native - Ampulla - Follicular phase - Ipsilateral - Decellul
## Ampulla - Luteal phase - Ipsilateral - Decellularized - Ampulla - Follicular phase - Ipsilateral - De
## Ampulla - Luteal phase - Ipsilateral - Native - Ampulla - Follicular phase - Ipsilateral - Decellula
## Isthmus - Follicular phase - Contralateral - Decellularized - Ampulla - Follicular phase - Ipsilatera
## Isthmus - Follicular phase - Contralateral - Native - Ampulla - Follicular phase - Ipsilateral - Dec
## Isthmus - Follicular phase - Ipsilateral - Decellularized - Ampulla - Follicular phase - Ipsilateral
## Isthmus - Follicular phase - Ipsilateral - Native - Ampulla - Follicular phase - Ipsilateral - Decell
## Isthmus - Luteal phase - Contralateral - Decellularized - Ampulla - Follicular phase - Ipsilateral -
## Isthmus - Luteal phase - Contralateral - Native - Ampulla - Follicular phase - Ipsilateral - Decellul
## Isthmus - Luteal phase - Ipsilateral - Decellularized - Ampulla - Follicular phase - Ipsilateral - De
## Isthmus - Luteal phase - Ipsilateral - Native - Ampulla - Follicular phase - Ipsilateral - Decellula
## Ampulla - Luteal phase - Contralateral - Decellularized - Ampulla - Follicular phase - Ipsilateral -
```

```
## Ampulla - Luteal phase - Contralateral - Native - Ampulla - Follicular phase - Ipsilateral - Native =
## Ampulla - Luteal phase - Ipsilateral - Decellularized - Ampulla - Follicular phase - Ipsilateral - Na
## Ampulla - Luteal phase - Ipsilateral - Native - Ampulla - Follicular phase - Ipsilateral - Native ==
## Isthmus - Follicular phase - Contralateral - Decellularized - Ampulla - Follicular phase - Ipsilatera
## Isthmus - Follicular phase - Contralateral - Native - Ampulla - Follicular phase - Ipsilateral - Nati
## Isthmus - Follicular phase - Ipsilateral - Decellularized - Ampulla - Follicular phase - Ipsilateral
## Isthmus - Follicular phase - Ipsilateral - Native - Ampulla - Follicular phase - Ipsilateral - Native
## Isthmus - Luteal phase - Contralateral - Decellularized - Ampulla - Follicular phase - Ipsilateral -
## Isthmus - Luteal phase - Contralateral - Native - Ampulla - Follicular phase - Ipsilateral - Native =
## Isthmus - Luteal phase - Ipsilateral - Decellularized - Ampulla - Follicular phase - Ipsilateral - Na
## Isthmus - Luteal phase - Ipsilateral - Native - Ampulla - Follicular phase - Ipsilateral - Native ==
## Ampulla - Luteal phase - Contralateral - Native - Ampulla - Luteal phase - Contralateral - Decellular
## Ampulla - Luteal phase - Ipsilateral - Decellularized - Ampulla - Luteal phase - Contralateral - Dece
## Ampulla - Luteal phase - Ipsilateral - Native - Ampulla - Luteal phase - Contralateral - Decellulari
## Isthmus - Follicular phase - Contralateral - Decellularized - Ampulla - Luteal phase - Contralateral
## Isthmus - Follicular phase - Contralateral - Native - Ampulla - Luteal phase - Contralateral - Decell
## Isthmus - Follicular phase - Ipsilateral - Decellularized - Ampulla - Luteal phase - Contralateral -
## Isthmus - Follicular phase - Ipsilateral - Native - Ampulla - Luteal phase - Contralateral - Decellul
## Isthmus - Luteal phase - Contralateral - Decellularized - Ampulla - Luteal phase - Contralateral - De
## Isthmus - Luteal phase - Contralateral - Native - Ampulla - Luteal phase - Contralateral - Decellular
## Isthmus - Luteal phase - Ipsilateral - Decellularized - Ampulla - Luteal phase - Contralateral - Dece
## Isthmus - Luteal phase - Ipsilateral - Native - Ampulla - Luteal phase - Contralateral - Decellulari
## Ampulla - Luteal phase - Ipsilateral - Decellularized - Ampulla - Luteal phase - Contralateral - Nati
## Ampulla - Luteal phase - Ipsilateral - Native - Ampulla - Luteal phase - Contralateral - Native == 0
## Isthmus - Follicular phase - Contralateral - Decellularized - Ampulla - Luteal phase - Contralateral
## Isthmus - Follicular phase - Contralateral - Native - Ampulla - Luteal phase - Contralateral - Native
## Isthmus - Follicular phase - Ipsilateral - Decellularized - Ampulla - Luteal phase - Contralateral -
## Isthmus - Follicular phase - Ipsilateral - Native - Ampulla - Luteal phase - Contralateral - Native =
## Isthmus - Luteal phase - Contralateral - Decellularized - Ampulla - Luteal phase - Contralateral - Na
## Isthmus - Luteal phase - Contralateral - Native - Ampulla - Luteal phase - Contralateral - Native ==
## Isthmus - Luteal phase - Ipsilateral - Decellularized - Ampulla - Luteal phase - Contralateral - Nati
## Isthmus - Luteal phase - Ipsilateral - Native - Ampulla - Luteal phase - Contralateral - Native == 0
## Ampulla - Luteal phase - Ipsilateral - Native - Ampulla - Luteal phase - Ipsilateral - Decellularized
## Isthmus - Follicular phase - Contralateral - Decellularized - Ampulla - Luteal phase - Ipsilateral -
## Isthmus - Follicular phase - Contralateral - Native - Ampulla - Luteal phase - Ipsilateral - Decellul
## Isthmus - Follicular phase - Ipsilateral - Decellularized - Ampulla - Luteal phase - Ipsilateral - De
## Isthmus - Follicular phase - Ipsilateral - Native - Ampulla - Luteal phase - Ipsilateral - Decellular
## Isthmus - Luteal phase - Contralateral - Decellularized - Ampulla - Luteal phase - Ipsilateral - Dece
## Isthmus - Luteal phase - Contralateral - Native - Ampulla - Luteal phase - Ipsilateral - Decellulari
## Isthmus - Luteal phase - Ipsilateral - Decellularized - Ampulla - Luteal phase - Ipsilateral - Decell
## Isthmus - Luteal phase - Ipsilateral - Native - Ampulla - Luteal phase - Ipsilateral - Decellularized
## Isthmus - Follicular phase - Contralateral - Decellularized - Ampulla - Luteal phase - Ipsilateral -
## Isthmus - Follicular phase - Contralateral - Native - Ampulla - Luteal phase - Ipsilateral - Native =
## Isthmus - Follicular phase - Ipsilateral - Decellularized - Ampulla - Luteal phase - Ipsilateral - Na
## Isthmus - Follicular phase - Ipsilateral - Native - Ampulla - Luteal phase - Ipsilateral - Native ==
## Isthmus - Luteal phase - Contralateral - Decellularized - Ampulla - Luteal phase - Ipsilateral - Nati
## Isthmus - Luteal phase - Contralateral - Native - Ampulla - Luteal phase - Ipsilateral - Native == 0
## Isthmus - Luteal phase - Ipsilateral - Decellularized - Ampulla - Luteal phase - Ipsilateral - Native
## Isthmus - Luteal phase - Ipsilateral - Native - Ampulla - Luteal phase - Ipsilateral - Native == 0
## Isthmus - Follicular phase - Contralateral - Native - Isthmus - Follicular phase - Contralateral - De
## Isthmus - Follicular phase - Ipsilateral - Decellularized - Isthmus - Follicular phase - Contralatera
## Isthmus - Follicular phase - Ipsilateral - Native - Isthmus - Follicular phase - Contralateral - Deco
## Isthmus - Luteal phase - Contralateral - Decellularized - Isthmus - Follicular phase - Contralateral
## Isthmus - Luteal phase - Contralateral - Native - Isthmus - Follicular phase - Contralateral - Decell
```

```
## Isthmus - Luteal phase - Ipsilateral - Decellularized - Isthmus - Follicular phase - Contralateral -
## Isthmus - Luteal phase - Ipsilateral - Native - Isthmus - Follicular phase - Contralateral - Decellul
## Isthmus - Follicular phase - Ipsilateral - Decellularized - Isthmus - Follicular phase - Contralatera
## Isthmus - Follicular phase - Ipsilateral - Native - Isthmus - Follicular phase - Contralateral - Nati
## Isthmus - Luteal phase - Contralateral - Decellularized - Isthmus - Follicular phase - Contralateral
## Isthmus - Luteal phase - Contralateral - Native - Isthmus - Follicular phase - Contralateral - Native
## Isthmus - Luteal phase - Ipsilateral - Decellularized - Isthmus - Follicular phase - Contralateral -
## Isthmus - Luteal phase - Ipsilateral - Native - Isthmus - Follicular phase - Contralateral - Native =
## Isthmus - Follicular phase - Ipsilateral - Native - Isthmus - Follicular phase - Ipsilateral - Decell
## Isthmus - Luteal phase - Contralateral - Decellularized - Isthmus - Follicular phase - Ipsilateral -
## Isthmus - Luteal phase - Contralateral - Native - Isthmus - Follicular phase - Ipsilateral - Decellul
## Isthmus - Luteal phase - Ipsilateral - Decellularized - Isthmus - Follicular phase - Ipsilateral - De
## Isthmus - Luteal phase - Ipsilateral - Native - Isthmus - Follicular phase - Ipsilateral - Decellula
## Isthmus - Luteal phase - Contralateral - Decellularized - Isthmus - Follicular phase - Ipsilateral -
## Isthmus - Luteal phase - Contralateral - Native - Isthmus - Follicular phase - Ipsilateral - Native =
## Isthmus - Luteal phase - Ipsilateral - Decellularized - Isthmus - Follicular phase - Ipsilateral - Na
## Isthmus - Luteal phase - Ipsilateral - Native - Isthmus - Follicular phase - Ipsilateral - Native ==
## Isthmus - Luteal phase - Contralateral - Native - Isthmus - Luteal phase - Contralateral - Decellula
## Isthmus - Luteal phase - Ipsilateral - Decellularized - Isthmus - Luteal phase - Contralateral - Dec
## Isthmus - Luteal phase - Ipsilateral - Native - Isthmus - Luteal phase - Contralateral - Decellulari
## Isthmus - Luteal phase - Ipsilateral - Decellularized - Isthmus - Luteal phase - Contralateral - Nati
## Isthmus - Luteal phase - Ipsilateral - Native - Isthmus - Luteal phase - Contralateral - Native == 0
## Isthmus - Luteal phase - Ipsilateral - Native - Isthmus - Luteal phase - Ipsilateral - Decellularized
##
## Ampulla - Follicular phase - Contralateral - Native - Ampulla - Follicular phase - Contralateral - De
## Ampulla - Follicular phase - Ipsilateral - Decellularized - Ampulla - Follicular phase - Contralatera
## Ampulla - Follicular phase - Ipsilateral - Native - Ampulla - Follicular phase - Contralateral - Dec
## Ampulla - Luteal phase - Contralateral - Decellularized - Ampulla - Follicular phase - Contralateral
## Ampulla - Luteal phase - Contralateral - Native - Ampulla - Follicular phase - Contralateral - Decell
## Ampulla - Luteal phase - Ipsilateral - Decellularized - Ampulla - Follicular phase - Contralateral -
## Ampulla - Luteal phase - Ipsilateral - Native - Ampulla - Follicular phase - Contralateral - Decellul
## Isthmus - Follicular phase - Contralateral - Decellularized - Ampulla - Follicular phase - Contralate
## Isthmus - Follicular phase - Contralateral - Native - Ampulla - Follicular phase - Contralateral - De
## Isthmus - Follicular phase - Ipsilateral - Decellularized - Ampulla - Follicular phase - Contralatera
## Isthmus - Follicular phase - Ipsilateral - Native - Ampulla - Follicular phase - Contralateral - Dec
## Isthmus - Luteal phase - Contralateral - Decellularized - Ampulla - Follicular phase - Contralateral
## Isthmus - Luteal phase - Contralateral - Native - Ampulla - Follicular phase - Contralateral - Decell
## Isthmus - Luteal phase - Ipsilateral - Decellularized - Ampulla - Follicular phase - Contralateral -
## Isthmus - Luteal phase - Ipsilateral - Native - Ampulla - Follicular phase - Contralateral - Decellul
## Ampulla - Follicular phase - Ipsilateral - Decellularized - Ampulla - Follicular phase - Contralatera
## Ampulla - Follicular phase - Ipsilateral - Native - Ampulla - Follicular phase - Contralateral - Nati
## Ampulla - Luteal phase - Contralateral - Decellularized - Ampulla - Follicular phase - Contralateral
## Ampulla - Luteal phase - Contralateral - Native - Ampulla - Follicular phase - Contralateral - Native
## Ampulla - Luteal phase - Ipsilateral - Decellularized - Ampulla - Follicular phase - Contralateral -
## Ampulla - Luteal phase - Ipsilateral - Native - Ampulla - Follicular phase - Contralateral - Native =
## Isthmus - Follicular phase - Contralateral - Decellularized - Ampulla - Follicular phase - Contralate
## Isthmus - Follicular phase - Contralateral - Native - Ampulla - Follicular phase - Contralateral - Na
## Isthmus - Follicular phase - Ipsilateral - Decellularized - Ampulla - Follicular phase - Contralatera
## Isthmus - Follicular phase - Ipsilateral - Native - Ampulla - Follicular phase - Contralateral - Nati
## Isthmus - Luteal phase - Contralateral - Decellularized - Ampulla - Follicular phase - Contralateral
## Isthmus - Luteal phase - Contralateral - Native - Ampulla - Follicular phase - Contralateral - Native
## Isthmus - Luteal phase - Ipsilateral - Decellularized - Ampulla - Follicular phase - Contralateral -
## Isthmus - Luteal phase - Ipsilateral - Native - Ampulla - Follicular phase - Contralateral - Native =
## Ampulla - Follicular phase - Ipsilateral - Native - Ampulla - Follicular phase - Ipsilateral - Decell
```

```
## Ampulla - Luteal phase - Contralateral - Decellularized - Ampulla - Follicular phase - Ipsilateral -
## Ampulla - Luteal phase - Contralateral - Native - Ampulla - Follicular phase - Ipsilateral - Decellul
## Ampulla - Luteal phase - Ipsilateral - Decellularized - Ampulla - Follicular phase - Ipsilateral - De
## Ampulla - Luteal phase - Ipsilateral - Native - Ampulla - Follicular phase - Ipsilateral - Decellular
## Isthmus - Follicular phase - Contralateral - Decellularized - Ampulla - Follicular phase - Ipsilatera
## Isthmus - Follicular phase - Contralateral - Native - Ampulla - Follicular phase - Ipsilateral - Dece
## Isthmus - Follicular phase - Ipsilateral - Decellularized - Ampulla - Follicular phase - Ipsilateral
## Isthmus - Follicular phase - Ipsilateral - Native - Ampulla - Follicular phase - Ipsilateral - Decell
## Isthmus - Luteal phase - Contralateral - Decellularized - Ampulla - Follicular phase - Ipsilateral -
## Isthmus - Luteal phase - Contralateral - Native - Ampulla - Follicular phase - Ipsilateral - Decellul
## Isthmus - Luteal phase - Ipsilateral - Decellularized - Ampulla - Follicular phase - Ipsilateral - De
## Isthmus - Luteal phase - Ipsilateral - Native - Ampulla - Follicular phase - Ipsilateral - Decellular
## Ampulla - Luteal phase - Contralateral - Decellularized - Ampulla - Follicular phase - Ipsilateral -
## Ampulla - Luteal phase - Contralateral - Native - Ampulla - Follicular phase - Ipsilateral - Native =
## Ampulla - Luteal phase - Ipsilateral - Decellularized - Ampulla - Follicular phase - Ipsilateral - Na
## Ampulla - Luteal phase - Ipsilateral - Native - Ampulla - Follicular phase - Ipsilateral - Native ==
## Isthmus - Follicular phase - Contralateral - Decellularized - Ampulla - Follicular phase - Ipsilatera
## Isthmus - Follicular phase - Contralateral - Native - Ampulla - Follicular phase - Ipsilateral - Nati
## Isthmus - Follicular phase - Ipsilateral - Decellularized - Ampulla - Follicular phase - Ipsilateral
## Isthmus - Follicular phase - Ipsilateral - Native - Ampulla - Follicular phase - Ipsilateral - Native
## Isthmus - Luteal phase - Contralateral - Decellularized - Ampulla - Follicular phase - Ipsilateral -
## Isthmus - Luteal phase - Contralateral - Native - Ampulla - Follicular phase - Ipsilateral - Native =
## Isthmus - Luteal phase - Ipsilateral - Decellularized - Ampulla - Follicular phase - Ipsilateral - Na
## Isthmus - Luteal phase - Ipsilateral - Native - Ampulla - Follicular phase - Ipsilateral - Native ==
## Ampulla - Luteal phase - Contralateral - Native - Ampulla - Luteal phase - Contralateral - Decellular
## Ampulla - Luteal phase - Ipsilateral - Decellularized - Ampulla - Luteal phase - Contralateral - Dece
## Ampulla - Luteal phase - Ipsilateral - Native - Ampulla - Luteal phase - Contralateral - Decellulariz
## Isthmus - Follicular phase - Contralateral - Decellularized - Ampulla - Luteal phase - Contralateral
## Isthmus - Follicular phase - Contralateral - Native - Ampulla - Luteal phase - Contralateral - Decell
## Isthmus - Follicular phase - Ipsilateral - Decellularized - Ampulla - Luteal phase - Contralateral -
## Isthmus - Follicular phase - Ipsilateral - Native - Ampulla - Luteal phase - Contralateral - Decellul
## Isthmus - Luteal phase - Contralateral - Decellularized - Ampulla - Luteal phase - Contralateral - De
## Isthmus - Luteal phase - Contralateral - Native - Ampulla - Luteal phase - Contralateral - Decellular
## Isthmus - Luteal phase - Ipsilateral - Decellularized - Ampulla - Luteal phase - Contralateral - Dece
## Isthmus - Luteal phase - Ipsilateral - Native - Ampulla - Luteal phase - Contralateral - Decellulariz
## Ampulla - Luteal phase - Ipsilateral - Decellularized - Ampulla - Luteal phase - Contralateral - Nati
## Ampulla - Luteal phase - Ipsilateral - Native - Ampulla - Luteal phase - Contralateral - Native == 0
## Isthmus - Follicular phase - Contralateral - Decellularized - Ampulla - Luteal phase - Contralateral
## Isthmus - Follicular phase - Contralateral - Native - Ampulla - Luteal phase - Contralateral - Native
## Isthmus - Follicular phase - Ipsilateral - Decellularized - Ampulla - Luteal phase - Contralateral -
## Isthmus - Follicular phase - Ipsilateral - Native - Ampulla - Luteal phase - Contralateral - Native =
## Isthmus - Luteal phase - Contralateral - Decellularized - Ampulla - Luteal phase - Contralateral - Na
## Isthmus - Luteal phase - Contralateral - Native - Ampulla - Luteal phase - Contralateral - Native ==
## Isthmus - Luteal phase - Ipsilateral - Decellularized - Ampulla - Luteal phase - Contralateral - Nati
## Isthmus - Luteal phase - Ipsilateral - Native - Ampulla - Luteal phase - Contralateral - Native == 0
## Ampulla - Luteal phase - Ipsilateral - Native - Ampulla - Luteal phase - Ipsilateral - Decellularized
## Isthmus - Follicular phase - Contralateral - Decellularized - Ampulla - Luteal phase - Ipsilateral -
## Isthmus - Follicular phase - Contralateral - Native - Ampulla - Luteal phase - Ipsilateral - Decellul
## Isthmus - Follicular phase - Ipsilateral - Decellularized - Ampulla - Luteal phase - Ipsilateral - De
## Isthmus - Follicular phase - Ipsilateral - Native - Ampulla - Luteal phase - Ipsilateral - Decellular
## Isthmus - Luteal phase - Contralateral - Decellularized - Ampulla - Luteal phase - Ipsilateral - Dece
## Isthmus - Luteal phase - Contralateral - Native - Ampulla - Luteal phase - Ipsilateral - Decellulariz
## Isthmus - Luteal phase - Ipsilateral - Decellularized - Ampulla - Luteal phase - Ipsilateral - Decell
## Isthmus - Luteal phase - Ipsilateral - Native - Ampulla - Luteal phase - Ipsilateral - Decellularized
```

```
## Isthmus - Follicular phase - Contralateral - Decellularized - Ampulla - Luteal phase - Ipsilateral -
## Isthmus - Follicular phase - Contralateral - Native - Ampulla - Luteal phase - Ipsilateral - Native =
## Isthmus - Follicular phase - Ipsilateral - Decellularized - Ampulla - Luteal phase - Ipsilateral - Na
## Isthmus - Follicular phase - Ipsilateral - Native - Ampulla - Luteal phase - Ipsilateral - Native ==
## Isthmus - Luteal phase - Contralateral - Decellularized - Ampulla - Luteal phase - Ipsilateral - Nati
## Isthmus - Luteal phase - Contralateral - Native - Ampulla - Luteal phase - Ipsilateral - Native == 0
## Isthmus - Luteal phase - Ipsilateral - Decellularized - Ampulla - Luteal phase - Ipsilateral - Native
## Isthmus - Luteal phase - Ipsilateral - Native - Ampulla - Luteal phase - Ipsilateral - Native == 0
## Isthmus - Follicular phase - Contralateral - Native - Isthmus - Follicular phase - Contralateral - De
## Isthmus - Follicular phase - Ipsilateral - Decellularized - Isthmus - Follicular phase - Contralatera
## Isthmus - Follicular phase - Ipsilateral - Native - Isthmus - Follicular phase - Contralateral - Dec
## Isthmus - Luteal phase - Contralateral - Decellularized - Isthmus - Follicular phase - Contralateral
## Isthmus - Luteal phase - Contralateral - Native - Isthmus - Follicular phase - Contralateral - Decell
## Isthmus - Luteal phase - Ipsilateral - Decellularized - Isthmus - Follicular phase - Contralateral -
## Isthmus - Luteal phase - Ipsilateral - Native - Isthmus - Follicular phase - Contralateral - Decellul
## Isthmus - Follicular phase - Ipsilateral - Decellularized - Isthmus - Follicular phase - Contralatera
## Isthmus - Follicular phase - Ipsilateral - Native - Isthmus - Follicular phase - Contralateral - Nati
## Isthmus - Luteal phase - Contralateral - Decellularized - Isthmus - Follicular phase - Contralateral
## Isthmus - Luteal phase - Contralateral - Native - Isthmus - Follicular phase - Contralateral - Native
## Isthmus - Luteal phase - Ipsilateral - Decellularized - Isthmus - Follicular phase - Contralateral -
## Isthmus - Luteal phase - Ipsilateral - Native - Isthmus - Follicular phase - Contralateral - Native =
## Isthmus - Follicular phase - Ipsilateral - Native - Isthmus - Follicular phase - Ipsilateral - Decell
## Isthmus - Luteal phase - Contralateral - Decellularized - Isthmus - Follicular phase - Ipsilateral -
## Isthmus - Luteal phase - Contralateral - Native - Isthmus - Follicular phase - Ipsilateral - Decellul
## Isthmus - Luteal phase - Ipsilateral - Decellularized - Isthmus - Follicular phase - Ipsilateral - De
## Isthmus - Luteal phase - Ipsilateral - Native - Isthmus - Follicular phase - Ipsilateral - Decellula
## Isthmus - Luteal phase - Contralateral - Decellularized - Isthmus - Follicular phase - Ipsilateral -
## Isthmus - Luteal phase - Contralateral - Native - Isthmus - Follicular phase - Ipsilateral - Native =
## Isthmus - Luteal phase - Ipsilateral - Decellularized - Isthmus - Follicular phase - Ipsilateral - Na
## Isthmus - Luteal phase - Ipsilateral - Native - Isthmus - Follicular phase - Ipsilateral - Native ==
## Isthmus - Luteal phase - Contralateral - Native - Isthmus - Luteal phase - Contralateral - Decellula
## Isthmus - Luteal phase - Ipsilateral - Decellularized - Isthmus - Luteal phase - Contralateral - Dec
## Isthmus - Luteal phase - Ipsilateral - Native - Isthmus - Luteal phase - Contralateral - Decellulari
## Isthmus - Luteal phase - Ipsilateral - Decellularized - Isthmus - Luteal phase - Contralateral - Nati
## Isthmus - Luteal phase - Ipsilateral - Native - Isthmus - Luteal phase - Contralateral - Native == 0
## Isthmus - Luteal phase - Ipsilateral - Native - Isthmus - Luteal phase - Ipsilateral - Decellularize
## (Adjusted p values reported -- single-step method)
```

```
tidied_results <- tidy(comp)
```

```
## Warning in RET$pfunction("adjusted", ...): Completion with error > abseps

## Warning in RET$pfunction("adjusted", ...): Completion with error > abseps

## Warning in RET$pfunction("adjusted", ...): Completion with error > abseps

## Warning in RET$pfunction("adjusted", ...): Completion with error > abseps

## Warning in RET$pfunction("adjusted", ...): Completion with error > abseps

## Warning in RET$pfunction("adjusted", ...): Completion with error > abseps

## Warning in RET$pfunction("adjusted", ...): Completion with error > abseps
```

```r
write.csv(tidied_results, file = "Tukey_results.csv", row.names = FALSE)


################## YM-Porosity correlation ######################################


# Read the dataset from a CSV file
df <- read.csv("Oviduct_YM.csv")

# Create a new column to define the X_position group
df$X_position_group <- ifelse(df$X_position <= 400, "Lumen-Stroma surface", "Muscle-Tunica surface")

# Update the labels for Position, Estrus_cycle, and Tissue
df$Position <- factor(df$Position, labels = c("Contralateral", "Ipsilateral"))
df$Estrus_cycle <- factor(df$Estrus_cycle, labels = c("Follicular phase", "Luteal phase"))
df$Tissue <- factor(df$Tissue, labels = c("Ampulla", "Isthmus"))

png(filename="Oviduct_correlation_Ampulla.png",
    width = 10, height = 15, units = "in",
    res = 600)

par(mgp=c(1.5,0.5,0),
    mar = c(4, #bottom
            4, #left
            3, #top
            2),
    cex.main = 0.8,
    family = "sans" #right
)


# Ampulla

# Subset the data for Ampulla
df_ampulla <- subset(df, Tissue == "Ampulla")

# Create scatter plot for Contra-lateral and Decellularized in the Follicular phase
p_contra_decell_follicular <- ggplot(df_ampulla, aes(x = Porosity, y = YM, color = Cellular)) +
  geom_point(data = subset(df_ampulla, Position == "Contralateral" & Estrus_cycle == "Follicular phase"
  geom_smooth(data = subset(df_ampulla, Position == "Contralateral" & Estrus_cycle == "Follicular phase
  labs(x = "Porosity (%)", y = "Young's Modulus (Pa)") +
  ggtitle("Contralateral - Decellularized (Follicular phase)") +
  stat_cor(data = subset(df_ampulla, Position == "Contralateral" & Estrus_cycle == "Follicular phase" &
  theme_bw() +
  theme(panel.grid.major = element_blank(),
        panel.grid.minor = element_blank(),
        legend.position = "none",
        axis.title.x = element_text(size = 16),
        axis.title.y = element_text(size = 16))

# Create scatter plot for Contra-lateral and Native in the Follicular phase
p_contra_native_follicular <- ggplot(df_ampulla, aes(x = Porosity, y = YM, color = Cellular)) +
  geom_point(data = subset(df_ampulla, Position == "Contralateral" & Estrus_cycle == "Follicular phase"
```

```
  geom_smooth(data = subset(df_ampulla, Position == "Contralateral" & Estrus_cycle == "Follicular phase
  labs(x = "Porosity (%)", y = "Young's Modulus (Pa)") +
  ggtitle("Contralateral - Native (Follicular phase)") +
  stat_cor(data = subset(df_ampulla, Position == "Contralateral" & Estrus_cycle == "Follicular phase" &
  theme_bw() +
  theme(panel.grid.major = element_blank(),
        panel.grid.minor = element_blank(),
        legend.position = "none",
        axis.title.x = element_text(size = 16),
        axis.title.y = element_text(size = 16))

# Create scatter plot for Ipsi-lateral and Decellularized in the Follicular phase
p_ipsi_decell_follicular <- ggplot(df_ampulla, aes(x = Porosity, y = YM, color = Cellular)) +
  geom_point(data = subset(df_ampulla, Position == "Ipsilateral" & Estrus_cycle == "Follicular phase" &
  geom_smooth(data = subset(df_ampulla, Position == "Ipsilateral" & Estrus_cycle == "Follicular phase" &
  labs(x = "Porosity (%)", y = "Young's Modulus (Pa)") +
  ggtitle("Ipsilateral - Decellularized (Follicular phase)") +
  stat_cor(data = subset(df_ampulla, Position == "Ipsilateral" & Estrus_cycle == "Follicular phase" & Ce
  theme_bw() +
  theme(panel.grid.major = element_blank(),
        panel.grid.minor = element_blank(),
        legend.position = "none",
        axis.title.x = element_text(size = 16),
        axis.title.y = element_text(size = 16))

# Create scatter plot for Ipsi-lateral and Native in the Follicular phase
p_ipsi_native_follicular <- ggplot(df_ampulla, aes(x = Porosity, y = YM, color = Cellular)) +
  geom_point(data = subset(df_ampulla, Position == "Ipsilateral" & Estrus_cycle == "Follicular phase" &
  geom_smooth(data = subset(df_ampulla, Position == "Ipsilateral" & Estrus_cycle == "Follicular phase" &
  labs(x = "Porosity (%)", y = "Young's Modulus (Pa)") +
  ggtitle("Ipsilateral - Native (Follicular phase)") +
  stat_cor(data = subset(df_ampulla, Position == "Ipsilateral" & Estrus_cycle == "Follicular phase" & Ce
  theme_bw() +
  theme(panel.grid.major = element_blank(),
        panel.grid.minor = element_blank(),
        legend.position = "none",
        axis.title.x = element_text(size = 16),
        axis.title.y = element_text(size = 16))

# Create scatter plot for Contra-lateral and Decellularized in the Luteal phase
p_contra_decell_luteal <- ggplot(df_ampulla, aes(x = Porosity, y = YM, color = Cellular)) +
  geom_point(data = subset(df_ampulla, Position == "Contralateral" & Estrus_cycle == "Luteal phase" & Ce
  geom_smooth(data = subset(df_ampulla, Position == "Contralateral" & Estrus_cycle == "Luteal phase" & C
  labs(x = "Porosity (%)", y = "Young's Modulus (Pa)") +
  ggtitle("Contralateral - Decellularized (Luteal phase)") +
  stat_cor(data = subset(df_ampulla, Position == "Contralateral" & Estrus_cycle == "Luteal phase" & Cell
  theme_bw() +
  theme(panel.grid.major = element_blank(),
        panel.grid.minor = element_blank(),
        legend.position = "none",
        axis.title.x = element_text(size = 16),
        axis.title.y = element_text(size = 16))
```

```r
# Create scatter plot for Contra-lateral and Native in the Luteal phase
p_contra_native_luteal <- ggplot(df_ampulla, aes(x = Porosity, y = YM, color = Cellular)) +
  geom_point(data = subset(df_ampulla, Position == "Contralateral" & Estrus_cycle == "Luteal phase" & Ce
  geom_smooth(data = subset(df_ampulla, Position == "Contralateral" & Estrus_cycle == "Luteal phase" & C
  labs(x = "Porosity (%)", y = "Young's Modulus (Pa)") +
  ggtitle("Contralateral - Native (Luteal phase)") +
  stat_cor(data = subset(df_ampulla, Position == "Contralateral" & Estrus_cycle == "Luteal phase" & Cel
  theme_bw() +
  theme(panel.grid.major = element_blank(),
        panel.grid.minor = element_blank(),
        legend.position = "none",
        axis.title.x = element_text(size = 16),
        axis.title.y = element_text(size = 16))

# Create scatter plot for Ipsi-lateral and Decellularized in the Luteal phase
p_ipsi_decell_luteal <- ggplot(df_ampulla, aes(x = Porosity, y = YM, color = Cellular)) +
  geom_point(data = subset(df_ampulla, Position == "Ipsilateral" & Estrus_cycle == "Luteal phase" & Cell
  geom_smooth(data = subset(df_ampulla, Position == "Ipsilateral" & Estrus_cycle == "Luteal phase" & Cel
  labs(x = "Porosity (%)", y = "Young's Modulus (Pa)") +
  ggtitle("Ipsilateral - Decellularized (Luteal phase)") +
  stat_cor(data = subset(df_ampulla, Position == "Ipsilateral" & Estrus_cycle == "Luteal phase" & Cellul
  theme_bw() +
  theme(panel.grid.major = element_blank(),
        panel.grid.minor = element_blank(),
        legend.position = "none",
        axis.title.x = element_text(size = 16),
        axis.title.y = element_text(size = 16))

# Create scatter plot for Ipsi-lateral and Native in the Luteal phase
p_ipsi_native_luteal <- ggplot(df_ampulla, aes(x = Porosity, y = YM, color = Cellular)) +
  geom_point(data = subset(df_ampulla, Position == "Ipsilateral" & Estrus_cycle == "Luteal phase" & Cell
  geom_smooth(data = subset(df_ampulla, Position == "Ipsilateral" & Estrus_cycle == "Luteal phase" & Cel
  labs(x = "Porosity (%)", y = "Young's Modulus (Pa)") +
  ggtitle("Ipsilateral - Native (Luteal phase)") +
  stat_cor(data = subset(df_ampulla, Position == "Ipsilateral" & Estrus_cycle == "Luteal phase" & Cellul
  theme_bw() +
  theme(panel.grid.major = element_blank(),
        panel.grid.minor = element_blank(),
        legend.position = "none",
        axis.title.x = element_text(size = 16),
        axis.title.y = element_text(size = 16))

# Combine the plots using cowplot
p_combined <- cowplot::plot_grid(
  cowplot::plot_grid(p_contra_decell_follicular, p_contra_native_follicular, nrow = 1),
  cowplot::plot_grid(p_ipsi_decell_follicular, p_ipsi_native_follicular, nrow = 1),
  cowplot::plot_grid(p_contra_decell_luteal, p_contra_native_luteal, nrow = 1),
  cowplot::plot_grid(p_ipsi_decell_luteal, p_ipsi_native_luteal, nrow = 1),
  nrow = 4
)
```

```
## `geom_smooth()` using formula = 'y ~ x'
## `geom_smooth()` using formula = 'y ~ x'
```

```
## Warning: Removed 6 rows containing non-finite values (`stat_smooth()`).

## Warning: Removed 6 rows containing non-finite values (`stat_cor()`).

## Warning: Removed 6 rows containing missing values (`geom_point()`).

## `geom_smooth()` using formula = 'y ~ x'
## `geom_smooth()` using formula = 'y ~ x'
## `geom_smooth()` using formula = 'y ~ x'
## `geom_smooth()` using formula = 'y ~ x'
## `geom_smooth()` using formula = 'y ~ x'
## `geom_smooth()` using formula = 'y ~ x'
```

```
# Display the combined plot
p_combined

dev.off()
```

```
## pdf
##   2
```

```
#Isthmus

png(filename="Oviduct_correlation_isthmus.png",
    width = 10, height = 15, units = "in",
    res = 600)

par(mgp=c(1.5,0.5,0),
    mar = c(4, #bottom
            4, #left
            3, #top
            2),
    cex.main = 0.8,
    family = "sans" #right
)

# Subset the data for Isthmus
df_isthmus <- subset(df, Tissue == "Isthmus")


# Create scatter plot for Contra-lateral and Decellularized in the Follicular phase
p_contra_decell_follicular <- ggplot(df_isthmus, aes(x = Porosity, y = YM, color = Cellular)) +
  geom_point(data = subset(df_isthmus, Position == "Contralateral" & Estrus_cycle == "Follicular phase"
  geom_smooth(data = subset(df_isthmus, Position == "Contralateral" & Estrus_cycle == "Follicular phase
  labs(x = "Porosity (%)", y = "Young's Modulus (Pa)") +
  ggtitle("Contralateral - Decellularized (Follicular phase)") +
  stat_cor(data = subset(df_isthmus, Position == "Contralateral" & Estrus_cycle == "Follicular phase" &
  theme_bw() +
  theme(panel.grid.major = element_blank(),
        panel.grid.minor = element_blank(),
        legend.position = "none",
        axis.title.x = element_text(size = 16),
        axis.title.y = element_text(size = 16))

# Create scatter plot for Contra-lateral and Native in the Follicular phase
p_contra_native_follicular <- ggplot(df_isthmus, aes(x = Porosity, y = YM, color = Cellular)) +
  geom_point(data = subset(df_isthmus, Position == "Contralateral" & Estrus_cycle == "Follicular phase"
```

```
  geom_smooth(data = subset(df_isthmus, Position == "Contralateral" & Estrus_cycle == "Follicular phase
  labs(x = "Porosity (%)", y = "Young's Modulus (Pa)") +
  ggtitle("Contralateral - Native (Follicular phase)") +
  stat_cor(data = subset(df_isthmus, Position == "Contralateral" & Estrus_cycle == "Follicular phase" &
  theme_bw() +
  theme(panel.grid.major = element_blank(),
        panel.grid.minor = element_blank(),
        legend.position = "none",
        axis.title.x = element_text(size = 16),
        axis.title.y = element_text(size = 16))

# Create scatter plot for Ipsi-lateral and Decellularized in the Follicular phase
p_ipsi_decell_follicular <- ggplot(df_isthmus, aes(x = Porosity, y = YM, color = Cellular)) +
  geom_point(data = subset(df_isthmus, Position == "Ipsilateral" & Estrus_cycle == "Follicular phase" &
  geom_smooth(data = subset(df_isthmus, Position == "Ipsilateral" & Estrus_cycle == "Follicular phase" &
  labs(x = "Porosity (%)", y = "Young's Modulus (Pa)") +
  ggtitle("Ipsilateral - Decellularized (Follicular phase)") +
  stat_cor(data = subset(df_isthmus, Position == "Ipsilateral" & Estrus_cycle == "Follicular phase" & Ce
  theme_bw() +
  theme(panel.grid.major = element_blank(),
        panel.grid.minor = element_blank(),
        legend.position = "none",
        axis.title.x = element_text(size = 16),
        axis.title.y = element_text(size = 16))

# Create scatter plot for Ipsi-lateral and Native in the Follicular phase
p_ipsi_native_follicular <- ggplot(df_isthmus, aes(x = Porosity, y = YM, color = Cellular)) +
  geom_point(data = subset(df_isthmus, Position == "Ipsilateral" & Estrus_cycle == "Follicular phase" &
  geom_smooth(data = subset(df_isthmus, Position == "Ipsilateral" & Estrus_cycle == "Follicular phase" &
  labs(x = "Porosity (%)", y = "Young's Modulus (Pa)") +
  ggtitle("Ipsilateral - Native (Follicular phase)") +
  stat_cor(data = subset(df_isthmus, Position == "Ipsilateral" & Estrus_cycle == "Follicular phase" & Ce
  theme_bw() +
  theme(panel.grid.major = element_blank(),
        panel.grid.minor = element_blank(),
        legend.position = "none",
        axis.title.x = element_text(size = 16),
        axis.title.y = element_text(size = 16))

# Create scatter plot for Contra-lateral and Decellularized in the Luteal phase
p_contra_decell_luteal <- ggplot(df_isthmus, aes(x = Porosity, y = YM, color = Cellular)) +
  geom_point(data = subset(df_isthmus, Position == "Contralateral" & Estrus_cycle == "Luteal phase" & Ce
  geom_smooth(data = subset(df_isthmus, Position == "Contralateral" & Estrus_cycle == "Luteal phase" & C
  labs(x = "Porosity (%)", y = "Young's Modulus (Pa)") +
  ggtitle("Contralateral - Decellularized (Luteal phase)") +
  stat_cor(data = subset(df_isthmus, Position == "Contralateral" & Estrus_cycle == "Luteal phase" & Cel
  theme_bw() +
  theme(panel.grid.major = element_blank(),
        panel.grid.minor = element_blank(),
        legend.position = "none",
        axis.title.x = element_text(size = 16),
        axis.title.y = element_text(size = 16))
```

```r
# Create scatter plot for Contra-lateral and Native in the Luteal phase
p_contra_native_luteal <- ggplot(df_isthmus, aes(x = Porosity, y = YM, color = Cellular)) +
  geom_point(data = subset(df_isthmus, Position == "Contralateral" & Estrus_cycle == "Luteal phase" & Ce
  geom_smooth(data = subset(df_isthmus, Position == "Contralateral" & Estrus_cycle == "Luteal phase" & C
  labs(x = "Porosity (%)", y = "Young's Modulus (Pa)") +
  ggtitle("Contralateral - Native (Luteal phase)") +
  stat_cor(data = subset(df_isthmus, Position == "Contralateral" & Estrus_cycle == "Luteal phase" & Cel
  theme_bw() +
  theme(panel.grid.major = element_blank(),
        panel.grid.minor = element_blank(),
        legend.position = "none",
        axis.title.x = element_text(size = 16),
        axis.title.y = element_text(size = 16))

# Create scatter plot for Ipsi-lateral and Decellularized in the Luteal phase
p_ipsi_decell_luteal <- ggplot(df_isthmus, aes(x = Porosity, y = YM, color = Cellular)) +
  geom_point(data = subset(df_isthmus, Position == "Ipsilateral" & Estrus_cycle == "Luteal phase" & Cel
  geom_smooth(data = subset(df_isthmus, Position == "Ipsilateral" & Estrus_cycle == "Luteal phase" & Ce
  labs(x = "Porosity (%)", y = "Young's Modulus (Pa)") +
  ggtitle("Ipsilateral - Decellularized (Luteal phase)") +
  stat_cor(data = subset(df_isthmus, Position == "Ipsilateral" & Estrus_cycle == "Luteal phase" & Cellu
  theme_bw() +
  theme(panel.grid.major = element_blank(),
        panel.grid.minor = element_blank(),
        legend.position = "none",
        axis.title.x = element_text(size = 16),
        axis.title.y = element_text(size = 16))

# Create scatter plot for Ipsi-lateral and Native in the Luteal phase
p_ipsi_native_luteal <- ggplot(df_isthmus, aes(x = Porosity, y = YM, color = Cellular)) +
  geom_point(data = subset(df_isthmus, Position == "Ipsilateral" & Estrus_cycle == "Luteal phase" & Cel
  geom_smooth(data = subset(df_isthmus, Position == "Ipsilateral" & Estrus_cycle == "Luteal phase" & Ce
  labs(x = "Porosity (%)", y = "Young's Modulus (Pa)") +
  ggtitle("Ipsilateral - Native (Luteal phase)") +
  stat_cor(data = subset(df_isthmus, Position == "Ipsilateral" & Estrus_cycle == "Luteal phase" & Cellu
  theme_bw() +
  theme(panel.grid.major = element_blank(),
        panel.grid.minor = element_blank(),
        legend.position = "none",
        axis.title.x = element_text(size = 16),
        axis.title.y = element_text(size = 16))

# Combine the plots using cowplot
p_combined <- cowplot::plot_grid(
  cowplot::plot_grid(p_contra_decell_follicular, p_contra_native_follicular, nrow = 1),
  cowplot::plot_grid(p_ipsi_decell_follicular, p_ipsi_native_follicular, nrow = 1),
  cowplot::plot_grid(p_contra_decell_luteal, p_contra_native_luteal, nrow = 1),
  cowplot::plot_grid(p_ipsi_decell_luteal, p_ipsi_native_luteal, nrow = 1),
  nrow = 4
)
```

```
## `geom_smooth()` using formula = 'y ~ x'
## `geom_smooth()` using formula = 'y ~ x'
## `geom_smooth()` using formula = 'y ~ x'
```

```
## `geom_smooth()` using formula = 'y ~ x'
## `geom_smooth()` using formula = 'y ~ x'

## Warning: Removed 10 rows containing non-finite values (`stat_smooth()`).

## Warning: Removed 10 rows containing non-finite values (`stat_cor()`).

## Warning: Removed 10 rows containing missing values (`geom_point()`).

## `geom_smooth()` using formula = 'y ~ x'
## `geom_smooth()` using formula = 'y ~ x'
## `geom_smooth()` using formula = 'y ~ x'
```

```r
# Display the combined plot
p_combined


dev.off()
```

```
## pdf
##   2
```
